# Comprehensive genomic analysis of *Trypanosoma rangeli* reveals key insights into the biology and evolution of this non-virulent American mammalian trypanosome

**DOI:** 10.64898/2026.02.11.700901

**Authors:** Guilherme Augusto Maia, Gabriel Machado Matos, Adriana Corrêa da Silva, Caibe Alves Pereira, Carime Lessa Mansur Pontes, Debora Denardin Lückmeyer, Rafaela Andrade do Carmo, Renato Simões Moreira, Thaynara Karoliny de Souza Pereira, Amábilli de Souza Rosar, Theo Cardozo Brascher, Guilherme Razzera, José Franco da Silveira Filho, Mário Steindel, Patrícia Hermes Stoco, Glauber Wagner, Björn Andersson, Edmundo Carlos Grisard

## Abstract

**Background:** *Trypanosoma rangeli* is a non-virulent hemoflagellate protozoan parasite that infects mammals, including humans, in Central and South America. It is primarily transmitted through the bites of triatomine bugs and shares an overlapping geographical distribution with *T. cruzi*, as well as triatomine vectors and mammalian hosts, and various shared surface antigens. The life cycle of *T. rangeli* differs from those of other human-infecting trypanosomes, such as *T. cruzi* and *T. brucei*, and the molecular mechanisms underlying host-parasite and host-vector interactions are not well understood, demanding improved molecular and genomic resources.

**Results:** The use of a hybrid approach to sequence and assemble the *T. rangeli* genome, complemented by transcriptomics and proteomics for functional gene annotation, led to the generation of the near-complete genome sequence of the parasite. Detailed intra- and inter-specific comparative genomics allowed analysis of polymorphisms, genome structure and improved resolution of genes coding for important surface molecules such as Mucins, TASV and GP63.

**Conclusions:** The improved *T. rangeli* genome assembly, combined with comparative genomics has yielded novel biological insights. These included the first description of a metalloprotease activity, attributed to specific GP63 genes that are absent in *Leishmania* species. In addition, a TASV gene family that is absent in *T. cruzi* was identified, which indicates a possible role in the *T. rangeli* infection process.

## 1. BACKGROUND

Trypanosomatids are a group of single-celled, mono-flagellated organisms belonging to the class Kinetoplastea. This group includes etiological agents of neglected tropical diseases of humans such as *Trypanosoma cruzi* (Chagas disease or American trypanosomiasis), *Trypanosoma brucei* (African sleeping sickness), and *Leishmania* spp. (Leishmaniasis) [1]. Collectively, these neglected and socially determined tropical diseases collectively affect more than 18 million people worldwide, mostly in the developing countries [2,3].

*Trypanosoma rangeli* [4], is a mammalian parasite species found in Central and South America, primarily transmitted via bite of triatomine vectors. Unlike *T. brucei* and *T. cruzi*, *T. rangeli* is considered non-virulent to mammalian hosts, despite the genetic and antigenic similarity with *T. cruzi* and its inoculative mode of transmission [5,6]. On the other hand, *T. rangeli* is harmful to insect vectors, particularly within the genus *Rhodnius* [7–9].

Reports of natural mixed infections by *T. rangeli* and *T. cruzi* have been documented in a variety of mammalian hosts and triatomine vectors species throughout Central and South America [4,7,10–12]. However, the prevalence of human infection by *T. rangeli* remains unclear. The sympatric occurrence, along with shared hosts, vectors and several antigens, leads to the possibility of a strong serological cross-reactivity [13–15].

*T. rangeli* infection biology is understudied. Previously, comparative analyses of isolated genes coding for surface antigens or related to cell infection or evasion of the immune system, partial genomic [5,16], transcriptomic [17], and proteomic datasets [18] have been produced.

Comparative biological and molecular studies have revealed the presence of trans-sialidase, one of the most important and well-studied virulence factors of *T. cruzi*, and the surface glycoprotein 63 (GP63) metalloprotease (also known as Leishmaniolysin), this major surface protease of *Leishmania* spp. plays a crucial role in cell infection and intracellular survival of both *Leishmania* [19], and *T. cruzi* [20].

The *T. rangeli* genome has been estimated to be around 21-24 million base pairs (Mbp) [5,16] in size, sharing over 90% of the core genome with the pathogenic trypanosomatids. However, these estimates are based on fragmented assemblies containing 3,436 (AM80) and 9,066 (SC58) scaffolds, respectively. While incomplete, these studies revealed that the *T. rangeli* genome is smaller, less repetitive and with shorter telomeric and sub-telomeric regions compared to the *T. cruzi* genome [21–23]. Additionally, it was proposed that the number of *T. rangeli* genes coding for virulence factors or proteins associated with pathogenesis varies between strains from different lineages and differs from those of *T. cruzi* [24,25]. For instance, compared to *T. cruzi*, *T. rangeli* has a similar number of genes coding for GP63, reduced numbers of genes coding for sialidases/trans-sialidases and Mucin-Associated Surface Proteins (MASP), and, interestingly, an increased number of genes coding for Kinetoplastid Membrane Protein 11 (KMP-11) [5,16].

Trypanosomatid genomes contain syntenic regions where genes are organized in directional transcription clusters, separated by strand-switch regions [5,25–28]. Gene families encoding surface proteins are mostly contained within repetitive regions [24,26,29,30]. Recent proposals suggested that regions of the *T. cruzi* genome that are rich in surface protein-coding genes are involved in generating antigenic diversity through gene recombination and conversion events due to the micro-homology of their flanking regions [23]. Additionally, it has been suggested that DNA replication origins in the *T. cruzi* genome are unevenly dispersed across the genome, as indicated by the enrichment of Orc1Cdc6, a component of the pre-replication complex [31].

Among several aspects of trypanosomatid biology and parasitism that remain to be fully understood, the survival and reproduction of *T. rangeli* within mammalian hosts are of particular interest [32]. For this purpose, the understanding of the complete structure and plasticity of the genome is essential for facilitating comparative genomics of virulent and non-virulent trypanosomatids of mammals. We here describe a far more accurate, near-complete genome sequence *of T. rangeli*, as well as a comparative study of other *T. rangeli* strains. Our findings reveal remarkable biological differences between virulent and non-virulent trypanosomatids, pointing out the *T. rangeli* genome structure, intraspecific variability and the repertoire of surface proteins, providing insights into the parasite biology and life cycle.

## 2. RESULTS

### 2.1 T. rangeli genome sequencing and assembly

Illumina sequencing of the *T. rangeli* SC58 strain provided 9,264,332 filtered reads (∼4.8Gb) and PacBio sequencing provided 361,938 filtered reads (∼6.9Gb), respectively, corresponding to approximately 20x and 192x coverage. The resulting assembly is here referred to as version 2 of the *T. rangeli* SC58 genome (“*T. rangeli* SC58 V.2”), and the fragmented assembly previously published by Stoco et al. (2014) as the “SC58 reference genome”. A comparison of general assembly metrics for the *T. rangeli* SC58 V.2 genome and the reference genome assembly is shown in Table 1. The *T. rangeli* SC58 V.2 genome is available in Genbank (Bioproject PRJNA1380247).

**Table 1.** Comparison of general assembly metrics of the *Trypanosoma rangeli* SC58 reference and V.2 genomes and *T. cruzi* (CL Brener Esmeraldo-like and Sylvio X10/1 strains) genomes. Footnote: * = Data retrieved from the TriTrypDB v.65. Mbp = Million base pairs. Kbp = Thousand base pairs. aa = amino acids. Scaffold N50 indicates the scaffold length where 50% of the entire assembly is contained in contigs equal to or larger than this value. Scaffold L50 is defined as the count of the smallest number of contigs whose length sum makes up half of the genome size. “Number of hypothetical CDS” represents CDS automatically annotated as “Hypothetical protein”. “ND”= Not determined.

| Metrics | *T. rangeli* |  | *T. cruzi* |  |
| --- | --- | --- | --- | --- |
|  | SC58 V.2 | SC58 Reference* | CL Brener Esmeraldo-like* | Sylvio X10/1* |
| Genome size (Mbp) | 23.68 | 14.02 | 32.53 | 41.38 |
| Number of scaffolds | 81 | 7,433 | 41 | 47 |
| G+C content (%) | 53.30 | 53.26 | 50.38 | 51.57 |
| Scaffolds > 50 Kbp | 67 | 0 | 41 | 47 |
| Longest scaffold (Mbp) | 1.37 | 0.25 | 2.37 | 3.12 |
| Scaffold N50 | 558,628 | 2,203 | 870,934 | 1,006,492 |
| Scaffold L50 | 14 | 2,096 | 12 | 14 |
| N bases / 100 Kbp | 0 | 21.31 | 20,604.70 | 3,242.42 |
| Number of rRNA | 58 | 16 | 9 | ND |
| Number of tRNA | 73 | 120 | 55 | 65 |
| Total number of CDS | 9,203 | 7,475 | 9,039 | 18,643 |
| Number of hypothetical CDS | 3,067 | 5,075 | 4,260 | 13,760 |
| Mean CDS size (aa) | 505 | 460 | 493 | 178 |
| CDS < 100 aa | 299 | 410 | 207 | 5,573 |

Cross-mapping analysis using the MUMmer4 software indicated that 99.52% of the nuclear genomic content previously described in the SC58 reference genome [5] is included in the *T. rangeli* SC58 V.2 genome. Also, 97.71%, 97.25% and 97.52% of the reads from the PIT10, Choachí and R1625 *T. rangeli* strains, respectively, were mapped to the *T. rangeli* SC58 V.2 genome. Unique copies of all 130 universal single-copy orthologs of the phylum Euglenozoa were identified, distributed across 43 different scaffolds of the *T. rangeli* SC58 V.2 genome.

A high level of synteny was observed among the *T. rangeli* genomes, with few structural variants. However, when comparing the *T. rangeli* SC58 V.2 genome assembly to the assemblies of the *T. cruzi* CL Brener Esmeraldo-like and Sylvio X10/1 strains, significant structural diversity was found (Additional file 1: Supplementary Figure S1).

The V.2 genome contained tRNA genes for all 20 amino acids and the canonical 5.8S, 18S and 28S rRNA genes (Table 1). The V.2 genome contains fewer tRNA genes than the previous reference genome [5], having a significantly higher number of rRNA genes compared to other trypanosomatid genomes (Table 1). Also, a total of 62 Spliced Leader (SL) RNA genes (Mini-Exon) exhibiting a high level of conservancy were identified (Additional file 2: Supplementary Figure S2A) in five distinct scaffolds, ranging from 9 to 16 copies/scaffold (Additional file 2: Supplementary Figure S2B).

The search for chromosome-specific probe sequences of *T. cruzi* within the scaffolds of the *T. rangeli* SC58 V.2 genome revealed unique mapping for nine out of 11 *T. cruzi* probes (Additional file 3: Supplementary Table S1). When comparing the TcCLB.510129.20 chromosomal probe, derived from a 1.38 Kb-long surface membrane protein identified as a single-copy gene in the *T. cruzi* genome, positive BLAST results were obtained with sequences annotated as “surface membrane proteins” on scaffolds 5 and 14 of the *T. rangeli* SC58 V.2 genome. These two scaffolds are of 0.73 and 0.45 Mbp, respectively, and exhibited 85.38% similarity to each other. Fourteen segments of the *T. cruzi* TcCLB.510129.20 sequence matched these scaffolds, displaying similarities ranging from 78.36 to 93.51% with 100% coverage. However, the regions located upstream and downstream of these matches were distinct, indicating that this gene is present in multiple copies within the *T. rangeli* genome.

Probe TcCLB.511041.40 targeting a gene coding for a 1.63 Kb-long hexose transporter gene in *T. cruzi* and showed sequence similarity with several transporter genes distributed in 10 scaffolds in the *T. rangeli* SC58 V.2 genome. This finding supports the observations from Cabrine-Santos et al. (2009) regarding the presence of duplicated genes encoding hexose transporters on different *T. rangeli* chromosomes. These chromosomes range in size from approximately 500 to 650 Kb, indicating the presence of paralogous genes that are specifically unique to *T. rangeli*.

Ten out of the 81 scaffolds of *T. rangeli* SC58 V.2 genome contain canonical telomeric repeats close to their extremities, possibly corresponding to chromosomal ends (Additional file 4: Supplementary Table S2). In accordance with the genome topology proposed by Stoco et al. (2014), subtelomeric regions adjacent to the telomeric repeats were found to contain genes from multicopy families such as (Trans-)Sialidase and GP63 (Fig. 2).

**Figure 1:**
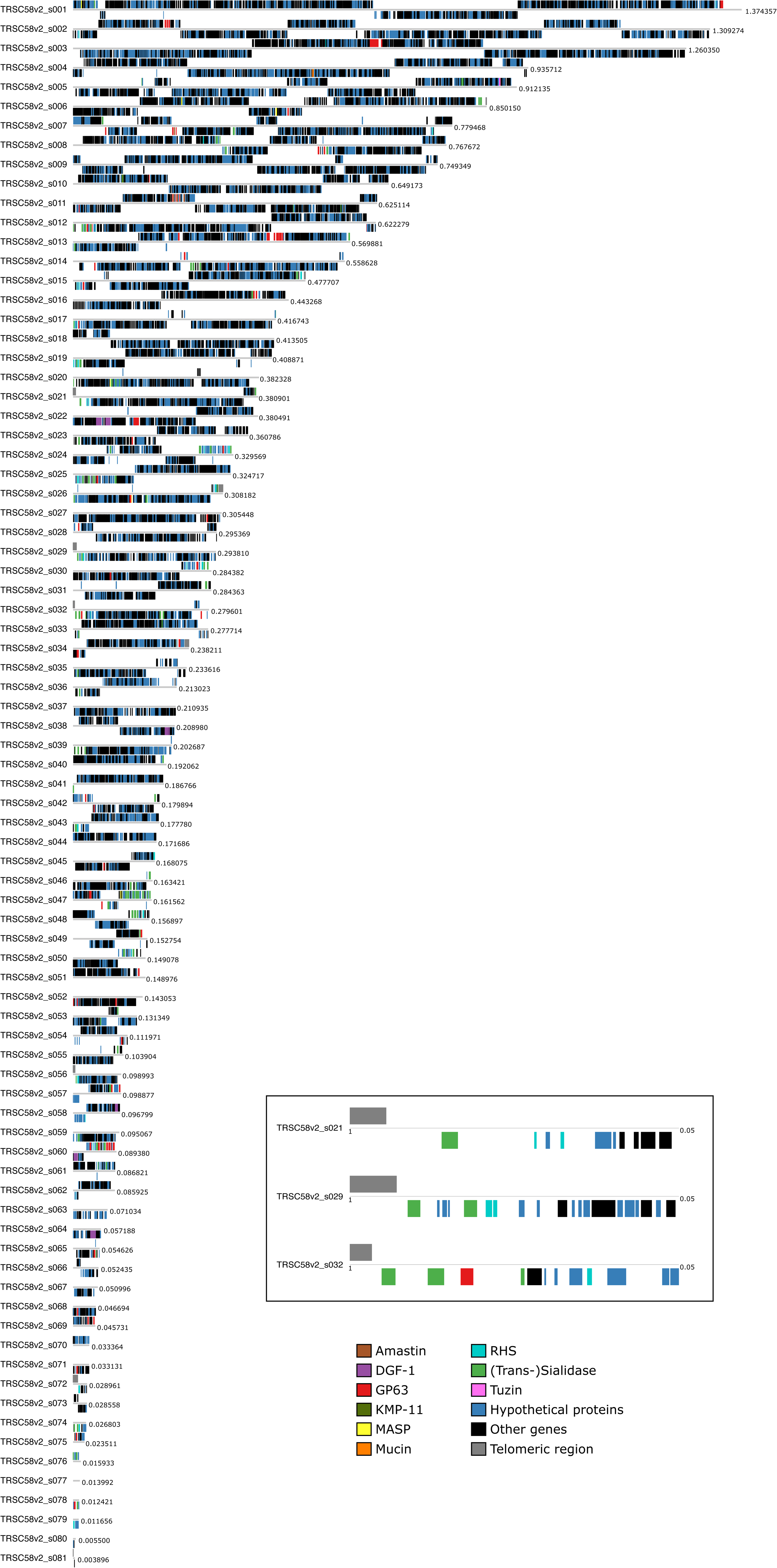
Distribution of multicopy gene families in the scaffolds of the *Trypanosoma rangeli* SC58 V.2 genome. Scaffolds were drawn in proportion to their size, indicated in million base pairs (Mbp). Core genes are represented by black boxes, and the hypothetical and multicopy gene families are represented by colored boxes as indicated. The position of the genes above or below the line corresponds to the direction of transcription, and scaffolds containing canonical telomeric repetitions are shown in detail within the black square. The highlighted region represents an enlarged view (0.05 Mb window) of the telomeric regions of scaffolds 22, 29, and 32.

**Figure 2.**
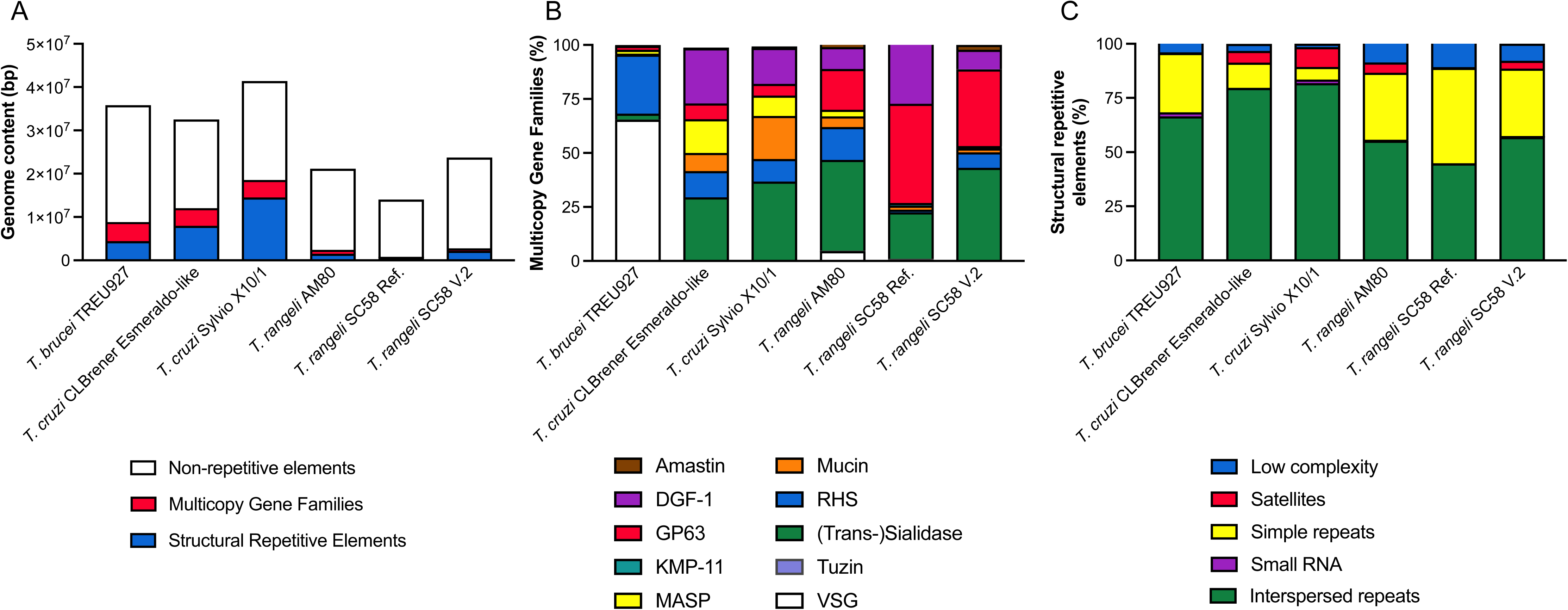
Comparison of repetitive genomic content in studied trypanosomatid species, discerning structural repetitions from multicopy gene families. Footnote: bp = base pairs. (A) Representation of repetitive genomic content in studied trypanosomatid species, in total number of base pairs. (B) Detailed composition of multicopy gene families in the genome of studied trypanosomatid species, relative to the respective genome size of the species. (C) Detailed composition of structural repetitive elements in the genome of studied trypanosomatid species, relative to the respective genome size of the species. “Ref.” points to the reference genome of *T. rangeli* (SC58 strain), obtained from TriTrypDB v65. “V.2” indicates the *T. rangeli* SC58 V.2 genome.

From the 9,454 CDSs identified in the *T. rangeli* SC58 V.2 genome, approximately 32.93% encode for hypothetical proteins. This proportion is comparable to that of *T. brucei* TREU927 (34.37%), but notably lower than for *T. cruzi*, for example, the Esmeraldo-like haplotype (47.13%) and the Sylvio X10/1 strain (73.81%). The use of a hybrid assembly and meticulous annotation allowed a reduction of hypothetical proteins compared to previous *T. rangeli* datasets, such as AM80 (56.72%) and the SC58 reference genome (67.89%). The annotation of the *T. rangeli* SC58 V.2 genome was supported by transcriptomic [17] and proteomic [18] data generated by our group, demonstrating evidence of transcription and expression for 96 (1.02%) of the hypothetical CDSs. The distribution of all predicted and annotated CDSs in the *T. rangeli* SC58 V.2 genome, including notable multigenic families and the 110 potential pseudogenes, is illustrated in Fig. 1. Further analysis of genes associated with the RNA interference machinery (RNAi) on the *T. rangeli* SC58 V.2 genome confirmed the non-functionality of this pathway, as described previously [5].

### 2.2 Repetitive genome content

Excepting telomeric repeats, 11.60% of the *T. rangeli* SC58 V.2 genome consists of repetitive sequences, corresponding to approximately 2.8 Mbp, of which 9.22% (∼2.2 Mbp) comprises structural repetitive elements, while roughly ∼2.38% (565,000 bp) includes members of multicopy gene families. Compared to the TriTryp genomes (*T. brucei*, *T. cruzi* and *L. major*), *T. rangeli* has the least repetitive genome when compared to *T. cruzi* CL Brener (37.03%) and Sylvio X10/1 (44.86%) and to *T. brucei* TREU927 strain (24.73%). This difference is primarily attributed to a lower number of genes from multicopy gene families, fewer structural repetitive elements and shorter telomeric sequences (Fig. 2a).

Comparative analysis of the diversity of multicopy gene families in trypanosomatid genomes showed that the *T. brucei* VSG family is the largest, comprising over 65% of the total number of nucleotides among studied multicopy genes, followed by the RHS family that is also present in *T. cruzi* (Fig. 2b). VSG genes have previously been described in the *T. rangeli* AM80 genome, but these genes were not found in the *T. rangeli* SC58 reference genome or in the V.2 genome (Fig. 2b).

Among these gene families, the (Trans-)Sialidases are highly represented across the studied genomes, varying from 29.37% and 36.66% of repeated genes in the *T. cruzi* CL Brener and Sylvio X10/1 strains, to 41.96% and 42.74% in the *T. rangeli* AM80 and SC58 V.2 genomes, respectively (Fig. 2b). While *T. cruzi* can transfer sialic acid from the host cell to its cellular membrane through a sialyltransferase (trans-sialidase) activity, *T. rangeli* sialidases differ regarding the hydrolytic substrates [33] and lacks the transferase activity [34,35]. We therefore referred to *T. rangeli* sialidases as “(Trans-)Sialidase”.

GP63 is the second largest multicopy gene family in the *T. rangeli* SC58 V.2 genome (35.47%) (Fig. 2b). Uniquely for the American trypanosomatids, *T. cruzi* and *T. rangeli,* the DGF-1 multicopy gene family constitutes a considerable percentage of the total repetitive content (Fig. 2b).

Interspersed repeats substantially contributed to the repetitive content in the *T. rangeli* SC58 V.2 genome (65.10%). In total, this is lower than in *T. cruzi* Sylvio X10/1 (81.83%), but considerably higher when compared to simple repeats in the SC58 V.2 genome (21.42%) (Fig. 2c). These interspersed repeats are composed of sequences that have been found to be present in all trypanosomatid genomes included in this study. Among these sequences, retroelements constitute 15.89% in the *T. rangeli* SC58 V.2 genome, which can be compared to 44.39% in the *T. brucei* TREU927 genome (Fig. 2c).

### 2.3 Analysis of orthology

Orthology analysis using 42,969 CDS from *L. amazonensis*, *L. infantum*, *T. brucei*, *T. cruzi*, and the *T. rangeli* SC58 V.2 and reference genomes resulted in 9,111 CDS clusters and demonstrated that 96.56% of all CDSs from these species are orthologous. The orthologous core for these species constituted 4,501 clusters and the distribution of these clusters between species, as well as the species-specific genes (single-CDS clusters), is shown in Fig. 3.

**Figure 3.**
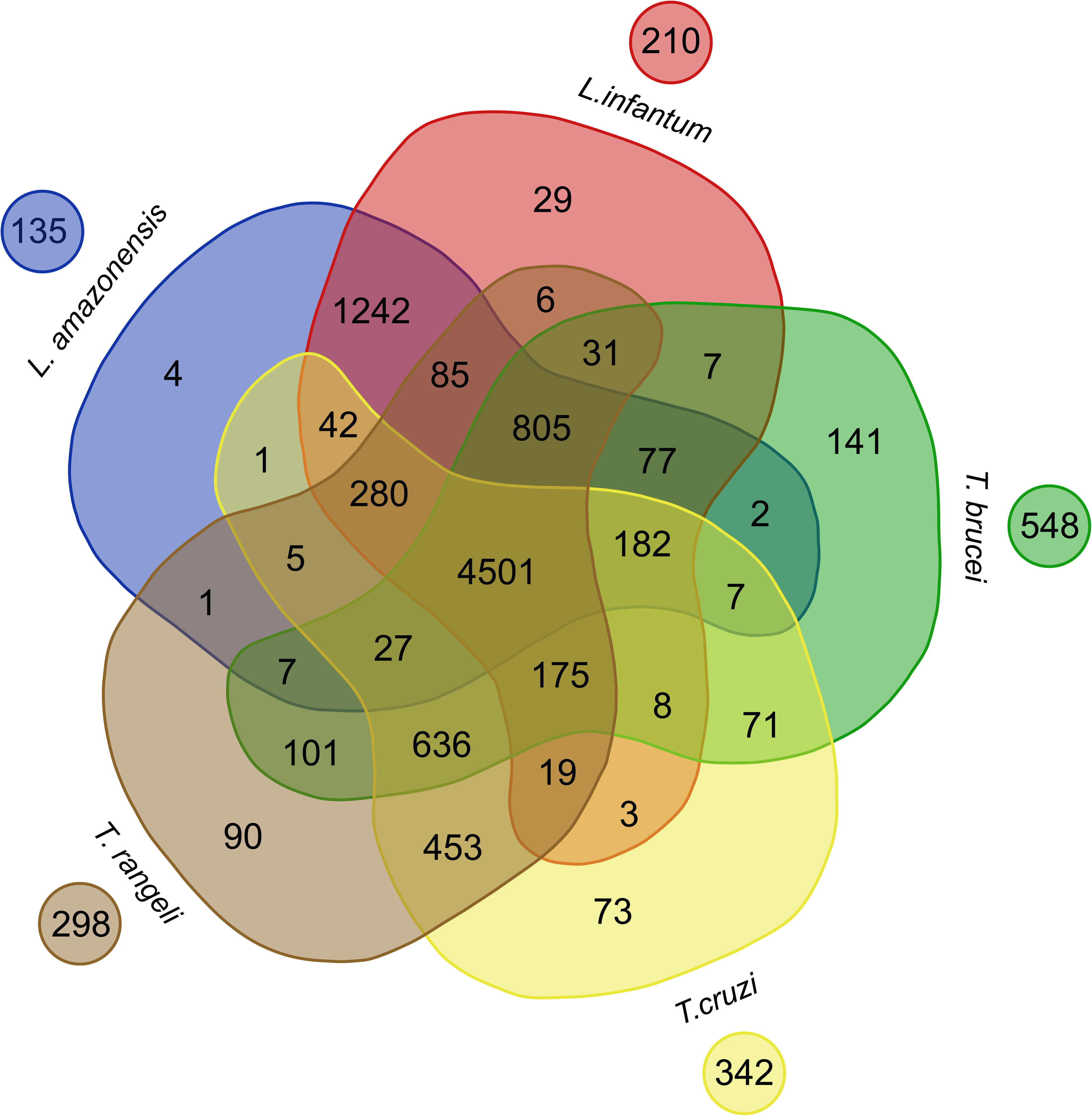
Distribution of clusters formed among the CDS of *Leishmania amazonensis, L. infantum, Trypanosoma brucei, T. cruzi*, the reference and the *T. rangeli* SC58 V.2 genomes, as well as their respective singletons. Footnote: The result of the orthology analysis among the CDS of five species of trypanosomatids is represented: *L. amazonensis* in blue, *L. infantum* in red, *T. brucei* in green, *T. cruzi* in yellow, and *T. rangeli* in brown. The numbers within the Venn diagram represent orthologous and paralogous clusters, which contain the analysed CDS. The singletons are represented as circles outside the Venn diagram.

Out of the 298 singletons identified in the *T. rangeli* SC58 V.2 genome, 166 (55.70%) were annotated as hypothetical proteins. From this subset, only six had a functional annotation through the assignment of GO terms, and one had experimental evidence of expression [18]. *In silico* characterization of this gene indicates that it encodes a protein of approximately 34.76 kDa that is addressed to the cell membrane of the parasite. This protein was predicted to have a transferase activity (GO:0016740), specifically hexapeptidase transferase (IPR:1837) due to the presence of a conserved amino acid site.

### 2.4 Chromosome copy number variation and somy estimation of T. rangeli strains

Whole genome ploidy was estimated by the determination of the proportion of alleles in heterozygous SNP positions in all scaffolds (Fig. 4a). In all four *T. rangeli* strains, a peak at 50% was observed, which is expected for diploid genomes, suggesting mostly disomic strains [29,36].

**Figure 4:**
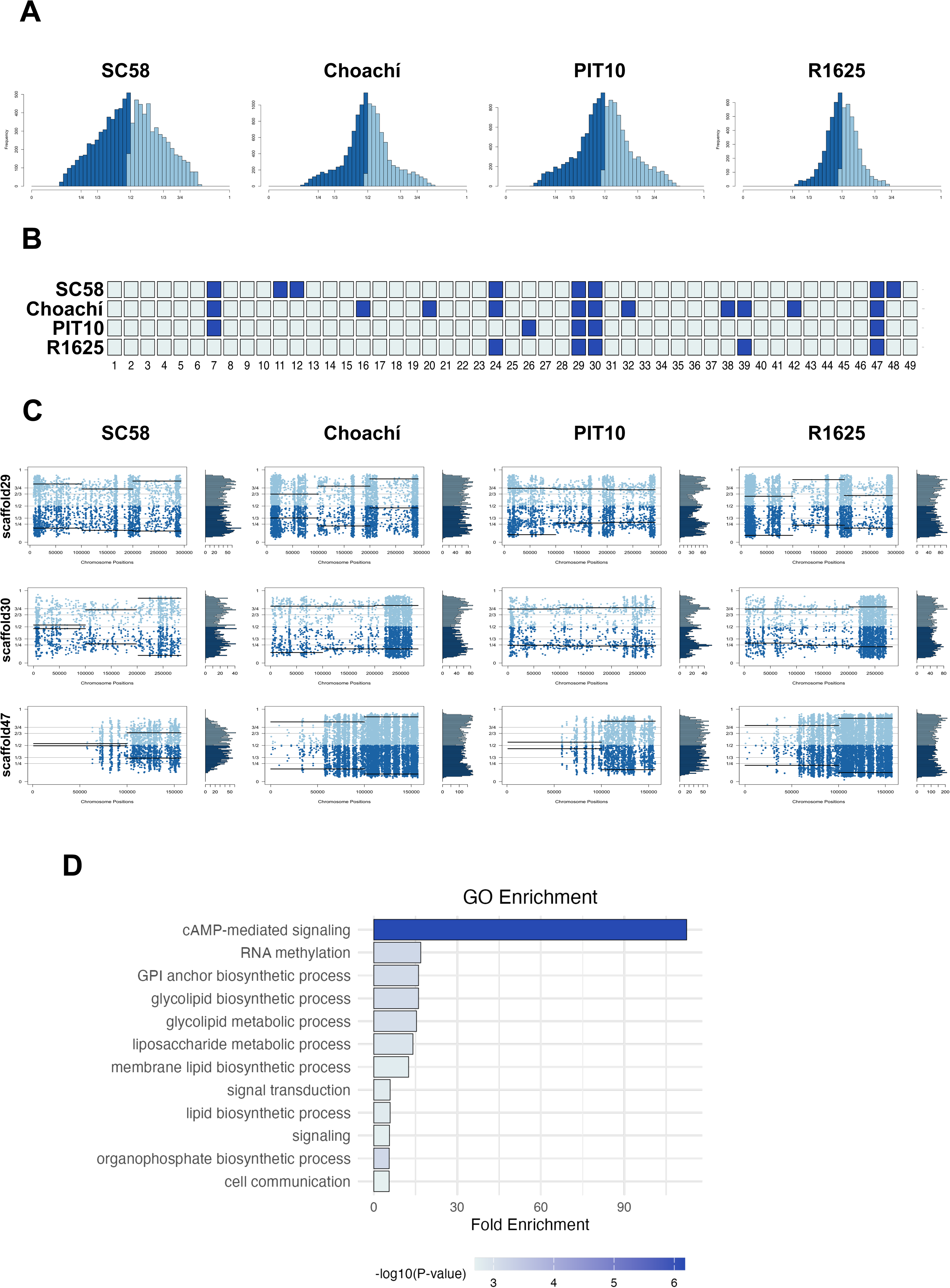
*Trypanosoma rangeli* strains are essentially diploid, with aneuploidies in scaffolds enriched in immune evasion genes. Footnote: (A) Ploidy estimation based on the proportion of alleles at heterozygous SNP positions across each scaffold. (B) Aneuploidy analysis using relative depth of coverage (RDC) across the 50 longest scaffolds. Dark blue indicates scaffolds where the ratio of median scaffold coverage to median genome-wide coverage exceeds 1.5. (C) Allelic proportions at heterozygous SNP positions for scaffolds 29, 31, and 47, supporting some estimations. Blue points represent allele proportions along each scaffold; black lines indicate median values within sliding windows. (D) Gene ontology (GO) enrichment analysis of scaffolds exhibiting shared aneuploidy across all strains.

Chromosome Copy Number Variation (CCNV) in each strain was investigated by combining Read Depth Coverage (RDC) and Allele Balance (AB) analyses [29,36]. Here, the RDC analysis was based on the ratio between the mean coverage of each scaffold and the mean coverage of all scaffolds combined (Fig. 4b). Due to the repetitive content of smaller scaffolds and how repetitive regions lead to erroneous somy estimates [36], CCNV estimation was restricted to longer scaffolds (0.15 Mb) (i.e. scaffolds 1 to 49). Despite applying the size cutoff, some scaffolds still contained too few SNPs for reliable AB estimation (Additional file 5: Supplementary Figure S3). Therefore, aneuploidies were only considered in scaffolds where both RDC and AB results were concordant, providing a two-step confirmation. Aneuploidies, in all cases, chromosome numbers >2, were observed in all strains. Our CCNV analysis combining RDC and AB estimations indicated aneuploidies in scaffolds 7, 11, 12, 24, 29, 30, and 47 in the reference assembly *T. rangeli* SC58 V.2 (Fig. 4b, Additional file 5: Supplementary Figure S3). Similarly, aneuploidies were identified in scaffolds 7, 16, 20, 24, 29, 30, 32, 38, 39, 42 and 47 for strain Choachí, in scaffolds 7, 26, 29, 30 and 47 for strain PIT10, and in scaffolds 24, 29, 30, 39, and 47 for strain R1625 (Fig. 4b, Additional file 5: Supplementary Figure S3). Interestingly, scaffolds 29, 30, and 47 were shown to have additional copies in all strains (Fig. 4c). Notably, these scaffolds are enriched with genes involved in immune evasion and parasite survival, including sialidases, GPI-anchored surface protein biosynthesis, cAMP signaling, RNA processing, and lipid metabolism (Fig. 4d, Additional file 5: Supplementary Figure S3).

### 2.5 T. rangeli GP63 family

A total of 76 complete GP63 gene sequences and 40 pseudogenes were identified in *T. rangeli* SC58 V.2, with 26 of the pseudogenes containing an internal stop codon and 14 exhibiting truncations. The complete *Tr*GP63 sequences include the full-length peptidase M8 domain (NCBI CDD code: cl19482) as detailed in Additional file 6 (Supplementary Table S3). Most of these sequences resemble the canonical metzincin class zinc proteases motif ‘HEXXHXXGX(∼60)H’, which features a conserved methionine at the base of the active site, referred to as the Met-turn. This motif was first described for GP63 in *Leishmania* spp. Furthermore, the sequences contain 19 cysteine residues distributed throughout [37] (Additional file 7: Supplementary Figure S4).

*Tr*GP63 gene sequences were classified based on a phylogenetic analysis that included sequences from different kinetoplastids (Fig. 5), along with two sequences from each of the 11 GP63 groups of *T. cruzi* formerly described [20]. This analysis led to the classification of at least 13 distinct GP63 groups in kinetoplastids, and *T. rangeli* GP63 sequences were included in 11 of these groups. Interestingly, group 12 was found to be exclusive to this species. *Tr*GP63 sequences do not cluster with the G0 and G1, which include sequences found in most kinetoplastids, where G0 mainly includes *Leishmania* spp. sequences and monoxenous invertebrate pathogens such as *Crithidia fasciculata* and *Leptomonas pyrrhocoris,* and G1 contains some functional *Leishmania* spp. sequences (represented by LmjF-10-0460) [38,39], sequences from *Bodo saltans*, *Angomonas deanei* and sequences from other *Trypanosoma* species. Thus, *Tr*GP63 did not group with *T. cruzi* GP63-G1, which intriguingly lacks the Met-turn, and this may reduce its activity. Synteny analysis with the corresponding genomic regions of *T. cruzi* DM28c confirmed the absence of these genes in *T. rangeli*, as the eight upstream and downstream genes are conserved. Except for GP63 group 2, members of all other groups were identified in *T. rangeli* SC58 V.2 as well as in the other three *T. rangeli* strains analysed (Additional file 6: Supplementary Table S3).

**Figure 5.**
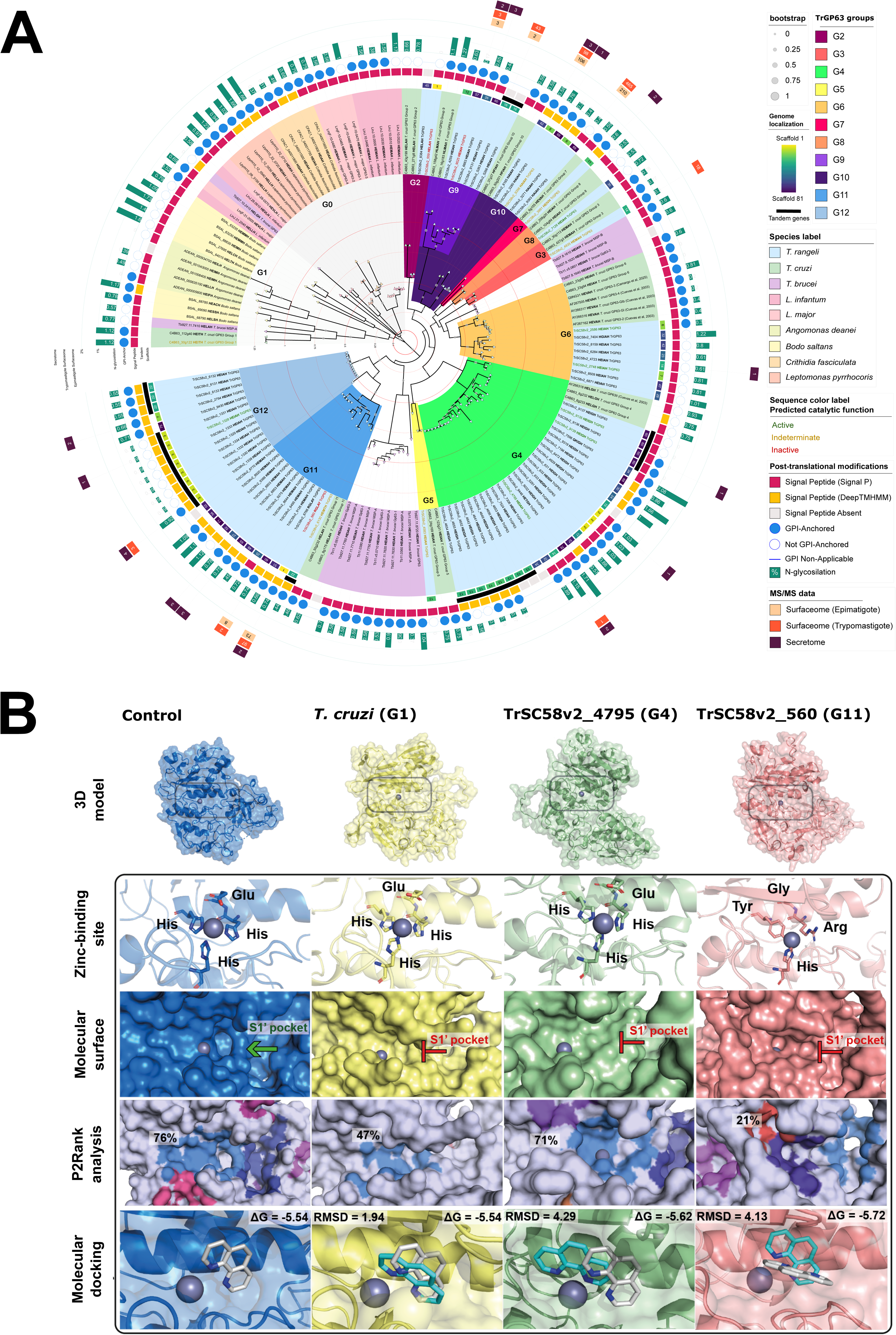
*Trypanosoma rangeli* displays functional secreted and surface-attached GP63 repertoire and species-specific groups from other Kinetoplastids. Footnote: (**A**) Orthology analysis of GP63 in the Kinetoplastea class. Maximum-likelihood tree inferred from full-length GP63 amino acid sequences. The branch scale is shown as red straight and dashed lines. Colored labels indicating metalloprotease inferred activity based on ligand cavity scores are shown as green (≥ 0.5), yellow (<0.5 to ≥ 0.4), and red (<0.4), representing active, indeterminate and inactive, respectively. Genes organized in tandem by at least three contiguous GP63-coding genes or pseudogenes are indicated by a black line on top of the scaffold number. In case of signal peptide prediction indicating absence in both analyses, GPI-anchor results are considered not applicable, indicated as blue lines. Detailed description of sequences and values is provided in Supplementary Table 5. (**B**) Structural and *in silico* functional analysis of *T. rangeli* GP63, in comparison to *Tc*GP63-G1 (C4B63_30g122) and *Leishmania major* (PDB: 1LML) as control. 3D Model: Box highlights the region corresponding to the Zinc-binding site, where structural and functional analyses were performed. The colors of the structures were assigned according to their classification: green for the highly promising group, yellow for the moderately promising group, and red for the less promising group according to ligand cavity scores established parameters. Zinc-binding site: Amino acid residues with close interaction with the zinc ion are shown in stick representation. Molecular surface: presence of the S1’ pocket is demonstrated by a green arrow or absence by a red blocked arrow. P2Rank analysis: Ligand-binding prediction showing predicted cavity positions by different colors. Boxes demonstrate predicted binding cavity probabilities. Molecular docking: Complexes of metalloproteases with the inhibitor 1,10-phenanthroline. The cyan ligand represents the highest-affinity conformation obtained for control structure, while the white ligands correspond to molecular docking results for other GP63. The boxes present the RMSD values (Å) and ΔG (kcal/mol) of the complexes resulting from molecular docking.

Inter-group *Tr*GP63 amino acid sequence identities are approximately 35%, whereas intra-group identities exceed 58%, with G11 being the most variable (Additional file 8: Supplementary Table S4). Four groups have a single representative gene in *T. rangeli* (Groups 3, 5, 7, and 8), while G4 is the largest, with 26 genes. Although *Tr*GP63 grouped with most of the *T. cruzi* groups [20], we did not identify active site residue conservation, except in G6, indicating that groups are formed by general sequence similarity. The classification correlated with the genomic distribution of *Tr*GP63, including the same classification of genes in tandem. All *Tr*GP63 in scaffold 3 belong to group G12, while the 13 *Tr*GP63 in tandem in scaffold 13 were from G4. One GP63 gene from G5, outside the tandem locus, was also identified in this scaffold.

Despite exhibiting a variable presence of both signal peptide and a GPI-anchor signature, most *Tr*GP63 were predicted to be N-glycosylated with an average of 0.62% (± 0.42) residues in the mature protein. The majority of *Tr*GP63 groups were found to be addressed to the parasite cell surface, except members of Groups 2, 6, and 8 that consistently lack a predicted GPI anchor, indicating that these may act extracellularly as free proteins. Some members of groups 3, 4, 9, 10, 11, and 12 showed experimental evidence of protein expression on the parasite surface of epimastigotes and/or trypomastigotes [18], whereas members from groups 4, 9, 10, 11, and 12 were potentially secreted, while members of Groups 2, 6, and 7 were exclusively found in the *T. rangeli* secretome (Additional file 6: Supplementary Table S3), corroborating the hypothesis that these groups have a role as secreted glycoproteins. Interestingly, proteomic evidence of expression from some pseudogenes remains to be addressed.

Metalloprotease activity was experimentally detected in both epimastigote and trypomastigote forms of *T. rangeli*, with higher activity observed at pH 5.5. The activity was considerably lower in *T. rangeli* epimastigotes compared to *T. cruzi* epimastigotes and *L. braziliensis* promastigotes (Additional file 9: Supplementary Figure S5). To assess the metalloprotease functionality in each *Tr*GP63 group, 15 sequences were chosen to generate three-dimensional models based on the available crystal structure of GP63 (leishmanolysin) of *L. major* (LmjF-10-0460/PDB: 1LML) [37]. Additionally, the *Tc*GP63-G1 (C4B63_30g122) was modelled for comparative purposes, since it was missing in *T. rangeli*. The models showed pTM scores ranging from 0.57 to 0.90, indicating a high predictive quality (Additional file 10: Supplementary Table S5). The *Tr*GP63 metalloproteases showed a molecular organization similar to the control structure, shown by global RMSD values between 9.11 and 24.79 Å, with lower variation observed when considering only helix B, located at the catalytic site of the metalloprotease. Overall, most of the *Tr*GP63 sequences displayed a conserved active/Zinc-binding site motif. Certain genes showed punctual substitutions or several alterations in the key residues as identified in, for example, TrSC58v2_9356 and TrSC58v2_560. Among the 15 modelled structures, only TrSC58v2_9135 (G4) and TrSC58v2_1328 (G12) exhibited the large active-site cleft (S1’ pocket) in the surface analysis (Additional file 11: Supplementary Figure S6). Between antiparallel β-sheets in the N-terminal domain, as part of the active site, we identified a loop region with ‘GXXAWA’ Motif (Additional file 5: Supplementary Figure S3), that is mimetic of the S-loop, a well-characterized conformation in vertebrate metalloproteases [40,41].

Considering that the motif conservation and S1’ pocket are important for activity inference, and that other parameters may also be important, the interaction potential of the active sites with ligands was assessed using P2Rank [42], which indicated that seven *Tr*GP63 had a ligand cavity probability above 50%, with TrSC58v2_4795 (G4) and TrSC58v2_2748 (G6) showing values close to the control structure (71%, 74%, and 76%, respectively). These also match the experimentally validated GP63 *T. cruzi* sequences (AY266316, AY266317, AY266318) [43]. These genes were classified as a highly promising group (green structures in Fig. 5b, Additional file 12: Supplementary Figure S6). Further six metalloproteases, including the *Tc*GP63-G1, displayed intermediary values and were classified as a moderately promising group (yellow structures). Finally, three metalloproteases showed values below 40% and were classified as a less promising group (red structures). Molecular docking of the Zinc-binding site region with the inhibitor 1,10-phenanthroline [44] was used to infer the metalloprotease activity. In the *L. major* structure used as control, the selected complex displayed the inhibitor oriented toward the Zn²⁺ ion with a probability of 34% and an affinity of -5.54 kcal/mol. The sequences classified as highly promising showed even higher probability of the inhibitor being faced to the cofactor (67 to 44 %) and low RMSD variations relative to the ligand position in the control structure. *Tc*GP63-G1 presented a behavior similar to that of the highly promising *Tr*GP63, regarding ligand orientation towards Zn²⁺ ion (probability: 56% Docking Affinity: -6.03 kcal/mol), while the structures classified as moderately and less promising did not. Fig. 5b, shows the parameters for the *L. major* structure (control), *Tc*GP63-G1, TrSC58v2_4795 (G4), a highly probable protease that was experimentally identified by two different techniques, and represents the biggest *TrGP63* group, and TrSC58v2_560 (G11) which is unlikely to be an active metalloprotease, despite the high level of expression shown by proteomic analysis in both epimastigotes and trypomastigotes (Fig. 5).

### 2.6 T. rangeli surface Mucins/TASV

A comparative analysis between the previous reference genome sequence and the SC58 V.2 genome revealed an increased number of genes belonging to the mucin-like family in the improved genome sequence, where 15 gene copies were found [5,17]. A detailed search for known signatures of mucin-like genes, such as the signal peptide, GPI-anchor signal, abundance of serine and threonine residues that are likely *O*-glycosylated, as well as orthology with members of the *T. cruzi* mucin-like gene family (*Tc*MUC) on the SC58 V.2 genome, resulted in the identification of seven genes (*Tr*MUC1 - *Tr*MUC7) that are dispersed in three scaffolds of the SC58 V.2 genome (Additional file 12: Supplementary Table S6). *Tr*MUC is slightly more similar to *Tc*MUC type I, sharing an identity of 56.88% ± 22.87, when compared to *Tc*MUC type II (44.34% ± 23.46). Interestingly, a conserved C-terminal region of 22 amino acids is present in members of the *Tr*MUC family, but not in mucins from any other kinetoplastid species present in the TriTryp database (Fig. 6a).

**Figure 6.**
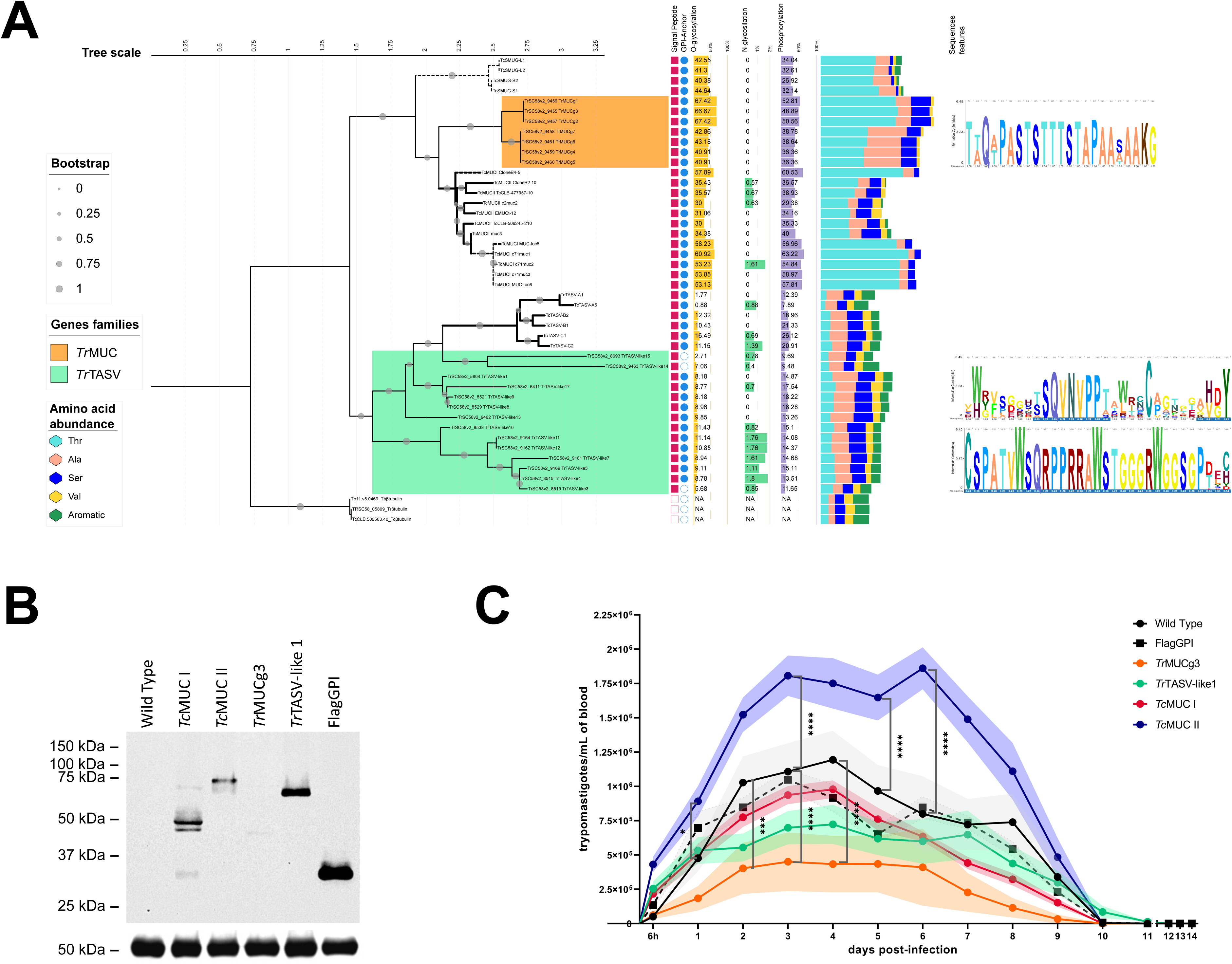
Overexpression of *Trypanosoma rangeli* glycoproteins homologous to *Trypanosoma cruzi* mucin-like and TASV impaired mammalian infection. Footnote: (**A**) Phylogenetic analysis of amino acid sequences of *Tr*MUC, *Tr*TASV-like in comparison to *Tc*MUC I, *Tc*MUC II, *Tc*SMUG and *Tc*TASV. β-tubulin sequences of *Trypanosoma* spp. were used as an outgroup. Bold branches indicate glycoproteins more expressed in *T. cruzi* mammalian forms (trypomastigote or amastigote), and dashed branches show *T. cruzi* replicative forms (epimastigote or amastigote). Bootstrap of 1000 replicates. PTM Percentage is shown internally in the bars considering mature protein size (excluding N- and C- terminal cleavaged sequences). NA stands for ’Non-Applicable’ due to a lack of signal peptide. Logo of exclusive *Tr*MUC motif and representation of two hypervariable regions of *Tr*TASV showing conserved aromatic residues. Logo Y-axis: observed information content (in bits), top: residues alignment position; bottom: occupancy values as gap penalty show in blue. Full description of all sequences and analysis is available in Additional file 12: Supplementary Table S6. (**B**) Molecular mass analysis of homologous and heterologous tagged-fused glycoproteins expressed by the *T. rangeli* Choachí strain. Western blot assay of 50 µg of total protein extract of epimastigotes. Parasites lineages: Wild-type = non-transfected Choachí strain; *Tc*MUCI = tagged-fused ‘clone B4-5’; *Tc*MUCII = tagged-fused ‘clone B2-10’; *Tr*MUC3 = tagged-fused *Tr*MUC3; *Tr*TASV-like1 = tagged-fused *Tr*TASV-like1; FlagGPI = synthetic gene not fused to glycoprotein. Top: glycoproteins recognized by anti-FlagTag monoclonal antibody (mAb). Bottom: Loading control recognized by anti-β-tubulin mAb. Molecular weight marker: Bio-Rad Dual Xtra. (**C**) Infection kinetics assessed by bloodstream parasite load in BALB/c mice (n= 5 animals/group). Intraperitoneal infection of 1x10^7^ *in vitro* trypomastigotes. Data are expressed as means (lines) and standard error of mean (shaded regions). Statistical analysis: two-way ANOVA and Post Hoc Bonferronís multiple comparisons test (*p≤0.05; ***p≤0.001; ****p≤0.0001). Main biological differences are shown in the graph, and full statistical differences are available in Additional file 13: Supplementary Figure S7.

A gene that encodes a 33 kDa surface protein was described in the *T. rangeli* proteome as a putative mucin-like [18], here we identified it as actually belonging to the *T. cruzi* Trypomastigote, Alanine, Serine and Valine rich protein glycoprotein family (*Tc*TASV), previously only found in *T. cruzi* [45].

Based on TASV canonical features that include post-translational modifications (PTM) such as *N*-glycosylation, phosphorylation, and abundance of other amino acids less common in mucin-like proteins, for instance, aromatic residues (phenylalanine, histidine, tryptophan, tyrosine) and cysteine, the SC58 V.2 genome contains 14 TASV-coding genes, and three TASV pseudogenes, dispersed in six scaffolds. Both glycoprotein families are also present in the Choachí and R1625 strains, while *Tr*MUC was not found in PIT10 assembly (Additional file 12: Supplementary Table S6).

A phylogenetic analysis using well described protein sequences of mucin-like families (*Tc*MUC type I and type II), *T. cruzi* small mucin-like family (*Tc*SMUG) [46,47], and *Tc*TASV, where certain members were previously annotated as *Tc*MUC [48] is shown in Fig. 6a.

An assessment of the biological function of *Tr*MUC and *Tr*TASV-like genes was performed by overexpressing selected homologous genes and by expressing *T. cruzi* type I and II mucins (*Tc*MUC I and *Tc*MUC II) in *T. rangeli,* adopting a similar strategy to that proposed by Cánepa et al. (2012). The expression analysis of *Tr*MUC, *Tr*TASV-like, *Tc*MUC I and *Tc*MUC II by transfected *T. rangeli* trypomastigotes and epimastigotes (Choachí strain) demonstrated an expression of hyperglycosylated molecules as indicated by a comparison of their molecular weight to native proteins (Fig. 6b). Variable glycosylation was observed, ranging from a 5X increment in *Tc*MUC I to 2.5x increment for *Tc*MUC II-expressing parasites (Additional file 12: Supplementary Table S6).

Interestingly, overexpression of homologous *Tr*MUC and *Tr*TASV-like resulted in less glycosylated proteins (1.6x and 1.3x, respectively) (Fig. 6b). In addition, overexpression of *Tr*MUC3 by Choachí strain was impaired as indicated by a very faint band around 45 kDa in a Western Blot analysis (Figure 6B), despite all sequences being integrated in the same genome locus under the control of the tubulin promoter and being attached to the parasite surface using the same GPI-anchor sequence.

Expression of these glycoproteins by *T. rangeli* altered the parasitemia profile in the mammalian host, as shown in comparative infection assays in BALB/c mice with wild-type and transfected trypomastigotes (Fig. 6c, Additional file 13: Supplementary Figure S7).

In contrast with *Tc*MUC I-expressing parasites that showed no difference in parasitemia from wild-type and Flag-GPI expressing parasites used as controls (*p* >0.05), *T. rangeli* expressing *Tc*MUC II were found earlier and in higher in number in the bloodstream of the experimentally infected mice from the first (*p*≤0.05) to the 7^th^ day post-infection (d.p.i.) (*p*<0.001) in comparison to controls (Fig. 6c). Overexpression of *Tr*MUC and *Tr*TASV-like1 genes in *T. rangeli*, resulted in a moderate decrease in the parasitemia of *Tr*TASV-like1 transfected parasites in mice between 2 (*p*<0.01) and 4 d.p.i. (*p*≤0.05) when compared to the controls. In contrast, overexpression of *Tr*MUC3 by *T. rangeli* was unsuccessful, as shown in Fig. 6b. Mice infected with *Tr*MUC3-transfected parasites showed variable parasitemia between infected animals, and a significantly reduced parasitemia between the 2^nd^ and 4^th^ d.p.i. (*p*<0.0001), to the 8^th^ d.p.i. (*p*<0.01) (Fig. 6c, Additional file 13: Supplementary Figure S7). The overall parasitemia in mice observed between *T. rangeli* strains from distinct genetic lineages (10-12 days) was not altered by the expression of any of the studied glycoproteins.

## 3. DISCUSSION

The use of a hybrid deep-sequencing approach (PacBio and Illumina) for a *de novo* assembly of the *T. rangeli* genome resulted in a more contiguous and improved resolved genome sequence, especially for repetitive and complex regions containing multicopy gene families and telomeric sequences. Although a large proportion of the gene content was identified in the previous reference genome [5], a reduction of over 90% of the number of scaffolds was achieved. The use of PacBio long reads has improved the contiguity of some of the complex *T. cruzi* strain genomes by resolving repetitive regions [49]. Among these, strains as Bug2148 [50]; Dm28c and TCC [26]; Brazil and Y [51], and Sylvio X10/1 [23] have achieved a near-complete chromosomal resolution. The additional use of Illumina reads in the present study have further increased the contiguity and the quality of the resulting genome assembly, as well as allowing intraspecific comparative analysis.

The identification of 10 possible telomeric regions represents a significant improvement compared to the three telomeric regions previously described in the *T. rangeli* SC58 reference genome, since sub-telomeric regions are key sites for the expression of multicopy genes in trypanosomatids. Telomeric regions were not identified at both ends of any of the scaffolds in the *T. rangeli* SC58 V.2 genome. In *T. brucei*, the sub-telomeric region is responsible for the expression of VSG [52,53]. In *T. cruzi*, the regions adjacent to the telomeres contain (Trans-)Sialidases genes, mostly type II (GP82, GP90, GP85, TSA-1, SA85, and ASP-2) [54], DGF-1, and RHS [52]. In *T. rangeli*, we have previously shown that the sub-telomeric region is shorter than that of *T. cruzi*, due to the absence of multicopy genes from the (Trans-)Sialidases, DGF-1, and other gene families [5].

Polycistronic transcription of trypanosomatids via RNA polymerase II leads to a pre-mRNA that requires maturation by addition of a 39bp-long spliced leader (SL) sequence at their 5’ ends, providing the SL RNA splice donor site and essential motifs for RNA stability and splicing recognition [55,56]. Despite such relevance, the genomic organization and the sequence of SL genes in trypanosomatids vary between species. The sequences of SL genes in *T. cruzi* are conserved but these genes differ widely in copy number and chromosomal location between strains, often appearing on multiple chromosomes [20,22,23]. In *T. brucei*, SL genes are also conserved, but far less dispersed across the genome [57]. Although having a similar structure formed by an exon and an intron of specific lengths as described for *T. cruzi* and *T. brucei*, the *T. rangeli* SL genes are distinctively interspersed with the 5S rRNA genes, a feature conserved among strains of distinct origins [6]. A total of 62 SL gene copies were found across five distinct scaffolds on the *T. rangeli* SC58 V.2 genome, all of them exhibiting the exon/intron structure and being interspersed with the 5S rRNA genes as formerly described.

The improved resolution of this *T. rangeli* genome sequence has revealed additional repetitive genomic content, but the *T. rangeli* SC58 V.2 genome still contains a lower number of genes from multicopy gene families and fewer repetitive genomic elements compared to *T. cruzi* and *T. brucei* overall.

In all studied *T. rangeli* strains, the Sialidase and GP63 families have the highest numbers of genes [5,16]. The large amount of genes coding for sialidases in the *T. rangeli* genome might be related to the involvement of this molecule during cell adhesion and invasion processes as described to other trypanosomatids. Thus, it was initially hypothesized that these enzymes would only be in their active states in epimastigote forms in the vector insect [58]. However, it has been shown through experiments using liquid chromatography coupled with mass spectrometry that sialidases are the most abundant proteins identified in both the epimastigote and trypomastigote forms of *T. rangeli* [18]

The description of VSG-coding genes in the *T. rangeli* AM80 genome by Bradwell et al. (2018) was unexpected, since these surface glycoproteins have only been exclusively described in African trypanosomes, such as *T. brucei*, *T. congolense* and *T. vivax*, but not in *T. cruzi* or *T. rangeli* [5,22,59]. No VSG-related genes were found in the assemblies or reads from all sequenced strains in this study (SC58, Choachí, PIT10 and R1625), nor on previous genomic, transcriptomic or proteomic studies of *T. rangeli* strains.

The results also show that DGF-1 is present in multiple copies in the genome of all studied species of American trypanosomatids. There is experimental evidence that *T. cruzi* trypomastigotes express DGF-1 proteins on the cell surface [60], as computational predictions suggesting an integrin-like activity [60,61]. It has been suggested that DGF-1 may play a role in the contaminative transmission of *T. cruzi* and other American trypanosomatids [61].

All analysed species contain a significant portion of interspersed repeats and a smaller portion of simple repeats. The number of interspersed repeats identified here may be inflated due to the number of unclassified repeats present in the database used [62]. The reliably classified interspersed repeats in the *T. rangeli* SC58 V.2 genome are mostly Long Terminal Repeats (LTR) and Long Interspersed Nuclear Elements (LINEs). The large number of unclassified sequences shows the need for further investigation of the repetitive sequences in trypanosomatids to address their possible biological functions.

Simple repeats were previously presumed to be “junk DNA”, but it is now known that these repetitive elements may function as facilitators of evolution, by providing gene-associated sites of genetic variation with minimal genetic load [63]. Indeed, in the latest version of the genome of the Sylvio X10/1 strain of *T. cruzi*, it was shown that simple repeat regions often flank multicopy genes that encode surface proteins [23]. This configuration could allow micro-homology processes to facilitate recombination between repetitive and similar sequences, resulting in new variants of surface molecules, as described in other parasites [23,64,65]. Thus, the resolution and identification of these regions in the *T. rangeli* SC58 V.2 genome allows recombination sites containing virulence factors to be studied in a non-pathogenic trypanosomatid model.

The orthology analysis showed a shared core genome in all studied trypanosomatids, which is represented by a cluster of 4,501 orthologous CDS groups. This result is consistent with previous studies indicating that trypanosomatid organisms share a significant portion of their genetic content [29,66]. In comparison, 403 orthologous clusters were exclusively shared between *T. cruzi* and *T. rangeli* in the previous genome study of the SC58 strain, whereas this study identified 453 such orthologous groups. This is not surprising, given the close phylogenetic relationship between these parasites [67]. Of the set of 298 unique genes identified in the *T. rangeli* SC58 V.2 genome, 166 (55.70%) are annotated as hypothetical proteins. From this subset of, only one has a functional annotation by assignment of GO terms, and experimental evidence of expression from MS/MS data. The *in silico* characterization of this gene indicates that the corresponding protein is located in the plasma membrane of the parasite, with a conserved site with hexapeptide transferase activity and a unique associated peptide. The identification of this CDS is important for studies of genetic markers with biotechnological applications, especially for the differential diagnosis of Chagas disease.

It is noteworthy to mention that sequences revealing relevant but partial similarities were annotated as pseudogenes on *T. rangeli* SC58 V.2 genome. These may represent remnants of formerly functional genes that underwent pseudogenization via a number of biological mechanisms, as proposed for *L. major*, *T. brucei*, and *T. cruzi* [20,68].

While aneuploidies have been widely reported in trypanosomatids, including *T. cruzi* [36,69,70] and *Leishmania* spp. [71–73], their occurrence in *T. rangeli* remains underexplored. Previous studies using pulsed-field gel electrophoresis (PFGE) documented chromosomal size polymorphisms across *T. rangeli* strains [5,74,75], but ploidy and CCNV analyses were not performed. By using our new reference assembly, with improved resolution of repetitive and complex regions, we estimated the somy of *T. rangeli* SC58 and three additional strains (Choachí, PIT10, and R1625). The CCNV analyses indicated that these genomes are mostly diploid, with aneuploidies that are shared among all strains and ones that are unique to individual strains.

Since some trypanosomatids lack a functional RNA interference pathway [76] and rely on polycistronic transcription, which limits promoter-driven regulation of individual genes, chromosomal duplications can directly influence gene expression by providing additional transcriptional templates [77]. The aneuploidies shared among *T. rangeli* strains in this study were associated with scaffolds enriched with genes involved in immune evasion and parasite survival, including sialidases, GPI-anchored surface protein biosynthesis, cAMP signaling, RNA processing, and lipid metabolism. These functions are likely essential mechanisms to evade both host and vector immune responses. At the same time, the presence of aneuploidies restricted to specific chromosomes in individual strains may reflect selective pressures linked to their geographic distribution and the distinct host- or vector-related environments.

The GP63 gene copy number varies greatly among kinetoplastids. *L. major* carries in total six GP63 genes, four of them arranged in tandem (Chr. 10) [78], while *L. braziliensis* has 39 copies [79]. *T. brucei* contains about 12 genes of the orthologous Major Surface Protease (MSP) divided into three families (seven MSP-A genes, four MSP-B in tandem, and a single MSP-C gene) [80,81]. In contrast, *T. cruzi* (DM28c strain) has 96 genes and 282 pseudogenes belonging to 11 groups [20]. We found that *T. rangeli* possesses an intermediary number of 76 genes and 40 pseudogenes, in 12 groups, in which G12 is a species-specific group, including four groups that are likely to be catalytically active and experimentally found to be secreted and/or attached to the parasite surface. The G2 genes were not identified in previous studies. Neither the phylogenetic nor the synteny analysis in SC58.V2 support the presence of G1 in *T. rangeli*. Additionally, no *T. rangeli* sequence was identified as belonging to G0, the group that includes the majority of *Leishmania* spp. GP63 genes, suggesting a divergence in function and activity.

The structural and functional computational analysis of *Tr*GP63 is consistent with the metalloprotease enzyme class, particularly when compared with leishmanolysin and *Tc*GP63-G1. We verified the ability of these proteins to coordinate a Zn²⁺ ion, an essential requirement for catalytic activity [41]. The presence of the S1’ pocket was only identified in a few *Tr*GP63 sequences. This pocket is located near the active site and has been extensively studied for its ability to accommodate inhibitory molecules and is also essential for the recognition of hydrophobic amino acids present in oligopeptides [40,82]. However, the absence of the active-site cleft cannot be interpreted as evidence of inactivity, as its formation may depend on dynamic structural rearrangements not explored in this study. Moreover, the absence of the S1’ pocket may indicate that these metalloproteases interact with distinct sets of ligands that do not rely on this region for recognition or binding. We identified several *Tr*GP63 structures with high functional potential to act as metalloproteases. This included structures with elevated ligand cavity probability values as predicted by P2Rank. Additionally, the set of structures classified as highly promising showed interaction modes with the inhibitor oriented toward the zinc ion, which is the expected interaction pattern for this inhibitor with metalloproteases [44]. These findings indicate that the active sites of *Tr*GP63 display a biochemical environment comparable to *L. major,* suggesting enzymatic functionality. Notably, the *T. rangeli* metalloproteases in the highly promising group exhibited stronger functional indicators than those observed for the *Tc*GP63-G4 and G6, which have already been characterized as functionally active. Therefore, *Tr*GP63 may perform equivalent functions and represent suitable candidates for functional substitution to verify them *in vitro* using the *T. cruzi* CL Brener strain [43].

The metalloprotease activity of GP63 plays various roles across kinetoplastids, including supporting parasite survival and adaptation to different hosts [43], as well as contributing to basic parasite biology, such as lifecycle progression in *T. brucei* through VSG release along with GPI-specific phospholipase C (GPI-PLC) [81]. In *Leishmania* spp., GP63 is known to act in adhesion, invasion, and immune evasion by interacting with host receptors, modulating complement, degrading signaling pathways to impair immune signaling and vesicular trafficking (phagolysosome biogenesis) and protecting the parasite against oxidative stress [83], via lipid raft–mediated entry, collectively enabling parasite survival inside macrophages [84]. Recently, this has been more clearly defined by identifying the host-target substrates for GP63 [38]. There is a possibility of a non-catalytic role of GP63, including the expression of inactive proteins as adhesins that adhere to the fibronectin-receptor [85]. In addition, GP63 proteins are also expressed by species that do not infect mammals such as *Phytomonas* spp., or even in monoxenic species that develop exclusively in insects such as *Leptomonas* spp., *Herpetomonas* spp., *Angomonas* spp., and *Crithidia* spp. [86]. This suggests a role of GP63 in the defense against the immune system of the invertebrate host, since this metalloprotease is also capable of hydrolyzing antimicrobial peptides produced by insects [86]. Thus, GP63 performs several different roles, which may explain the occurrence of different groups within the same species. The functions of *T. rangeli* GP63 are still largely unknown. A comparative view could suggest a role in macrophage survival since it is known that amastigote-like forms remain viable after internalization by promonocytes [87].

It is known that *T. rangeli* encodes sialidases with hydrolytic activity but lacks enzymes with sialic acid transglycosylase activity [88]. Thus, the finding of a GPI-anchored, mucin-like protein of 33 kDa expressed by *T. rangeli* trypomastigotes [18] led us to investigate the biological function of *T. rangeli* mucins as receptors of sialic acid transferred from the host cell glycoconjugates to the *T. cruzi* surface [89].

In the *T. rangeli* SC58 V.2 genome, it has been possible to determine that this protein belongs to a previously undetected glycoprotein family in *T. rangeli*: *Tr*TASV-like, consisting of 13 complete genes and three pseudogenes. TASV has previously only been described in *T. cruzi* [48,90]. The *Tc*TASV family is conserved among *T. cruzi* strains with around 40 genes in three subgroups (A, B, and C) [45,48,91]. *Tc*TASV-C is the most studied subgroup of this family due to immunoreactivity in natural and experimental infections [45,91,92]. The proteins are mainly shed through extracellular vesicles by bloodstream trypomastigotes, but also attached to the parasite cell [45,93]. Trypomastigotes express *Tc*TASV-C as 60 kDa glycoproteins and, to a lesser degree, as 45 kDa proteins by amastigotes and epimastigotes, while the theoretical molecular mass corresponds to ∼36 kDa. Phosphorylation and glycosylation of *Tc*TASV-C has been experimentally confirmed [91]. The difference in molecular mass in *T. cruzi* trypomastigotes similar to that exhibited by the Choachí strain overexpressing *Tr*TASV-like1, which strongly indicates that *Tr*TASV-like has the same PTMs as *Tc*TASV.

It was possible to show that the overexpression of *Tr*TASV-like1 during *T. rangeli* infection in mice causes a slight decrease in parasitemia at the beginning of the infection. In *T. cruz*i, even more virulent strains exhibit high levels of *Tc*TASV. *Tc*TASV-C delays and decreases parasitemia peaks in the acute mice model [45]. Therefore, new data obtained in *T. rangeli* indicate a possible mechanism of control of parasite number in the bloodstream, possibly through stimulus of phagocytosis and subsequent antigen presentation, which does not necessarily lead to parasite elimination but to intracellular survival instead of exposure in the bloodstream.

In the present genome assembly, a more accurate estimation on the copy number variation of multicopy genes was achieved. We showed that *T. rangeli* has only seven genes that encode mucin-like glycoproteins (*Tr*MUC), which is very different from the high copy number in *T. cruzi* (e.g., 700 in the Brazil A4 strain and 797 in Y strain) [51]. A very tight expression control of *the Tr*MUC3 gene product was also observed, even under the control of a vector that enforces their expression (pROCK plasmid). Despite the low expression level, the infection was impacted by the overexpression of *Tr*MUC3 in trypomastigotes, decreasing the parasitemia in BALB/c mice. The expression of *Tr*MUC thus causes the opposite effect compared to *Tc*MUC II in *T. rangeli*, apparently decreasing the presence of parasites in the bloodstream, while TcMUC II causes an increase. While the expression of *Tc*MUC I by the Choachí strain does not affect the number of parasites in the blood, it is not possible to conclude that there is no influence on the parasite load since this glycoprotein is preferentially expressed in *T. cruzi* amastigotes [94,95], a parasite form that is still unexplored in *T. rangeli* during mammalian infection [87], along with the incomplete understanding of the tissue tropism of this avirulent trypanosomatid in mice models [32,96]. Moreover, *T. cruzi* amastigotes do not express *Tc*TS group I (with trans-sialidase activity) [54,97], alike is well-known for *T. rangeli* [98,99]. Therefore, the best conditions to express *Tr*MUC might be found during an alleged amastigote phase, playing a role with sialidases in an uncharted way, even in the *T. cruzi,* where these enzymes/substrates are abundant. Further studies, mainly focused on characterizing the *T. rangeli* glycoproteins, are needed to comprehend which glycoproteins and how this parasite can acquire sialic acid from the media. Taking together the low copy number, the tight expression control, and the absence of detection of any peak in the proteome studies so far, we also hypothesize that *Tr*MUC is a gene family in the suppression process, possibly a trace of the evolutionary speciation event.

## 4. CONCLUSIONS

The complexity of the trypanosomatid genomes correlates with their intricate biology, presenting intra and interspecific variability mostly due to regions of highly repetitive content such as sub-telomeric sequences. The combination of PacBio and Illumina reads allowed for an improved version of the *T. rangeli* genome, and was also useful for resolving complex and repetitive regions in the genome of the parasite, which is the smallest and least repetitive among the mammalian infective trypanosomatids. Among these repetitive regions, the results presented here demonstrate that the portion of structural repetitive elements is large compared to the quantity of multicopy genes, which is different from other trypanosomatid genomes studied. We highlight the complexity of the *T. rangeli* multigene families of mucins-TASV and GP63 and their involvement in the biology of the parasite.

## 5. METHODS

### 5.1 Parasite culture, DNA extraction and sequencing

Epimastigotes of *T. rangeli* strains Choachí, PIT10, R1625, and SC58 [100] (Additional file 14: Supplementary Table S7) were cultured at 27°C in liver infusion tryptose (LIT) medium supplemented with 10% fetal calf serum (FCS), as described previously [5]. Total DNA was extracted from exponential growth phase epimastigotes using the ReliaPrep™ gDNA Tissue Miniprep System Kit (Promega). Library generation and sequencing were performed at the National Genomics Infrastructure (NGI) hosted by the Science for Life Laboratory, Stockholm (Sweden). *T. rangeli* DNA from all strains was used to prepare paired-end libraries for Illumina (MiSeq or HiSeq 2500) sequencing. Additionally, a SMRT library for PacBio (RSII) sequencing was prepared only for the SC58 strain. Library preparation and DNA sequencing were carried out according to the respective manufacturer’s protocol.

*T. rangeli* Illumina reads with Q-score mean quality <30 were removed from the analysis, following removal of the adapters from the high-quality reads using the Trimmomatic v.0.39 software [101]. PacBio reads were trimmed based on sequence length (>1Kb) using the FiltLong v.0.2.1 software (https://github.com/rrwick/Filtlong).

### 5.2 Hybrid assembling of T. rangeli SC58 strain genome

Hybrid assembly of the SC58 strain genome was performed *de novo* using high quality PacBio reads by the software CANU v.2.2 [102], following incorporation of Illumina reads into the assembly during the scaffolding and gap-filling steps, using the SSPACE Standard v.3.0 [103] and GapFiller v.1-10 software [104], respectively. Both steps were run through multiple iterations to compensate for base calling errors of PacBio reads. Polishing of the assembled genome was performed using Illumina reads with Q-score mean quality >34 using the Pilon v.1.23 software [105]. Finally, automated allelic content reassignment was performed by the Purge Haplotigs software v.1.1.2 [106].

General assembly metrics of *T. rangeli* SC58 V.2 were evaluated using the QUAST v.5.0.2 software [107] and mapping with the SC58 reference genome was performed using the “dnadiff” algorithm of the MUMmer4 v.1.4 software [108].

The sequences corresponding to *T. rangeli* kDNA maxicircles and minicircles were removed prior assembling using the BLASTn and BLASTx algorithms from the BLAST+ software v.2.9.0 [109], against a database of kDNA sequences from *T. brucei*, *T. cruzi*, and *T. rangeli*.

Further validation of the *T. rangeli* SC58 V.2 genome assembly was conducted by mapping 130 universal single-copy orthologs of the Euglenozoa phylum using BUSCO v.5.5.0 software [110], along with *in silico* mapping of *T. cruzi* probe sequences specific to chromosomes 4, 37 and 39 [111] onto the SC58 V.2 genome assembly (Additional file 3: Supplementary Table S1). For this later, a high-stringency (E-value >1e-10, coverage >60%, identity >60%) similarity analysis of the *T. cruzi* probes against the *T. rangeli* SC58 V.2 genome scaffolds were carried out using the BLASTn and tBLASTx algorithms.

### 5.3 Structural and functional annotation of the T. rangeli SC58 V.2 genome

CDS prediction and automatic annotation were performed using the AnnotaPipeline v1.0 software [112], using default parameters for similarity analysis (E-value cutoff = 1e-5) and peptide identification (Q-value cut-off = 0.05). Gene prediction was performed using AUGUSTUS v.3.3.3 [113] and an in-house training set composed of *T. rangeli* SC58 expressed sequence tags (EST) [17] and deduced amino acid sequences from the reference genome [5]. Automated annotation of CDS was based on sequence similarity analysis using the BLASTp algorithm, matching with the protein sequence dataset from the Swiss-Prot v.2022 [114] and TriTrypDB v.65 [115] databases. CDS showing no similarity were automatically annotated as hypothetical. Search for protein domains and families was performed using InterProScan v.5.61-93.0 [116] allowing assignment of Gene Ontology (GO) classification based on the identified protein domains. Transcriptomic [17] and proteomic [18] datasets generated from epimastigotes and trypomastigotes forms of *T. rangeli* were used to assess evidence of gene transcription by Kallisto v.0.48.0 [117], and evidence of expression by Comet v.2021.01 [118] and Percolator v.3.05.0 [119]. A profile for each protein composing the final set of predicted and annotated CDS was generated using FastProtein v.1.0 [120]. Furthermore, CDS separated by less than 100 base pairs, along with adjacent ones sharing similar annotations from the previous analysis, were manually reviewed to assess the potential presence of pseudogenes.

Prediction of ribosomal RNA (rRNA) and transfer RNA (tRNA) was based on sequence similarity (Identity >90%) with the Rfam database v.14.2 [121] using the software Infernal v.1.1.2 [122] and validated based on coverage and identity via BLASTn using the RNAcentral database v.20 [123].

Analysis of structural repetitive elements was performed using the software RepeatMasker v.4.1.5 [124], disregarding insertions of bacterial origin and using a custom library containing sequences specific to trypanosomatids retrieved from msRepDB [62].

The presence of telomeric regions on the extremities (∼10Kbp) of each scaffold of the *T. rangeli* SC58 V.2 genome was assessed by search of the canonical telomeric repeat of eukaryotes (TTAGGG)n [1,52], using the Seqtk v.1.4 (https://github.com/lh3/seqtk) and the Tidk v.0.2.31 software [125].

### 5.4 Multicopy gene families

Comparative analysis of multicopy gene families (amastins, DGF-1, GP63, KMP-11, mucins, MASP, RHS, sialidase/trans-sialidases, tuzins, and Variant Surface Glycoproteins - VSG) was carried out between the *T. rangeli* (SC58 V.2, SC58 reference genome, AM80), *T. cruzi* (CL Brener Esmeraldo-like and Sylvio X10/1), and *T. brucei* (TREU927) genome.

Initially, a search for keywords across CDS annotations of each species genome was performed using the software SeqKit v2.5.1 [126] using the following definitions as queries: *amastin*, *dgf*, *dispersed, dispersed gene family*, *gp63*, *leishmanolysin*, *kmp*, *kinetoplastid*, *kinetoplastid membrane protein*, *masp*, *mucin-associated*, *mucin-associated surface protein*, *muc*, *mucin*, *rhs*, *retrotransposon*, *retrotransposon hot spot*, *sialidase*, *vsg*, *surface glycoprotein*, *variant surface glycoprotein*, and *tuzin*. Related terms, such as “MASP” and “Mucin-Associated Surface Proteins”, for example, were considered as belonging to the same multicopy gene family to avoid redundancy. The annotation of each CDS was manually curated to remove false positives and to assign correct description of their products. In this sense, all genes whose automatic annotation were “trans-sialidase” or “leishmaniolysin” were corrected to “sialidase”, since *T. rangeli* lacks trans-sialidase activity [34], and to “GP63”, since members of this multicopy gene family differs from *Leishmania* species, respectively.

Furthermore, a similarity analysis was then carried out between the predicted CDS of each multicopy gene family and the corresponding transcript data of trypanosomatid species on TriTrypDB. This step was performed by the BLASTx algorithm of the software BLAST+ v.2.14.1 [109], considering an E-value cutoff of 1e-5. Results from this analysis were only considered if the corresponding match presented a coverage of over 30%, at least 50% identity and 50% positivity. Hence, redundancy was removed once more, to ensure no sequence duplicates between the dataset obtained from the similarity analysis and the recovered annotated CDS. Finally, the total number of nucleotides for each multicopy gene family was obtained using the “stats’’ algorithm of the software SeqKit.

### 5.4.1 T. rangeli GP63

Sequences identified by BLAST as GP63 underwent further examination using the NCBI’s Conserved Domain Database (CDD v3.21). Sequences shorter than 200 amino acids and classified as “incomplete”, “frameshift” or “interrupted Peptidase M8 Superfamily (cl19482)” were considered pseudogenes. Fragmented CDS with CDD e-value > 1x10^-32^ for the Peptidase M8 Superfamily were artificially combined with upstream or downstream coding sequences that contain complementary domains in the most repetitive nucleotides at the break points, primarily occurring in glycine or proline residues. The identification of the zinc-binding and active site was accomplished by recognizing the “ZnMc astacin-like” domain (cd04280) and conducting a manual search using Jalview version 2.11.4.1 [127]. The characterization of the signal peptide, glycosylphosphatidylinositol (GPI) anchoring, and N-glycosylation was performed through the use of SignalP v.6.0 [128], DeepTMHMM v.1.0 [129], NetGPI v.1.1 [130], NetN-Glyc v.1.0 predictors, respectively.

Analysis of orthology of GP63 genes among trypanosomatids used a dataset of 156 sequences from *T. cruzi* strains Dm28c (21) [20] and CL Brener (5) [43] , *L. infantum* JPCM5 strain (7), *L. major* Friedlin strain (6) [38,131,132], *T. brucei* TREU927 strain (17) [78]. In addition, sequences from *Angomonas deanei* strain Carvalho ATCC PRA-265 (5), *Leptomonas pyrrhocoris* strain H10 (5), *Crithidia fasciculata* - strain Cf-Cl (5), and the free-living kinetoplastid *Bodo saltans* strain Lake Konstanz (8) were obtained by BLASTp search in TriTrypDB using the *L. infantum* GP63 entries (LinJ.10.0500 and LinJ.10.0520) as query [78,79]. Phylogenetic analysis was carried out using the workflow provided by NGPhylogeny.fr [133] with default parameters of each included tool. Data was then integrated using the Interactive Tree of Life (iTOL) v.7.2 online tool [134]. Identity matrix analysis was performed using Clustal Omega (EMBL-EBI Job Dispatcher [135]) and SIAS tool (http://imed.med.ucm.es/Tools/sias.html). Active site motif was describe using ScanProsite tool [136].

Metalloprotease activity assays were carried out using a total of 1 × 10⁸ parasites, which included *Leishmania braziliensis* (MHOM/BR/75/M2904 strain promastigotes), *T. cruzi* (Y strain epimastigotes), and *T. rangeli* (Choachí strain epimastigotes and trypomastigotes). The parasites were lysed in a buffer containing 1% Triton X-100 and 10 mM Tris-HCl at pH 6.8. SDS-PAGE gels were prepared with 0.2% gelatin, and the samples were neither heated nor reduced prior to analysis. Following electrophoresis, the gels were washed twice with a 0.1 M sodium acetate buffer (pH 5.5) that contained 2.5% Triton X-100. Subsequently, the gels were incubated for 48 hours at 37 °C in a buffer consisting of 0.1M sodium acetate and 1mM DTT at pH 5.5. In some cases, 10mM 1,10-phenanthroline (Sigma-Aldrich) was included in the incubation.

Finally, the gels were stained with Coomassie Brilliant Blue R-250. To evaluate the impact of pH, additional experiments were conducted by incubating the gels at 37°C for 48 hours in reaction buffers with varying pH values: 3.5, 5.5, 7.0, and 9.0.

Building on the proteomic evidence of GP63 expression in *T. rangeli* [18], we conducted further analyses of the raw MS/MS data from the surface-enriched protein fraction (SEP) of *T. rangeli* SC58 strain for detailed identification of GP63 expression. This analysis employed MSFragger [137] and, also incorporated unpublished, in-house generated secretome datasets from *T. rangeli*.

The final dataset comprises all 11 GP63 sequences, supplemented with contaminant proteins from cRAP (https://thegp_m.org/crap/index.html), including porcine trypsin, *Saccharomyces* spp. alcohol dehydrogenase, and human keratin. Decoy database searches were performed automatically. MSFragger parameters specified tryptic peptides allowing one missed cleavage, carbamidomethylation of cysteine, methionine oxidation as a variable modification, and precursor and fragment tolerances of 10 and 5 p.p.m., respectively. Protein identification and scoring were conducted using the ProteinProphet algorithm within Philosopher [138]. Only proteins supported by at least two unique peptides, a false discovery rate (FDR) below 0.1%, and peptide/protein probabilities exceeding 95% and 99%, respectively, were considered as valid.

#### 5.4.2 Structural and functional analysis of T. rangeli metalloproteases

To investigate the presence of metalloprotease activity in *T. rangeli*, we conducted *in silico* structural and functional analyses of these proteins. Initially, we generated 3D models of *T. rangeli* metalloproteases using AlphaFold 3 [139]. For comparative purposes, we also obtained the model of the group 1 metalloprotease from *T. cruzi* (TriTrypDB: C4B63_30g122). All models were generated using the default parameters of AlphaFold 3, as outlined in its user guide. Quality assessment of the model was carried out using the confidence scores provided by the program, including pLDDT (per-residue confidence) and pTM (predicted template modelling). Specifically, pTM values above 0.5 were considered indicative of adequate modelling, while values above 0.8 reflected high predictive quality [139]. All generated models were visualized, analysed, and figures were created using PyMOL v.2.5.0 (https://www.pymol.org/).

For the structural and functional analyses of metalloproteases, leishmanolysin - a metalloprotease from *L. major* (PDB: 1LML) - was utilized as the control structure [37]. The structural analysis focused on the identification of the active site or zinc-binding motif (HEXXHXXG[∼60]H) located in helix B, which is a conserved characteristic of metalloproteases, was evaluated [41,44]. Surface representations of the models were created to identify the S1’ pocket, a common feature of these proteins extensively studied for its importance in interactions with various inhibitors [140]. The loop region preceding helix B, located between antiparallel β-sheets in the N-terminal domain, was examined due to its presence in the active site of the metalloproteases. Furthermore, we assessed the structural similarities between the metalloprotease models of *T. rangeli*, *T. cruzi* and the control *L. major* models through structural alignments. This process was followed by RMSD (Root Mean Square Deviation) calculations for both the overall protein and for helix B specifically, considering only the alpha carbons in each case.

To evaluate the interaction potential of the modeled metalloprotease active sites with ligands, the proteins were submitted to the P2Rank web server [42]. P2Rank is a machine-learning tool that identifies potential binding sites by analysing surface features, shape, and electrostatic potential. The server provides a ranked list of potential binding sites, derived from the P2Rank score and the ligand cavity probability. The metalloprotease models were categorized into three groups based on the ligand cavity probability value: (i) highly promising, when probability is ≥ 50%; (ii) moderately promising, when it falls between < 50% and ≥ 40%; and (iii) less promising, when it is < 40%.

The metalloprotease models underwent molecular docking with 1,10-phenanthroline, an established inhibitor of metalloproteinases [38,44]. This compound typically interacts with the active site of these enzymes, specifically targeting the Zn²⁺ ion, which serves as an indicator of a catalytic environment comparable to that of metalloproteinases. Docking simulations were carried out using AutoDock Vina [141]. An optimized grid box was strategically positioned over the active site to generate nine poses for each protein, while all other parameters adhered to the program’s default settings. Initially, the control structure was docked with 1,10-phenanthroline, and the pose with the lowest affinity value and the most favourable orientation toward the Zn²⁺ ion was selected as the reference model. Subsequently, docking was conducted with the metalloproteases from *T. rangeli* and *T. cruzi*, focusing on poses that displayed the highest similarity to the control complex. To accomplish this, the proteins were aligned, and the RMSD between the ligands was calculated, with the pose showing the lowest RMSD chosen for further analysis.

#### 5.4.3 T. rangeli mucin-like glycoproteins

Identification of *T. rangeli* mucin-like glycoproteins was carried out by BLASTn using the first 93 nt of *T. rangeli* mucin-likes TRSC58_06796 (TriTrypDB) and KC544919 (GenBank) genes described in the reference genome [5] and proteome [18], respectively. Also, the presence of N- and C-terminal domains of *T. cruzi* mucin-like genes signatures [142,143] was assessed. Hits that did not match to a CDS automatically annotated were evaluated by analysis ORFs obtained from 3 Kbp flanking regions of identified hits using the GetORF tool (https://www.bioinformatics.nl/cgi-bin/emboss/getorf), considering the following parameters: 150 nt minimum length; and only Met as start codon. Only sequences encoding for valid signal peptides according to SignalP v.6.0 predictor [128] or DeepTMHMM v.1.0 [129] were considered as putative *T. rangeli* mucins (*Tr*MUC). The presence of GPI-anchor, *N*- and *O*-glycosylation sites, in addition to phosphorylation (ATM, CKI, CKII, CaM-II, DNAPK, EGFR, GSK3, INSR, PKA, PKB, PKC, PKG, RSK, SRC, cdc2, cdk5, and p38MAPK) recognition sites was evaluated using NetGPI v.1.1 [130], NetN-Glyc v.1.0, NetO-glyc v.4.0 [144], and NetPhos v.3.1 [145] servers, respectively. Prediction of amino acid abundance and molecular weight was performed by Pepstats tool available at EMBL-EBI Job Dispatcher [135]. The conservative regions were evaluated by Gblocks [146] applying Clustal Omega [147] alignments as inputs, as well as to obtain logos representations through the Skylign tool [148], and overall similarity analysis performed by SIAS tool. Phylogenetic analysis based on maximum-likelihood tree inferred amino acid sequences identified from the *T. rangeli* SC58 V.2 genome, as well as members of the *T. cruzi* mucin-like (MUC) and Trypomastigote Alanine-Serine-Valine (TASV) glycoprotein families retrieved from TryTripDB and Genbank databases was conducted as described for GP63 analysis.

#### 5.4.4 Biological assessment of T. rangeli mucins and TASV

We designed a synthetic gene (purchased from Biomatik) hereby called ‘FlagGPI’ encoding a 3xFlagTag^®^ (Sigma-Aldrich) sequence, followed by the mNeonGreen fluorescent protein gene [149], and a *T. rangeli* GPI-anchor sequence (KC544919-GenBank) under control of *T. cruzi* gp82 3’ untranslated region (UTR). The GFP gene present in the integrative plasmid pROCK-Neo [150] was replaced by the FlagGPI gene through *Xba*I and *Xho*I restriction enzyme cleavage, generating the FlagGPI-pROCK vector.

The *T. rangeli* (Choachí strain) genes MUCg3 and TASV-like1, and the *T. cruzi* (Y strain) genes MUC I (clone B4-5), and MUC II (clone B2-10), were PCR amplified from genomic DNA. The amplicons, that excluded the GPI-anchor addition sequence, were gel purified using the QIAquick Purification kit (Qiagen), cloned in pGEM T-Easy vectors (Promega), sequenced (ABI3500 - Applied Biosystems), and subcloned in the FlagGPI-pROCK vector, resulting in tagged-fused constructs. A complete list of primers and restriction enzymes is provided in Additional file 15: Supplementary Table S8.

Epimastigotes of *T. rangeli* (Choachí strain) were transfected with 10 µg of *Not*I-linearized FlagGPI-pROCK vector containing the distinct tagged-fused constructs or with the FlagGPI synthetic gene, using the Nucleofactor^®^ II (Lonza) U-33 program. The transfected parasites were cultivated in LIT+NNN and selection was performed by increasing concentrations of geneticin (G418) up to 500 μg/mL. Plasmid integration into the parasite β-tubulin locus was confirmed by PCR using a set of primers designed to anneal exclusively in the genomic β-tubulin region upstream of the homologous recombination site and the HX1 5’ UTR present particularly in FlagGPI-pROCK vector (Additional file 15: Supplementary Table S8). Once the integration into the parasite genome was confirmed, parasites were cultivated in the absence of selective drugs.

To assess the expression of the FlagGPI-fused glycoproteins, total protein extracts of *T. rangeli* epimastigotes and trypomastigotes from each transfected lineage and wild-type parasites were obtained by lysis in proper buffer (50 mM NaCl, 200 mM Tris-HCl pH 8.0, 1% (v/v) Triton X-100), supplemented with a protease inhibitor cocktail (Sigma-Aldrich). A total of 50 μg of each extract were resolved in 12% SDS-Page gels and then wet-transferred to Hybond^®^ PVDF membranes (Amersham). Membranes were incubated with primary antibodies (mouse anti-FlagTag M2 or rabbit anti-β-tubulin), then with HRP anti-IgG conjugates (anti-mouse or anti-rabbit) (Sigma-Aldrich). The expression analysis was periodically assessed.

Comparative *in vivo* assessment of the infectivity of all transfected lineages was performed using 6 weeks old female BALB/c mice, maintained in ventilated cages under a 12-hour light/dark cycle and having water and food *ad libitum*. A total 1x10^7^ *in vitro* differentiated *T. rangeli* trypomastigotes were injected by intraperitoneal (i.p.) route and parasitemia was assessed daily by the Pizzi-Brener method from 6h post-infection [151]. *In vivo* experiments were approved by the UFSC Animal Ethics Committee (Protocol 9923170516). Graph and statistical analysis were performed using GraphPad Prism v.8.4.0 software. The expression of *T. rangeli* mucins and TASV was also evaluated by search on *T. rangeli* MS/MS proteomic data as performed for GP63 (Item 2.4.1.).

### 5.5 Analysis of orthologous groups

Orthology analysis was performed with OrthoFinder v.2.3.11 [152] using the predicted proteins from *T. rangeli* SC58 V.2 genome and proteins from *T. brucei* (TREU927 strain), *T. cruzi* (CL Brener Esmeraldo-like and Sylvio X10/1 strains), *T. rangeli* (SC58 reference strain), *Leishmania amazonensis* (MHOM/BR/71973/M2269 strain) and *Leishmania infantum* (JPCM5 strain), obtained from TriTrypDB version 65 [115], and also *T. rangeli* (AM80 strain) retrieved from GenBank (Access code: GCF_003719475.1).

### 5.6 Comparative intra-specific genomics of T. rangeli strains

Illumina paired-end libraries of distinct *T. rangeli* strains (SC58, Choachí, PIT10, and R1625) were quality-filtered using Trimmomatic v0.40 [101] and the filtered reads were mapped against the *T. rangeli* SC58 V.2 genome using the Burrows-Wheeler Aligner v. 0.7.17 [153].

For SNP calling, the mapping files were sorted with Samtools v.1.17 [154], PCR-duplicates were marked with Picard v.2.27.5 (http://broadinstitute.github.io/picard/), and SNPs were identified using the Genome Analysis Toolkit v.3.8 (https://gatk.broadinstitute.org/hc/en-us) using the “HaplotypeCaller” algorithm with a minimum quality value of 30 and a minimum depth of coverage of 10. A strict mapping quality filter was applied to remove the influence of low mapping quality in repeated regions and any SNP with mapping quality <50 was removed using BCFtools v.1.9 [154].

CCNV and ploidy estimation were performed using a combination of RDC and AB methods as previously described [36]. Briefly, the estimation of CCNV was based on the ratio between the mean coverage in each *T. rangeli* SC58 V.2 scaffold and the mean coverage of all scaffolds in the assembly. If the ratio between the median chromosome coverage and the median genome coverage was approximately one, the chromosome had the same copy as the genome overall, while fluctuations in this ratio (<0.75 or >1.25) were considered as putative aneuploidies. Heterozygous SNPs with a mapping quality higher than 50 were selected for AB analyses to confirm aneuploidies and to estimate chromosome somy and whole genome ploidy. Whole genome ploidy was estimated by analysing the proportion of the alleles in heterozygous SNP positions of all scaffolds. To confirm the RDC somy estimations, the proportion of each allele with heterozygous SNPs per chromosome position was plotted for the 82 scaffolds. All RDC and AB graphs were generated in R with ggplot2 (available at https://ggplot2.tidyverse.org), pheatmap (available at https://CRAN.R-project.org/package=pheatmap), and vcfR [155] packages. Gene Ontology Enrichment analyses in CCNV regions were performed using TriTryp tools (https://tritrypdb.org/tritrypdb/app/) based on orthologs with the *T. cruzi* reference CL Brener Esmeraldo-like.

## ADDITIONAL FILES

**Additional file 1: Supplementary Figure S1**. Comparative alignment of the assemblies of *Trypanosoma rangeli* SC58 V.2 genome with other *T. rangeli* and *T. cruzi* genomes.

Footnote: Dot-plot representation of the alignment of the *T. rangeli* SC58 V.2 genome assembly with the reference genome (A), *T. rangeli* Choachí (B), PIT10 strain (C) and R1625 (D) strains genomes, *T. cruzi* CL Brener Esmeraldo-like (E), and Sylvio X10/1 strains genomes using comparative dot-plots generated by D-Genies (Cabanettes et al., 2018).

**Additional file 2: Supplementary Figure S2.** *Trypanosoma rangeli* five loci of the Spliced Leader RNA genes interspersed with 5S RNA genes.

Footnote: (**A**) Alignment of 62 *T. rangeli* Spliced-Leader (SL) RNA genes highlight the exon region (light green), intron (light red), and splicing site at position 39. (**B**) Five loci show SL genes interspersed with 5S RNA genes, varying from 9 to 16 SL gene copies.

**Additional file 3: Supplementary Table S1**. Mapping of *Trypanosoma cruzi* (CL Brener strain) chromosome-specific probes onto the *Trypanosoma rangeli* SC58 V.2 genome.

Footnote: *Access codes of *T. cruzi* genes in the TriTrypDB v.65 database. **Genomic location and “P” and “S” haplotypes of *T. cruzi* CL Brener genome (Weatherly; Boehlke; Tarleton, 2009).

**Additional file 4: Supplementary Table S2**. Telomeric regions identified in the

*Trypanosoma rangeli* SC58 V.2 genome scaffolds.

Footnote: bp = base pairs.

**Additional file 5: Supplementary Figure S3 -** Allelic proportions at heterozygous SNP positions for the 81 scaffolds of each *Trypanosoma rangeli* strain (SC58, Choachí, PIT10, R1625). Blue points represent allele proportions along each scaffold; black lines indicate median values within sliding windows.

**Additional file 6: Supplementary Table S3**. Summary of GP63 genes of *T. rangeli* and other Kinetoplastids species are presented in the phylogenetic analysis.

Footnote: ‘GPI-Anchored’ in red= sequence forecasted as GPI-anchored, however biologically not possible if signal peptide prediction is correct. ‘BNP’ = biologically not possible; SP = Signal Peptide; GLOB = globular; TM = transmembrane, ----- = presence of gap in active site region; XXxxX = Non-conserved amino acids in zinc-binding site/active site (other than HExxH). NE: Non-evaluated. Peptidase_M8 Superfamily (NCBI CDD code: cl19482).

**Additional file 7:Supplementary Figure S4.** *Trypanosoma rangeli* Surface Glycoprotein 63 (GP63) presents the main classical leishmanolysin motifs.

Footnote: Logo representation of residue conservation as weighted counts of Zinc-binding site. Logo Y-axis = observed information content (in bits), top = residues position in the alignment; bottom = occupancy values.

**Additional file 8: Supplementary Table S4**. Amino Acid Percent Identity Matrix of *Tr*GP63 complete genes.

Footnote: Colors highlight the *Tr*GP63 groups categorization as shown in the header of the matrix.

**Additional file 9:Supplementary Figure S5.** *Trypanosoma rangeli* exhibits metalloprotease activity, and it is biochemically inhibited by chelating agents.

Footnote: (**A**) Total protein extracts from epimastigotes (E) and culture-derived trypomastigotes (T) of *T. rangeli* (10⁸ cells) were analysed by SDS-PAGE (10%) using a 0.2% gelatin-containing gel, without heating or reducing the samples before electrophoresis. Gels were incubated at 37 °C for 48 h in reaction buffer, without (-) or with (+) specific inhibitor 1,10-phenanthroline. Proteolytic activity is visualized as clear bands against the Coomassie brilliant blue-stained gelatin background. (**B**) Total protein extracts from axenic forms of *T. rangeli* Choachí strain (Tr), *T. cruzi* Y strain (Tc), and promastigotes of *L. braziliensis* MHOM/BR/75/M2904 strain (10⁸ cells) were analysed in gels incubated with reaction buffers at different pH conditions.

**Additional file 10: Supplementary Table S5**. Values from the structural and *in silico* functional analyses of *Trypanosoma rangeli* GP63 metalloproteases.

Footnote: Metalloprotease from *L. major* (PDB: 1LML) was used as a control, and a single-sequence from *T. cruzi* group 1 (TriTrypDB:C4B63_30g122) was included for comparative analysis. Colors indicate metalloprotease inferred activity based on ligand cavity scores: Green for highly promising (≥ 0.5), yellow for moderately promising (<0.5 to ≥ 0.4), and red for moderately promising (<0.4). pTM = predicted template modelling. RMSD = root mean square deviation.

**Additional file 11: Supplementary Figure S6.** Structure of *Trypanosoma rangeli* GP63 metalloproteases from distinct groups.

Footnote: **(A)** Structural and *in silico* functional analysis of *T. rangeli* GP63 classified as a highly promising group regarding inferred activity. (**B**) *Tr*GP63 is classified as a moderately promising group. (**C**) *Tr*GP63 was predicted as a less promising group. 3D Model: Three-dimensional models demonstrating the region corresponding to the active/Zinc-binding site (Highlighted by a box), where structural and functional analyses were performed. Zinc-binding site: Visualization of the conserved amino acid residues are shown in stick representation. Molecular surface: representation of the proteins indicating the presence or absence of the S1’ pocket representing the position which interacts with inhibitors. Presence of S1’ pocket is indicated by a green arrow, and absence by a red arrow. P2Rank analysis: Ligand-cavity prediction shown by colours demonstrates different predicted cavity positions. Predicted binding cavity probability is shown in the boxes. Molecular docking analysis: Complexes of metalloproteases with the inhibitor 1,10-phenanthroline. The cyan ligand represents the highest-affinity conformation obtained for the control structure, while the white ligands correspond to molecular docking results for the other GP63. Blue indicates nitrogen in both representations. The boxes present the RMSD values (Å) and ΔG (kcal/mol) of the complexes resulting from molecular docking.

**Additional file 12: Supplementary Table S6**. Description of *Tr*MUC and *Tr*TASV genes and predicted glycoprotein characteristics.

Footnote: Yellow background= *T. rangeli* glycoproteins selected for biological characterization. “MW”= Molecular Weight marker, “WB”= Western Blot. *excluded mature FlagGPI construct 30.39 kDa. #GPI-anchor cleavage did not occur in the synthetic gene due to a lack of signal peptide. Aromatic residues (Phe, His, Trp, Tyr). NP = Non-provided by DeepTMHMM Signal peptide predictor.

**Additional file 13: Supplementary Figure S7:** Comparative parasitemia between wild-type *Trypanosoma rangeli* and transfectants expressing T*r*MUC, *Tr*TASV-like1, or *Tc*MUC genes.

Footnote: Graphs expressed as means and standard error of mean by lines and shaded regions, respectively. Statistical analysis: Normal distribution of data evaluated by Kolmogorov-Smirnov test (ɑ=0.05), followed by two-way ANOVA and Post Hoc Bonferronís multiple comparisons test. Differences between groups per time point are shown in the graphs. Unique statistical symbols demonstrate the same statistical difference in all time points between 1 to 10 d.p.i. (ns *p>*0.05; \**p*≤0.05; \*\**p*≤0.01;*** *p*≤0.001; **** *p*≤0.0001). The overall evaluation between the transfected parasite lineages, regardless of time point, is provided in the detailed table.

**Additional file 14: Supplementary Table S7**. *Trypanosoma rangeli* strains used in this study, showing their original hosts and geographical origins, as well as their distinct genetic classifications based on kDNA and ribosomal markers.

Footnote: *kDNA-based classification as proposed by Vallejo et al. (2002). **Ribosomal markers-based (SSU and ITS) classification as proposed by Maia da Silva et al. (2004). ND= Not determined.

**Additional file 15: Supplementary Table S8:** Primers used in PCR amplification and cloning of *Trypanosoma rangeli* and *Trypanosoma cruzi* mucins into the FlagGPI-pROCK vector and on assessment of genomic integration.

Footnote: ’F’ and ’R’ stand for Forward and Reverse primers, respectively. Restriction sites are highlighted in bold/italics at the 5’-end of each primer sequence. *= Amplification product generated with primer HX1 R for integration evaluation by PCR.

## ABBREVIATIONS

AB: Allele Balance
Bp: Base pairs
CCNV: Chromosome Copy Number Variation
CDS: Coding sequence
DGF-1: Disperse Gene Family 1
GO: Gene Onthology
GP: Glycoprotein
GPI: Glycosylphosphatidylinositol
Kb: Kilobases
Kbp: Thousand base pairs
kcal/mol: Kilocalories per mole
kDa: Kilodaltons
KMP: Kinetoplastid Membrane Protein
mAb: Monoclonal antibody
MASP: Mucin-Associated Surface Proteins
Mb: Megabase
Mbp: Million base-pairs
pTM: Predicted template modelling
RDC: Read Depth Coverage
RHS: Retrotransposon Hot Spot
RMSD: Root Mean Square Deviation
RNAi: RNA interference machinery
SNP: Single Nucleotide Polymorphism
TASV: Trypomastigote, Alanine, Serine and Valine rich protein glycoprotein family
VSG: Variable Surface Glycoprotein.

## ACKNOWLEDGMENTS

The authors would like to thank the Laboratório de Estudos em Biologia (LAMEB/UFSC, Brazil) and the Science for Life Laboratory (KI, Stockholm) for technical support. Authors are indebted to Dr Ariel Silber (USP, Brazil) for critical reading and valuable comments on this manuscript.

## AUTHOR’S CONTRIBUTIONS

GAM, GMM and ACS: Equally contributed to Formal analysis, Investigation, Methodology, Validation, Visualization and Writing – original draft.

CAP, CLMP, DDL, RAC, RSM and TKSP: Formal analysis, Investigation, Methodology and Validation. ASR: Formal analysis and Investigation.

TCB and GRM: Formal analysis, Investigation, Methodology, Validation and Writing – original draft.

JFSF, MS: Data curation, Formal analysis, Investigation, Resources, Supervision, Validation and Writing – original draft.

PHS and GW: Data curation, Formal analysis, Investigation, Methodology, Validation, Resources, Supervision and Writing – original draft.

BA and ECG: Conceptualization, Data curation, Formal analysis, Funding acquisition, Investigation, Project administration, Resources, Supervision, Validation and Writing – original draft.

## AVAILABILITY OF DATA AND MATERIALS

Data generated during the current study is available in the GenBank database (Bioproject PRJNA1380247). Further datasets or subsets used and/or analysed during the current study are available from the corresponding authors on reasonable request. DNA transfer from UFSC to KI was carried out exclusively for sequencing purposes and was registered in the Brazilian National System for the Management of Genetic Heritage and Associated Traditional Knowledge (SISGEN A206000).

## ETHICS APPROVAL AND CONSENT TO PARTICIPATE

Not applicable.

## CONSENT FOR PUBLICATION

Not applicable.

