## Supplementary figures and images for "Comprehensive genomic analysis of *Trypanosoma rangeli* reveals key insights into the biology and evolution of this non-virulent American mammalian trypanosome"

### Supplementary figure S1

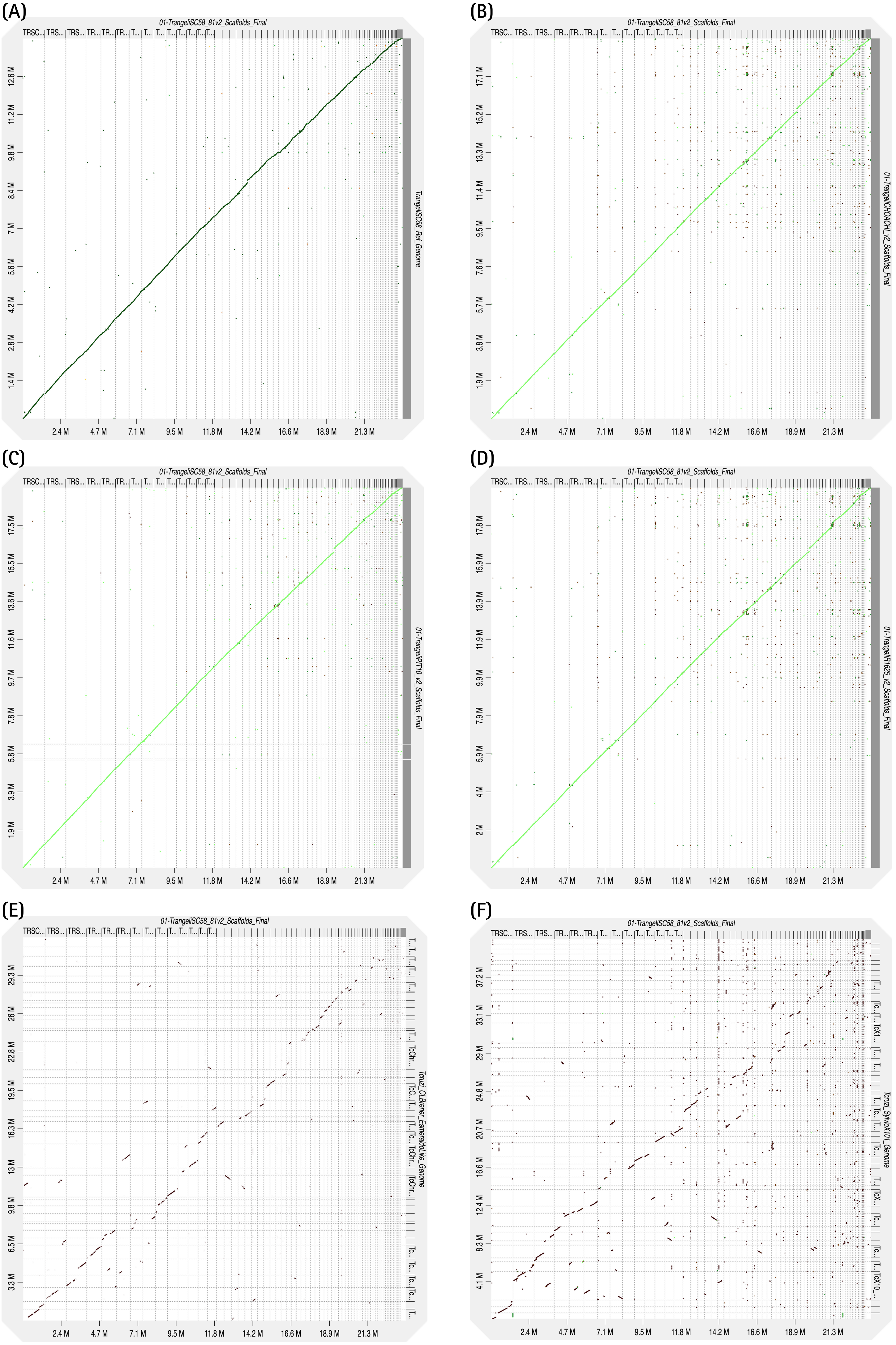

### Supplementary figure S2

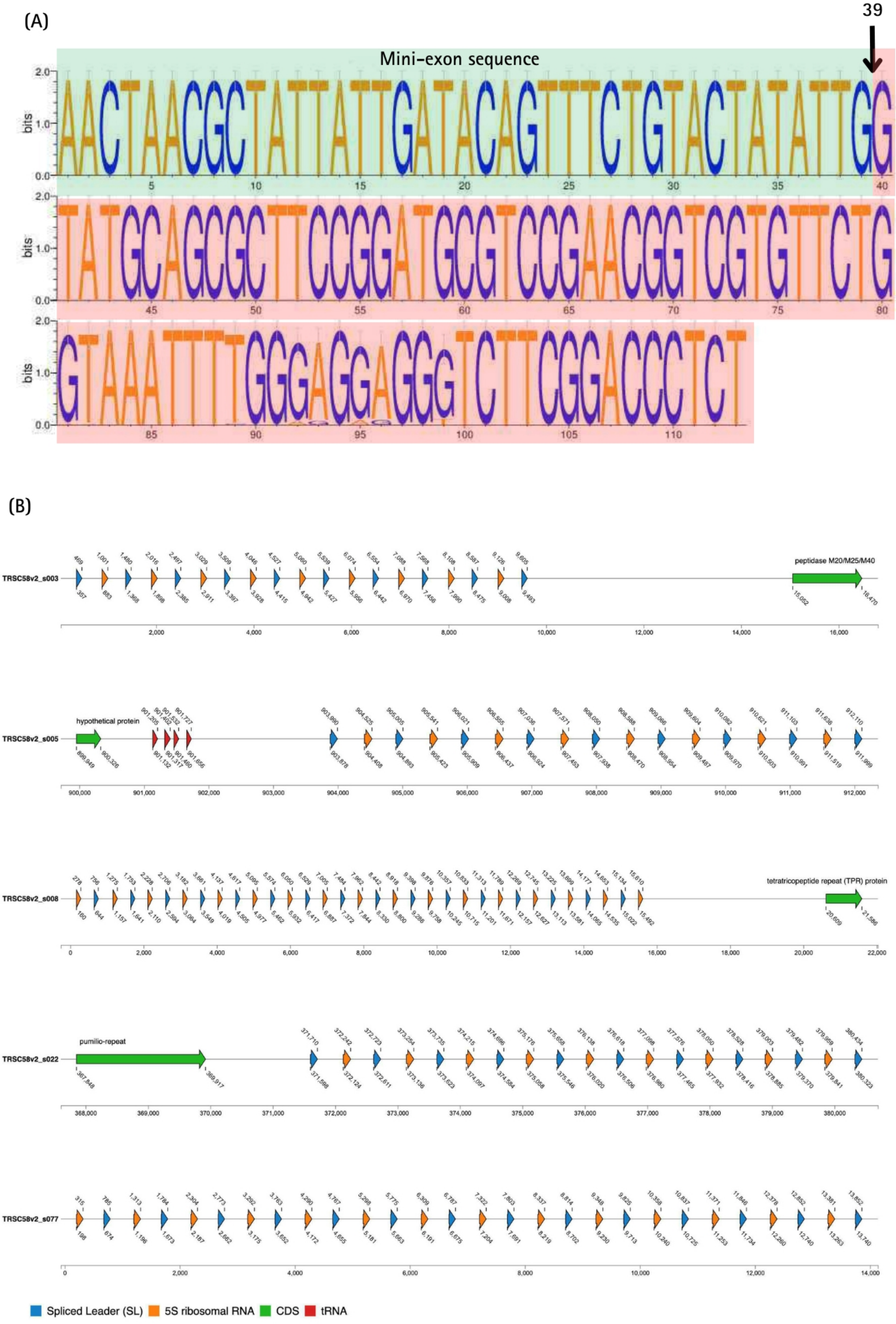

### Supplementary figure S4

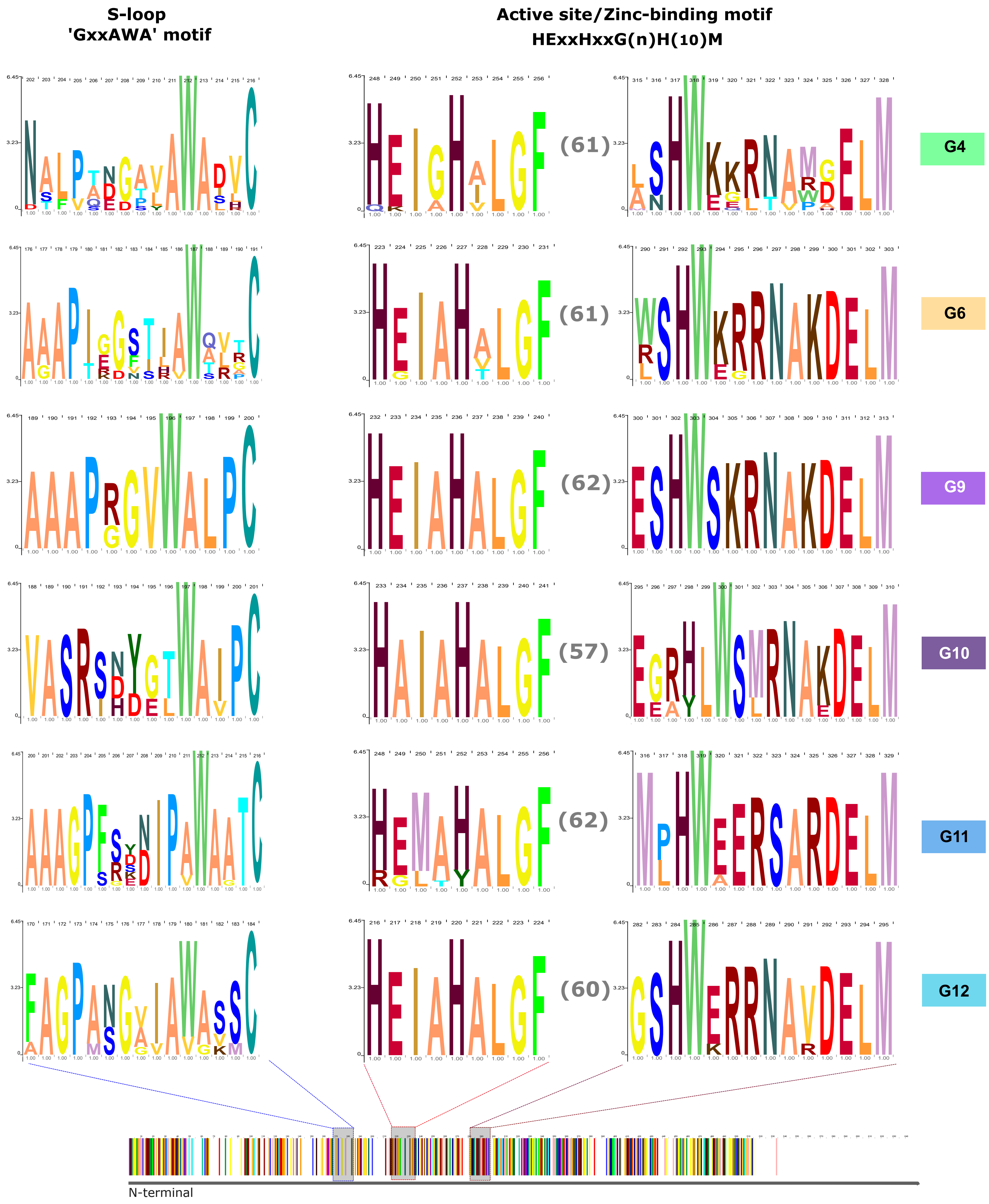

### Supplementary figure S5

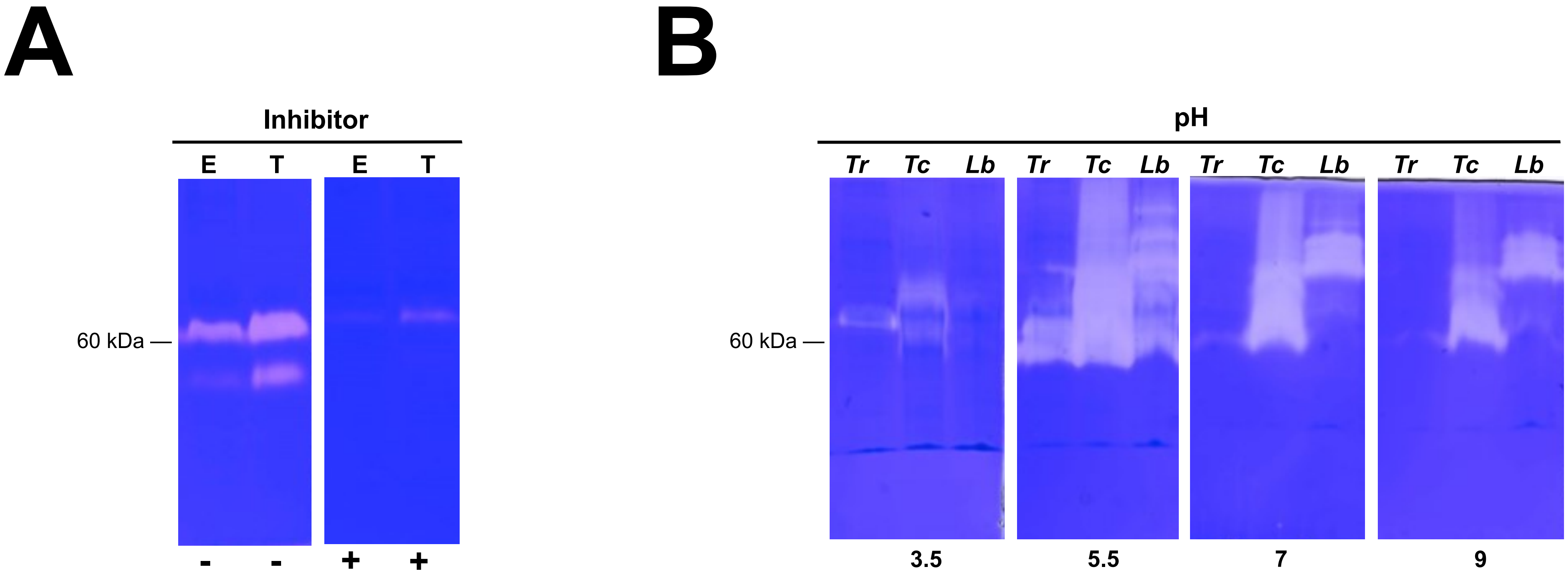

### Supplementary figure S6

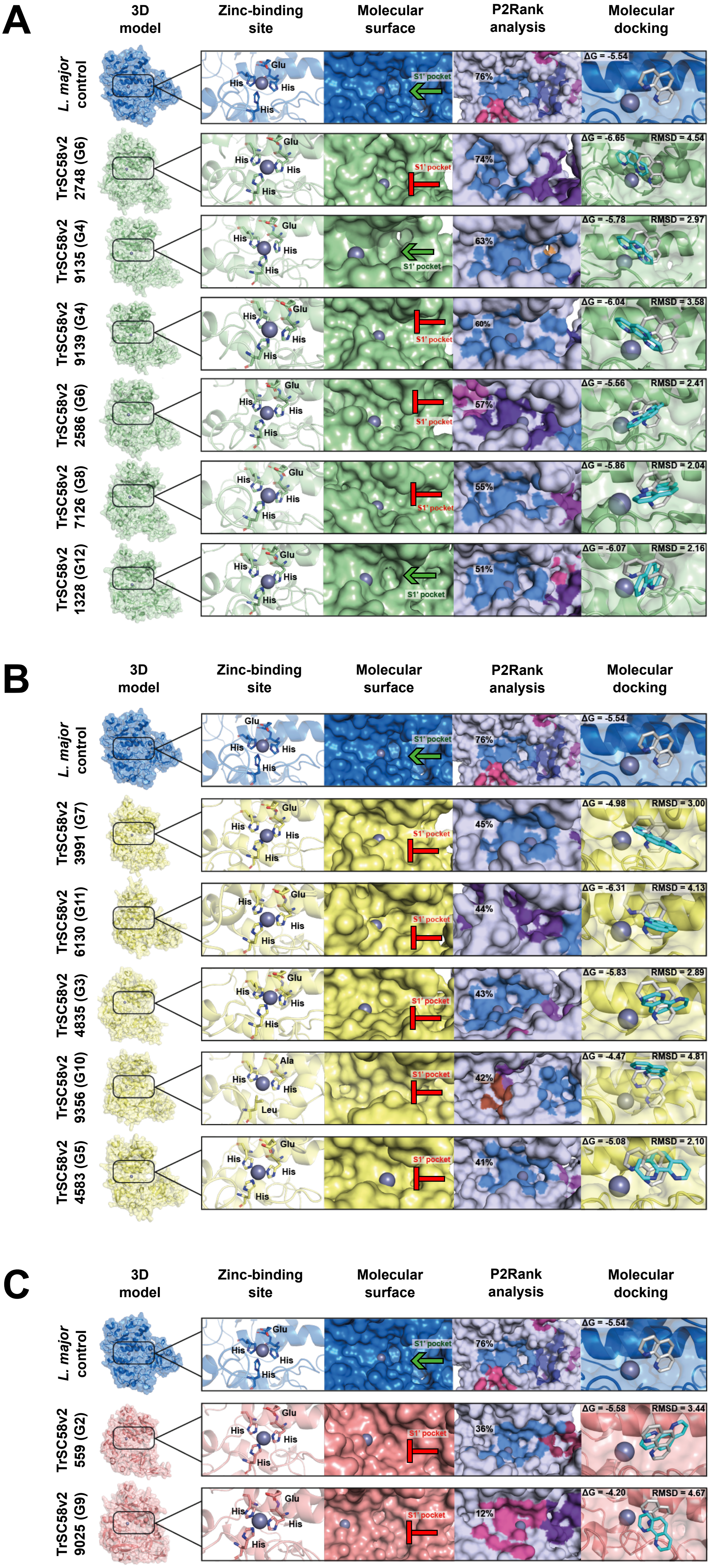
