## Supplementary figure S3 for "Comprehensive genomic analysis of *Trypanosoma rangeli* reveals key insights into the biology and evolution of this non-virulent American mammalian trypanosome"

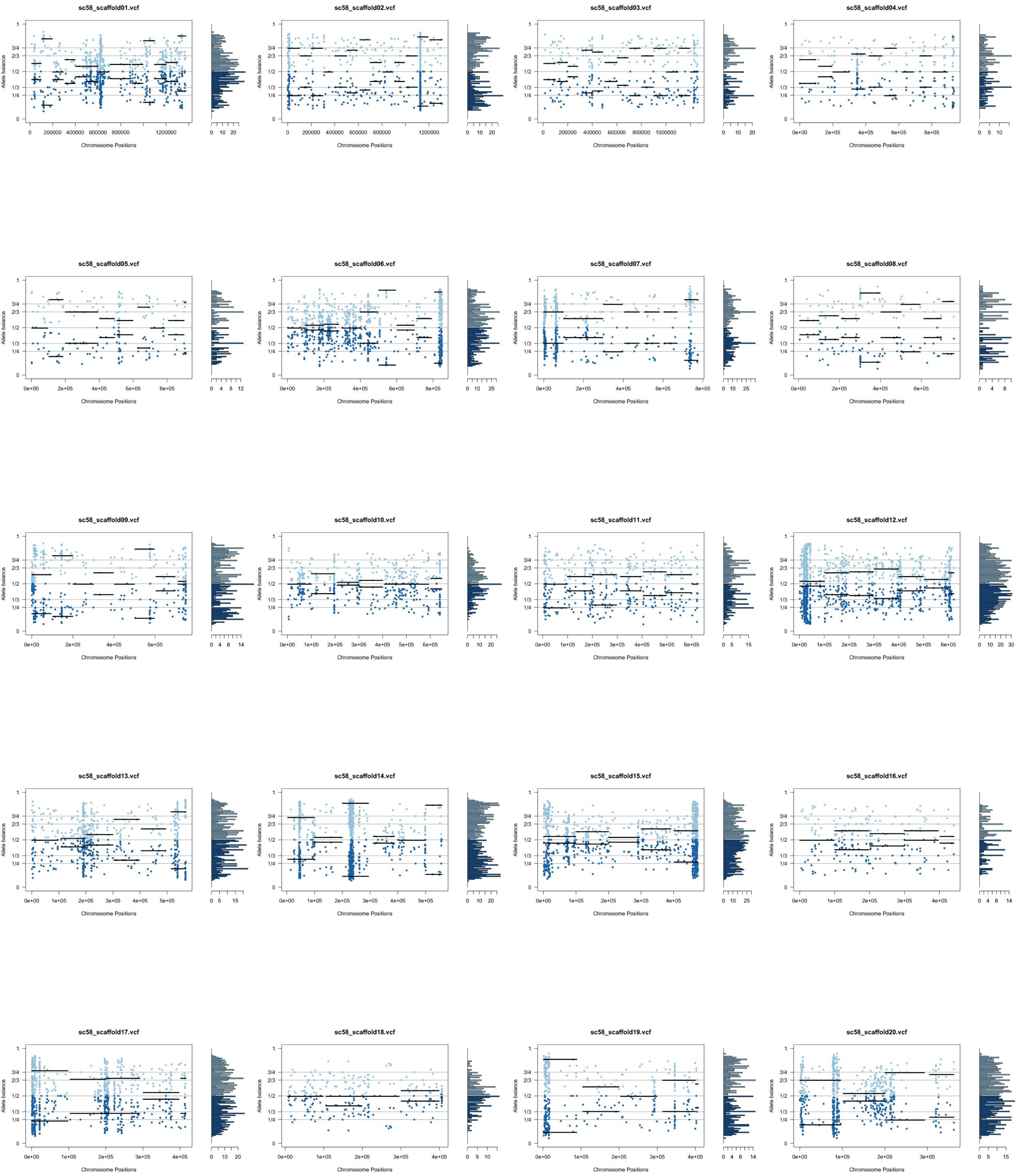

sc58_scaffold01.vcf
sc58_scaffold02.vcf
sc58_scaffold03.vcf
sc58_scaffold04.vcf
sc58_scaffold05.vcf
sc58_scaffold06.vcf
sc58_scaffold07.vcf
sc58_scaffold08.vcf
sc58_scaffold09.vcf
sc58_scaffold10.vcf
sc58_scaffold11.vcf
sc58_scaffold12.vcf
sc58_scaffold13.vcf
sc58_scaffold14.vcf
sc58_scaffold15.vcf
sc58_scaffold16.vcf
sc58_scaffold17.vcf
sc58_scaffold18.vcf
sc58_scaffold19.vcf
sc58_scaffold20.vcf
Allele balance
Chromosome Positions

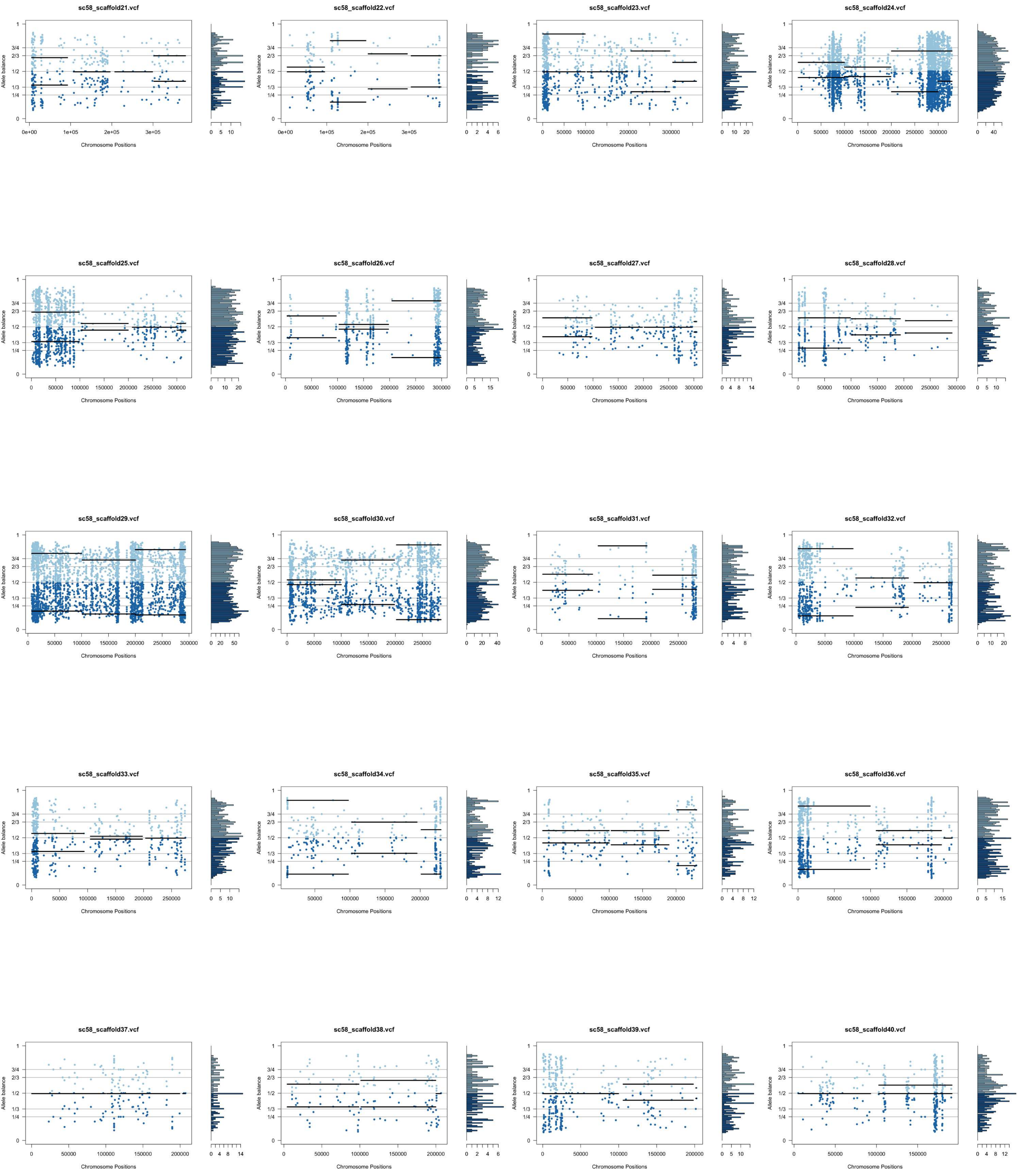
sc58_scaffold21.vcf
sc58_scaffold22.vcf
sc58_scaffold23.vcf
sc58_scaffold24.vcf
sc58_scaffold25.vcf
sc58_scaffold26.vcf
sc58_scaffold27.vcf
sc58_scaffold28.vcf
sc58_scaffold29.vcf
sc58_scaffold30.vcf
sc58_scaffold31.vcf
sc58_scaffold32.vcf
sc58_scaffold33.vcf
sc58_scaffold34.vcf
sc58_scaffold35.vcf
sc58_scaffold36.vcf
sc58_scaffold37.vcf
sc58_scaffold38.vcf
sc58_scaffold39.vcf
sc58_scaffold40.vcf
Allele balance
Chromosome Positions
1
3/4
2/3
1/2
1/3
1/4
0

sc58_scaffold41.vcf

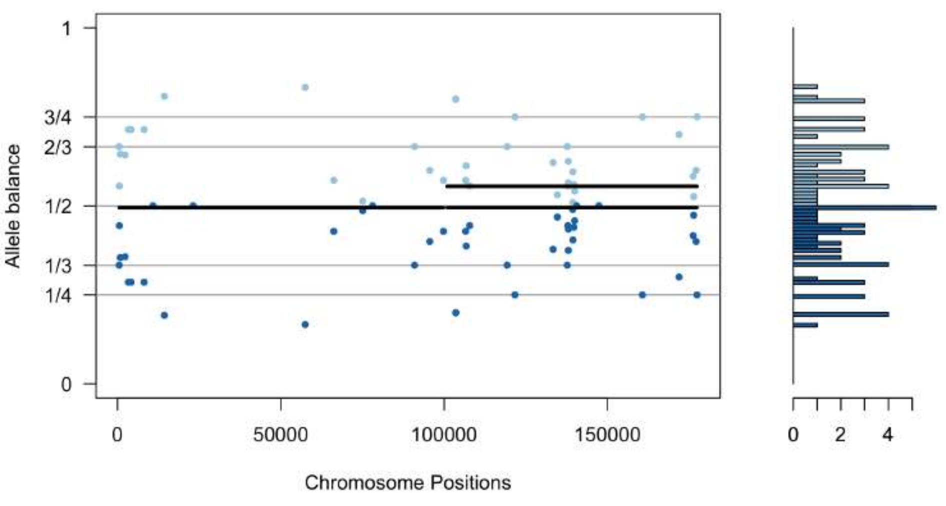


sc58_scaffold42.vcf

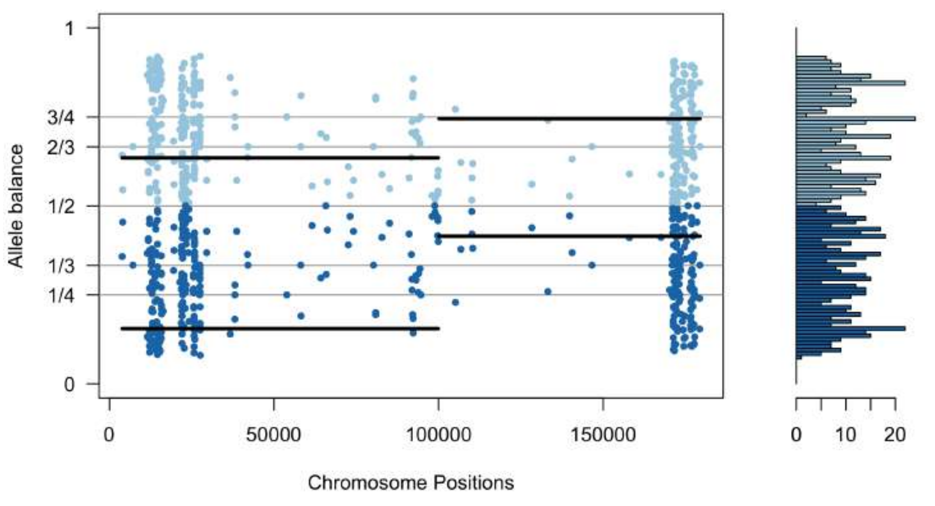


sc58_scaffold43.vcf

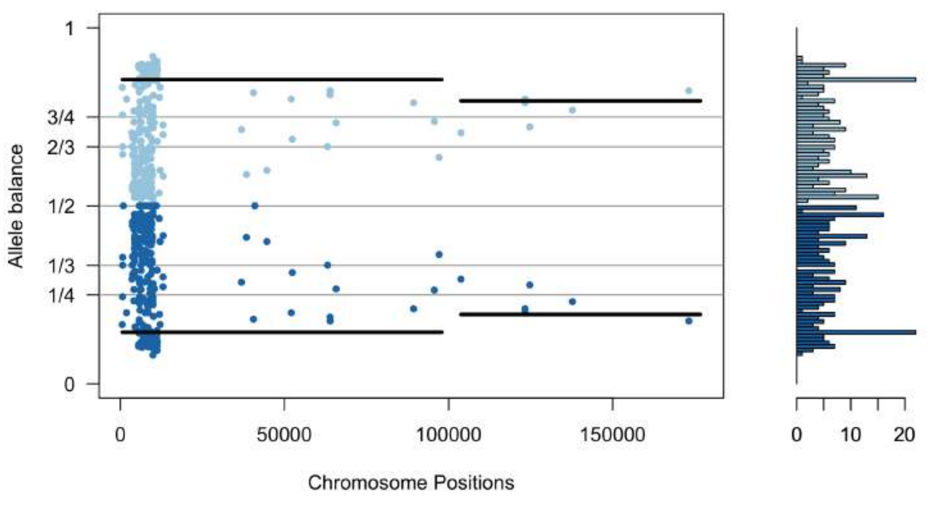


sc58_scaffold44.vcf

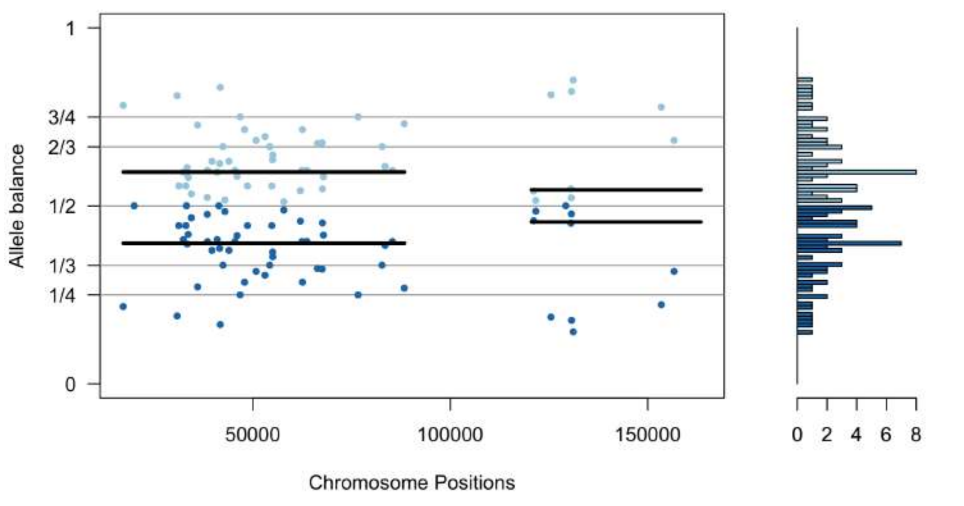


sc58_scaffold45.vcf

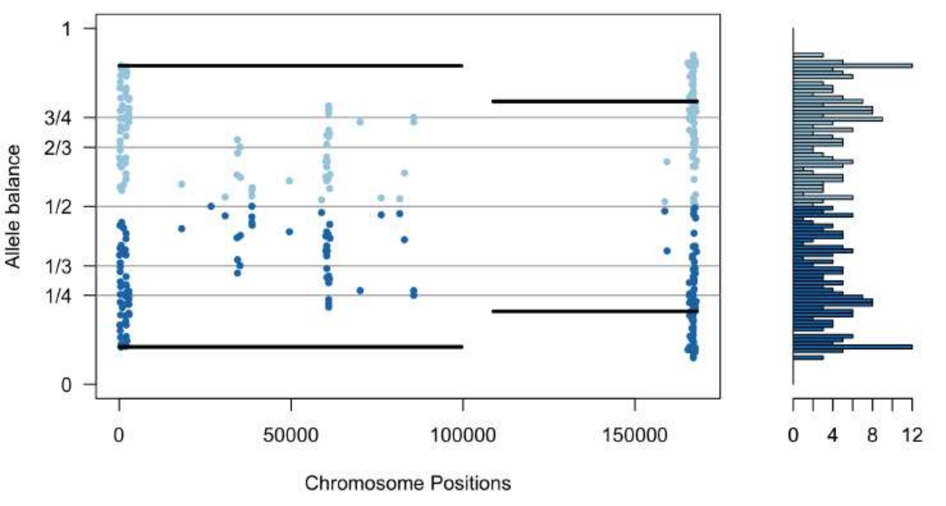


sc58_scaffold46.vcf

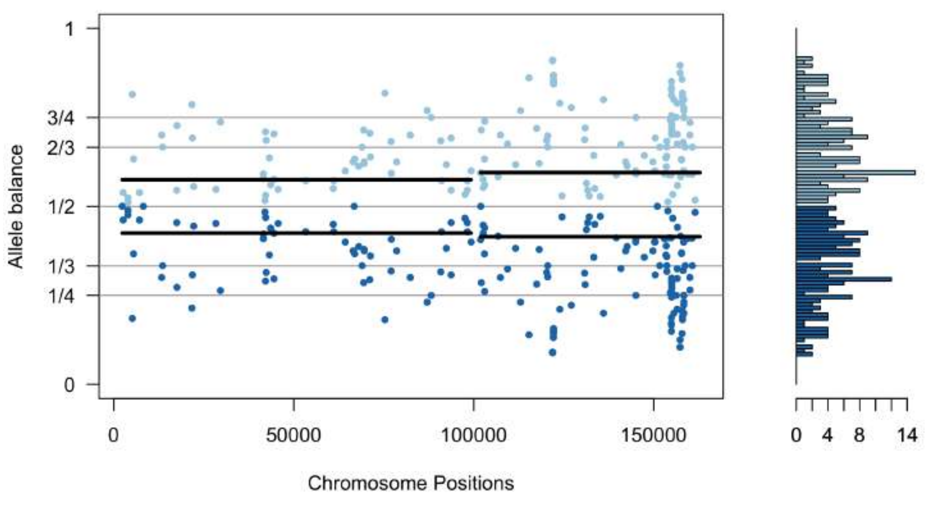


sc58_scaffold47.vcf

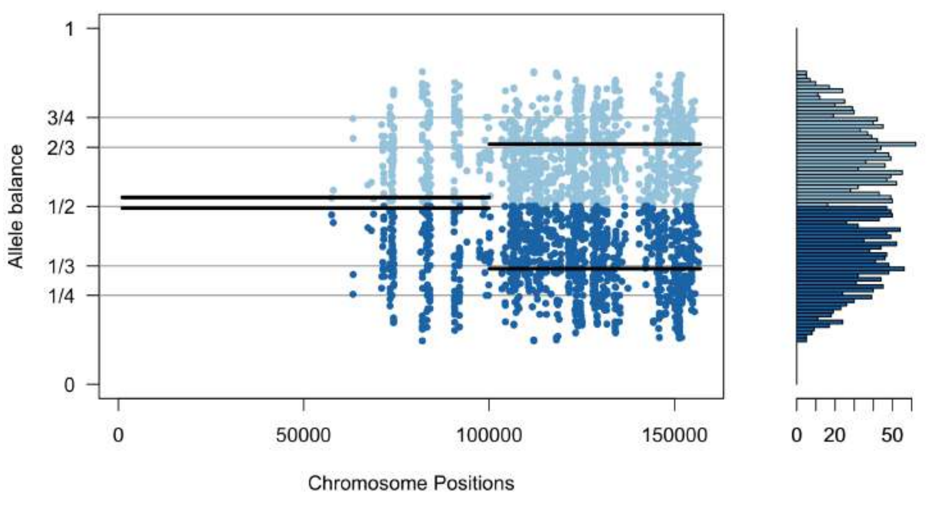


sc58_scaffold48.vcf

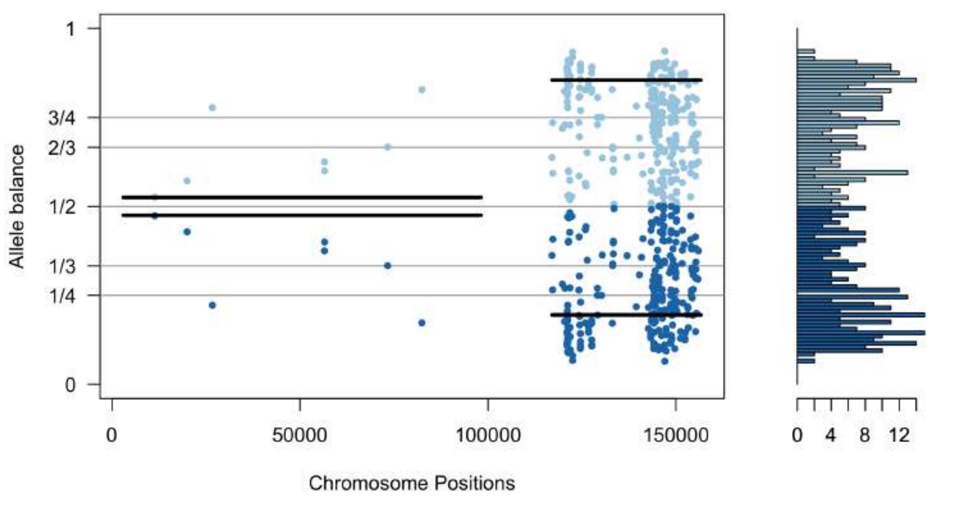


sc58_scaffold49.vcf

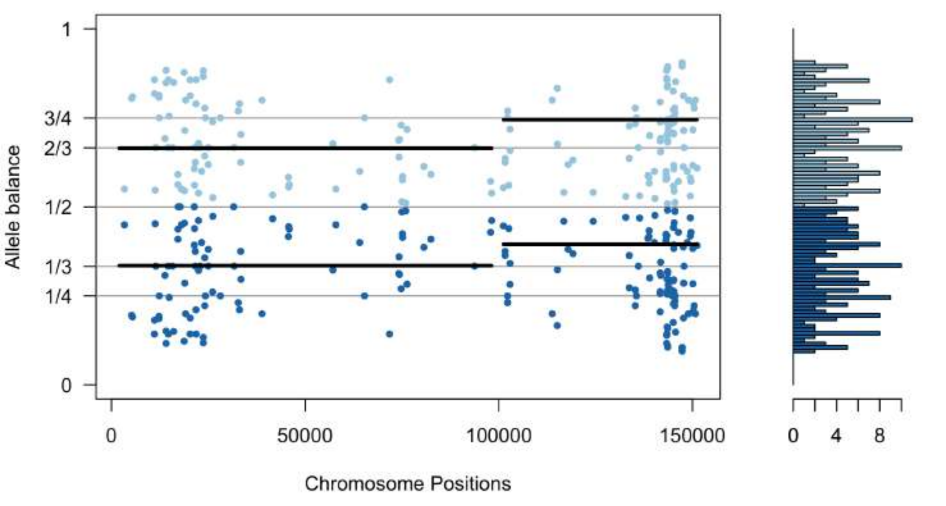


sc58_scaffold50.vcf

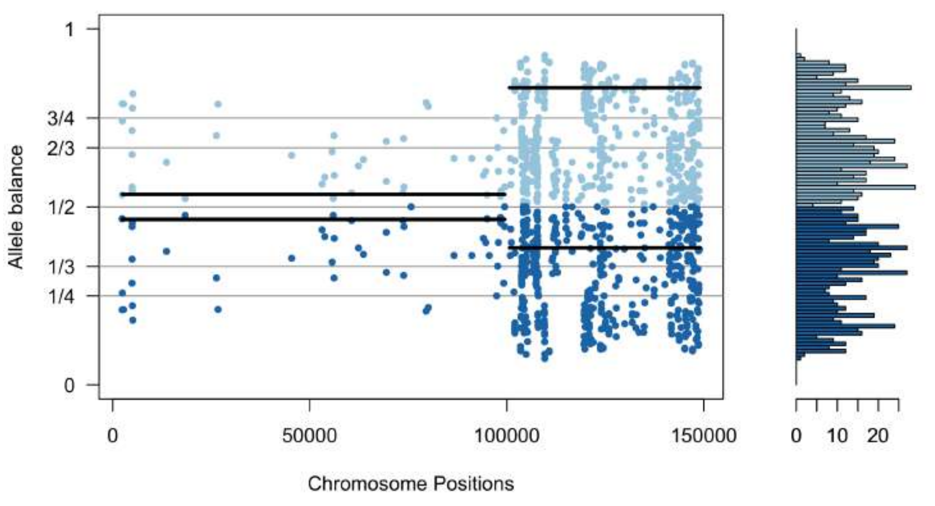


sc58_scaffold51.vcf

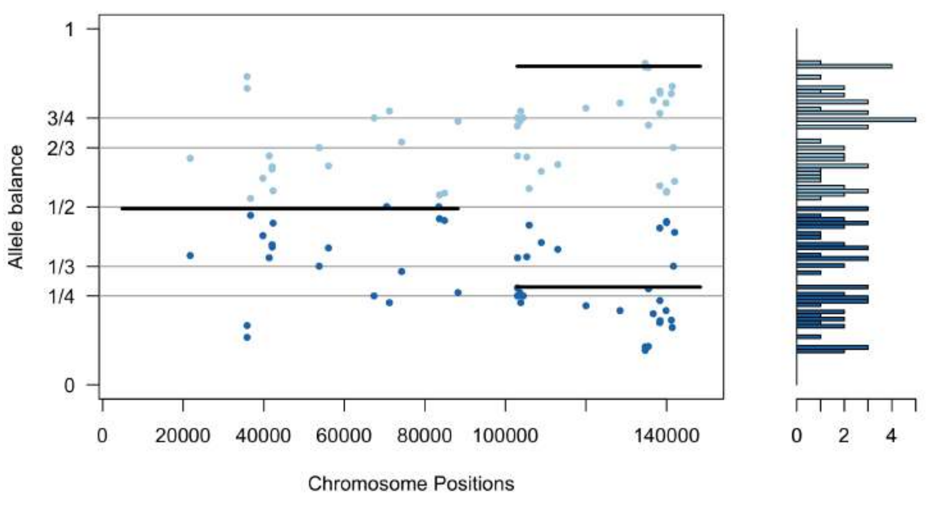


sc58_scaffold52.vcf

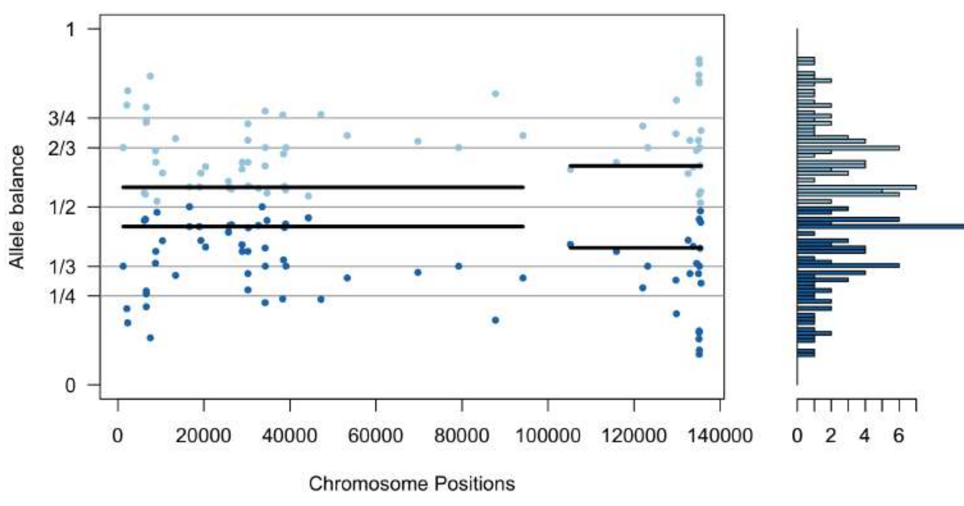


sc58_scaffold53.vcf

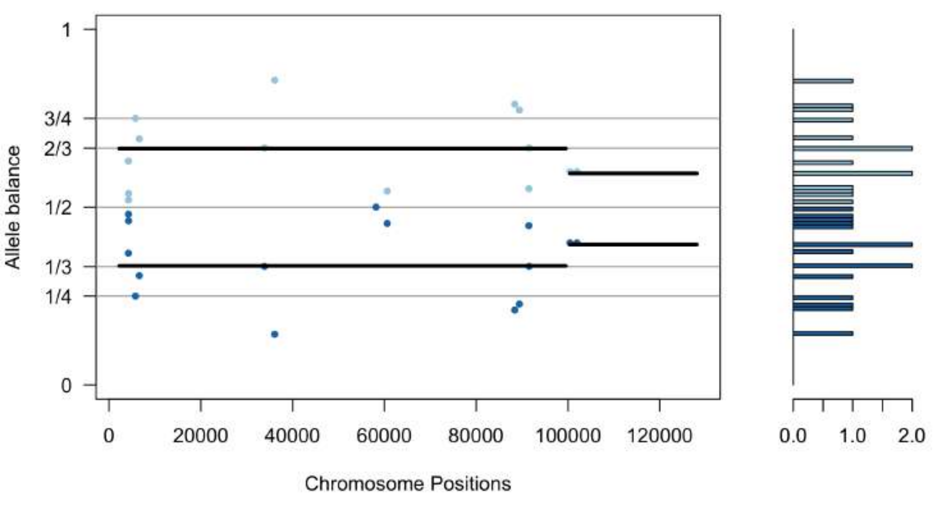


sc58_scaffold54.vcf

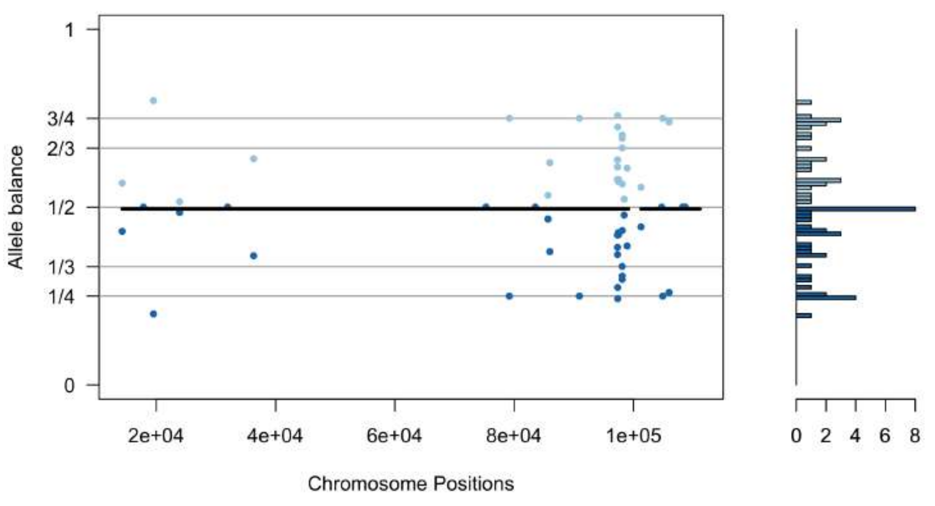


sc58_scaffold55.vcf

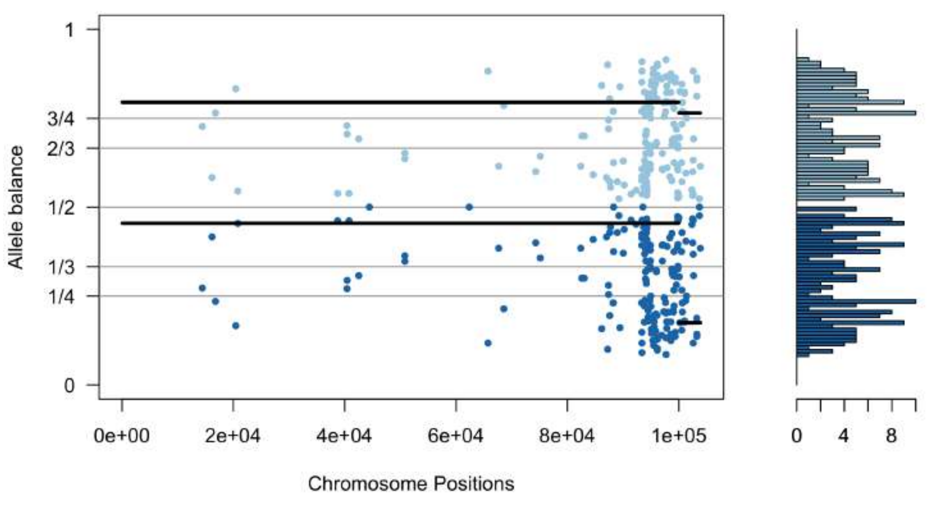


sc58_scaffold56.vcf

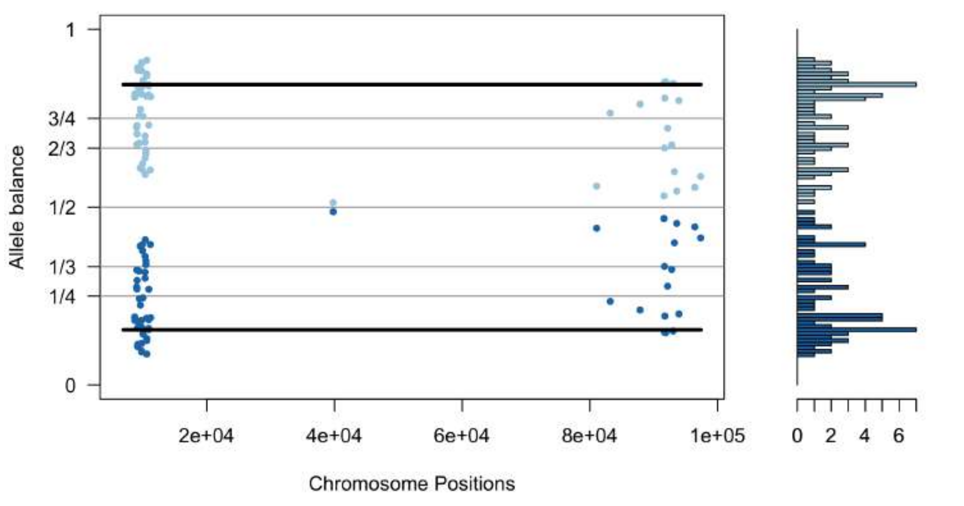


sc58_scaffold57.vcf

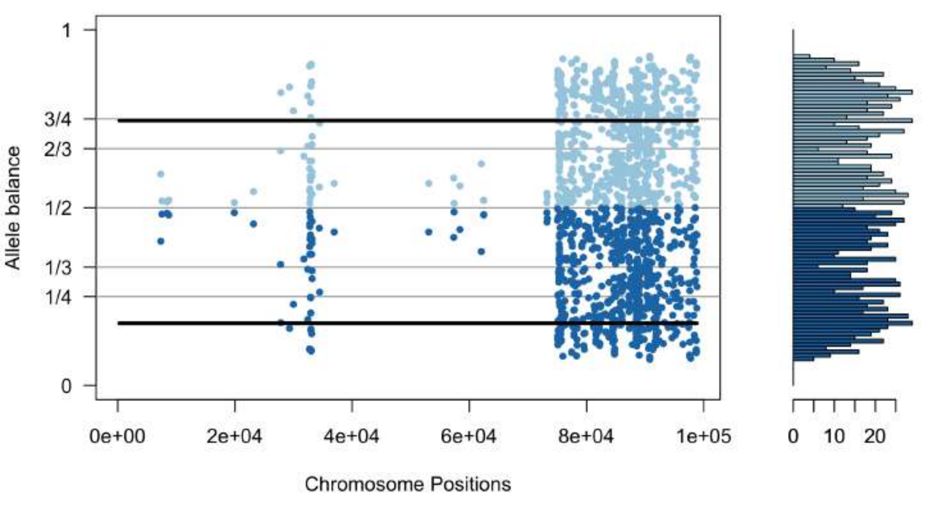


sc58_scaffold58.vcf

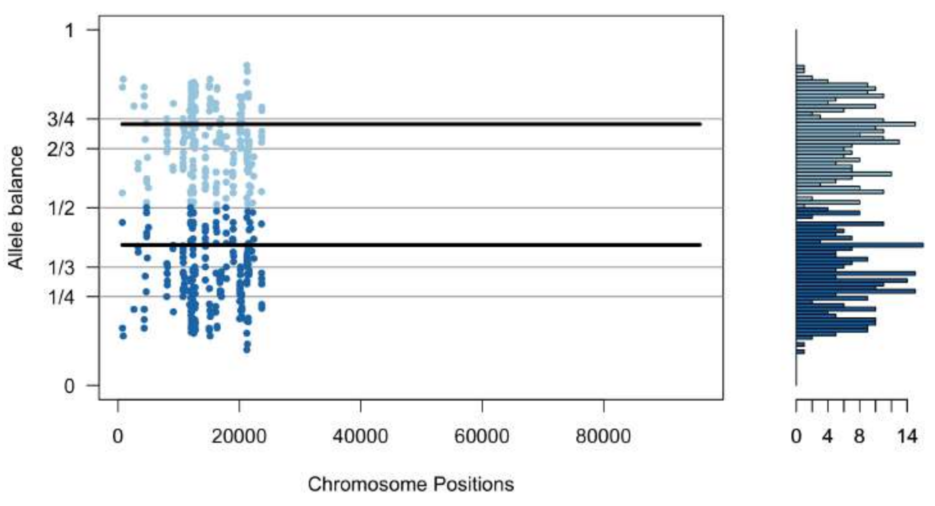


sc58_scaffold59.vcf

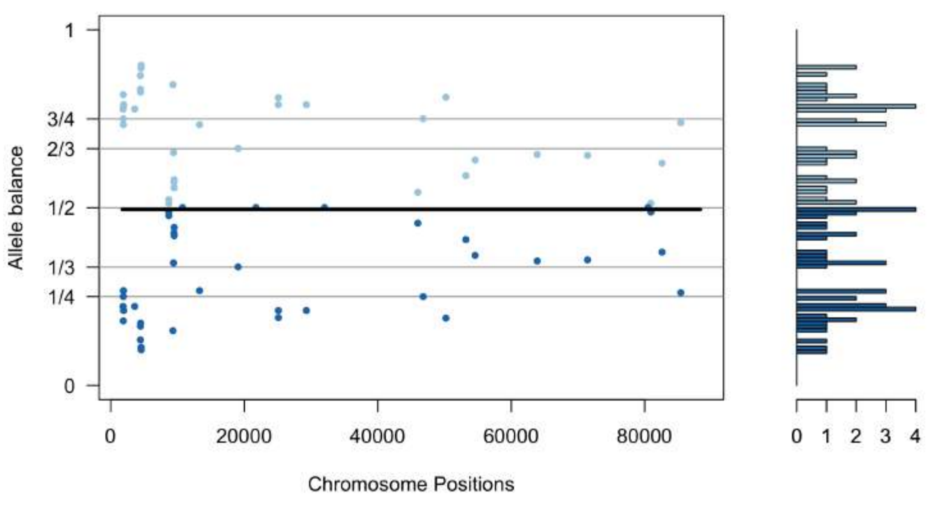


sc58_scaffold60.vcf

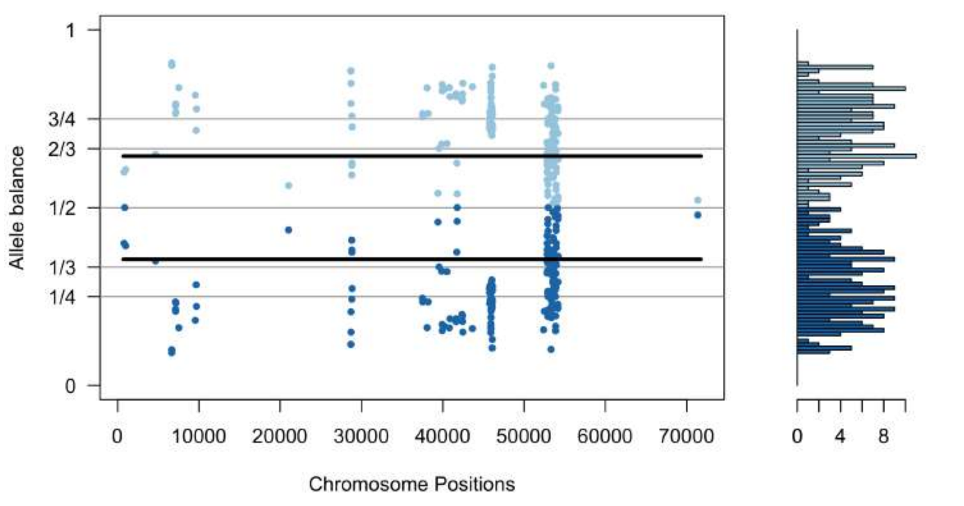

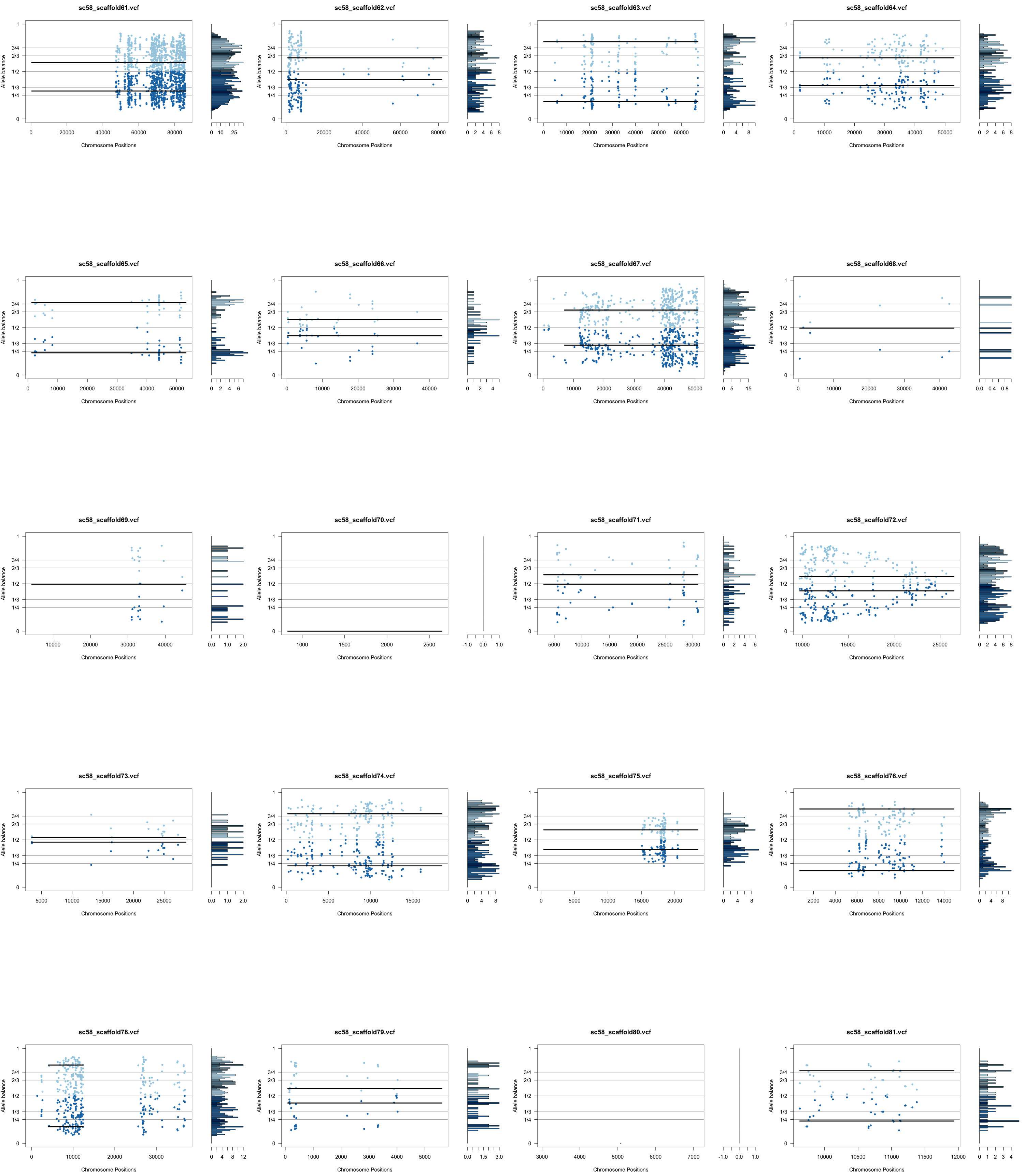
sc58_scaffold61.vcf
sc58_scaffold62.vcf
sc58_scaffold63.vcf
sc58_scaffold64.vcf
sc58_scaffold65.vcf
sc58_scaffold66.vcf
sc58_scaffold67.vcf
sc58_scaffold68.vcf
sc58_scaffold69.vcf
sc58_scaffold70.vcf
sc58_scaffold71.vcf
sc58_scaffold72.vcf
sc58_scaffold73.vcf
sc58_scaffold74.vcf
sc58_scaffold75.vcf
sc58_scaffold76.vcf
sc58_scaffold78.vcf
sc58_scaffold79.vcf
sc58_scaffold80.vcf
sc58_scaffold81.vcf
Allele balance
Chromosome Positions

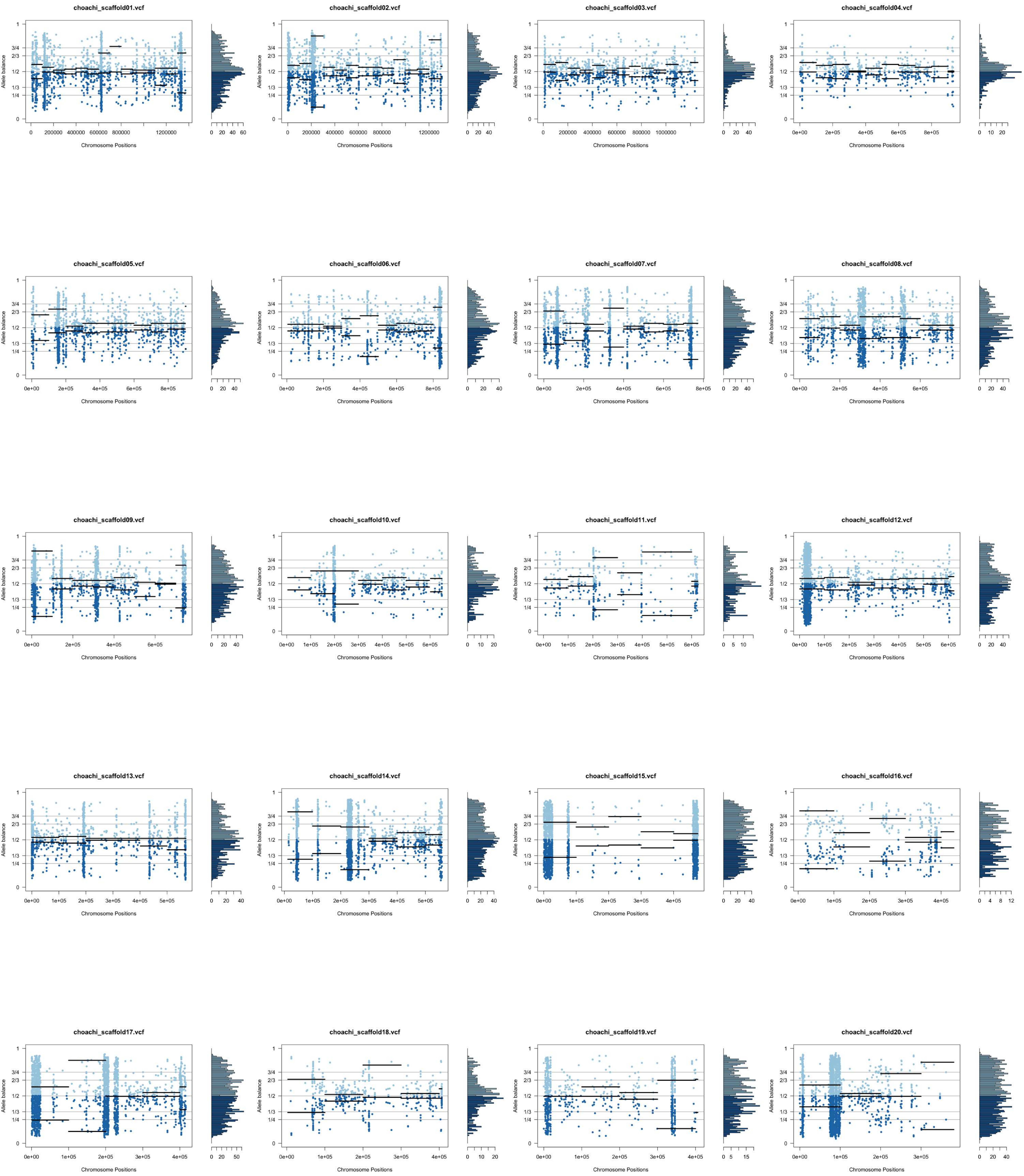
choachi_scaffold01.vcf
choachi_scaffold02.vcf
choachi_scaffold03.vcf
choachi_scaffold04.vcf
choachi_scaffold05.vcf
choachi_scaffold06.vcf
choachi_scaffold07.vcf
choachi_scaffold08.vcf
choachi_scaffold09.vcf
choachi_scaffold10.vcf
choachi_scaffold11.vcf
choachi_scaffold12.vcf
choachi_scaffold13.vcf
choachi_scaffold14.vcf
choachi_scaffold15.vcf
choachi_scaffold16.vcf
choachi_scaffold17.vcf
choachi_scaffold18.vcf
choachi_scaffold19.vcf
choachi_scaffold20.vcf
Allele balance
Chromosome Positions

**choachi_scaffold21.vcf**

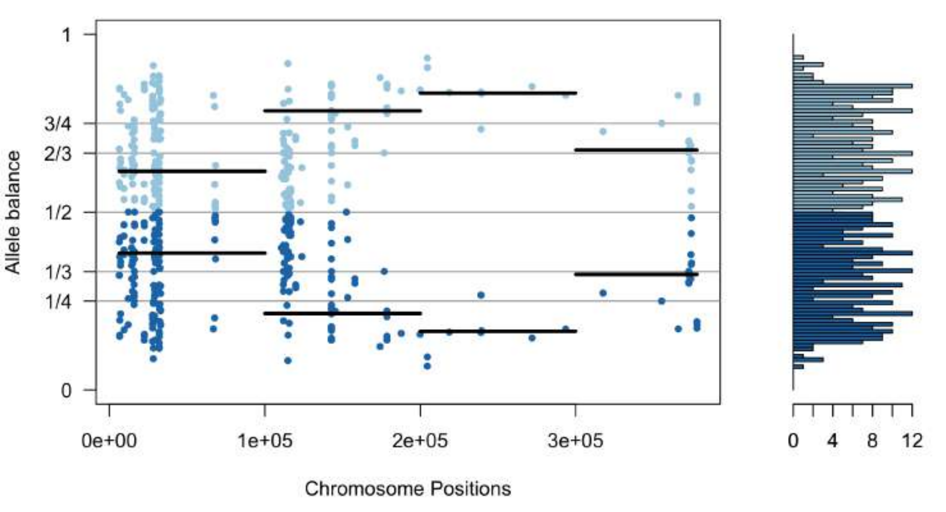


**choachi_scaffold22.vcf**

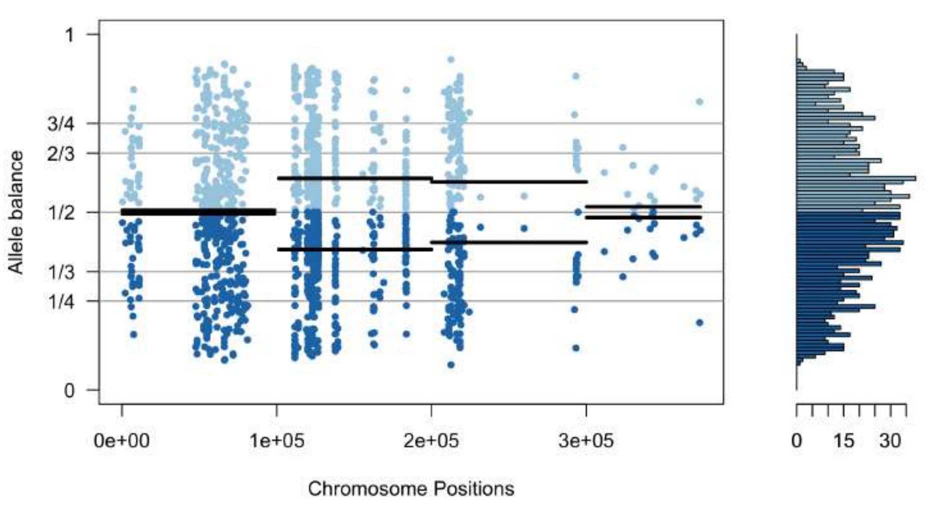


**choachi_scaffold23.vcf**

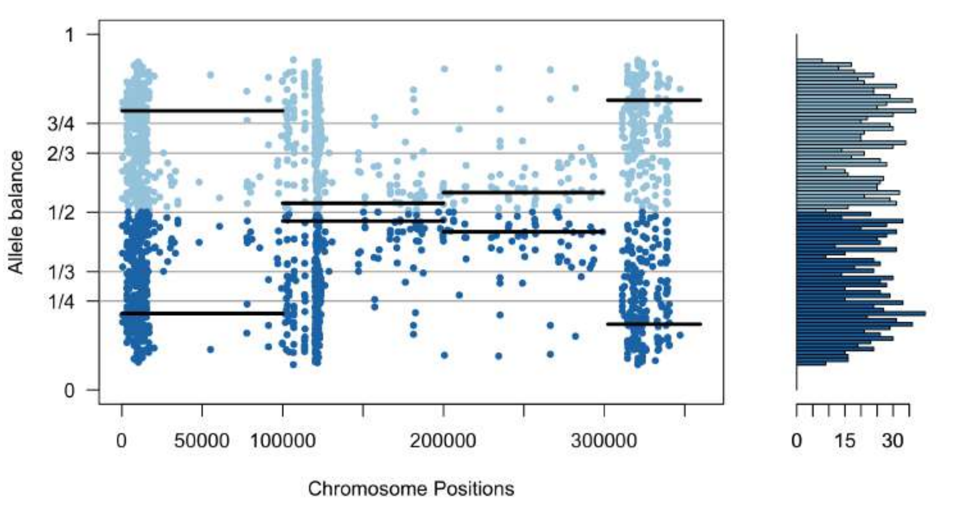


**choachi_scaffold24.vcf**

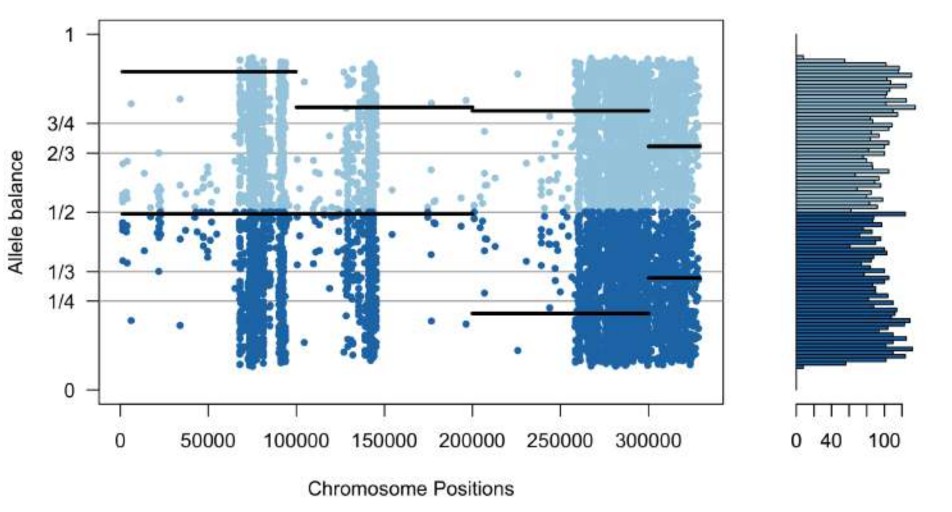


**choachi_scaffold25.vcf**

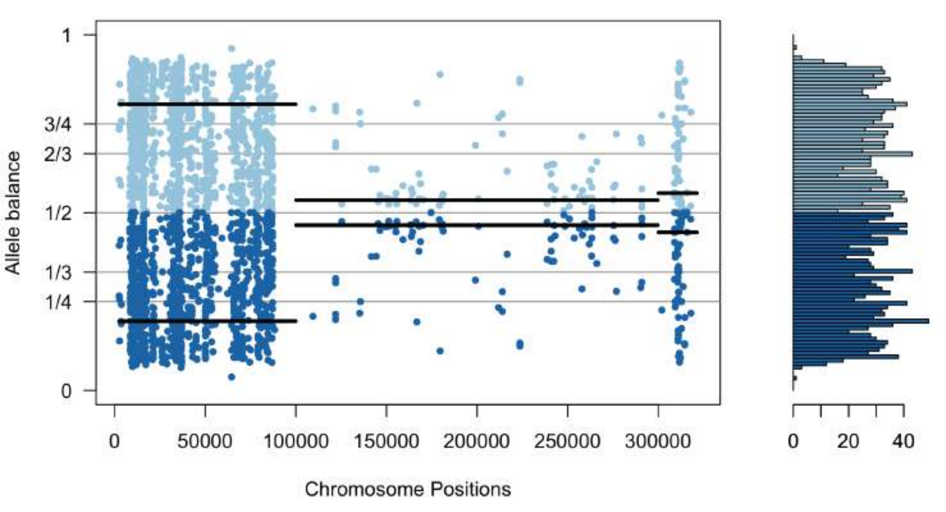


**choachi_scaffold26.vcf**

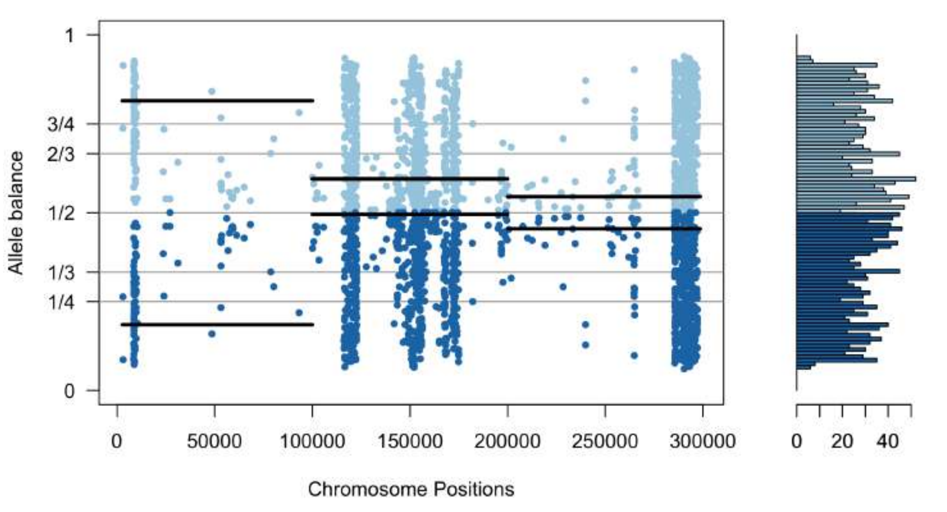


**choachi_scaffold27.vcf**

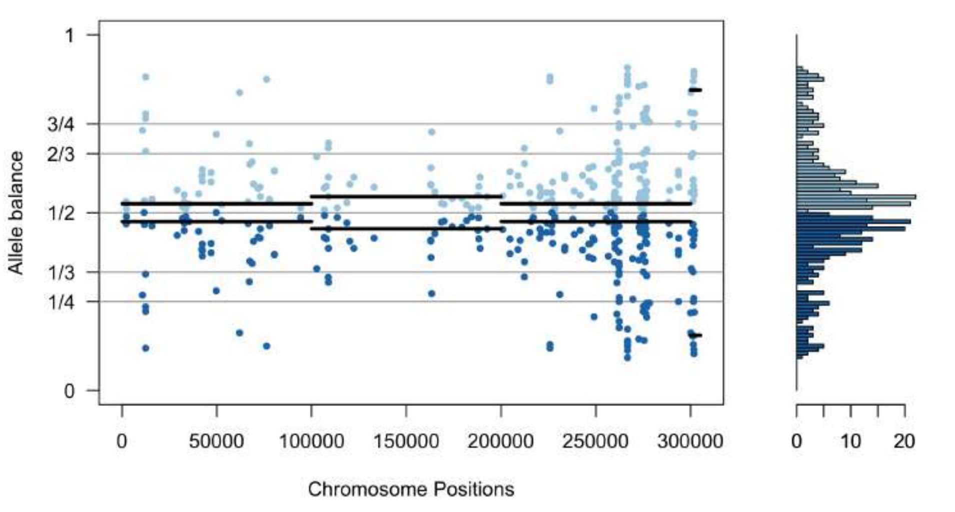


**choachi_scaffold28.vcf**

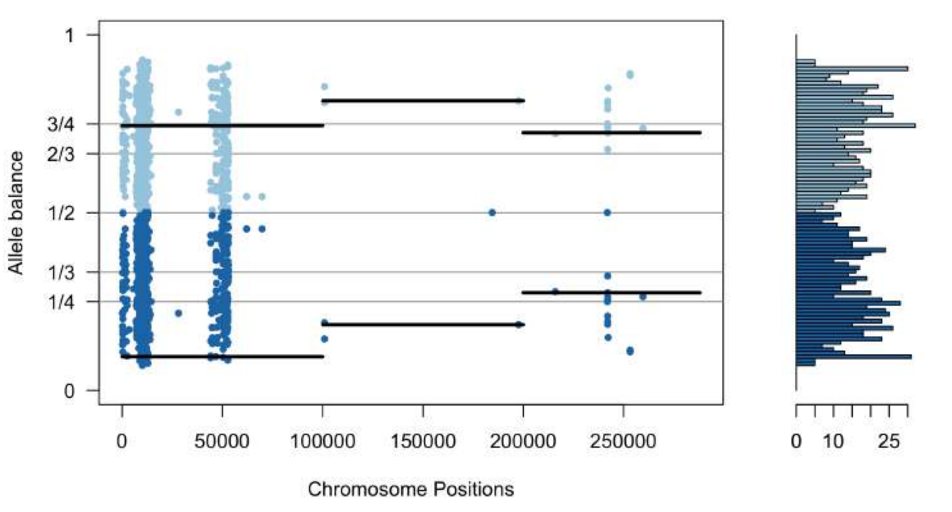


**choachi_scaffold29.vcf**

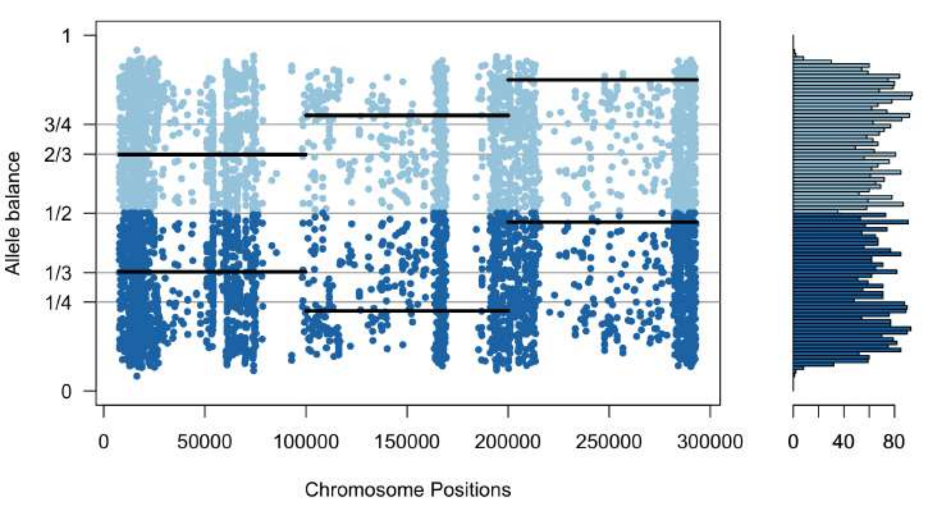


**choachi_scaffold30.vcf**

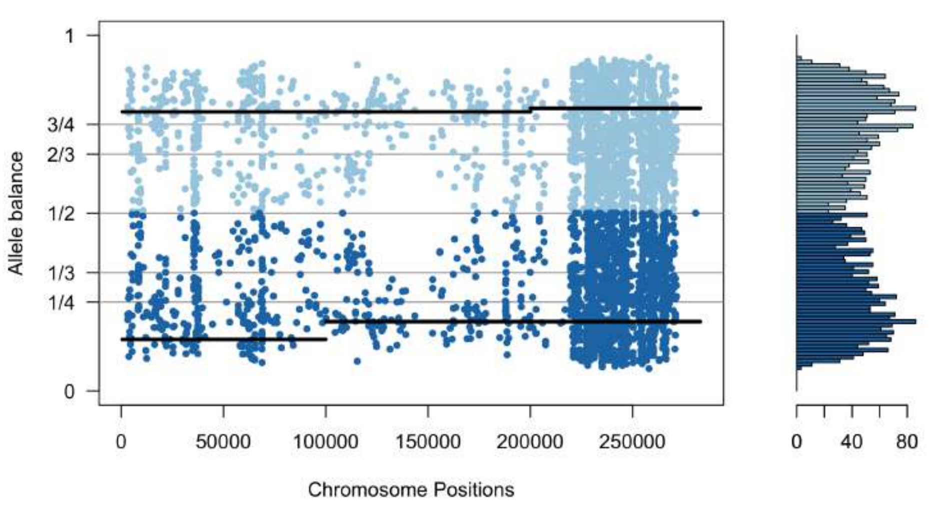


**choachi_scaffold31.vcf**

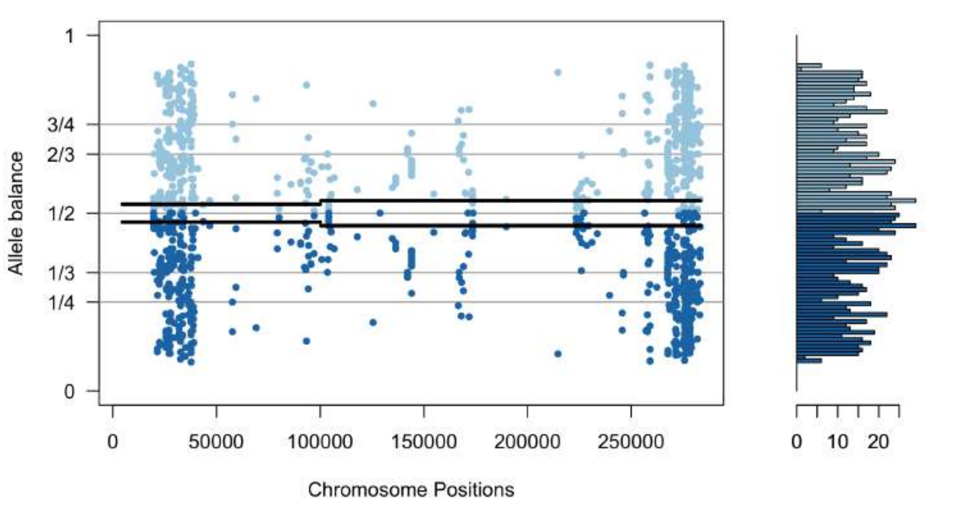


**choachi_scaffold32.vcf**

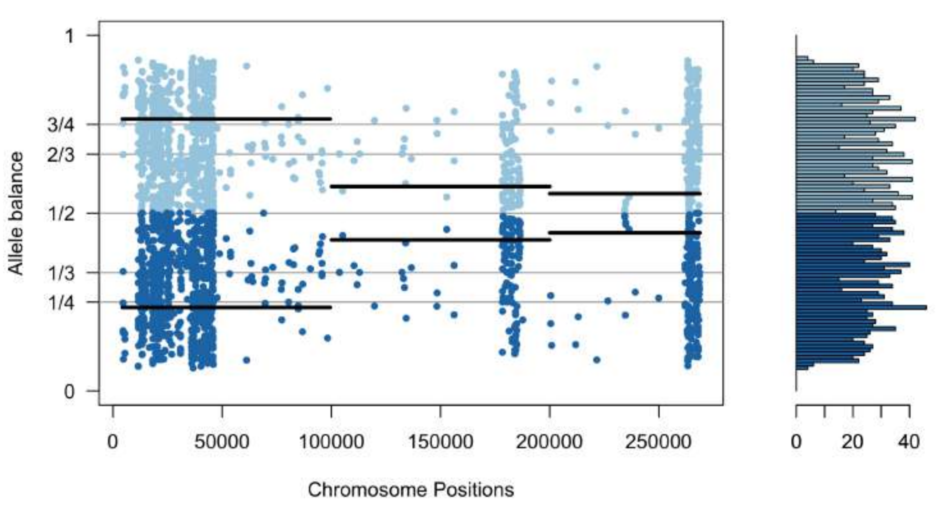


**choachi_scaffold33.vcf**

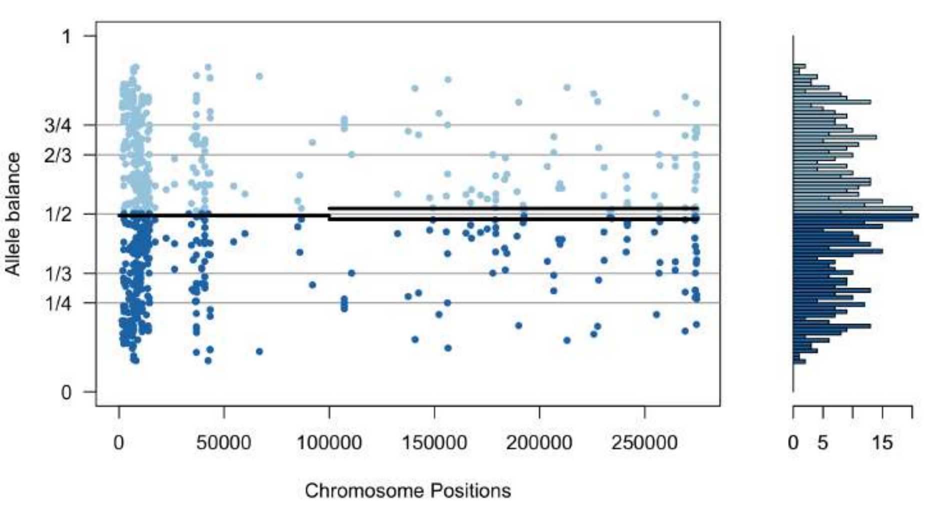


**choachi_scaffold34.vcf**

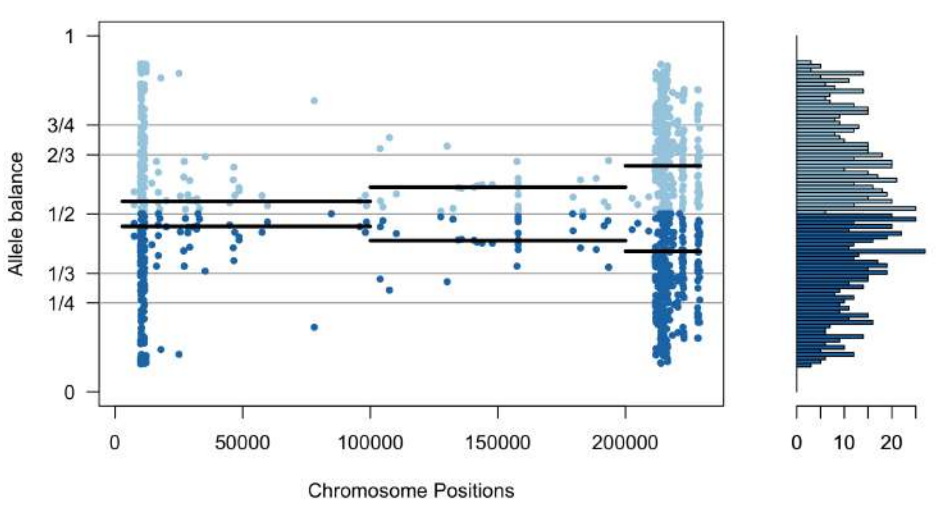


**choachi_scaffold35.vcf**

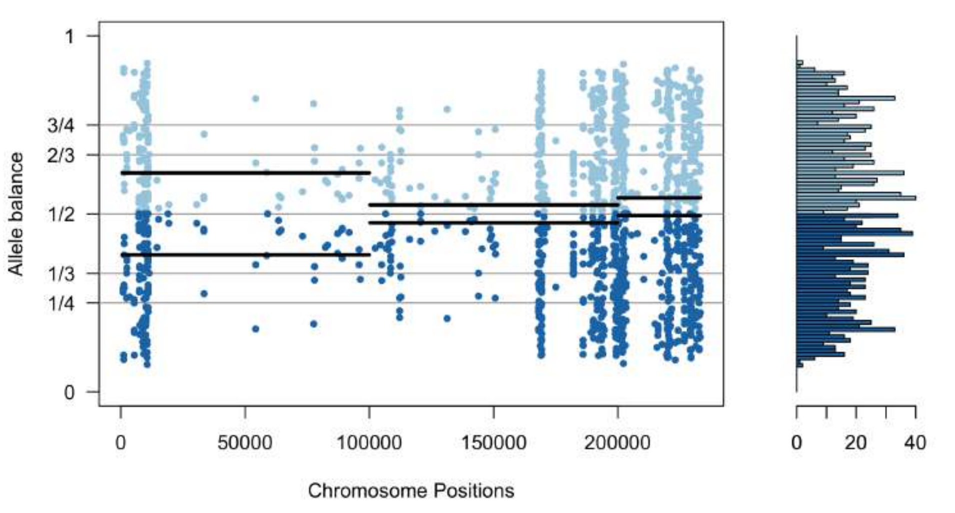


**choachi_scaffold36.vcf**

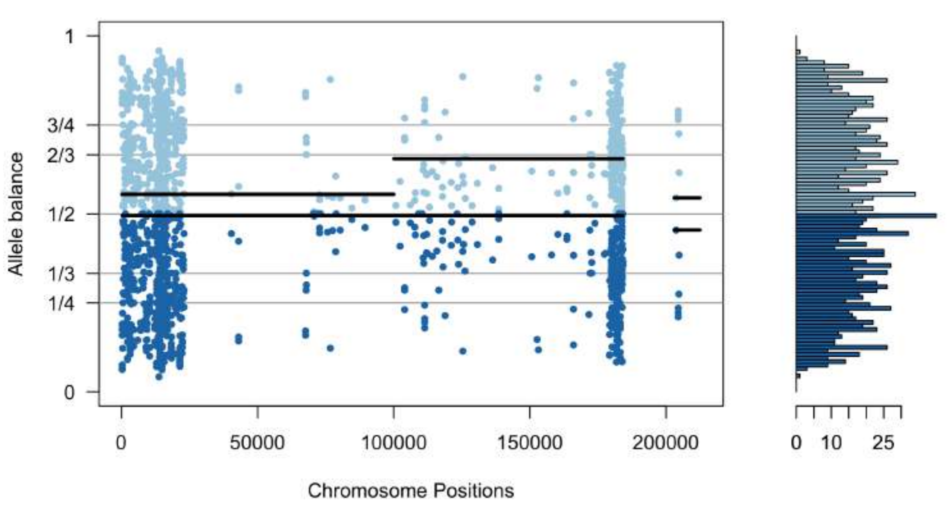


**choachi_scaffold37.vcf**

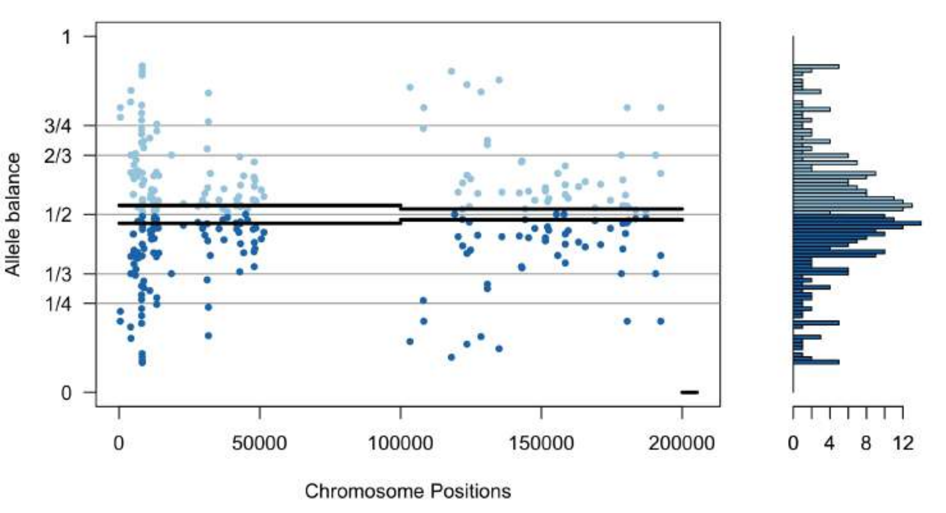


**choachi_scaffold38.vcf**

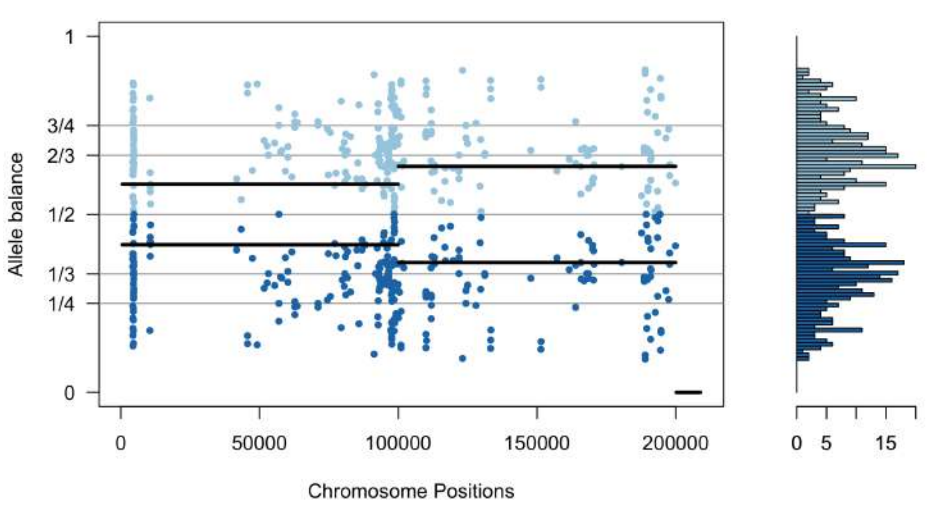


**choachi_scaffold39.vcf**

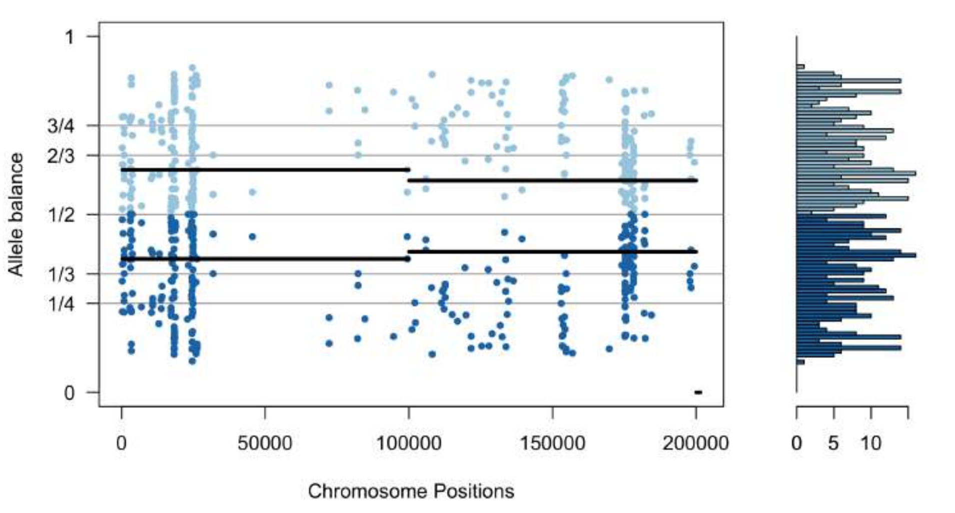


**choachi_scaffold40.vcf**

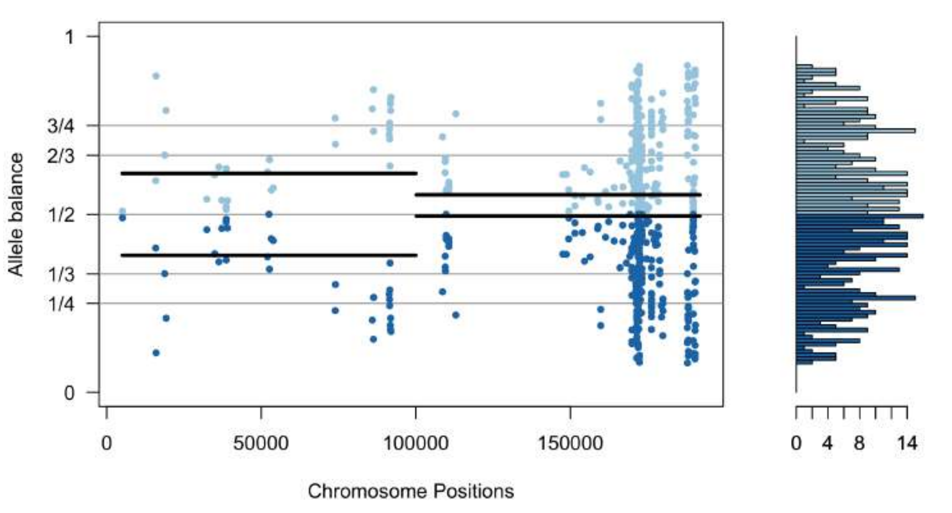

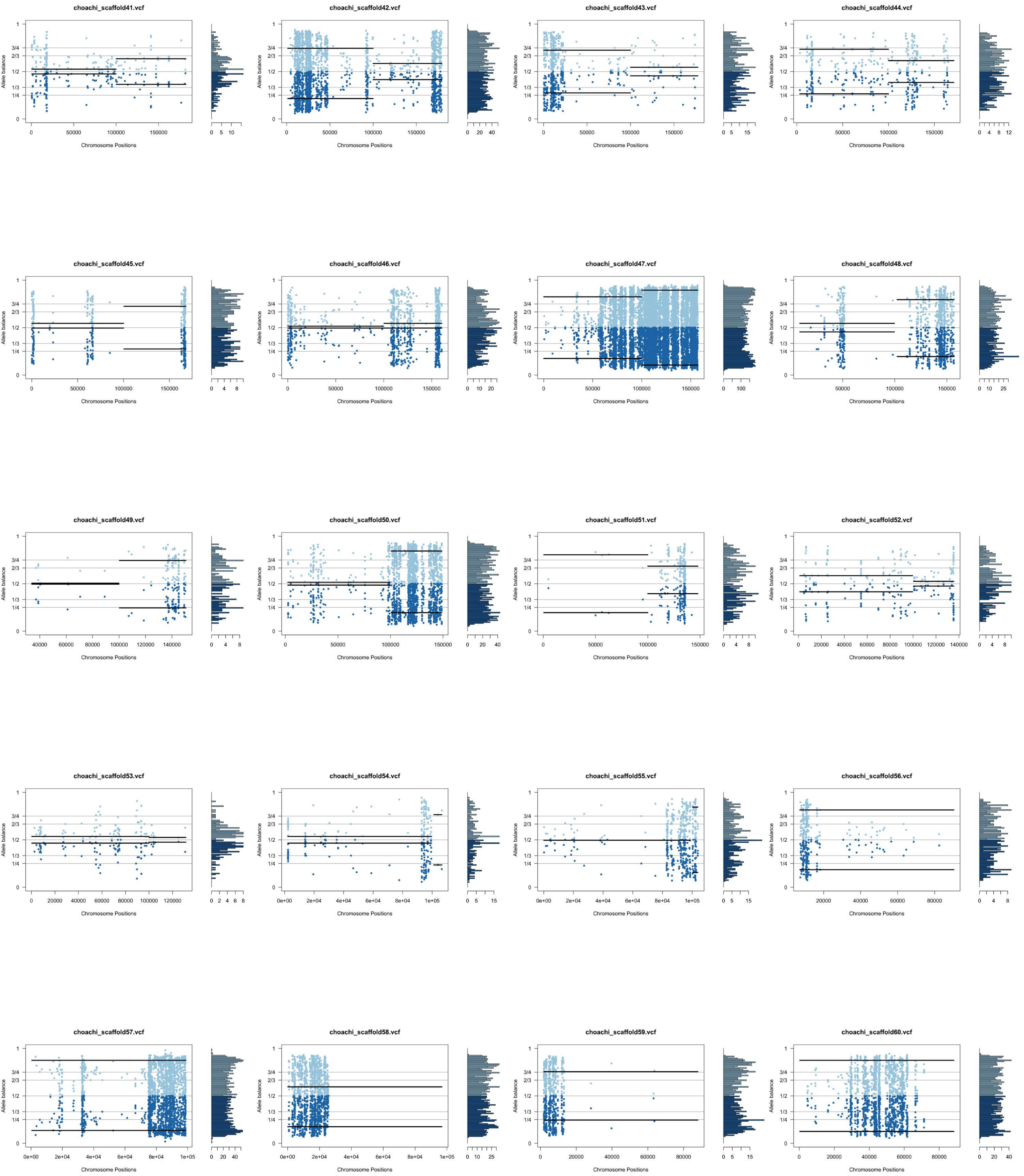
choachi_scaffold41.vcf
choachi_scaffold42.vcf
choachi_scaffold43.vcf
choachi_scaffold44.vcf
choachi_scaffold45.vcf
choachi_scaffold46.vcf
choachi_scaffold47.vcf
choachi_scaffold48.vcf
choachi_scaffold49.vcf
choachi_scaffold50.vcf
choachi_scaffold51.vcf
choachi_scaffold52.vcf
choachi_scaffold53.vcf
choachi_scaffold54.vcf
choachi_scaffold55.vcf
choachi_scaffold56.vcf
choachi_scaffold57.vcf
choachi_scaffold58.vcf
choachi_scaffold59.vcf
choachi_scaffold60.vcf
Allele balance
Chromosome Positions

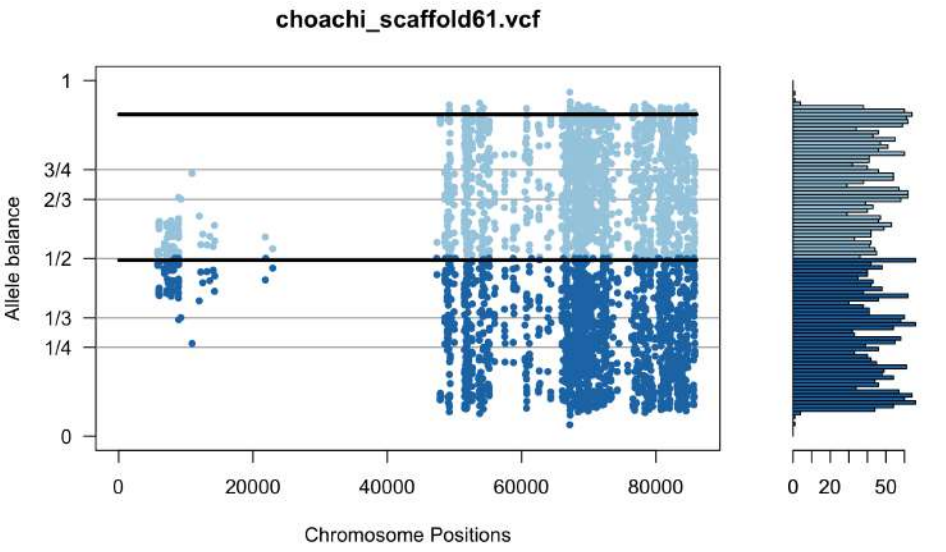
choachi_scaffold61.vcf
Allele balance
Chromosome Positions
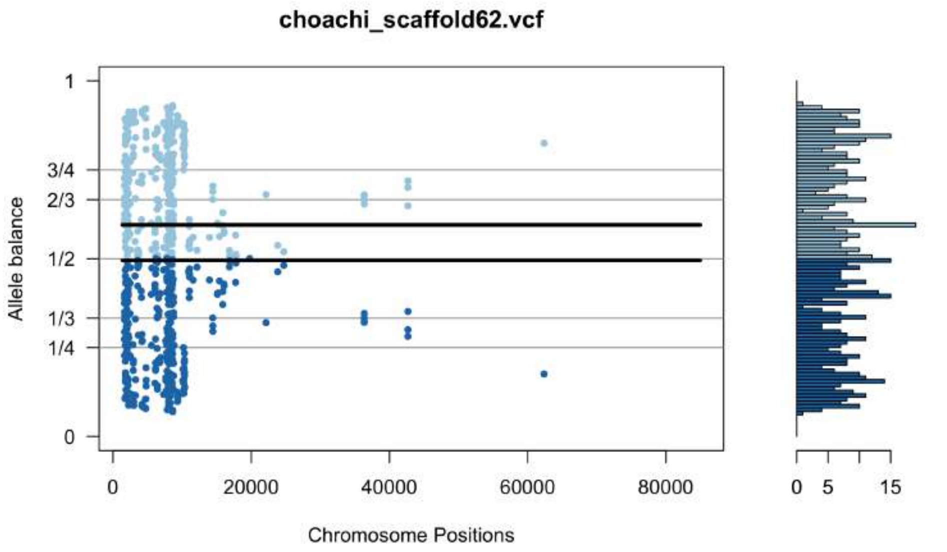
choachi_scaffold62.vcf
Allele balance
Chromosome Positions
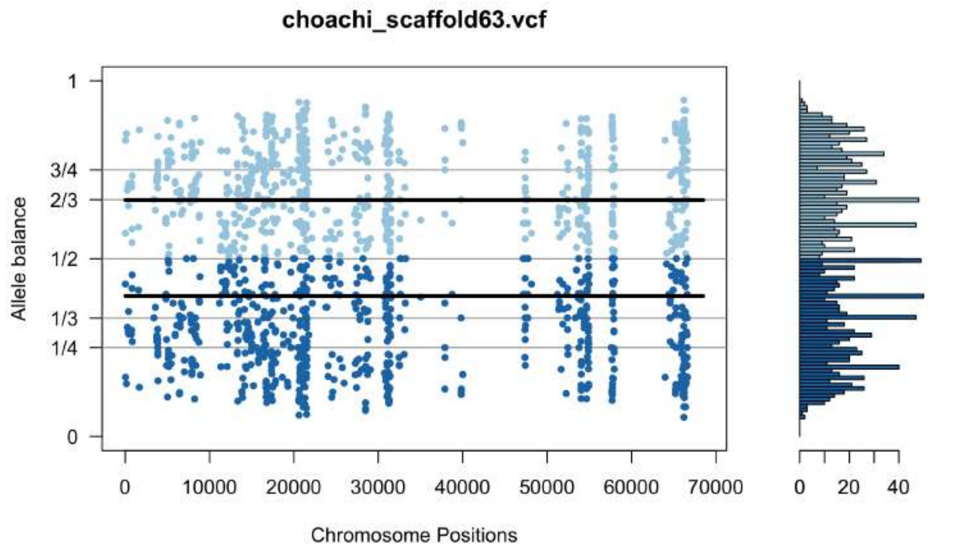
choachi_scaffold63.vcf
Allele balance
Chromosome Positions
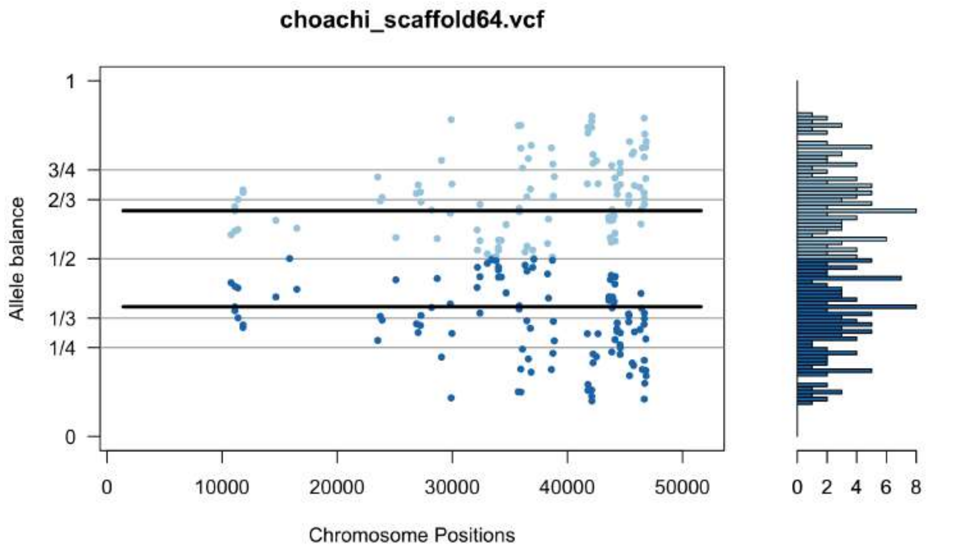
choachi_scaffold64.vcf
Allele balance
Chromosome Positions
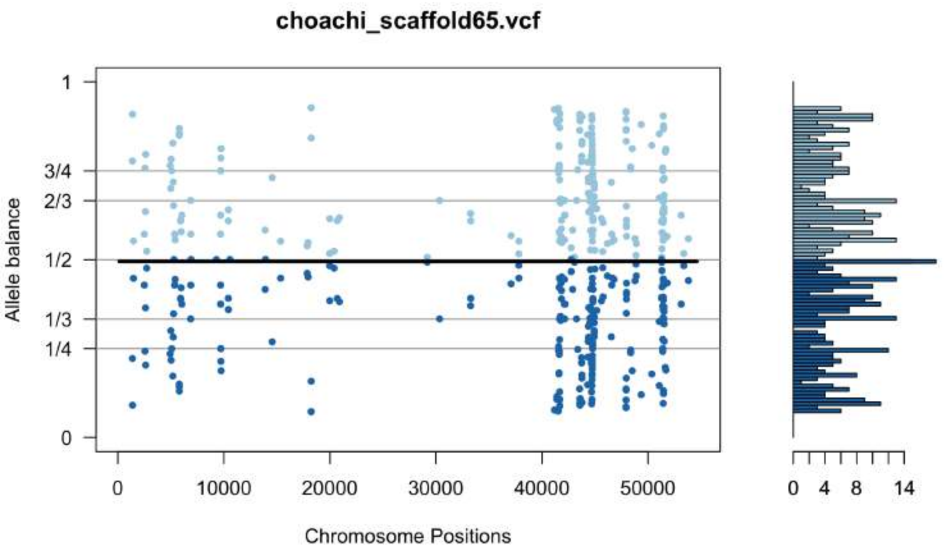
choachi_scaffold65.vcf
Allele balance
Chromosome Positions
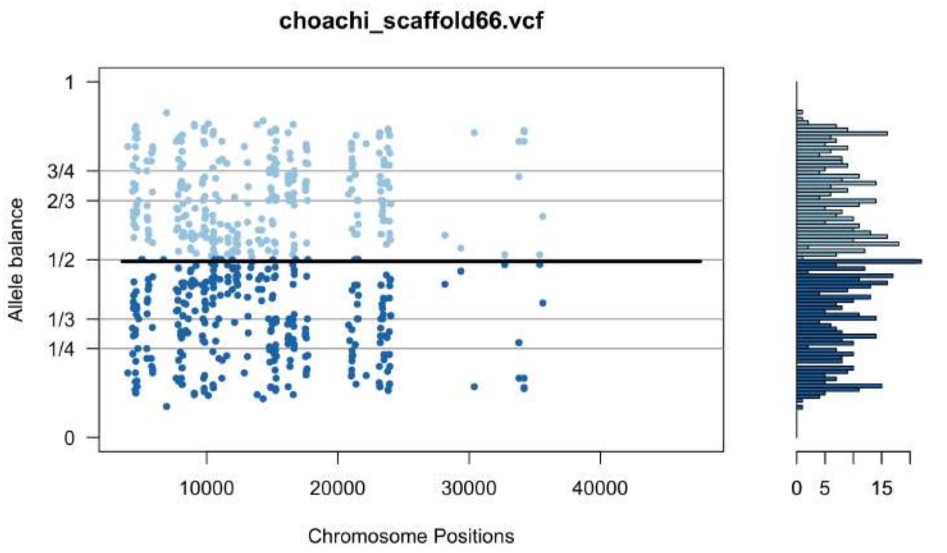
choachi_scaffold66.vcf
Allele balance
Chromosome Positions
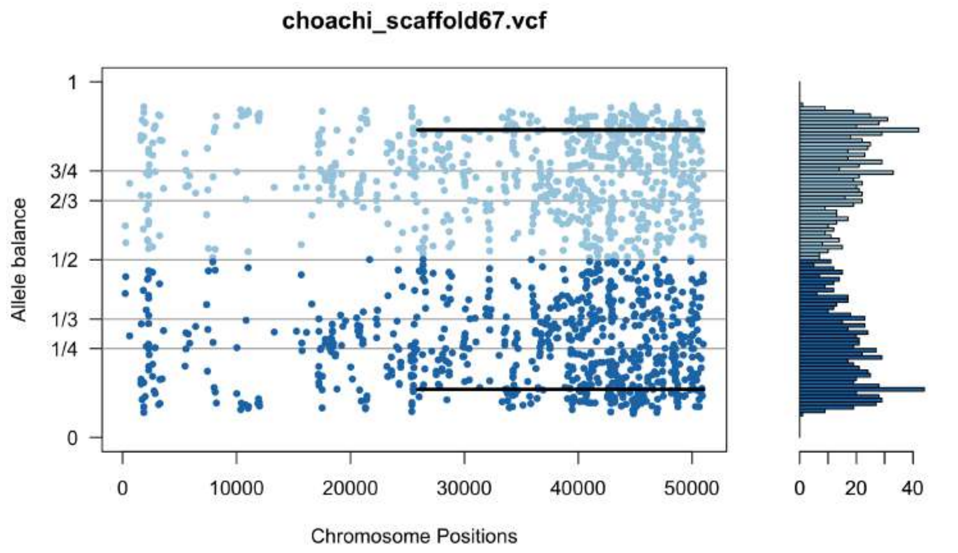
choachi_scaffold67.vcf
Allele balance
Chromosome Positions
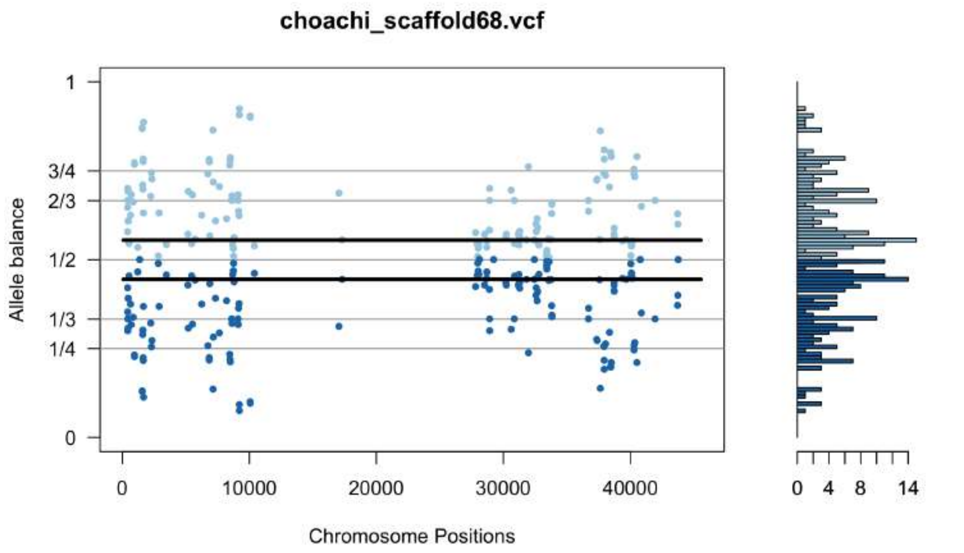
choachi_scaffold68.vcf
Allele balance
Chromosome Positions
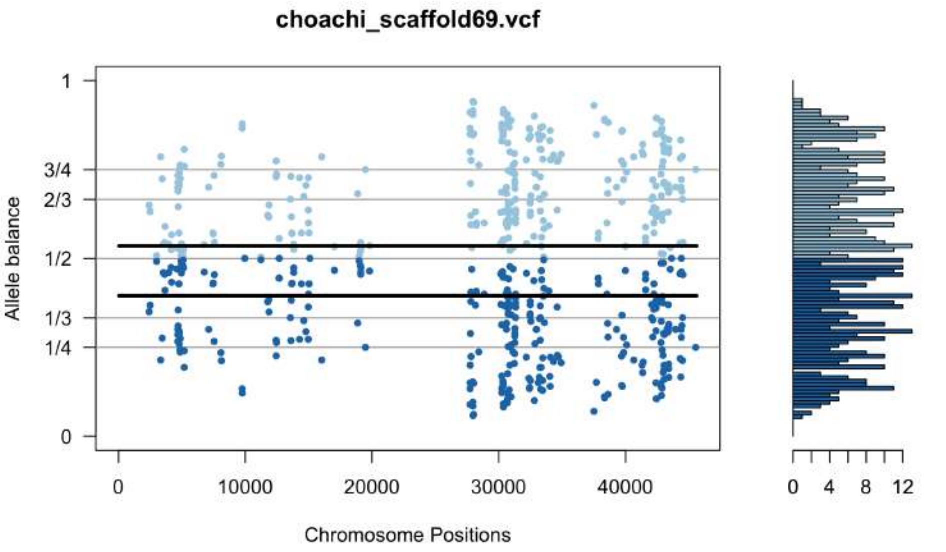
choachi_scaffold69.vcf
Allele balance
Chromosome Positions
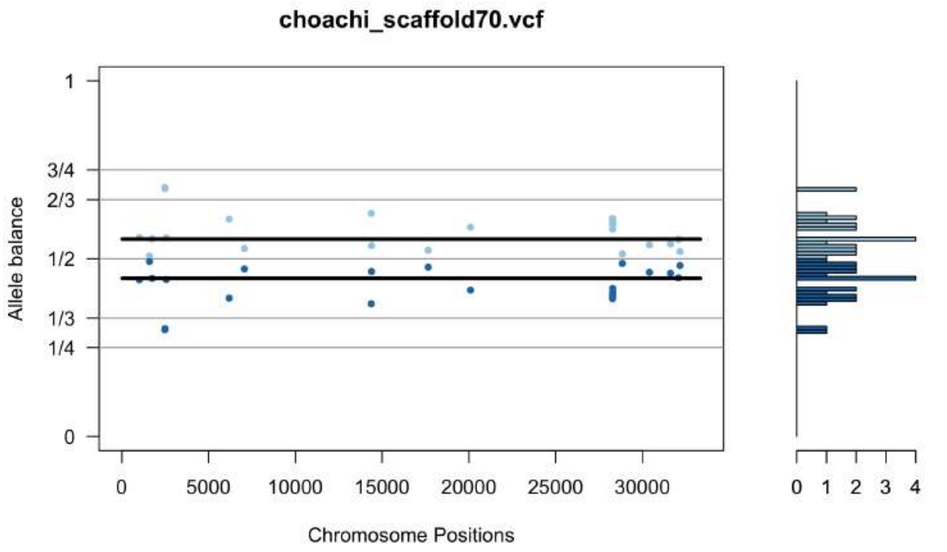
choachi_scaffold70.vcf
Allele balance
Chromosome Positions
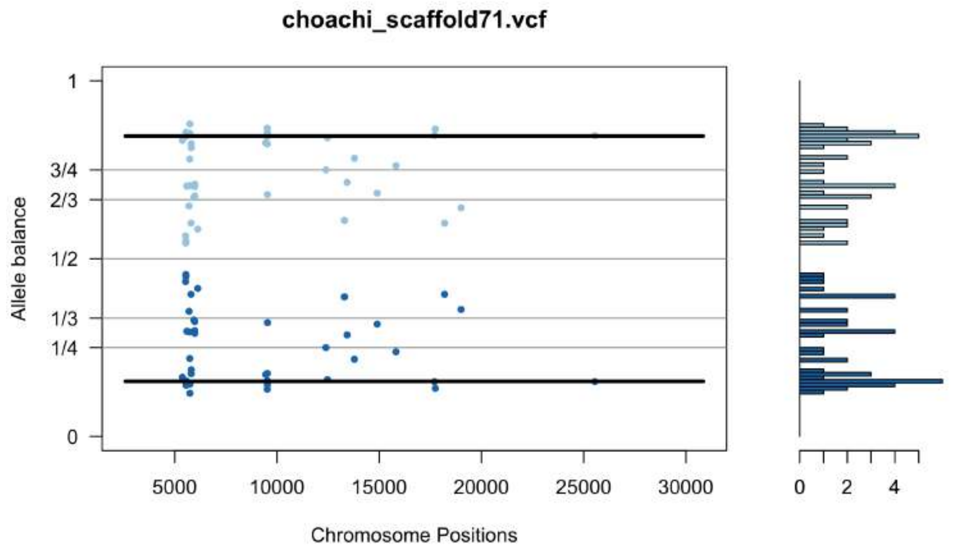
choachi_scaffold71.vcf
Allele balance
Chromosome Positions
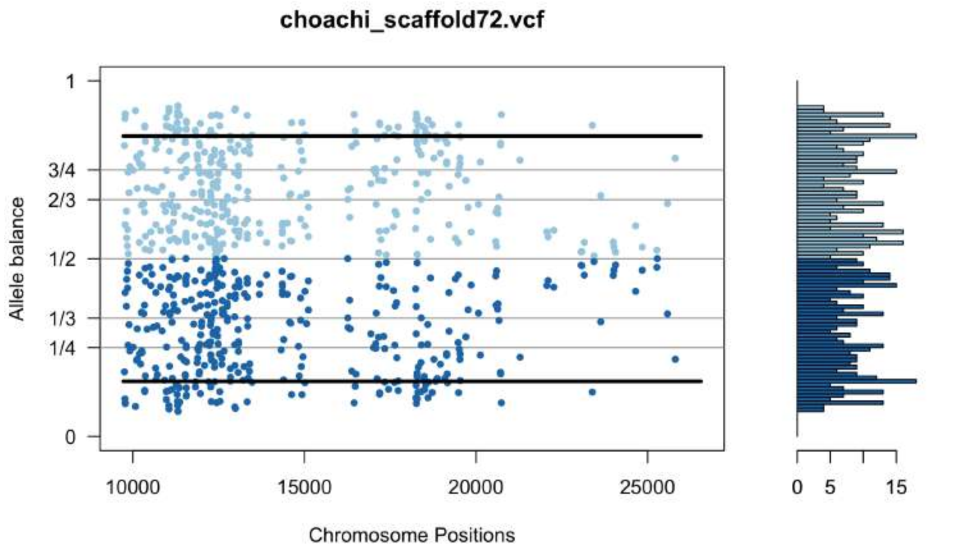
choachi_scaffold72.vcf
Allele balance
Chromosome Positions
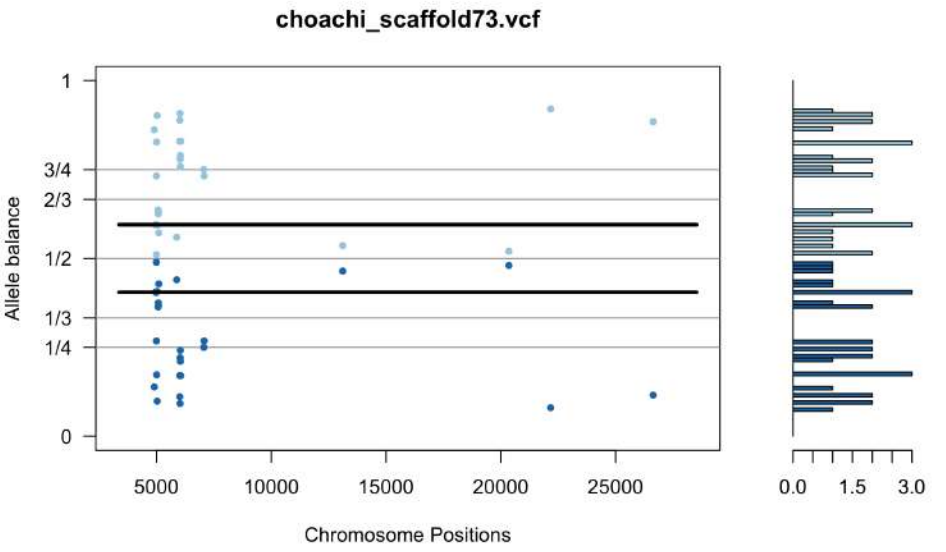
choachi_scaffold73.vcf
Allele balance
Chromosome Positions
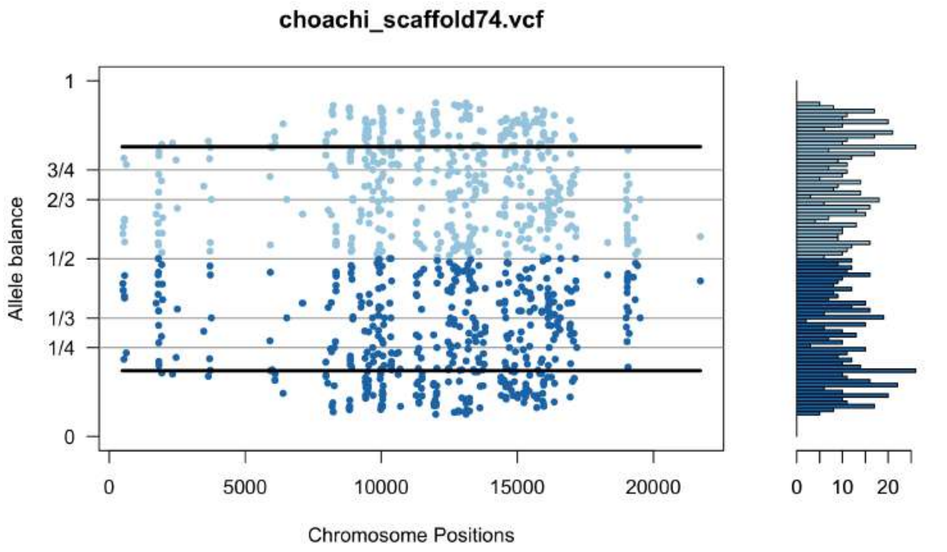
choachi_scaffold74.vcf
Allele balance
Chromosome Positions
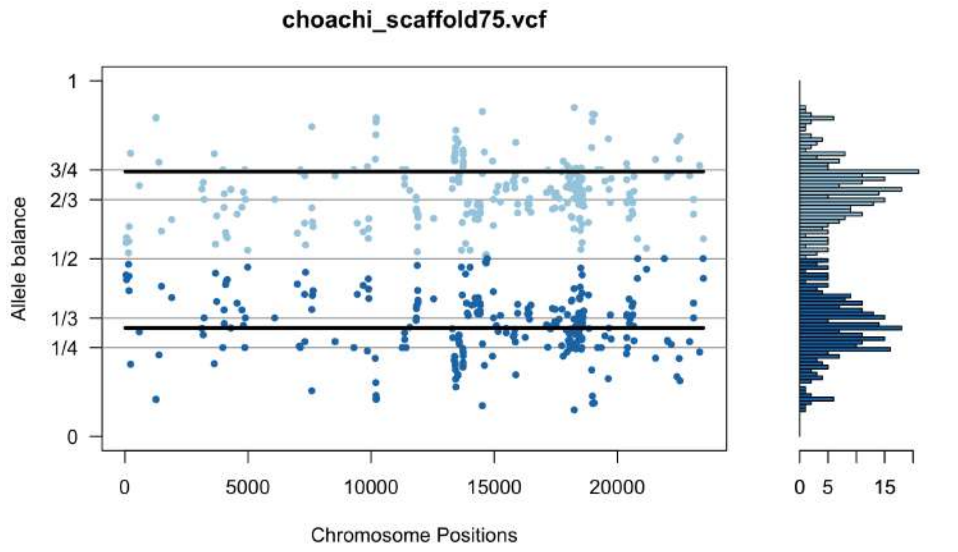
choachi_scaffold75.vcf
Allele balance
Chromosome Positions
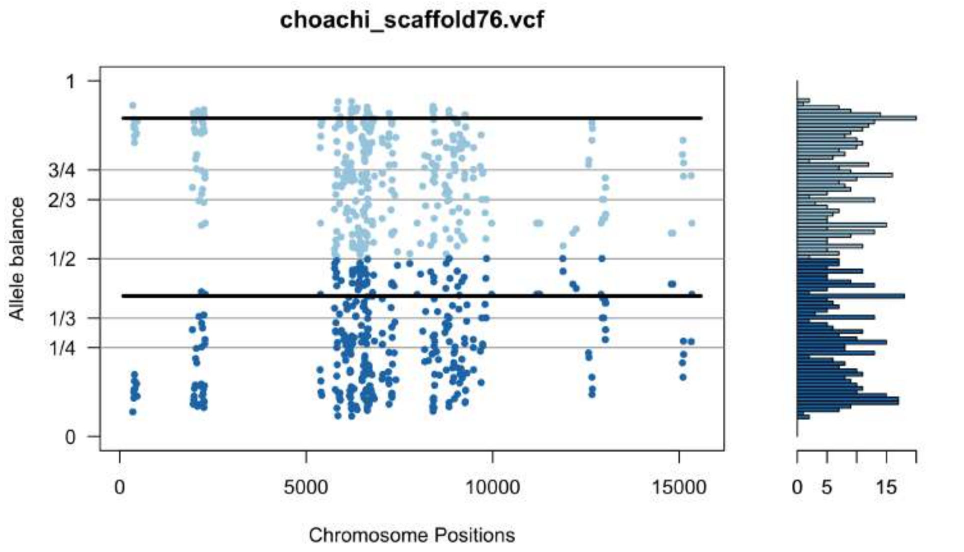
choachi_scaffold76.vcf
Allele balance
Chromosome Positions
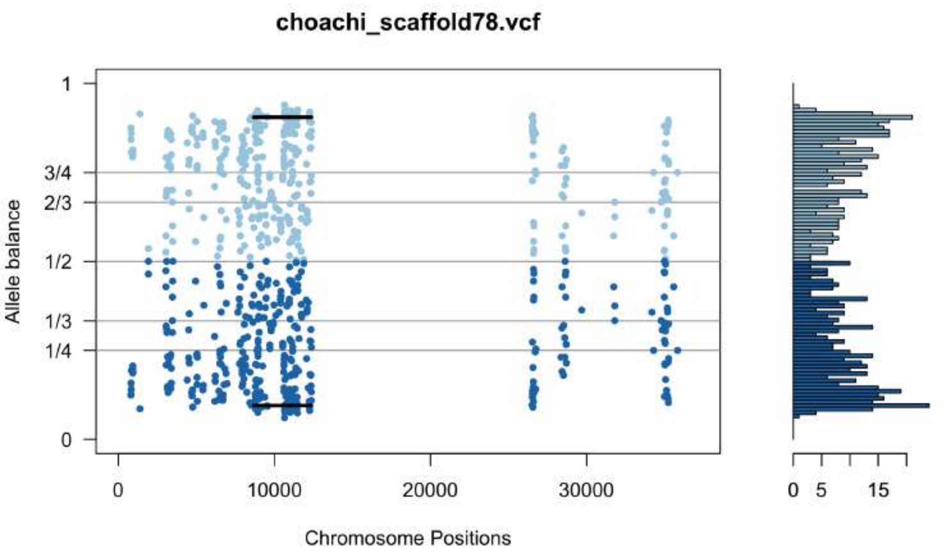
choachi_scaffold78.vcf
Allele balance
Chromosome Positions
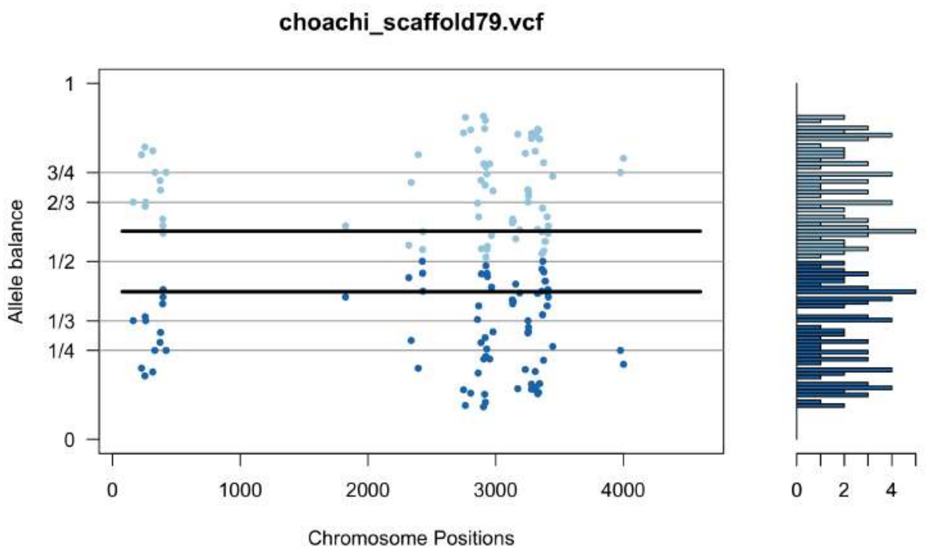
choachi_scaffold79.vcf
Allele balance
Chromosome Positions
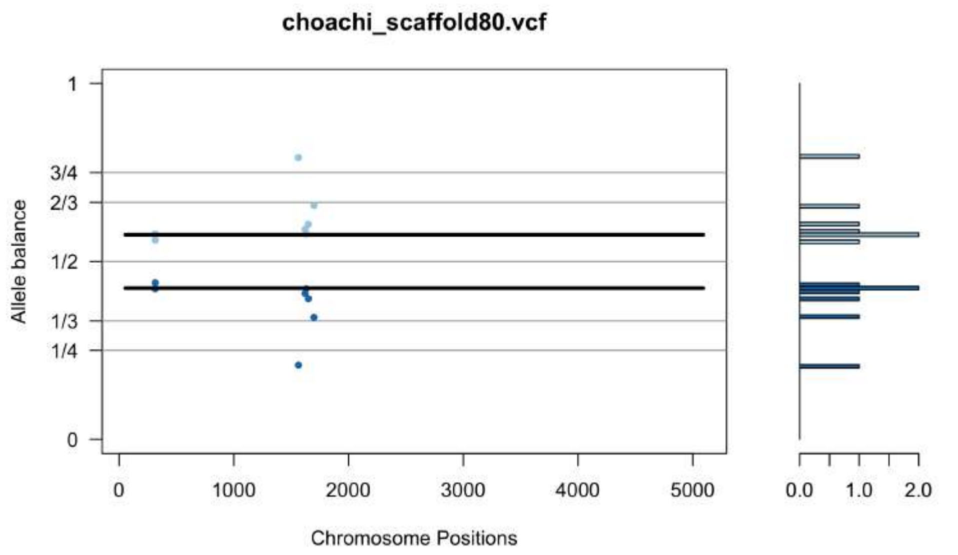
choachi_scaffold80.vcf
Allele balance
Chromosome Positions
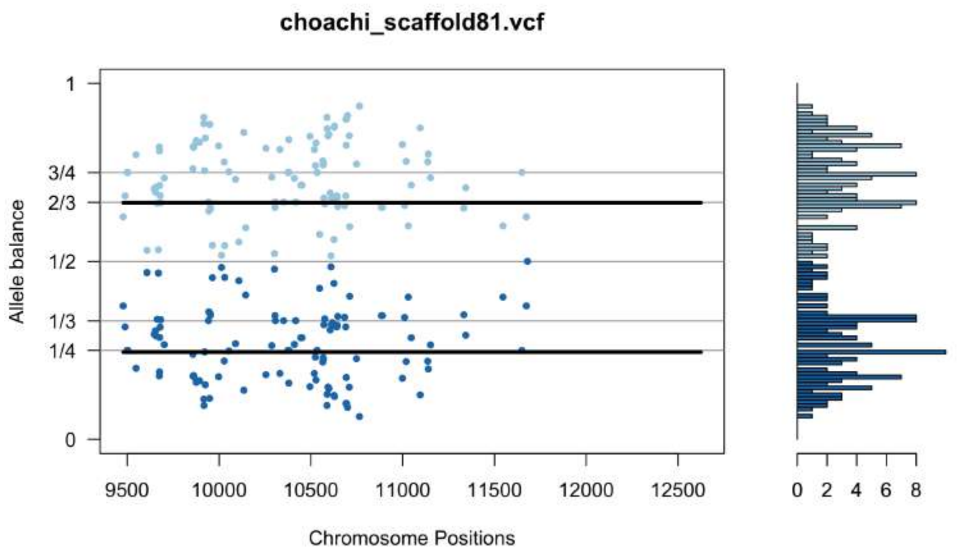
choachi_scaffold81.vcf
Allele balance
Chromosome Positions

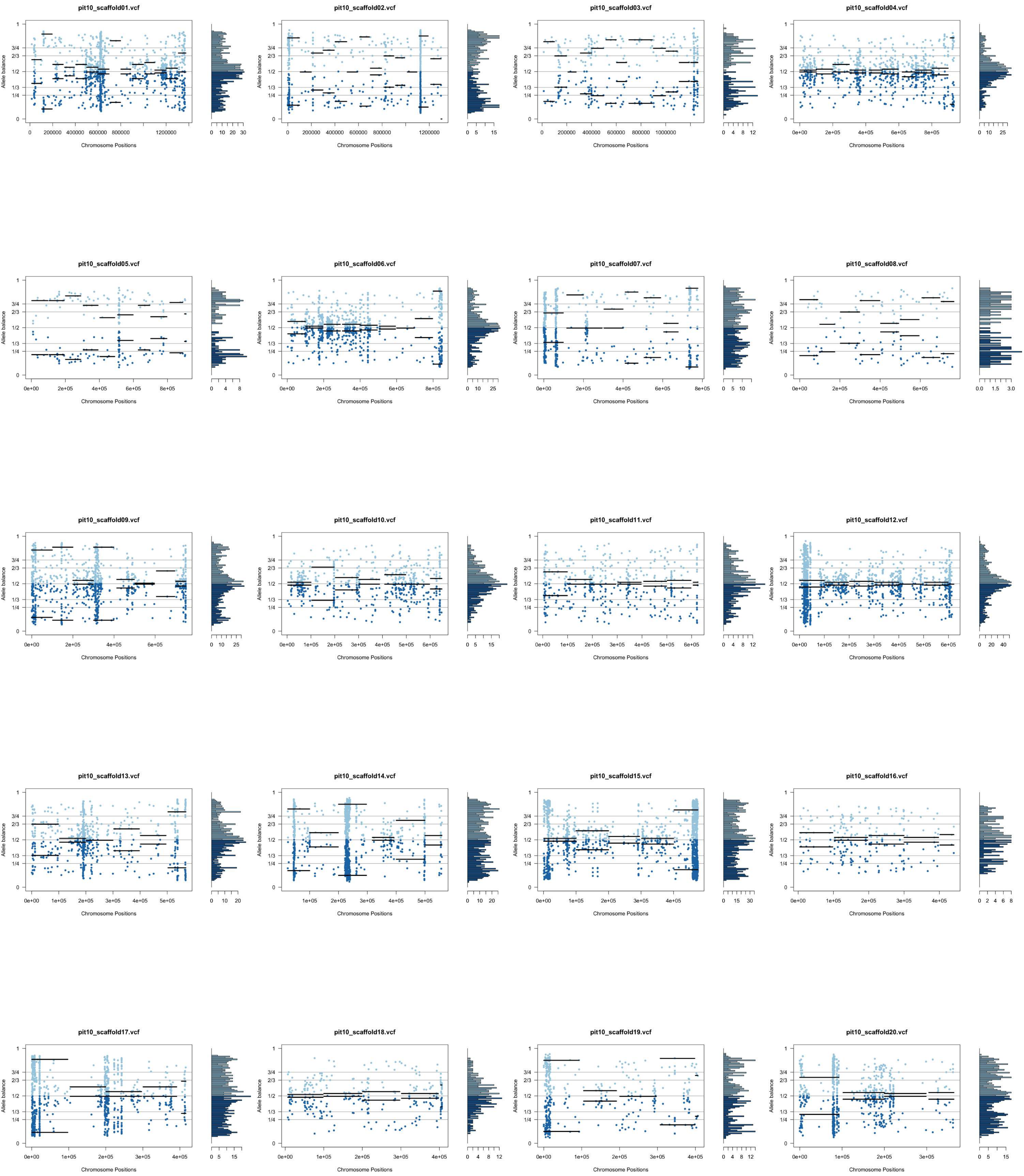
pit10_scaffold01.vcf
pit10_scaffold02.vcf
pit10_scaffold03.vcf
pit10_scaffold04.vcf
pit10_scaffold05.vcf
pit10_scaffold06.vcf
pit10_scaffold07.vcf
pit10_scaffold08.vcf
pit10_scaffold09.vcf
pit10_scaffold10.vcf
pit10_scaffold11.vcf
pit10_scaffold12.vcf
pit10_scaffold13.vcf
pit10_scaffold14.vcf
pit10_scaffold15.vcf
pit10_scaffold16.vcf
pit10_scaffold17.vcf
pit10_scaffold18.vcf
pit10_scaffold19.vcf
pit10_scaffold20.vcf
Allele balance
Chromosome Positions

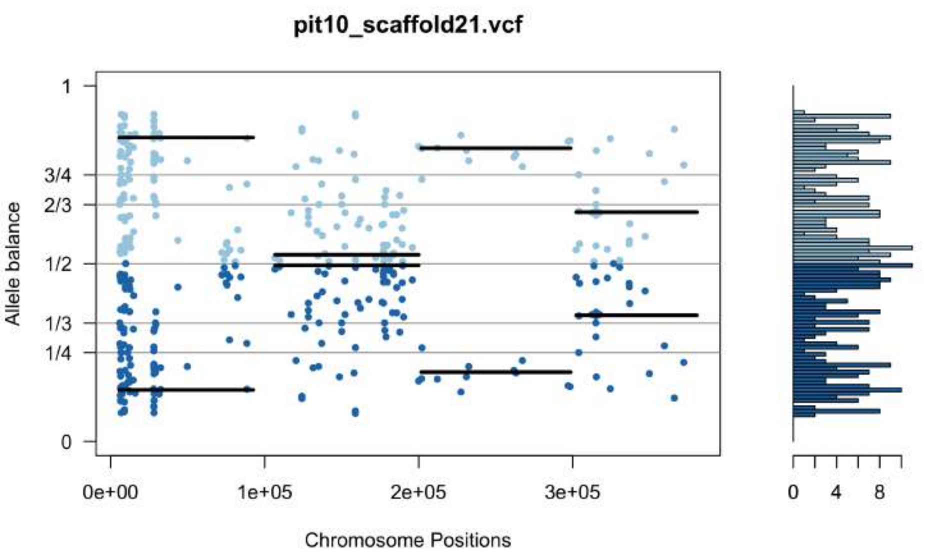
pit10_scaffold21.vcf
Allele balance
Chromosome Positions

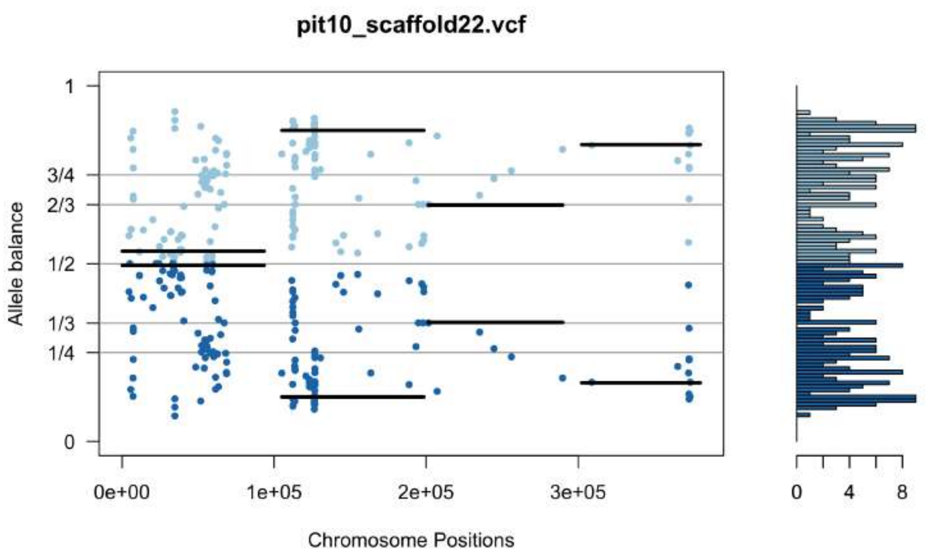
pit10_scaffold22.vcf
Allele balance
Chromosome Positions

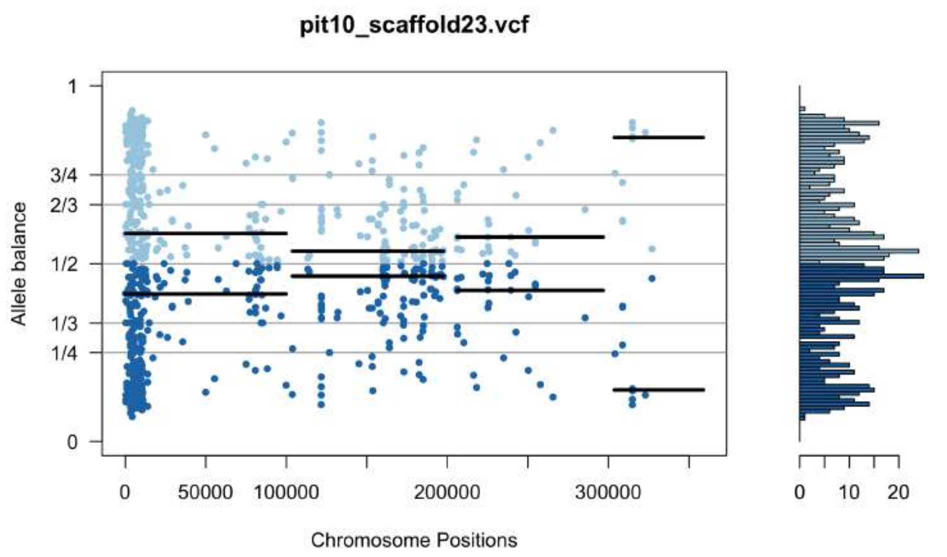
pit10_scaffold23.vcf
Allele balance
Chromosome Positions

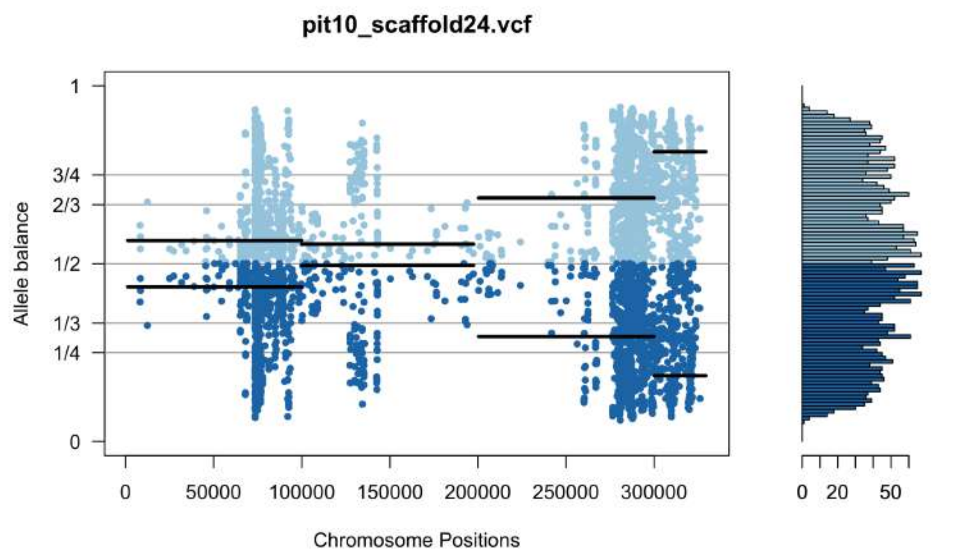
pit10_scaffold24.vcf
Allele balance
Chromosome Positions

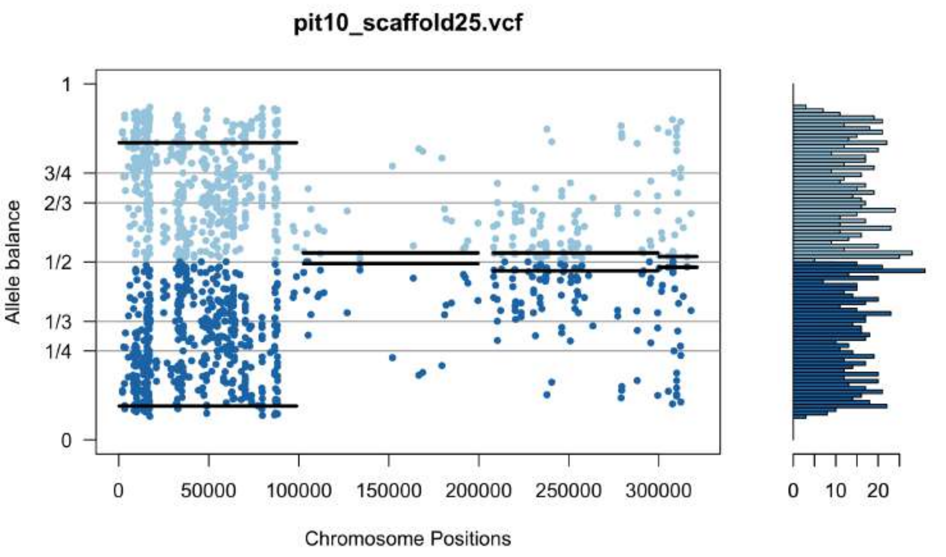
pit10_scaffold25.vcf
Allele balance
Chromosome Positions

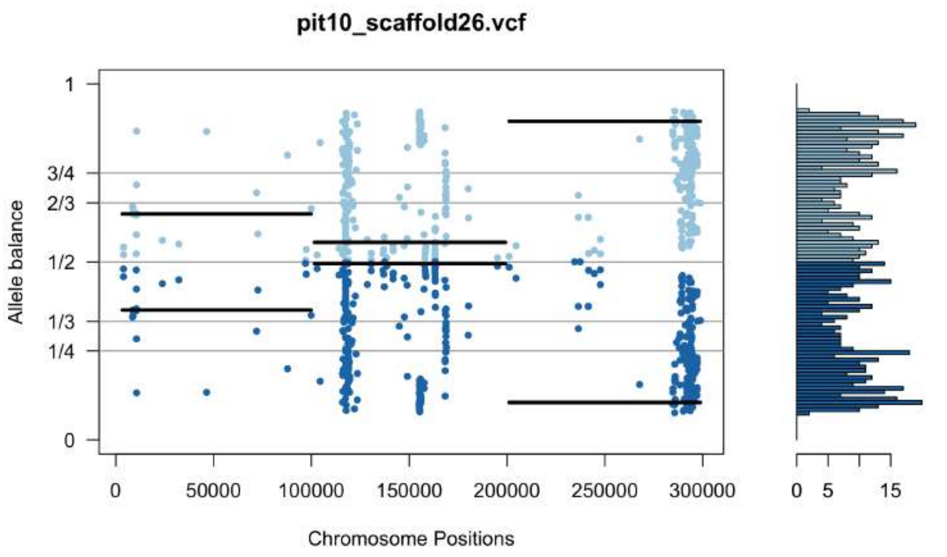
pit10_scaffold26.vcf
Allele balance
Chromosome Positions

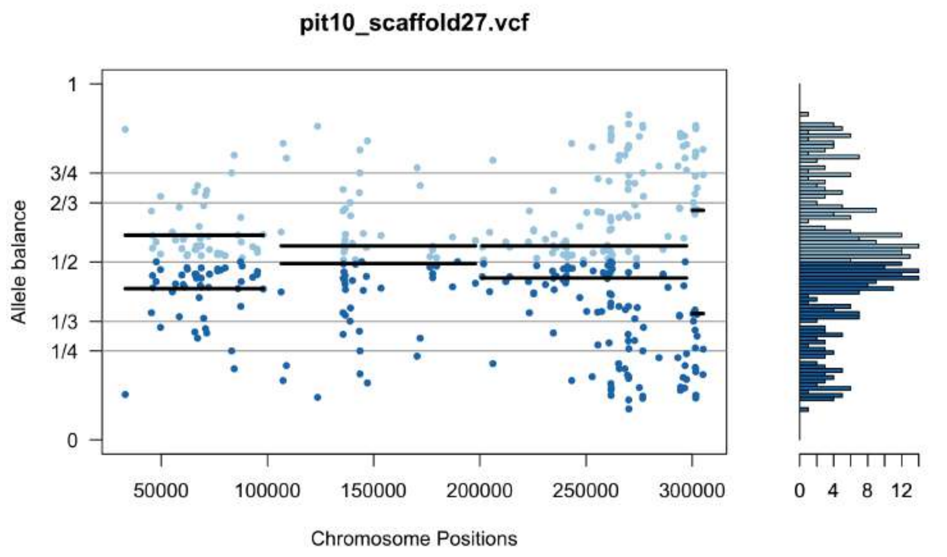
pit10_scaffold27.vcf
Allele balance
Chromosome Positions

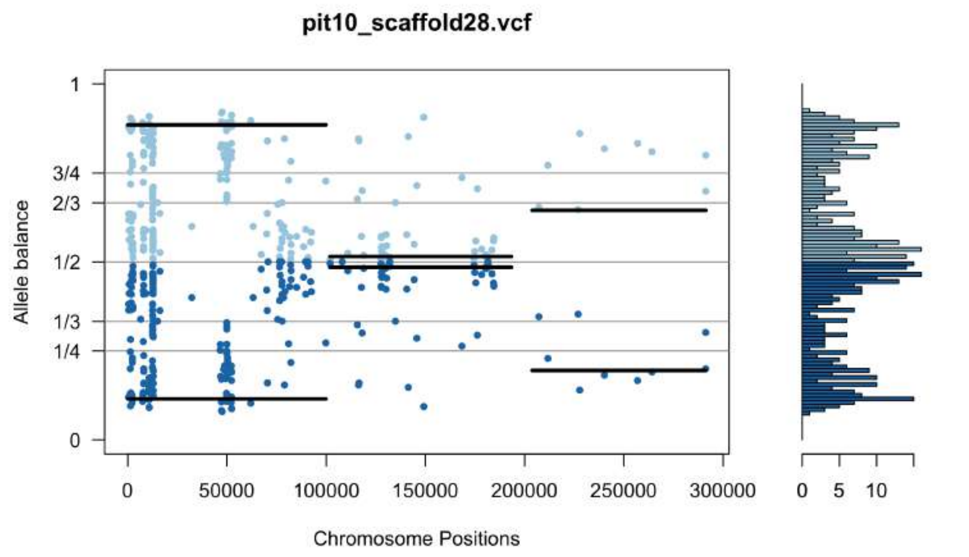
pit10_scaffold28.vcf
Allele balance
Chromosome Positions

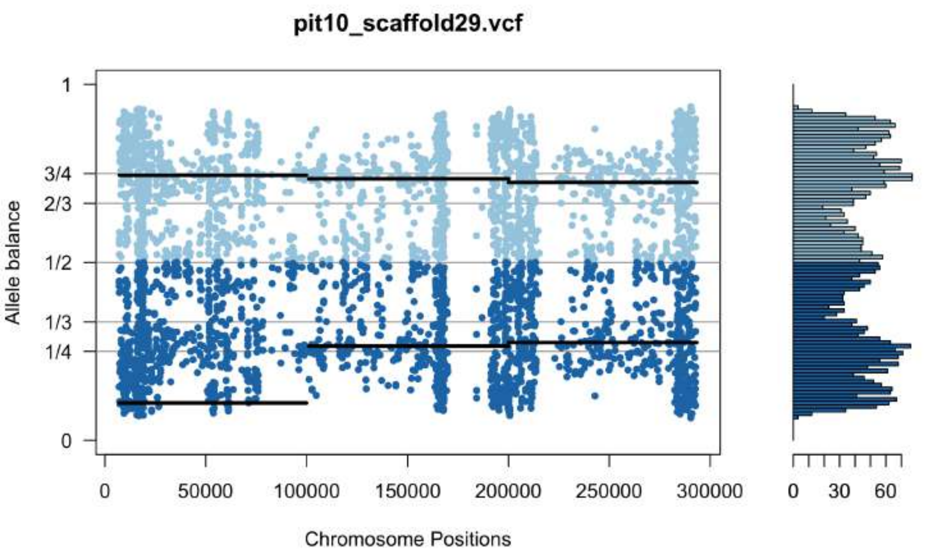
pit10_scaffold29.vcf
Allele balance
Chromosome Positions

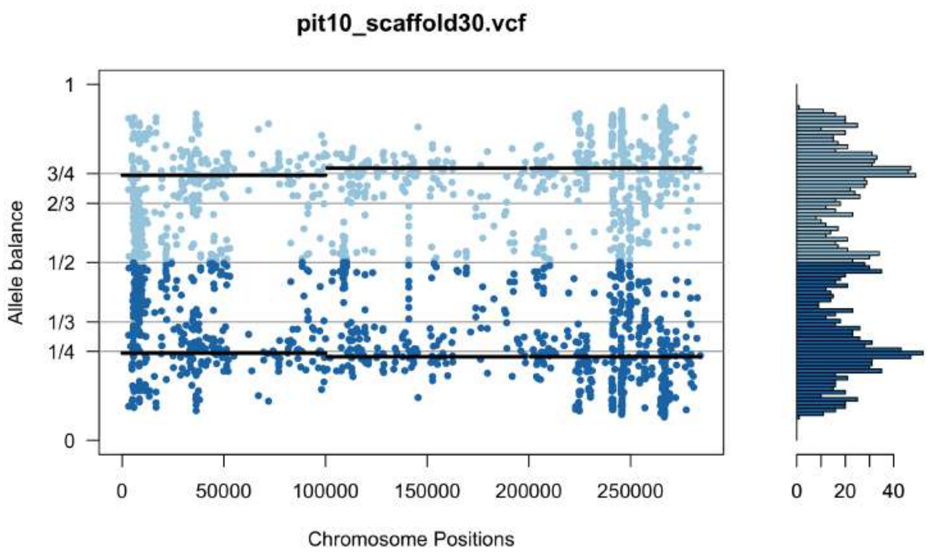
pit10_scaffold30.vcf
Allele balance
Chromosome Positions

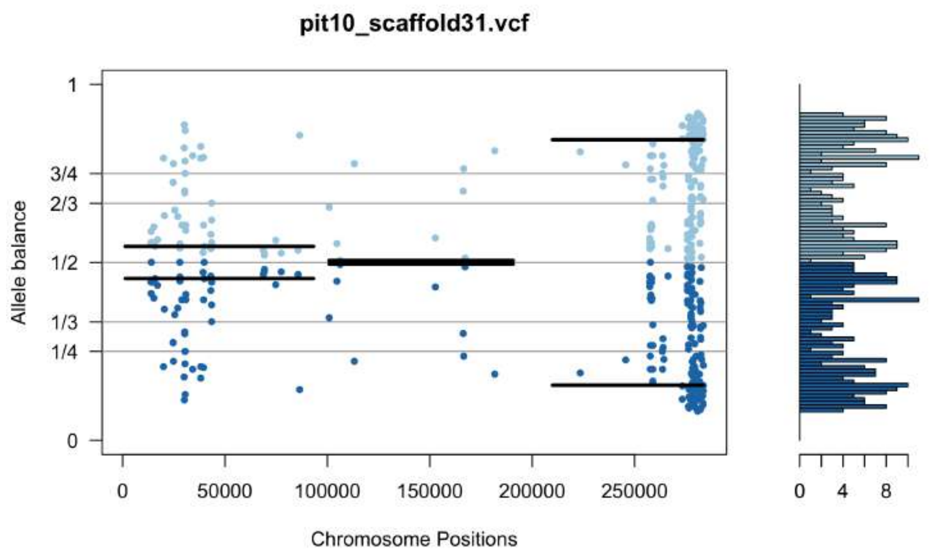
pit10_scaffold31.vcf
Allele balance
Chromosome Positions

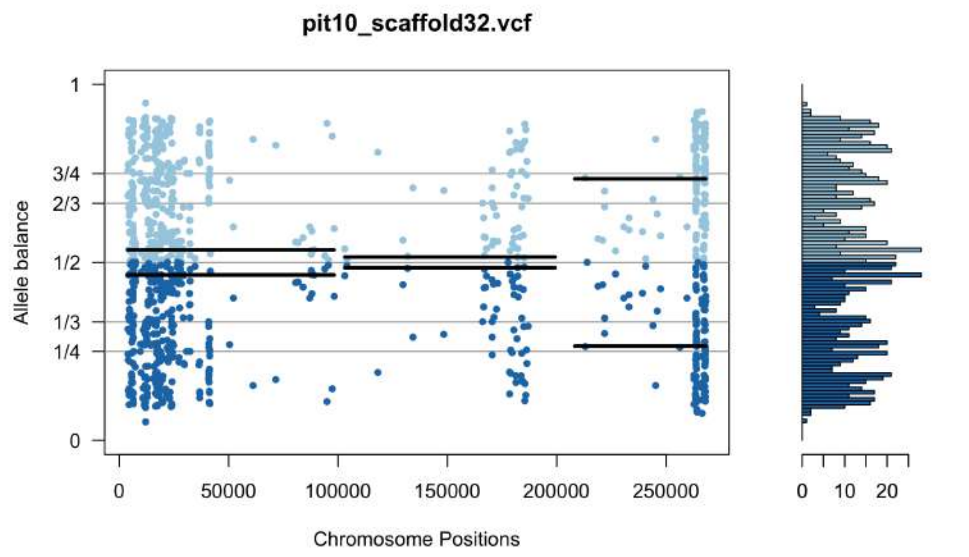
pit10_scaffold32.vcf
Allele balance
Chromosome Positions

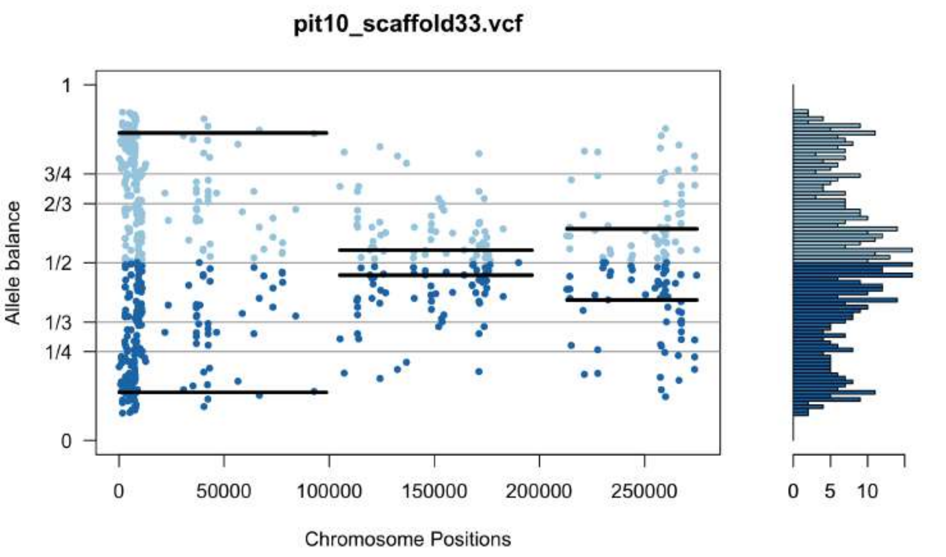
pit10_scaffold33.vcf
Allele balance
Chromosome Positions

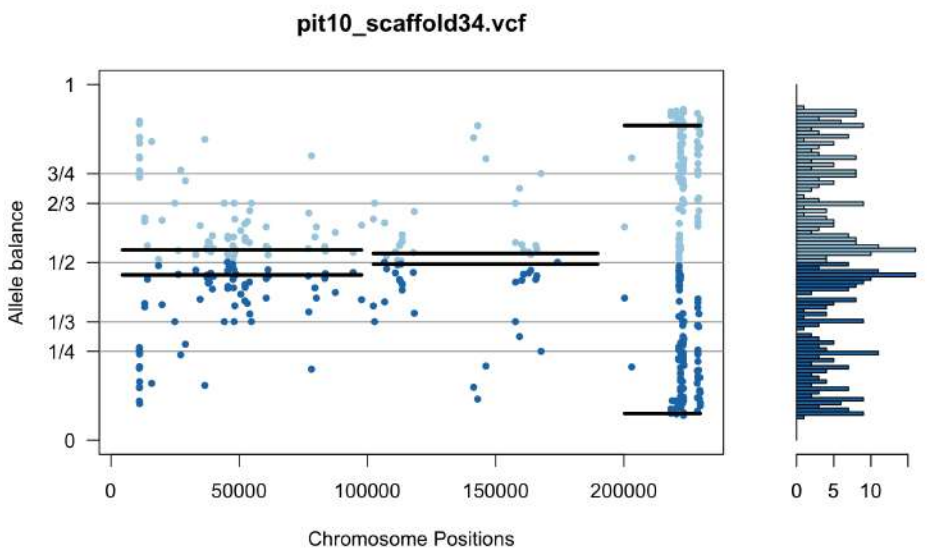
pit10_scaffold34.vcf
Allele balance
Chromosome Positions

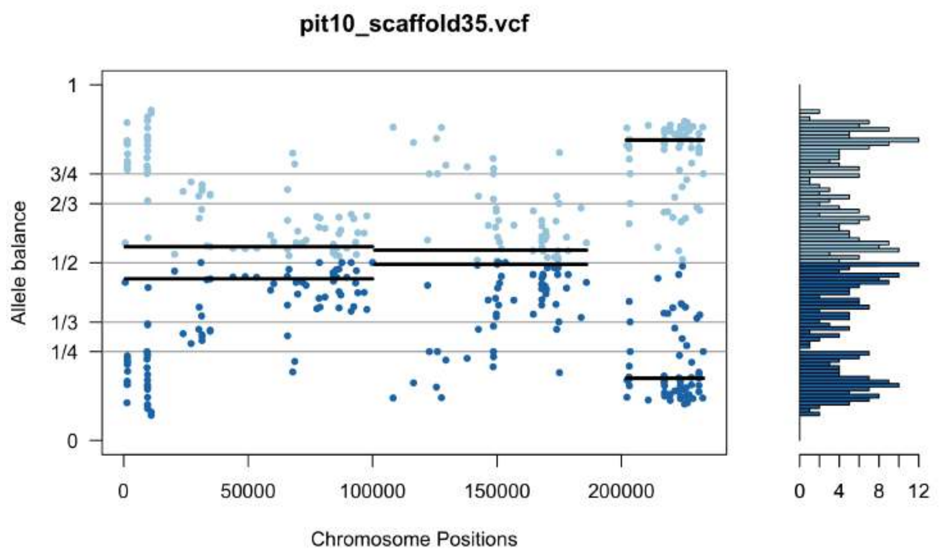
pit10_scaffold35.vcf
Allele balance
Chromosome Positions

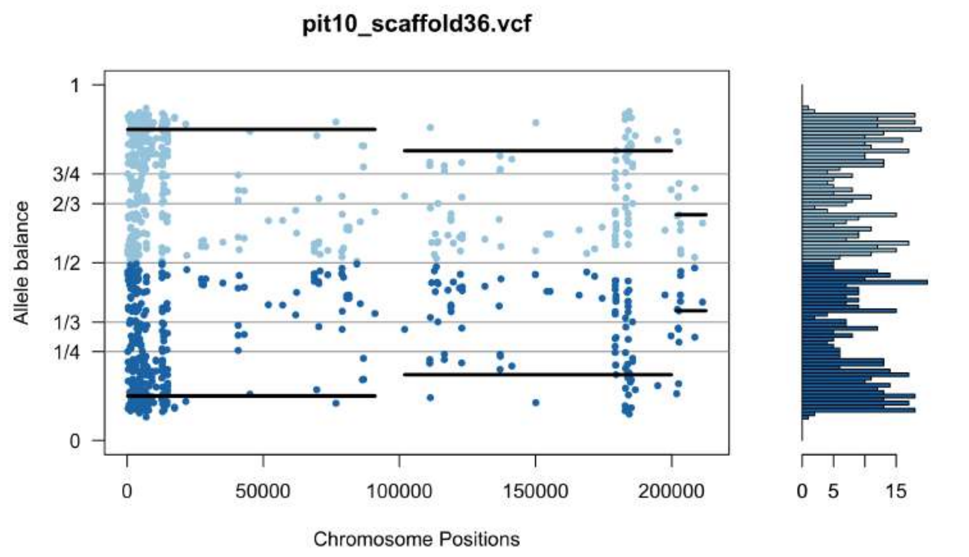
pit10_scaffold36.vcf
Allele balance
Chromosome Positions

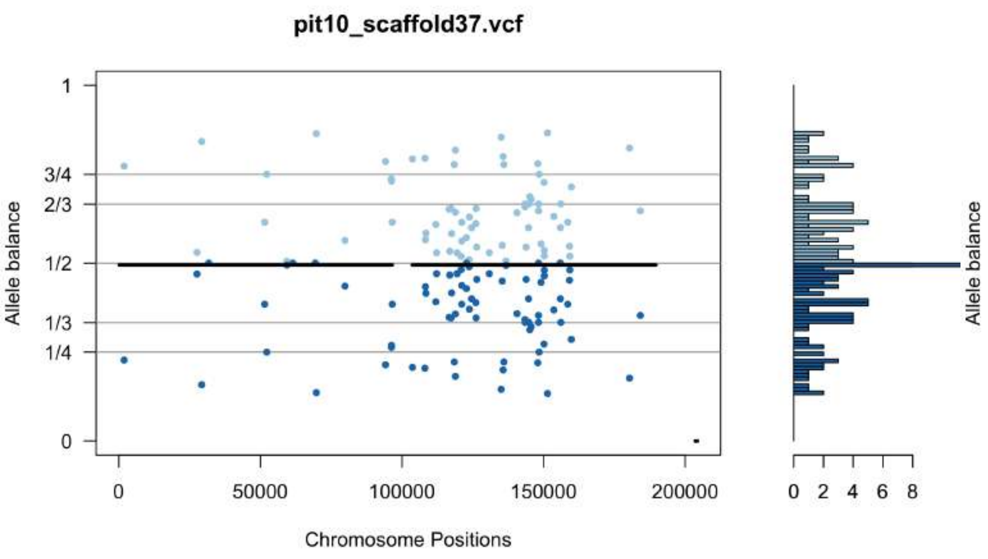
pit10_scaffold37.vcf
Allele balance
Chromosome Positions

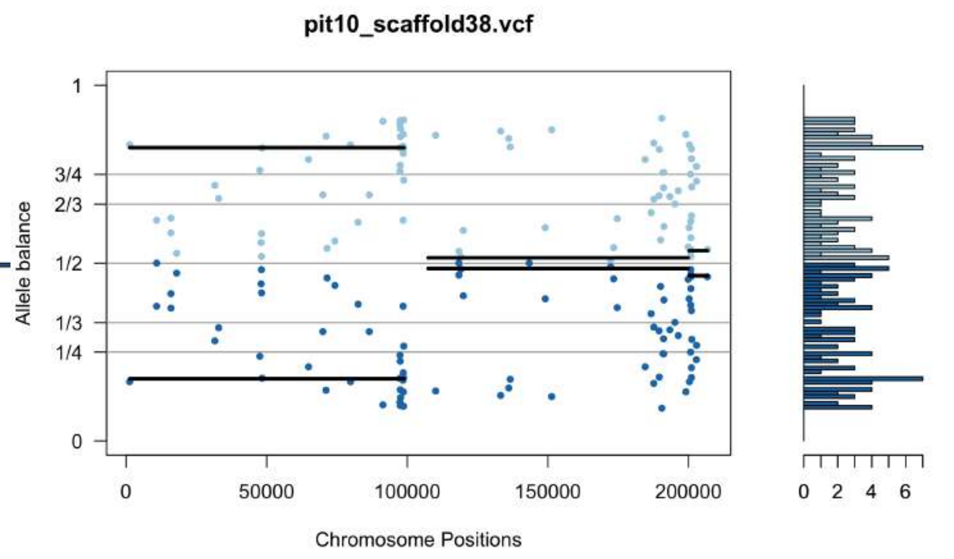
pit10_scaffold38.vcf
Allele balance
Chromosome Positions

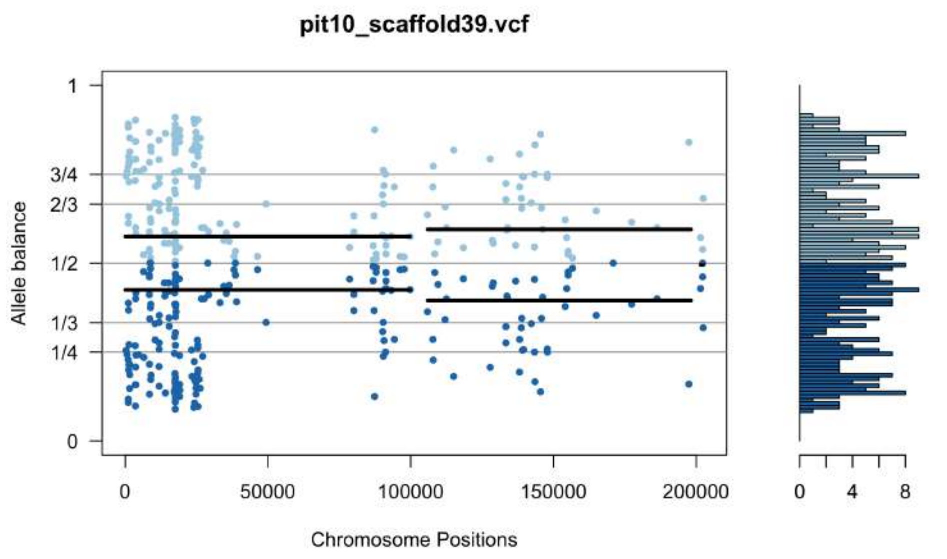
pit10_scaffold39.vcf
Allele balance
Chromosome Positions

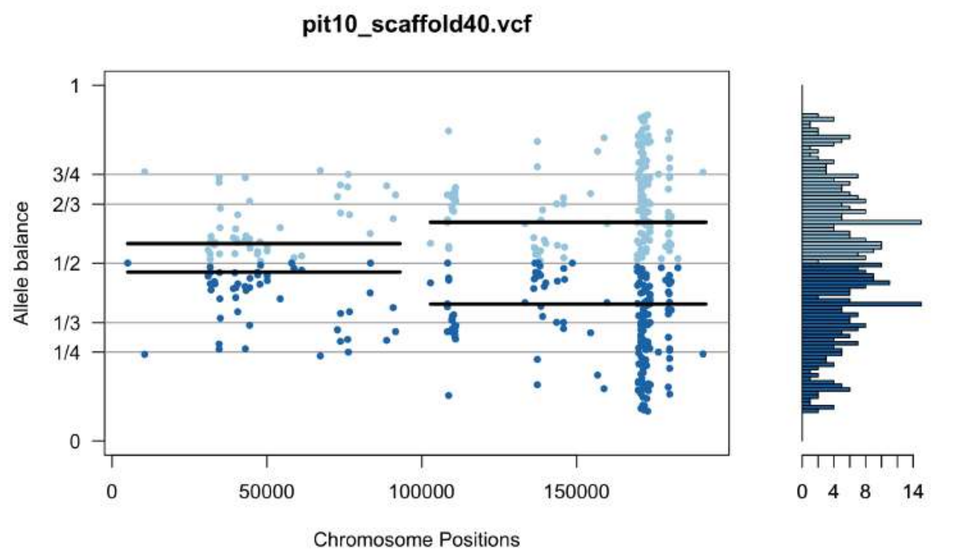
pit10_scaffold40.vcf
Allele balance
Chromosome Positions

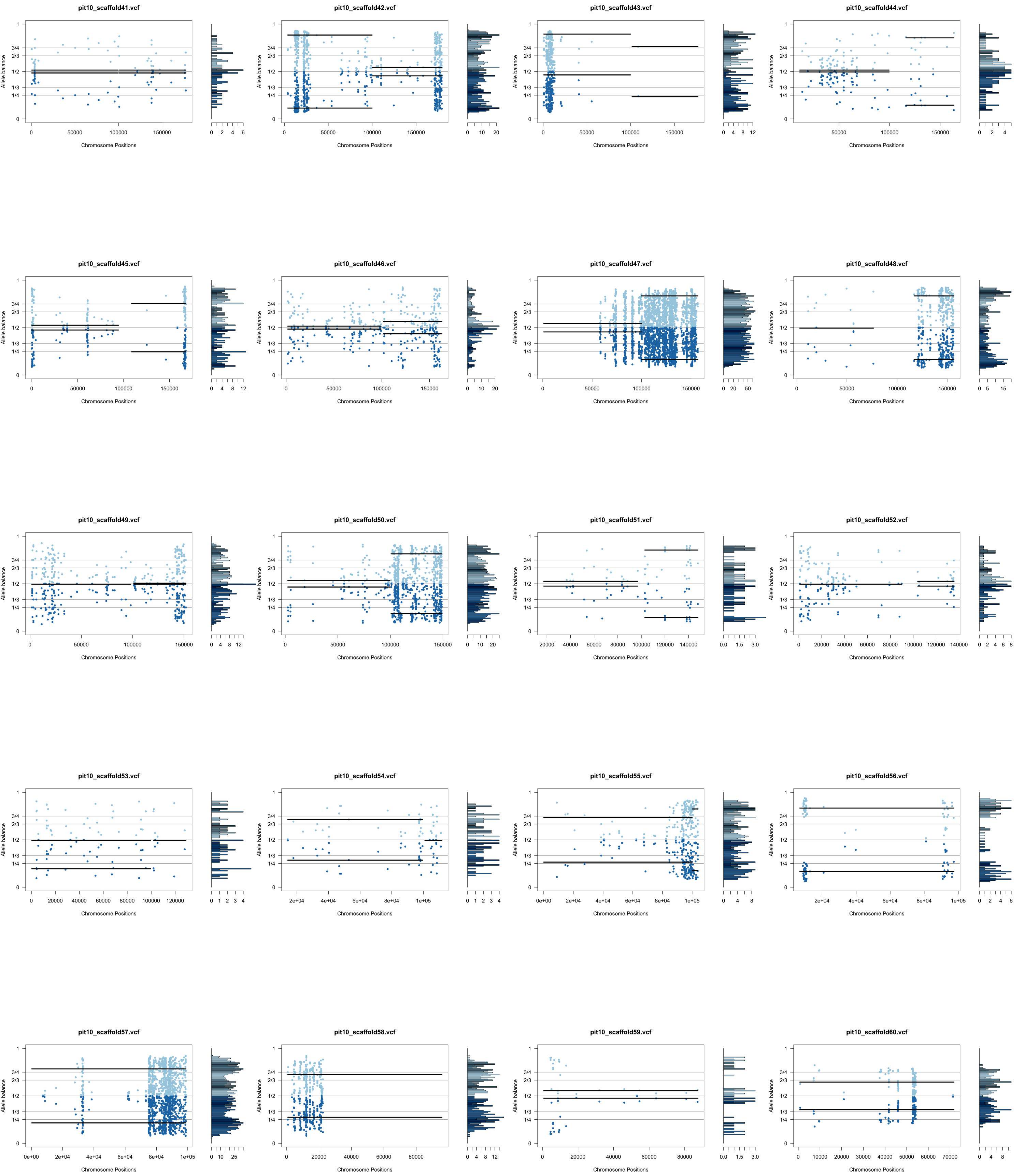
pit10_scaffold41.vcf
pit10_scaffold42.vcf
pit10_scaffold43.vcf
pit10_scaffold44.vcf
pit10_scaffold45.vcf
pit10_scaffold46.vcf
pit10_scaffold47.vcf
pit10_scaffold48.vcf
pit10_scaffold49.vcf
pit10_scaffold50.vcf
pit10_scaffold51.vcf
pit10_scaffold52.vcf
pit10_scaffold53.vcf
pit10_scaffold54.vcf
pit10_scaffold55.vcf
pit10_scaffold56.vcf
pit10_scaffold57.vcf
pit10_scaffold58.vcf
pit10_scaffold59.vcf
pit10_scaffold60.vcf
Allele balance
Chromosome Positions

pit10_scaffold61.vcf

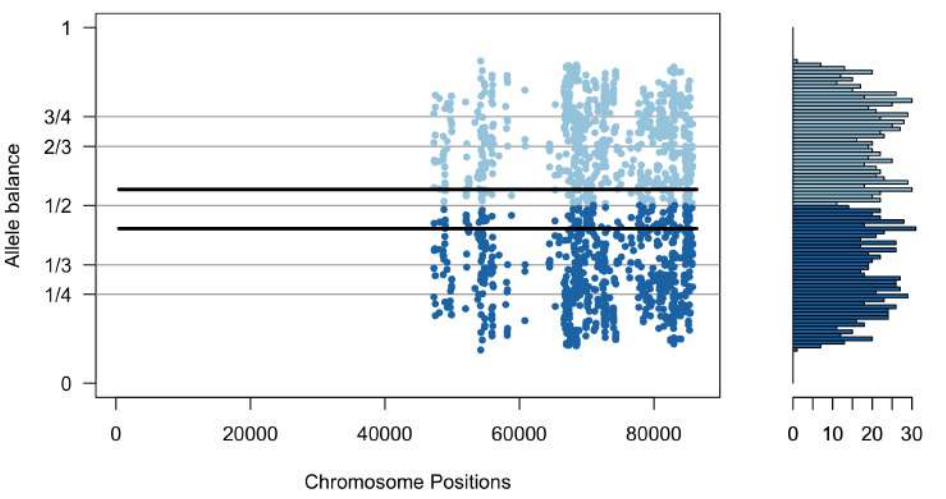


pit10_scaffold62.vcf

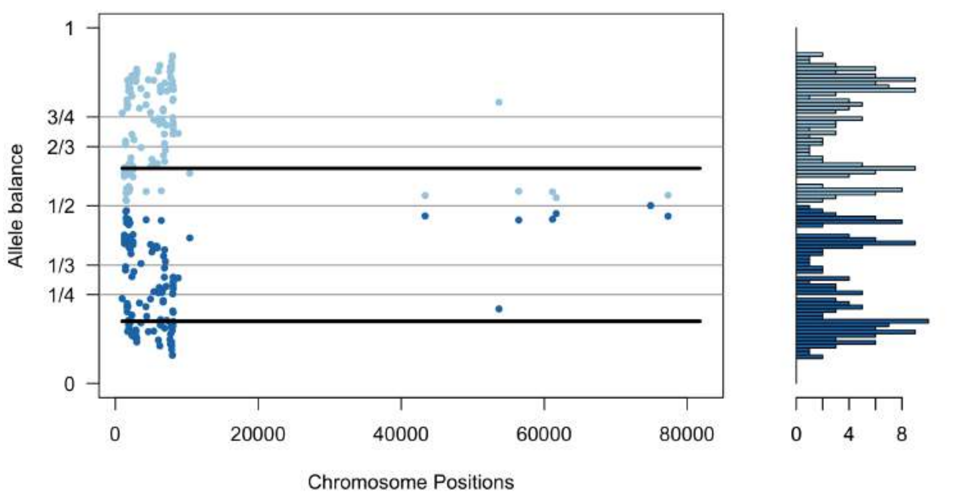


pit10_scaffold63.vcf

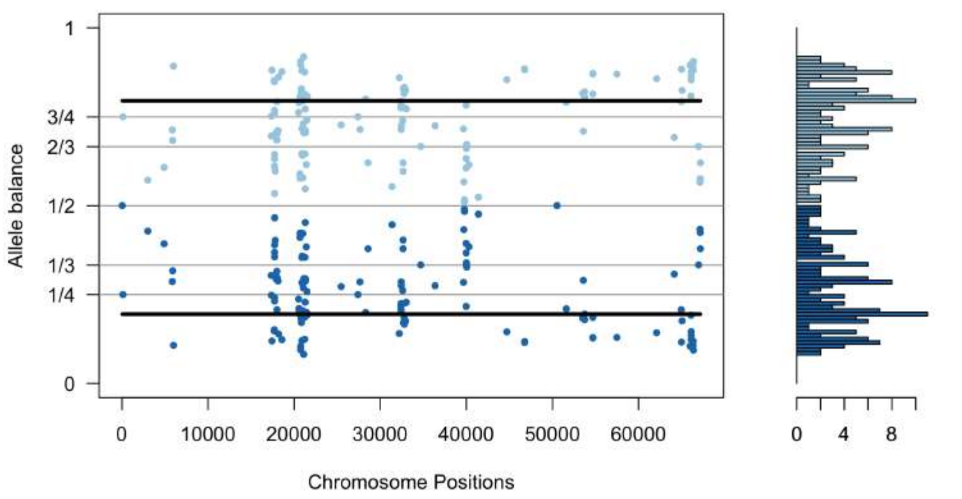


pit10_scaffold64.vcf

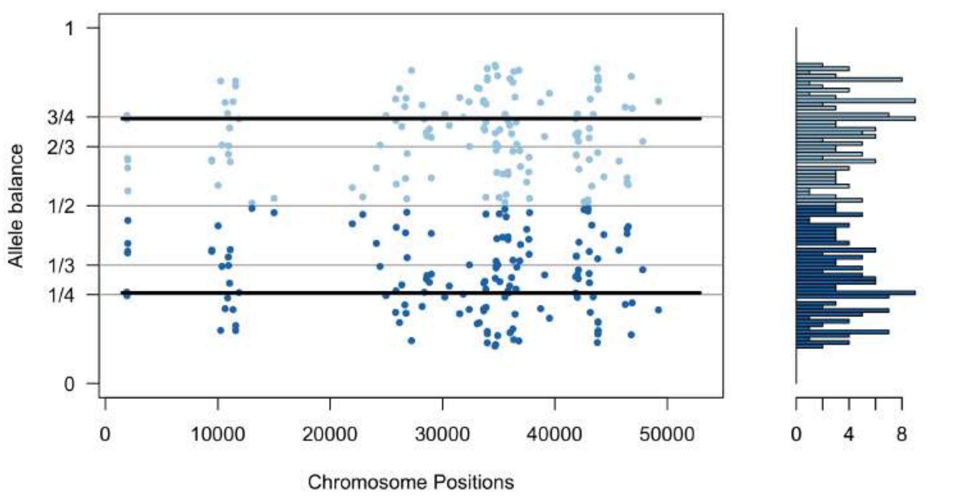


pit10_scaffold65.vcf

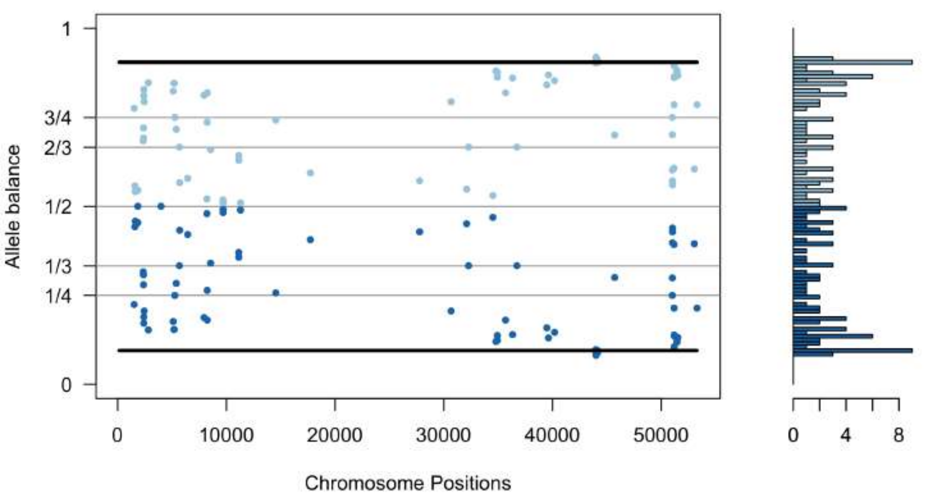


pit10_scaffold66.vcf

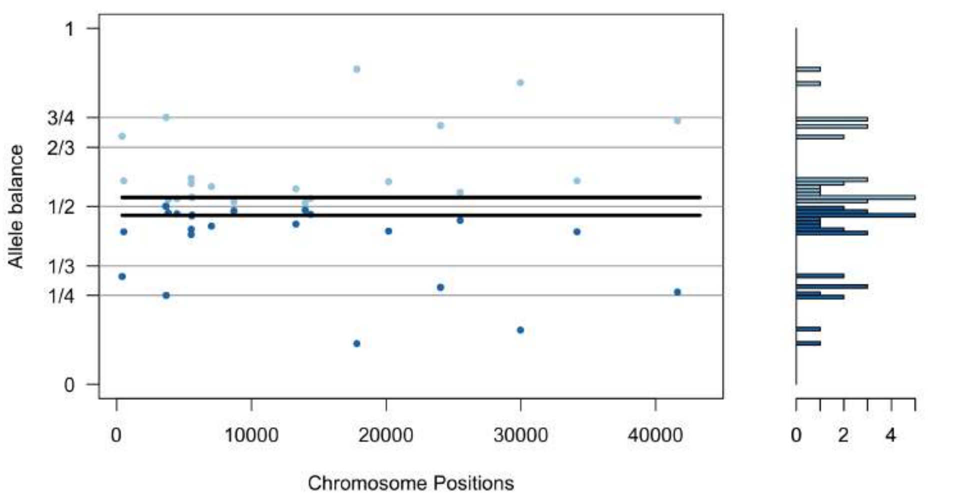


pit10_scaffold67.vcf

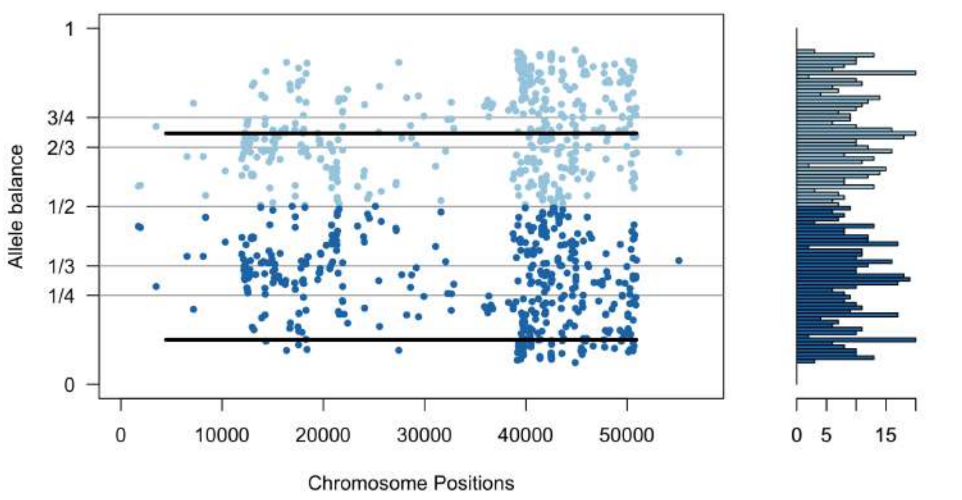


pit10_scaffold68.vcf

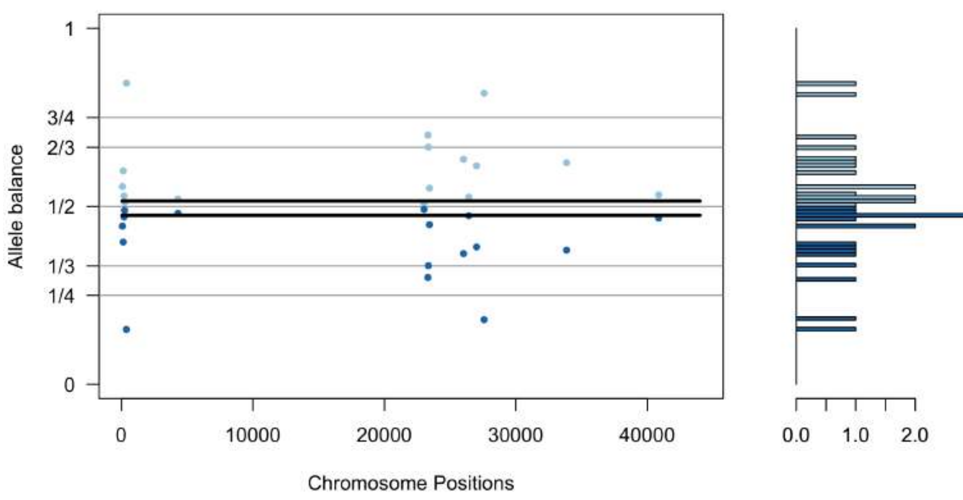


pit10_scaffold69.vcf

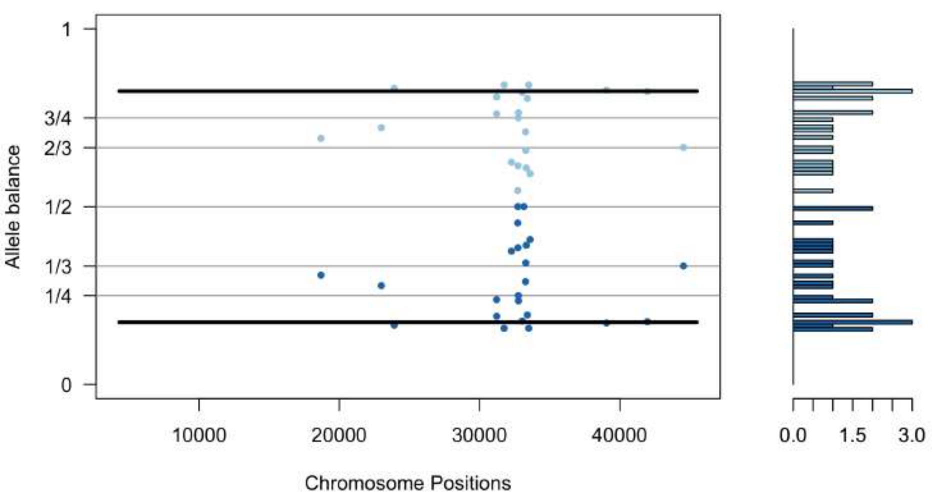


pit10_scaffold70.vcf

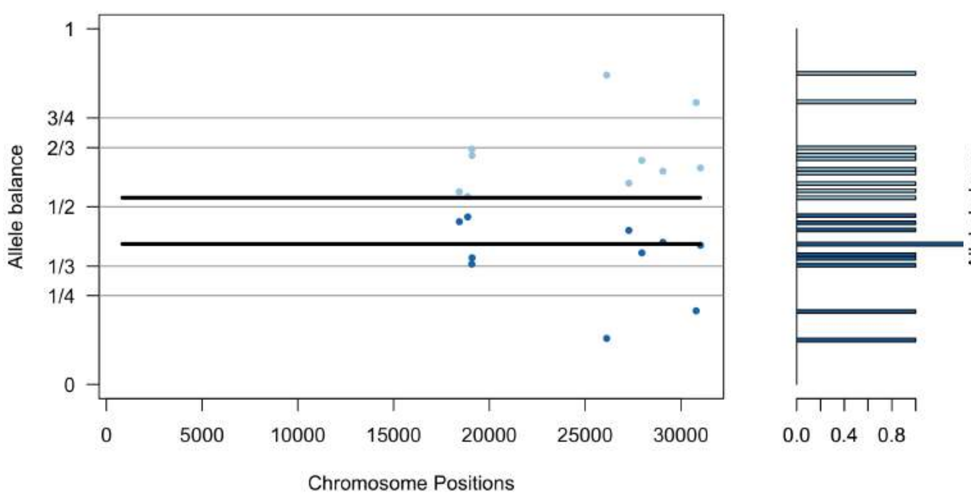


pit10_scaffold71.vcf

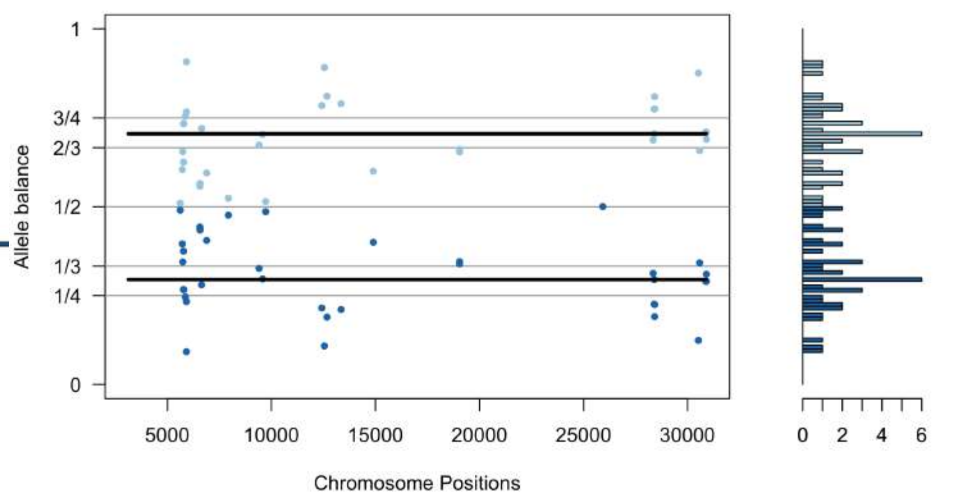


pit10_scaffold72.vcf

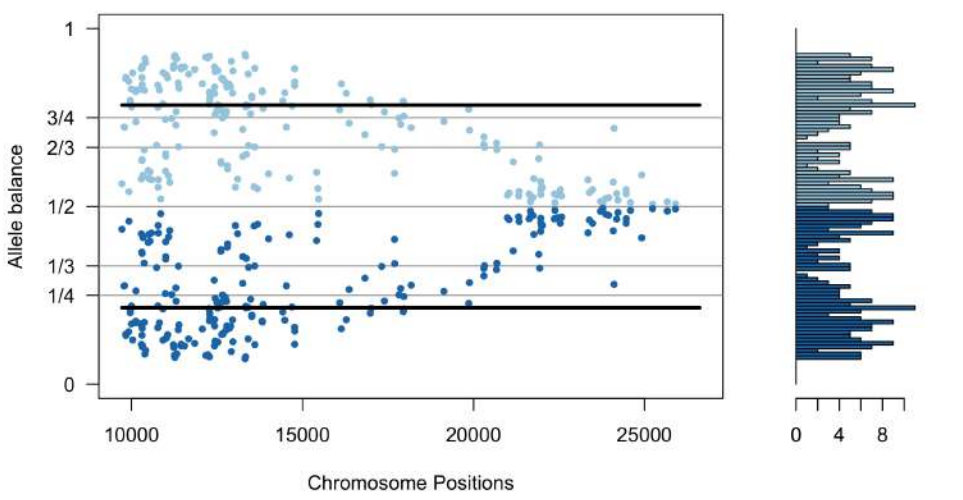


pit10_scaffold73.vcf

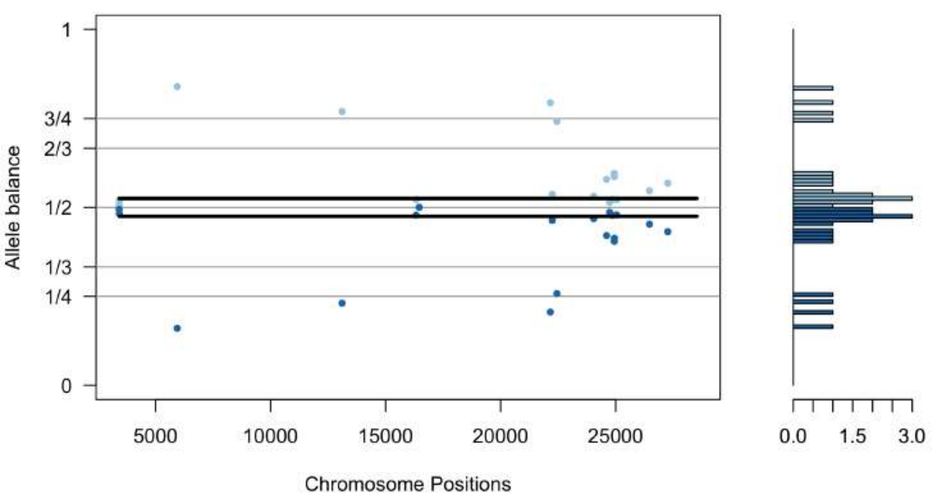


pit10_scaffold74.vcf

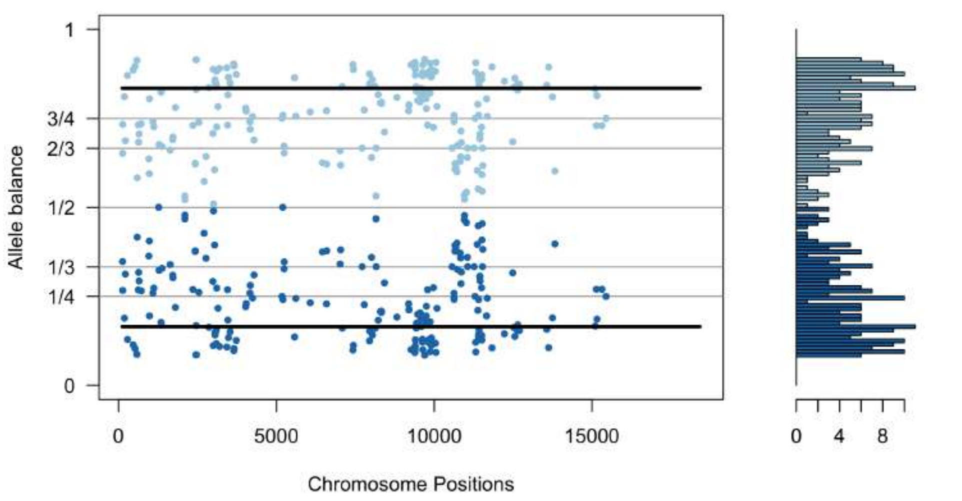


pit10_scaffold75.vcf

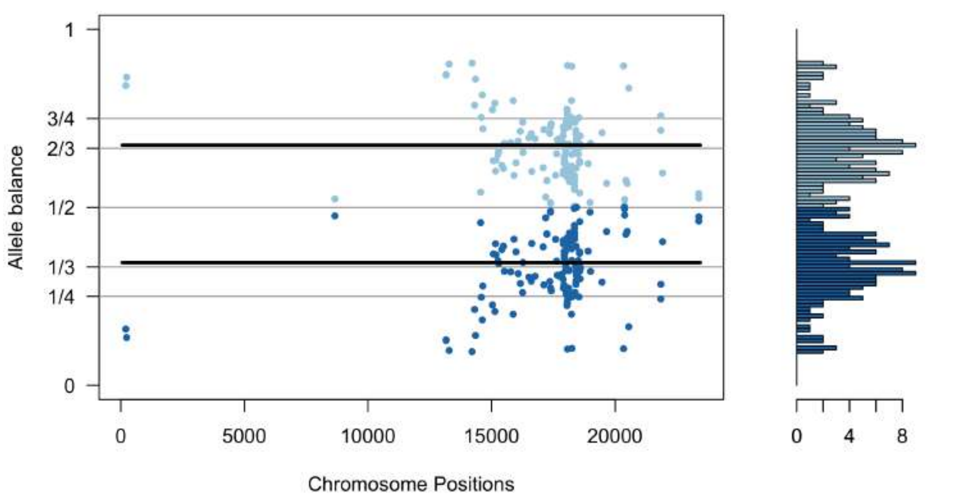


pit10_scaffold76.vcf

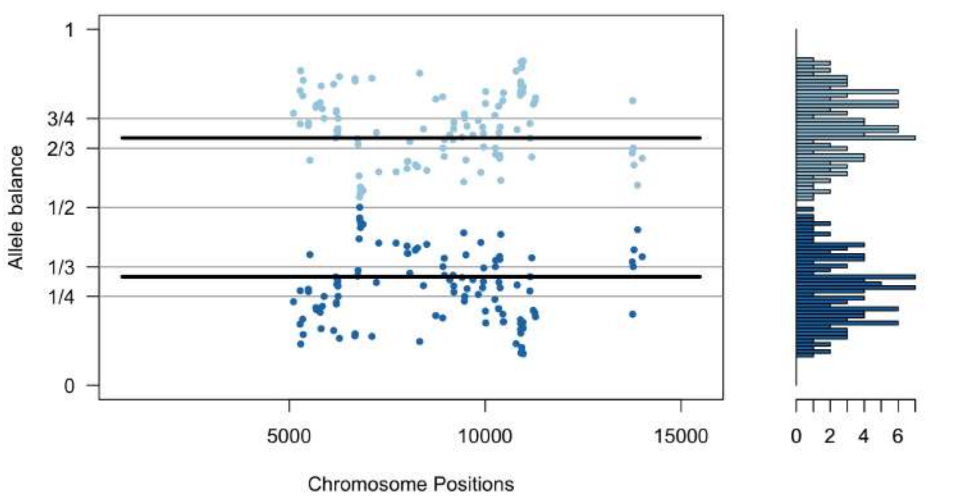


pit10_scaffold78.vcf

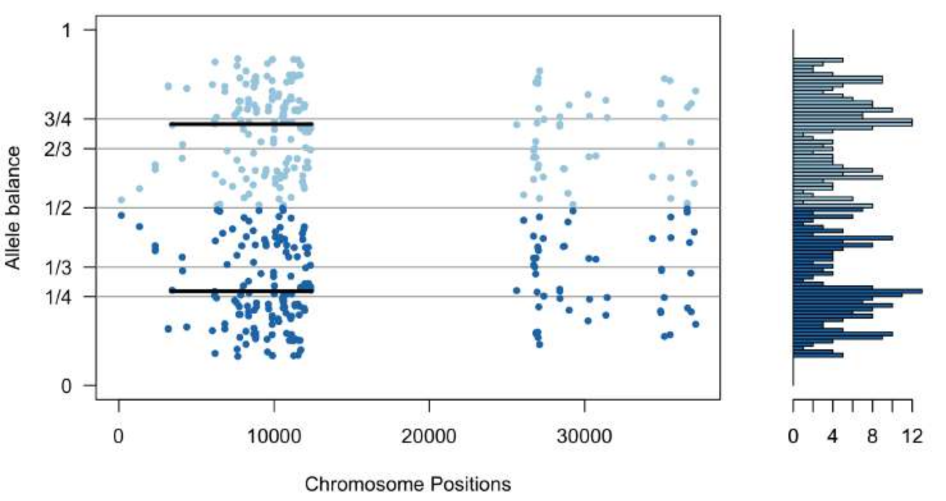


pit10_scaffold79.vcf

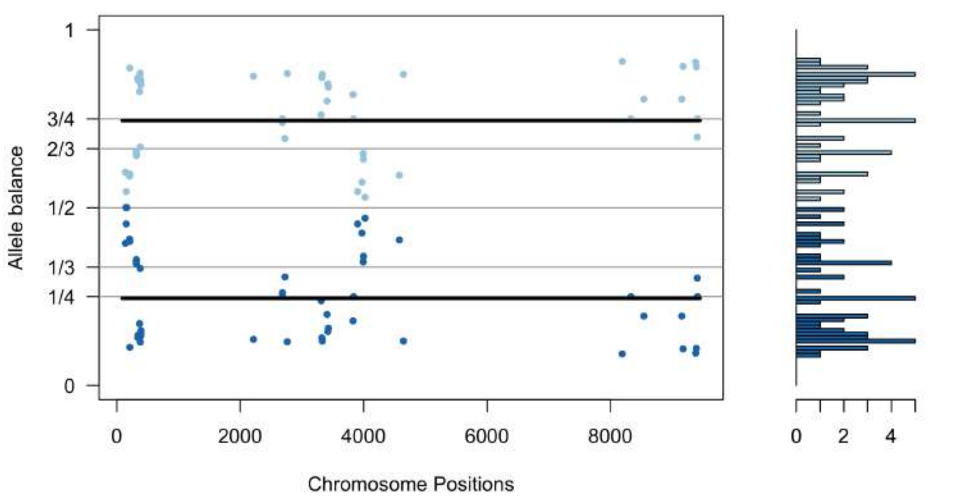


pit10_scaffold80.vcf

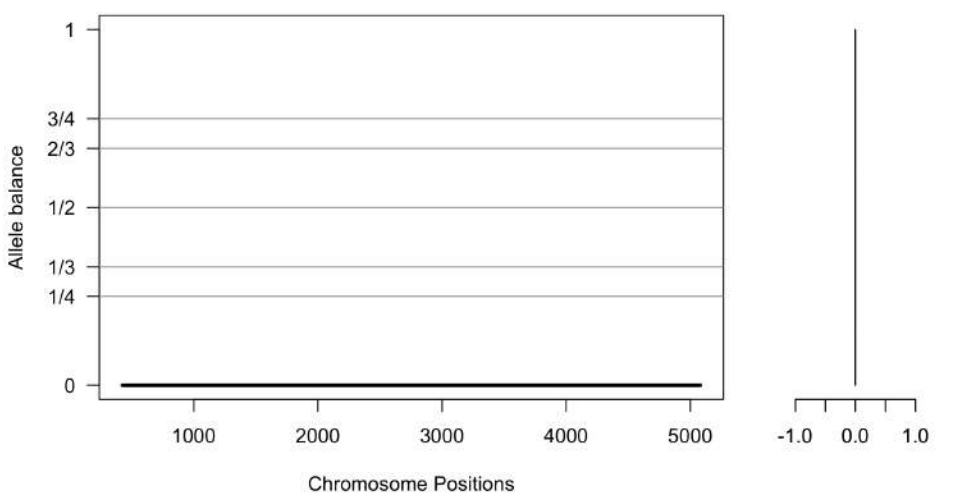


pit10_scaffold81.vcf

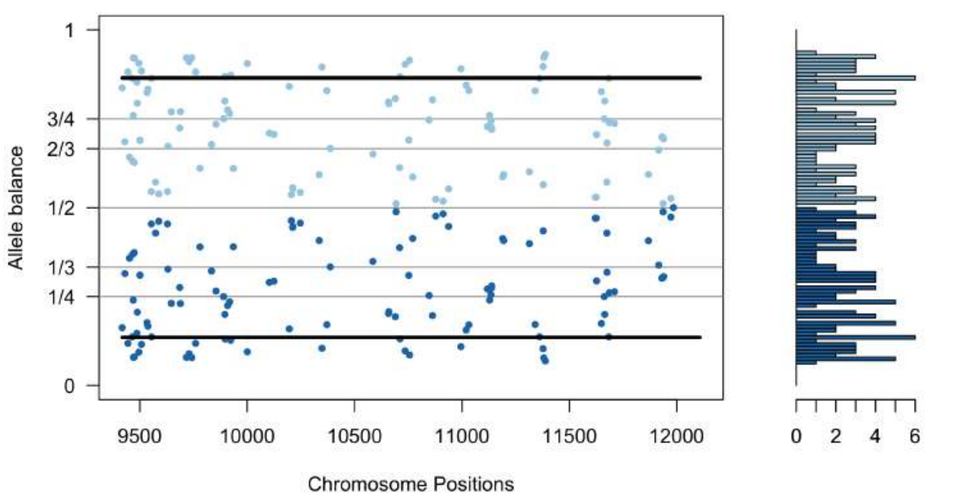

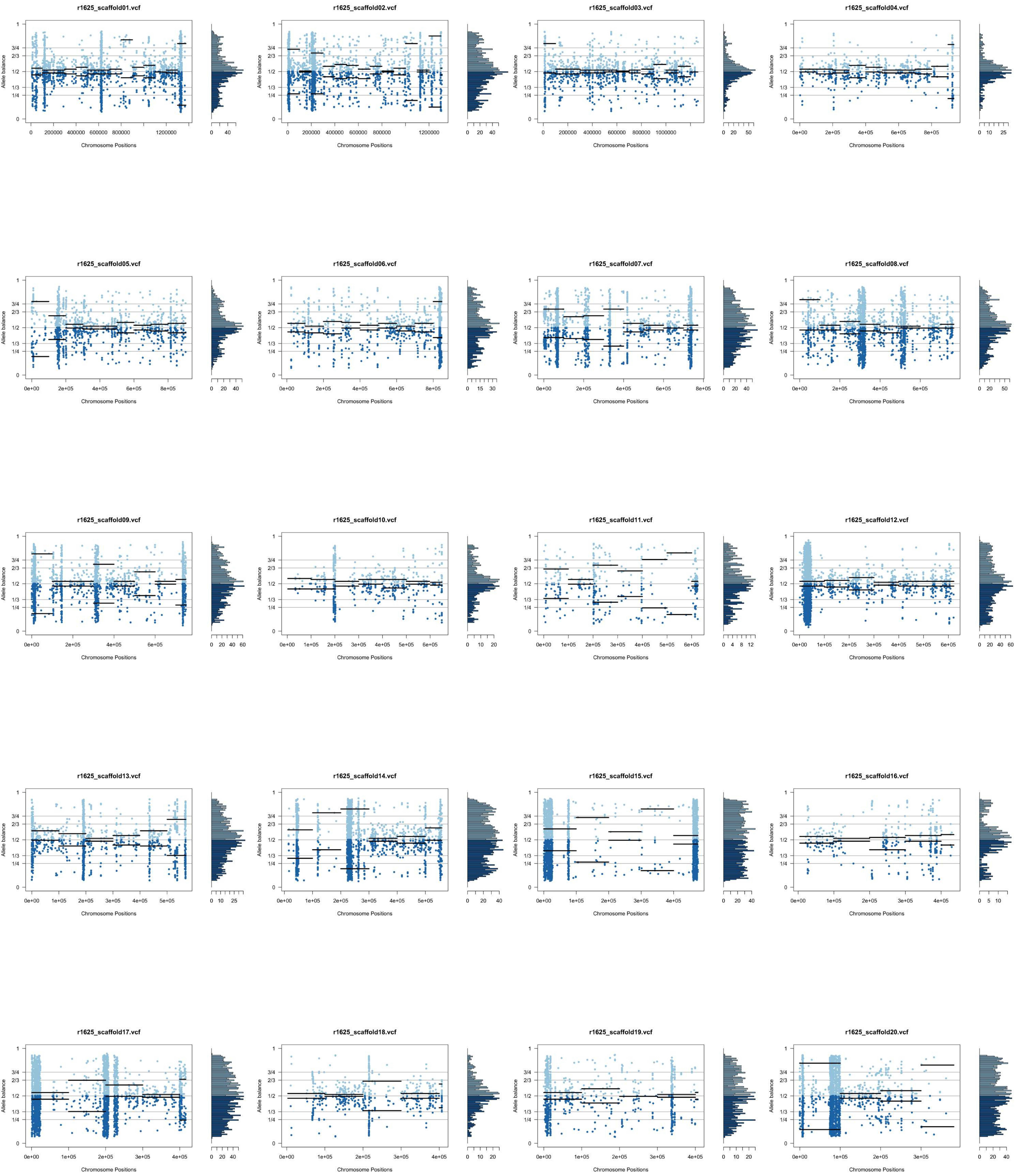
r1625_scaffold01.vcf
r1625_scaffold02.vcf
r1625_scaffold03.vcf
r1625_scaffold04.vcf
r1625_scaffold05.vcf
r1625_scaffold06.vcf
r1625_scaffold07.vcf
r1625_scaffold08.vcf
r1625_scaffold09.vcf
r1625_scaffold10.vcf
r1625_scaffold11.vcf
r1625_scaffold12.vcf
r1625_scaffold13.vcf
r1625_scaffold14.vcf
r1625_scaffold15.vcf
r1625_scaffold16.vcf
r1625_scaffold17.vcf
r1625_scaffold18.vcf
r1625_scaffold19.vcf
r1625_scaffold20.vcf
Allele balance
Chromosome Positions

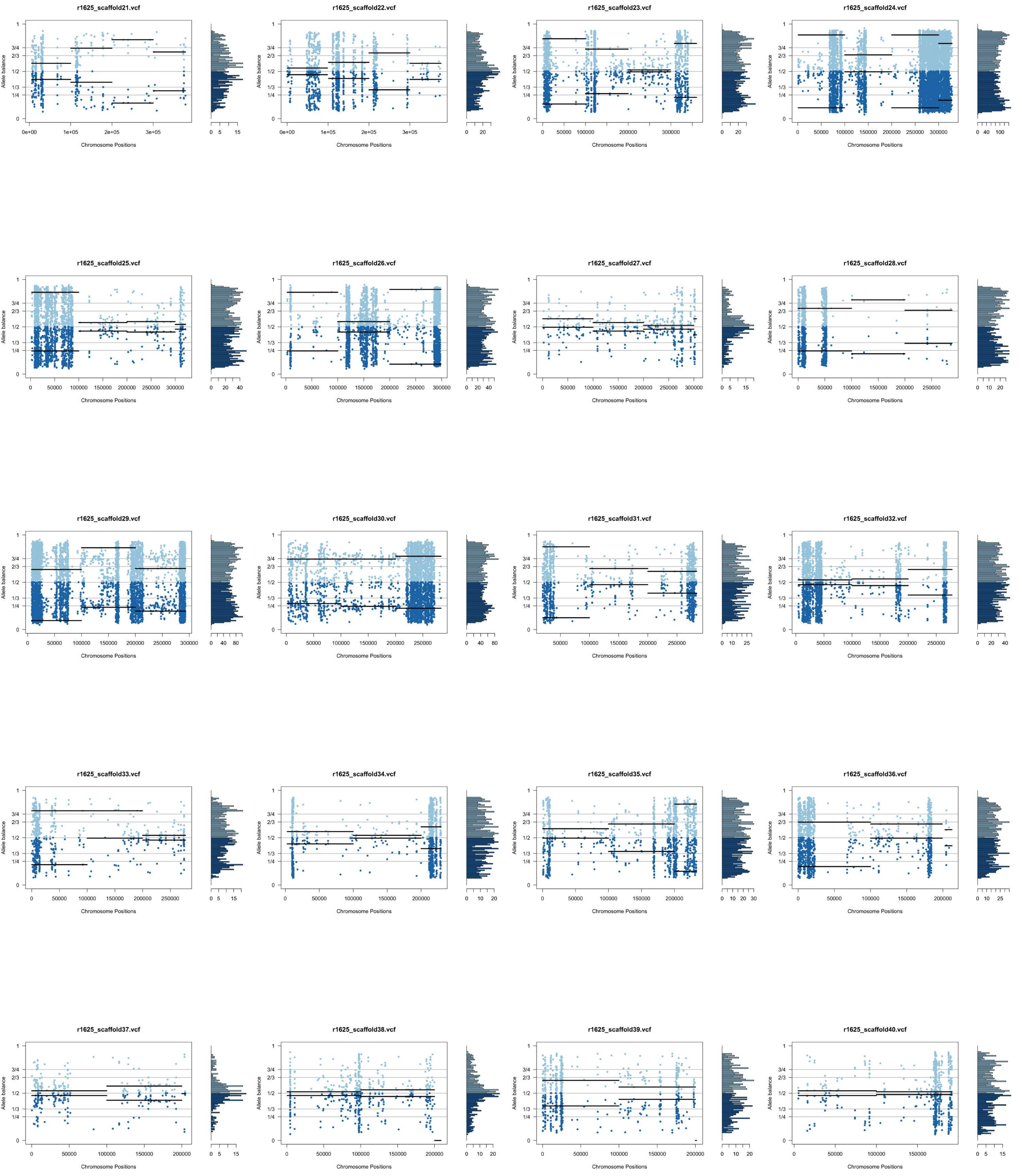
r1625_scaffold21.vcf
r1625_scaffold22.vcf
r1625_scaffold23.vcf
r1625_scaffold24.vcf
r1625_scaffold25.vcf
r1625_scaffold26.vcf
r1625_scaffold27.vcf
r1625_scaffold28.vcf
r1625_scaffold29.vcf
r1625_scaffold30.vcf
r1625_scaffold31.vcf
r1625_scaffold32.vcf
r1625_scaffold33.vcf
r1625_scaffold34.vcf
r1625_scaffold35.vcf
r1625_scaffold36.vcf
r1625_scaffold37.vcf
r1625_scaffold38.vcf
r1625_scaffold39.vcf
r1625_scaffold40.vcf
Allele balance
Chromosome Positions

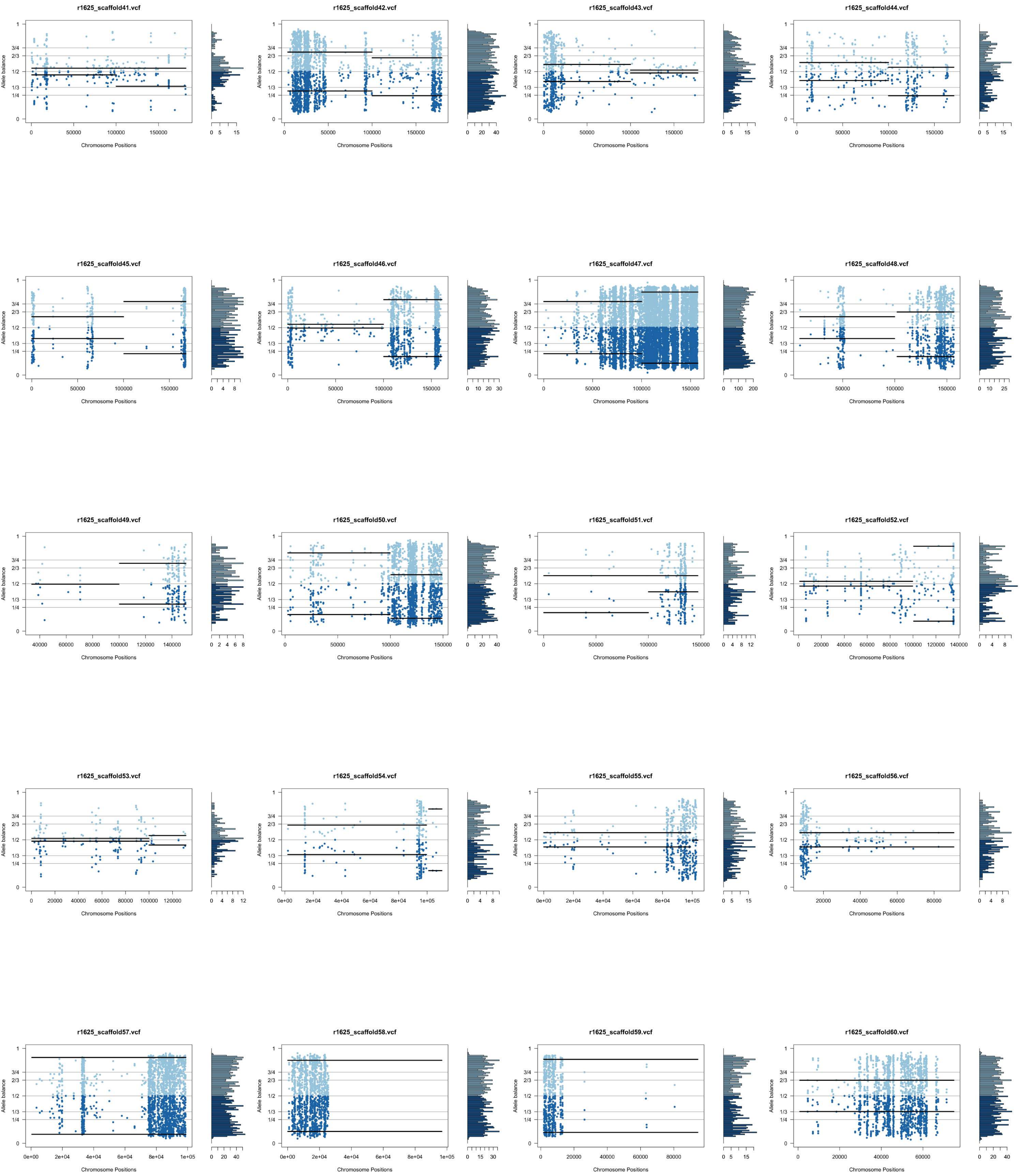
r1625_scaffold41.vcf
r1625_scaffold42.vcf
r1625_scaffold43.vcf
r1625_scaffold44.vcf
r1625_scaffold45.vcf
r1625_scaffold46.vcf
r1625_scaffold47.vcf
r1625_scaffold48.vcf
r1625_scaffold49.vcf
r1625_scaffold50.vcf
r1625_scaffold51.vcf
r1625_scaffold52.vcf
r1625_scaffold53.vcf
r1625_scaffold54.vcf
r1625_scaffold55.vcf
r1625_scaffold56.vcf
r1625_scaffold57.vcf
r1625_scaffold58.vcf
r1625_scaffold59.vcf
r1625_scaffold60.vcf
Allele balance
Chromosome Positions

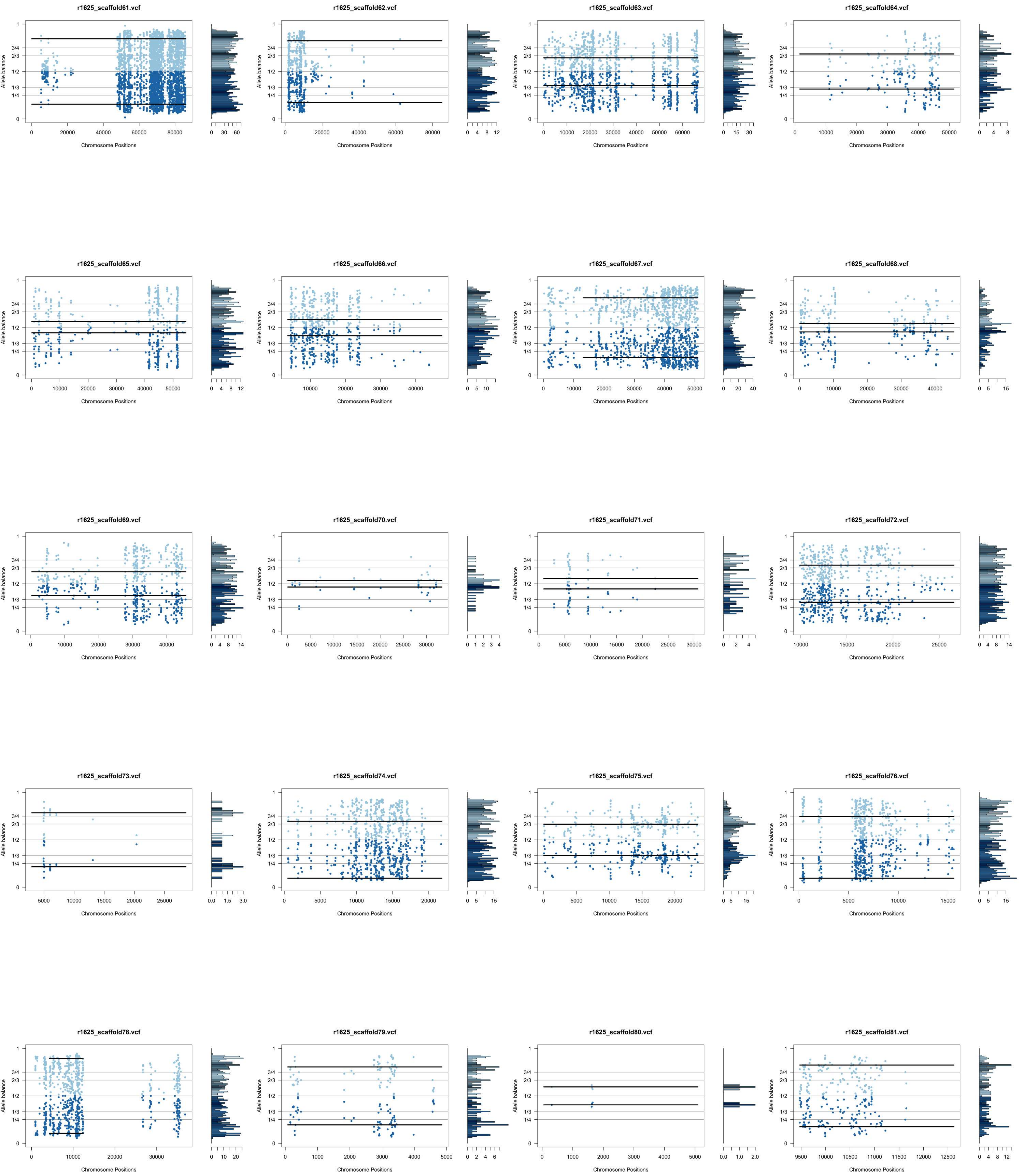
r1625_scaffold61.vcf
r1625_scaffold62.vcf
r1625_scaffold63.vcf
r1625_scaffold64.vcf
r1625_scaffold65.vcf
r1625_scaffold66.vcf
r1625_scaffold67.vcf
r1625_scaffold68.vcf
r1625_scaffold69.vcf
r1625_scaffold70.vcf
r1625_scaffold71.vcf
r1625_scaffold72.vcf
r1625_scaffold73.vcf
r1625_scaffold74.vcf
r1625_scaffold75.vcf
r1625_scaffold76.vcf
r1625_scaffold78.vcf
r1625_scaffold79.vcf
r1625_scaffold80.vcf
r1625_scaffold81.vcf
Allele balance
Chromosome Positions
1
3/4
2/3
1/2
1/3
1/4
0
