## Supplementary figure S7 for "Comprehensive genomic analysis of *Trypanosoma rangeli* reveals key insights into the biology and evolution of this non-virulent American mammalian trypanosome"

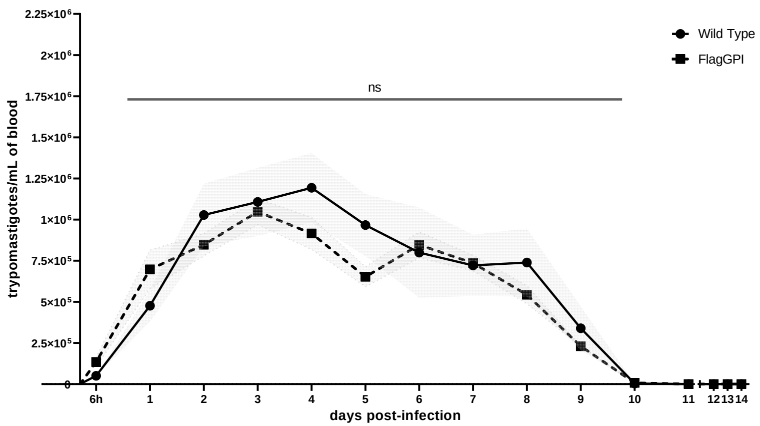

| Bonferroni's multiple comparisons test | Summary | Adjusted P Value |
|---|---|---|
| Wild Type vs. FlagGPI | ns | >0.9999 |
| Wild Type vs. *Tc*MUC I | * | 0.0499 |
| Wild Type vs. *Tc*MUC II | **** | <0.0001 |
| Wild Type vs. *Tr*MUC3 | **** | <0.0001 |
| Wild Type vs. *Tr*TASV-like1 | * | 0.0146 |

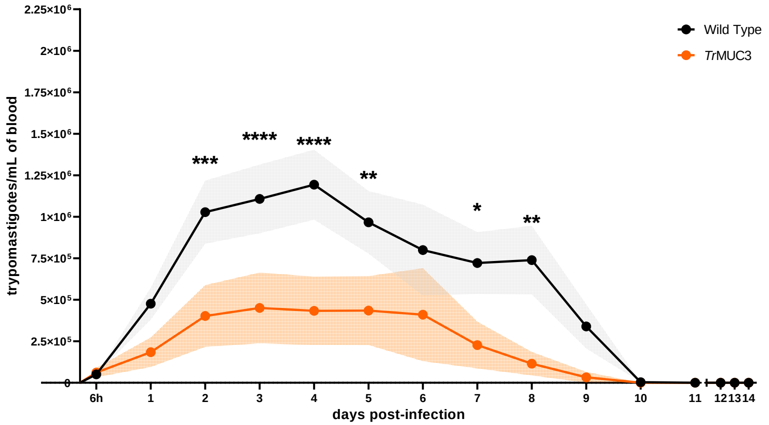

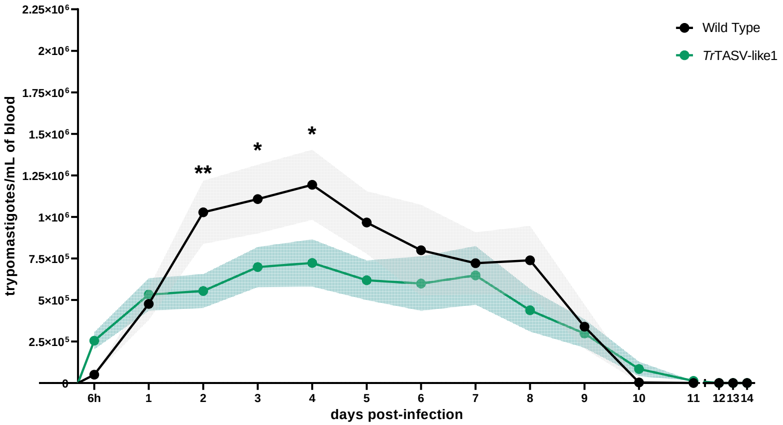

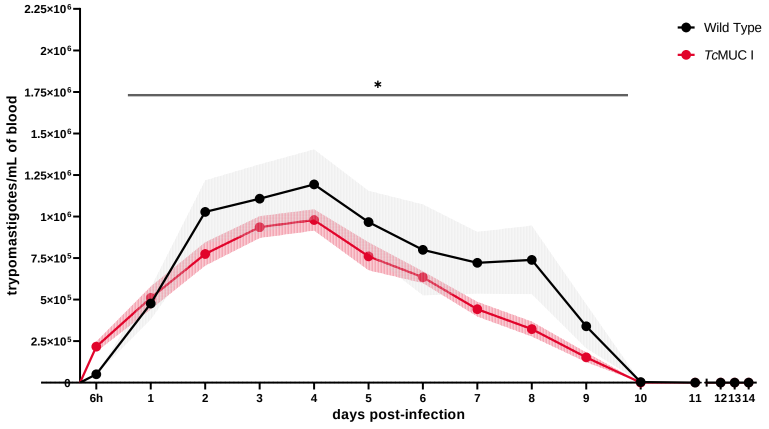

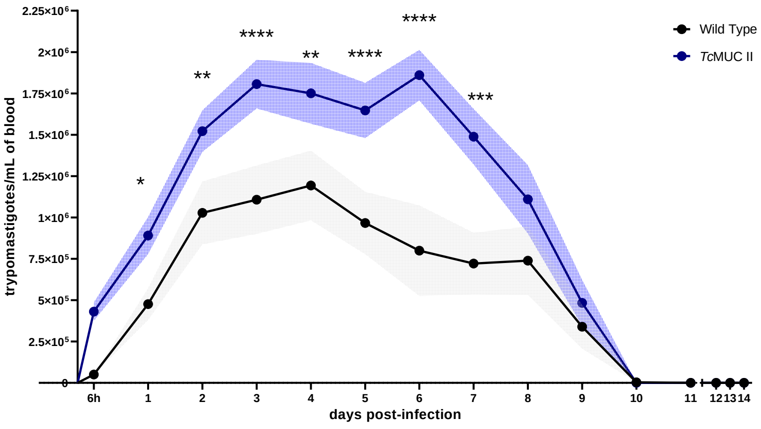
