## Supplementary table S1 for "Comprehensive genomic analysis of *Trypanosoma rangeli* reveals key insights into the biology and evolution of this non-virulent American mammalian trypanosome"

| Probe | | | | Genome location | |
|---|---|---|---|---|---|
| ID* | CDS annotation | Genomic coordinates (Strand)** | Size (bp) | *T. cruzi* CL Brener chromosome** | *T. rangeli* SC58 V.2 scaffolds |
| TcCLB.503793.20 | Phosphatidylinositol (3,5) kinase, putative (fragment) | 379,142 .. 380,494 (+) | 510 | 39-S | 16 |
| TcCLB.504109.170 | Hypothetical protein, conserved | 10,454 .. 11,866 (-) | 334 | 39-P | 16 |
| TcCLB.506127.90 | Hereditary spastic paraplegia protein strumpellin, putative | 612,942 .. 616,493 (+) | 373 | 39-S | 2 |
| TcCLB.507009.90 | Iron-sulfur cluster assembly protein, putative | 1,325,049 .. 1,325,540 (+) | 482 | 39-S | 10 |
| TcCLB.508465.120 | Syntaxin, putative | 1,536,810 .. 1,537,718 (+) | 763 | 39-S | 10 |
| TcCLB.507713.30 | Heat shock protein 85, putative | 793,891 .. 796,005 (-) | 479 | 37-S | 5 |
| TcCLB.507641.120 | ATP-dependent DEAD/H RNA helicase, putative | 1,167,741 .. 1,169,567 (-) | 693 | 37-P | 6 |
| TcCLB.507609.40 | Delta-4 fatty acid desaturase, putative | 1,316,342 .. 1,317,616 (+) | 366 | 37-P | 18 |
| TcCLB.511041.40 | Hexose transporter, putative | 292,428 .. 294,062 (+) | 678 | 37-S | Multiple |
| TcCLB.510129.20 | Surface membrane protein | 870,126 .. 871,505 (-) | 619 | 37-P | 5, 14 |
| TcCLB.507681.160 | 40S ribosomal protein S24E, putative | 38,962 .. 39,375 (-) | 380 | 4-P | 30 |
