## Supplementary table S2 for "Comprehensive genomic analysis of *Trypanosoma rangeli* reveals key insights into the biology and evolution of this non-virulent American mammalian trypanosome"

| Scaffold number | Scaffold size (bp) | Telomeric region position | |
|---|---|---|---|
| | | Start (bp) | End (bp) |
| 6 | 850,150 | 848,419 | 850,150 |
| 15 | 477,707 | 474,975 | 477,707 |
| 22 | 380,901 | 0 | 5,529 |
| 26 | 308,182 | 299,410 | 308,182 |
| 29 | 293,810 | 0 | 7,111 |
| 32 | 279,601 | 0 | 3,344 |
| 34 | 238,211 | 229,787 | 238,211 |
| 57 | 98,993 | 0 | 4,261 |
| 91 | 28,961 | 0 | 9,699 |
| 324 | 12,696 | 0 | 9,385 |
