## Supplementary table S3 for "Comprehensive genomic analysis of *Trypanosoma rangeli* reveals key insights into the biology and evolution of this non-virulent American mammalian trypanosome"

| | | SC58 V2 genome position | | | | | | | Signal Peptide | | | | GPI Anchor | | | | N-glycosilation | | | Peptidase_M8 superfamily | | | | Surfaceome MS/MS | | | | | | | Secretome MS/MS | | | | | |
|---|---|---|---|---|---|---|---|---|---|---|---|---|---|---|---|---|---|---|---|---|---|---|---|---|---|---|---|---|---|---|---|---|---|---|---|---|
| ID (Access code) | GP63 Group | Status | Tandem organization | Scaffold | Start | End | Strand | Seq-length (aa) | Signal P | Likelihood | DeepTMHMM | Clivage site position and probability | Pred. GPI-Anchored | ω-site Position | ω-residue anchor | Likelihood | residues | % | Core length (aa) | Domain status | Active site motif | Identity (e-value) | Identity (bitscore) | Protein Probability | Top Peptide Probability | Combined Total Peptides | Combined Spectral Count | Epi Spectral Count | Trypo Spectral Count | Indistinguishable Proteins | Coverage | Protein Probability | Top Peptide Probability | Total Peptides | Unique Peptides | Indistinguishable Proteins |
| TrSC58v2_1328_HEIAH TrGP63 | 12 | gene | Tandem | 3 | 610.368 | 611.969 | plus | 533 | Other | 1,000 | SP | 10 | Not GPI-Anchored | - | * | 0,992 | 1 | 0,19 | 523 | Complete | HEIAHALG(60)HW(9)M | 1,83E-130 | 391,571 | | | | | | | | 2,44 | 0,9836 | 0,9977 | 1 | 0 | g1329_HEIAH, g1330_HEIAH, g1331_HEIAH, g1332_HEIAH, g1333_HEIAH, g1334_HEIAH, g1336 |
| TrSC58v2_1329_HEIAH TrGP63 | 12 | gene | Tandem | 3 | 612.934 | 614.535 | plus | 533 | Other | 1,000 | SP | 10 | Not GPI-Anchored | - | * | 0,992 | 1 | 0,19 | 523 | Complete | HEIAHALG(60)HW(9)M | 1,83E-130 | 391,571 | | | | | | | | | | | | | |
| TrSC58v2_1330_HEIAH TrGP63 | 12 | gene | Tandem | 3 | 615.497 | 617.098 | plus | 533 | Other | 1,000 | SP | 10 | Not GPI-Anchored | - | * | 0,992 | 1 | 0,19 | 523 | Complete | HEIAHALG(60)HW(9)M | 1,83E-130 | 391,571 | | | | | | | | | | | | | |
| TrSC58v2_1331_HEIAH TrGP63 | 12 | gene | Tandem | 3 | 618.062 | 619.663 | plus | 533 | Other | 1,000 | SP | 10 | Not GPI-Anchored | - | * | 0,992 | 1 | 0,19 | 523 | Complete | HEIAHALG(60)HW(9)M | 1,83E-130 | 391,571 | | | | | | | | | | | | | |
| TrSC58v2_1332_HEIAH TrGP63 | 12 | gene | Tandem | 3 | 620.627 | 622.228 | plus | 533 | Other | 1,000 | SP | 10 | Not GPI-Anchored | - | * | 0,992 | 1 | 0,19 | 523 | Complete | HEIAHALG(60)HW(9)M | 1,83E-130 | 391,571 | | | | | | | | | | | | | |
| TrSC58v2_1333_HEIAH TrGP63 | 12 | gene | Tandem | 3 | 623.193 | 624.794 | plus | 533 | Other | 1,000 | SP | 10 | Not GPI-Anchored | - | * | 0,992 | 1 | 0,19 | 523 | Complete | HEIAHALG(60)HW(9)M | 1,81E-130 | 391,571 | | | | | | | | | | | | | |
| TrSC58v2_1334_HEIAH TrGP63 | 12 | gene | Tandem | 3 | 625.758 | 627.359 | plus | 533 | Other | 1,000 | SP | 10 | Not GPI-Anchored | - | * | 0,992 | 1 | 0,19 | 523 | Complete | HEIAHALG(60)HW(9)M | 1,83E-130 | 391,571 | | | | | | | | | | | | | |
| TrSC58v2_1339_HEIAH TrGP63 | 11 | gene | Tandem | 3 | 633.452 | 635.194 | plus | 580 | Other | 1,000 | SP | 10 | GPI-Anchored | 555 | S | 0,44 | 5 | 0,92 | 545 | Complete | HEIAHALG(60)HW(9)M | 7,74E-136 | 406,979 | 1 | 0,999 | 2 | 3 | 0 | 3 | | 13,1 | 10.000 | 0,999 | 4 | 0 | |
| TrSC58v2_2586_HEIAH TrGP63 | 6 | gene | | 6 | 441.240 | 442.943 | minus | 567 | SP | 1,000 | SP | 21-22 / 0.8063 | GPI-Anchored | 532 | G | 0,417 | 2 | 0,39 | 511 | Complete | HEIAHTLG(61)HW(9)M | 2,29E-176 | 510,212 | | | | | | | | | | | | | |
| TrSC58v2_2748_HEIAH TrGP63 | 6 | gene | | 7 | 66.356 | 67.903 | minus | 515 | SP | 1,000 | SP | 22-23 / 0.6551 | Not GPI-Anchored | - | * | 0,994 | 4 | 0,81 | 493 | Complete | HEIAHVLG(61)HW(9)M | 2,55E-172 | 497,886 | | | | | | | | | | | | | |
| TrSC58v2_2794_HEIAH TrGP63 | 12 | gene | Tandem | 7 | 204.272 | 206.194 | minus | 640 | Other | 1,000 | SP | 10 | GPI-Anchored | 616 | C | 0,51 | 4 | 0,66 | 606 | Complete | HEIAHALG(60)HW(9)M | 2,55E-127 | 387,333 | | | | | | | | 10,16 | 1 | 0,9813 | 2 | 0 | |
| TrSC58v2_3271_HAIAH TrGP63 | 10 | gene | | 8 | 503.482 | 505.512 | minus | 676 | Other | 1,000 | SP | 10 | GPI-Anchored | 650 | D | 0,279 | 3 | 0,47 | 640 | Complete | HAIAHALG(57)HLW(9)M | 7,45E-115 | 354,927 | | | | | | | | | | | | | |
| TrSC58v2_3280_HAIAH TrGP63 | 10 | gene | | 8 | 515.982 | 517.667 | minus | 561 | Other | 1,000 | SP | 10 | GPI-Anchored | 535 | D | 0,279 | 4 | 0,76 | 525 | Complete | HAIAHALG(57)HLW(9)M | 4,60E-118 | 359,549 | | | | | | | | 7,66 | 0,99 | 0,9986 | 1 | 1 | |
| TrSC58v2_3991_HEIVH TrGP63 | 7 | gene | | 11 | 12.100 | 13.704 | minus | 534 | Other | 1,000 | SP | 10 | GPI-Anchored | 513 | S | 0,582 | 3 | 0,6 | 503 | Complete | HEIVHILG(61)HW(9)M | 0 | 529,857 | | | | | | | | 11,8 | 0,9958 | 0,999 | 2 | 2 | |
| TrSC58v2_4259_HEIAH TrGP63 | 9 | gene | | 12 | 28.539 | 31.058 | minus | 840 | SP | 0,666 | SP | 17-18 / 0.6196 | GPI-Anchored | 814 | S | 0,239 | 5 | 0,63 | 797 | Complete | HEIAHALG(62)HW(9)M | 5,27E-109 | 344,527 | | | | | | | | | | | | | |
| TrSC58v2_4583_HEIAH TrGP63 | 5 | gene | | 13 | 216.252 | 218.276 | plus | 674 | SP | 1,000 | SP | 20-21 / 0.8840 | Not GPI-Anchored | - | * | 0,82 | 0 | 0 | 654 | Complete | HEIAHALG(62)HW(9)M | 1,02E-163 | 482,092 | | | | | | | | | | | | | |
| TrSC58v2_4641_HEIGH TrGP63 | 4 | gene | Tandem | 13 | 398.551 | 400.848 | plus | 765 | Other | 1,000 | SP | 10 | GPI-Anchored | 742 | S | 0,565 | 2 | 0,27 | 732 | Complete | HEIGHILG(61)HW(9)M | 0 | 668,914 | | | | | | | | | | | | | |
| TrSC58v2_4642_HEIGH TrGP63 | 4 | gene | Tandem | 13 | 401.627 | 403.774 | plus | 715 | Other | 1,000 | SP | 10 | GPI-Anchored | 692 | S | 0,565 | 2 | 0,29 | 682 | Complete | HEIGHILG(61)HW(9)M | 0 | 676,233 | | | | | | | | | | | | | |
| TrSC58v2_4649_HEIGH TrGP63 | 4 | gene | Tandem | 13 | 412.966 | 414.888 | plus | 640 | Other | 1,000 | SP | 10 | GPI-Anchored | 617 | S | 0,565 | 2 | 0,33 | 607 | Complete | HEIGHILG(61)HW(9)M | 0 | 678,159 | | | | | | | | | | | | | |
| TrSC58v2_4652_HEIGH TrGP63 | 4 | gene | Tandem | 13 | 418.173 | 420.530 | plus | 785 | Other | 1,000 | SP | 10 | Not GPI-Anchored | - | * | 0,488 | 2 | 0,26 | 775 | Complete | HEIGHILG(61)HW(9)M | 0 | 612,675 | | | | | | | | | | | | | |
| TrSC58v2_4653_HEIGH TrGP63 | 4 | gene | Tandem | 13 | 421.453 | 422.565 | plus | 370 | Other | 1,000 | GLOB | - | Not GPI-Anchored | - | * | 0,99 | 2 | BNP | - | Complete | HEIGHILG(61)HW(9)M | 2,17E-160 | 461,677 | | | | | | | | | | | | | |
| TrSC58v2_4654_HEIGH TrGP63 | 4 | gene | Tandem | 13 | 423.888 | 426.260 | plus | 790 | Other | 1,000 | SP | 10 | GPI-Anchored | 767 | S | 0,565 | 2 | 0,26 | 757 | Complete | HEIGHILG(61)HW(9)M | 0 | 671,225 | | | | | | | | | | | | | |
| TrSC58v2_4655_HEIGH TrGP63 | 4 | gene | Tandem | 13 | 426.921 | 428.546 | plus | 541 | Other | 1,000 | SP | 10 | Not GPI-Anchored | - | * | 0,995 | 3 | 0,56 | 531 | Complete | HEIGHILG(61)HW(9)M | 0 | 622,305 | | | | | | | | | | | | | |
| TrSC58v2_4656_HEIGH TrGP63 | 4 | gene | Tandem | 13 | 430.236 | 432.008 | plus | 590 | Other | 1,000 | GLOB | - | Not GPI-Anchored | - | * | 0,491 | 2 | BNP | - | Complete | HEIGHILG(61)HW(9)M | 0 | 522,924 | | | | | | | | | | | | | |
| TrSC58v2_4723_HEIAH TrGP63 | 6 | gene | | 14 | 44.040 | 45.587 | minus | 515 | SP | 1,000 | SP | 22-23 / 0.6523 | Not GPI-Anchored | - | * | 0,995 | 3 | 0,61 | 493 | Complete | HEIAHALG(61)HW(9)M | 3,48E-174 | 501,303 | | | | | | | | 6,99 | 0,8773 | 0,981 | 1 | 1 | |
| TrSC58v2_4795_HEIAH TrGP63 | 4 | gene | | 14 | 228.343 | 230.373 | plus | 676 | SP | 0,993 | SP | 42-43 / 0.9702 | GPI-Anchored | 653 | S | 0,21 | 7 | 1,15 | 611 | Complete | HEIAHALG(61)HW(9)M | 0 | 658,129 | 0,9863 | 0,999 | 1 | 1 | 0 | 1 | | 9,76 | 0,9944 | 0,9942 | 2 | 2 | |
| TrSC58v2_4835_HEMAH TrGP63 | 3 | gene | | 14 | 318.266 | 320.434 | minus | 722 | SP | 0,999 | SP | 22-23 / 0.8854 | GPI-Anchored | 699 | S | 0,53 | 2 | 0,3 | 677 | Complete | HEMAHALG(234)HW(9)M | 3,94E-117 | 362,246 | 1,000 | 0,999 | 3 | 18 | 0 | 18 | g4845 | | | | | | |
| TrSC58v2_4929_HEIGH TrGP63 | 4 | gene | | 15 | 20.504 | 22.453 | minus | 649 | Other | 1,000 | GLOB | - | GPI-Anchored | 624 | D | 0,376 | 6 | BNP | - | Complete | HEIGHALG(61)HW(9)M | 0 | 599,578 | | | | | | | | | | | | | |
| TrSC58v2_5263_HEIAH TrGP63 | 9 | gene | Tandem | 16 | 370.825 | 372.678 | plus | 617 | Other | 1,000 | GLOB | - | GPI-Anchored | 589 | D | 0,239 | 4 | BNP | - | Complete | HEIAHALG(62)HW(9)M | 1,69E-109 | 339,134 | 1,000 | 0,999 | 4 | 45 | 2 | 43 | g5266 | | | | | | |
| TrSC58v2_5266_HEIAH TrGP63 | 9 | gene | Tandem | 16 | 376.036 | 377.892 | plus | 618 | Other | 1,000 | GLOB | - | GPI-Anchored | 590 | D | 0,239 | 4 | BNP | - | Complete | HEIAHALG(62)HW(9)M | 4,97E-110 | 340,675 | | | | | | | | | | | | | |
| TrSC58v2_5291_HEIGH TrGP63 | 4 | gene | | 17 | 11.190 | 13.448 | minus | 752 | SP | 0,747 | SP | 42-43 / 0.8060 | GPI-Anchored | 727 | D | 0,224 | 6 | 0,88 | 685 | Complete | HEIGHILG(61)HW(9)M | 0 | 630,394 | | | | | | | | | | | | | |
| TrSC58v2_559_HELAH TrGP63 | 2 | gene | | 1 | 1.330.975 | 1.332.531 | plus | 576 | SP | 1,000 | SP | 33-34 / 0.7894 | Not GPI-Anchored | - | * | 0,995 | 0 | 0 | 543 | Complete | HELAHVLG(62)HW(9)M | 4,96E-112 | 344,527 | | | | | | | | | | | | | |
| TrSC58v2_560_RGLAY TrGP63 | 11 | gene | | 1 | 1.333.555 | 1.335.684 | plus | 709 | Other | 1,000 | SP | 11 | GPI-Anchored | 685 | A | 0,203 | 5 | 0,74 | 674 | Complete | RGLAYALG(61)HW(9)M | 1,45E-121 | 373,417 | 1,000 | 0,999 | 7 | 170 | 73 | 97 | | 6,63 | 0,9917 | 0,999 | 2 | 1 | |
| TrSC58v2_6130_HEMTH TrGP63 | 11 | gene | Tandem | 22 | 124.862 | 127.042 | minus | 726 | Other | 1,000 | SP | 10 | GPI-Anchored | 697 | S | 0,222 | 3 | 0,44 | 687 | Complete | HEMTHALG(61)HW(9)M | 4,75E-124 | 381,555 | | | | | | | | | | | | | |
| TrSC58v2_6131_HEIAH TrGP63 | 12 | gene | Tandem | 22 | 127.933 | 129.675 | minus | 580 | Other | 1,000 | SP | 10 | GPI-Anchored | 555 | S | 0,516 | 3 | 0,55 | 545 | Complete | HEIAHALG(60)HW(9)M | 1,14E-124 | 378,474 | | | | | | | | | | | | | |
| TrSC58v2_6132_HEIAH TrGP63 | 12 | gene | Tandem | 22 | 130.540 | 132.282 | minus | 580 | Other | 1,000 | SP | 10 | GPI-Anchored | 555 | S | 0,516 | 3 | 0,55 | 545 | Complete | HEIAHALG(60)HW(9)M | 1,14E-124 | 378,474 | | | | | | | | | | | | | |
| TrSC58v2_6133_HEIAH TrGP63 | 12 | gene | Tandem | 22 | 133.171 | 134.913 | minus | 580 | Other | 1,000 | SP | 10 | GPI-Anchored | 555 | S | 0,516 | 3 | 0,55 | 545 | Complete | HEIAHALG(60)HW(9)M | 1,14E-124 | 378,474 | | | | | | | | | | | | | |
| TrSC58v2_6284_HEIAH TrGP63 | 6 | gene | | 23 | 121.042 | 122.586 | minus | 514 | SP | 1,000 | SP | 21-22 / 0.3337 | Not GPI-Anchored | - | * | 0,994 | 3 | 0,61 | 493 | Complete | HEIAHALG(61)HW(9)M | 0 | 521,768 | | | | | | | | | | | | | |
| TrSC58v2_631_HEIGH TrGP63 | 4 | gene | | 2 | 188.862 | 190.814 | plus | 650 | SP | 0,997 | SP | 42-43 / 0.6467 | GPI-Anchored | 625 | D | 0,394 | 6 | 1,03 | 583 | Complete | HEIGHVLG(61)HW(9)M | 0 | 661,981 | | | | | | | | 6,31 | 0,9657 | 0,9951 | 1 | 0 | g632_HEIGH, g633_HEIGH |
| TrSC58v2_632_HEIGH TrGP63 | 4 | gene | | 2 | 192.778 | 194.730 | plus | 650 | SP | 0,999 | SP | 42-43 / 0.6478 | GPI-Anchored | 625 | D | 0,394 | 6 | 1,03 | 583 | Complete | HEIGHVLG(61)HW(9)M | 0 | 661,981 | | | | | | | | | | | | | |
| TrSC58v2_633_HEIGH TrGP63 | 4 | gene | | 2 | 196.689 | 198.641 | plus | 650 | SP | 0,999 | SP | 42-43 / 0.6478 | GPI-Anchored | 625 | D | 0,394 | 6 | 1,03 | 583 | Complete | HEIGHVLG(61)HW(9)M | 0 | 661,981 | | | | | | | | | | | | | |
| TrSC58v2_6472_HEIGH TrGP63 | 4 | gene | | 24 | 307.290 | 309.509 | plus | 739 | Other | 1,000 | SP | 10 | GPI-Anchored | 712 | S | 0,53 | 7 | 1 | 702 | Complete | HEIGHVLG(58)HW(9)M | 0 | 669,299 | | | | | | | | | | | | | |
| TrSC58v2_6495_HEMAH TrGP63 | 11 | gene | | 25 | 48.882 | 51.140 | minus | 752 | SP | 1,000 | SP | 30-31 / 0.6745 | GPI-Anchored | 724 | D | 0,228 | 5 | 0,72 | 694 | Complete | HEMAHALG(60)HW(9)M | 1,17E-120 | 372,261 | | | | | | | | | | | | | |
| TrSC58v2_6663_HEIAH TrGP63 | 9 | gene | | 26 | 116.516 | 119.014 | minus | 832 | SP | 1,000 | SP | 30-31 / 0.7669 | GPI-Anchored | 804 | S | 0,238 | 5 | 0,65 | 774 | Complete | HEIAHALG(62)HW(9)M | 1,37E-103 | 329,889 | | | | | | | | 12,62 | 1 | 0,9941 | 3 | 2 | |
| TrSC58v2_6853_HEMAH TrGP63 | 11 | gene | | 27 | 274.870 | 276.906 | minus | 678 | Other | 1,000 | SP | 11 | GPI-Anchored | 652 | S | 0,571 | 5 | 0,78 | 641 | Complete | HEMAHALG(61)HW(9)M | 2,35E-116 | 358,779 | | | | | | | | 9,59 | 1 | 0,999 | 3 | 0 | |
| TrSC58v2_6871_HEIAH TrGP63 | 6 | gene | | 28 | 10.839 | 12.386 | plus | 515 | SP | 1,000 | SP | 19-20 / 0.9505 | Not GPI-Anchored | - | * | 0,994 | 5 | 1,01 | 496 | Complete | HEIAHALG(61)HW(9)M | 1,36E-175 | 504,77 | | | | | | | | | | | | | |
| TrSC58v2_7126_HEIAH TrGP63 | 8 | gene | | 30 | 75.721 | 77.460 | minus | 579 | SP | 1,000 | SP | 26-27 / 0.9747 | Not GPI-Anchored | - | * | 0,985 | 4 | 0,72 | 553 | Complete | HEIAHALG(61)HW(9)M | 0 | 523,309 | | | | | | | | | | | | | |
| TrSC58v2_7196_HEIGH TrGP63 | 4 | gene | | 30 | 254.109 | 256.940 | plus | 943 | Other | 1,000 | SP | 10 | GPI-Anchored | 916 | S | 0,525 | 15 | 1,66 | 906 | Complete | HEIGHALG(61)HW(9)M | 0 | 653,506 | | | | | | | | | | | | | |
| TrSC58v2_7306_HEIAH TrGP63 | 4 | gene | | 32 | 16.864 | 18.747 | minus | 627 | SP | 0,742 | SP | 42-43 / 0.7679 | GPI-Anchored | 604 | S | 0,241 | 5 | 0,89 | 562 | Complete | HEIAHALG(61)HW(9)M | 0 | 676,233 | | | | | | | | | | | | | |
| TrSC58v2_7404_HGIAH TrGP63 | 6 | gene | | 32 | 262.834 | 264.378 | minus | 514 | SP | 1,000 | SP | 23-24 / 0.3171 | Not GPI-Anchored | - | * | 0,995 | 6 | 1,22 | 491 | Complete | HGIAHALG(61)HW(9)M | 1,97E-176 | 508,286 | | | | | | | | | | | | | |
| TrSC58v2_7524_HEIAH TrGP63 | 4 | gene | | 34 | 8.006 | 11.227 | minus | 1095 | Other | 1,000 | GLOB | - | GPI-Anchored | 1072 | S | 0,352 | 23 | BNP | - | Complete | HEIAHALG(61)HW(9)M | 0 | 610,749 | | | | | | | | | | | | | |
| TrSC58v2_7598_QKIGH TrGP63 | 4 | gene | | 34 | 217.444 | 220.527 | plus | 1027 | SP | 0,914 | SP | 42-43 / 0.9680 | GPI-Anchored | 1004 | S | 0,397 | 20 | 2,08 | 962 | Complete | QKIGHVLG(61)HW(9)M | 0 | 681,626 | | | | | | | | | | | | | |
| TrSC58v2_8159_HEIAH TrGP63 | 6 | gene | | 42 | 25.172 | 26.749 | plus | 525 | SP | 1,000 | SP | 23-24 / 0.5267 | Not GPI-Anchored | - | * | 0,989 | 4 | 0,8 | 502 | Complete | HEIAHALG(61)HW(9)M | 2,43E-176 | 508,671 | | | | | | | | | | | | | |
| TrSC58v2_8165_HAIAH TrGP63 | 10 | gene | | 42 | 42.954 | 44.636 | minus | 560 | Other | 0,670 | SP | 19 | GPI-Anchored | 536 | C | 0,34 | 3 | 0,58 | 517 | Complete | HAIAHALG56YLW(9)M | 1,64E-118 | 360,705 | 1,000 | 0,999 | 7 | 204 | 106 | 98 | | 7,32 | 1 | 0,999 | 3 | 2 | |
| TrSC58v2_8384_HELAH TrGP63 | 2 | gene | | 45 | 62.938 | 64.494 | minus | 518 | Other | 1,000 | GLOB | - | Not GPI-Anchored | - | * | 0,995 | 2 | BNP | - | Complete | HELAHVLG(62)HW(9)M | 1,08E-109 | 336,437 | | | | | | | | | | | | | |
| TrSC58v2_8509_HEIAH TrGP63 | 6 | gene | | 47 | 58.593 | 60.137 | minus | 514 | SP | 1,000 | SP | 21-22 / 0.4911 | Not GPI-Anchored | - | * | 0,994 | 4 | 0,81 | 493 | Complete | HEIAHVLG(61)HW(9)M | 6,03E-176 | 507,13 | | | | | | | | | | | | | |
| TrSC58v2_8644_HEMAH TrGP63 | 11 | gene | | 49 | 139.502 | 141.631 | plus | 709 | Other | 0,834 | SP | 11 | GPI-Anchored | 681 | D | 0,213 | 3 | 0,45 | 670 | Complete | HEMAHALG(60)HW(9)M | 4,31E-120 | 369,565 | | | | | | | | | | | | | |
| TrSC58v2_8766_RGLAY_TrGP63 | 11 | gene | | 51 | 134.134 | 136.071 | plus | 645 | Other | 1,000 | SP | 10 | GPI-Anchored | 616 | G | 0,286 | 4 | 0,66 | 606 | Complete | RGLAYALG(61)HW(9)M | 3,16E-124 | 378,039 | | | | | | | | | | | | | |
| TrSC58v2_8770_HEMAH TrGP63 | 11 | gene | | 52 | 8.304 | 10.115 | minus | 603 | Other | 1,000 | GLOB | - | GPI-Anchored | 577 | S | 0,564 | 5 | BNP | - | Complete | HEMAHALG(61)HW(9)M | 6,93E-116 | 355,312 | | | | | | | | | | | | | |
| TrSC58v2_8803_HEMAH TrGP63 | 11 | gene | | 52 | 87.363 | 89.276 | minus | 637 | Other | 1,000 | SP+TM | 10 | Not GPI-Anchored | - | * | 0,964 | 7 | 1,12 | 627 | Complete | HEMAHALG(61)HW(9)M | 9,29E-121 | 370,385 | | | | | | | | | | | | | |
| TrSC58v2_8920_HEMAH TrGP63 | 11 | gene | | 54 | 99.039 | 101.072 | minus | 677 | Other | 1,000 | SP | 11 | GPI-Anchored | 651 | S | 0,569 | 7 | 1,09 | 640 | Complete | HEMAHALG(60)HW(9)M | 3,47E-120 | 369,999 | | | | | | | | | | | | | |
| TrSC58v2_9025_HEIAH TrGP63 | 9 | gene | | 57 | 78.640 | 80.610 | plus | 656 | SP | 1,000 | SP | 31-32 / 0.9704 | GPI-Anchored | 637 | A | 0,247 | 1 | 0,17 | 606 | Complete | HEIAHALG(62)HW(9)M | 5,74E-118 | 362,246 | 0,9986 | 0,999 | 3 | 6 | 3 | 3 | | 7,16 | 0,9954 | 0,999 | 2 | 0 | |
| TrSC58v2_9032_HEIGH TrGP63 | 4 | gene | | 57 | 95.036 | 96.802 | plus | 588 | Other | 1,000 | SP | 42 | GPI-Anchored | 565 | S | 0,559 | 3 | 0,57 | 523 | Complete | HEIGHALG(61)HW(9)M | 0 | 688,174 | | | | | | | | | | | | | |
| TrSC58v2_9126_HEMAH TrGP63 | 11 | gene | | 60 | 34.529 | 36.715 | plus | 728 | Other | 0,579 | SP | 27 | GPI-Anchored | 699 | S | 0,206 | 5 | 0,74 | 672 | Complete | HEMAHALG(62)HW(9)M | 2,54E-118 | 365,713 | 0,9957 | 0,999 | 1 | 10 | 8 | 2 | | | | | | | |
| TrSC58v2_9131_HEIAH TrGP63 | 9 | gene | | 60 | 51.626 | 54.298 | plus | 890 | SP | 1,000 | SP | 31-32 / 0.9780 | GPI-Anchored | 863 | S | 0,254 | 4 | 0,48 | 832 | Complete | HEIAHALG(62)HW(9)M | 1,23E-102 | 329,119 | | | | | | | | | | | | | |
| TrSC58v2_9135_HEIGH TrGP63 | 4 | gene | Tandem | 60 | 68.813 | 71.134 | plus | 773 | SP | 0,608 | SP | 42-43 / 0.6714 | Not GPI-Anchored | - | * | 0,987 | 0 | 0 | 731 | Complete | HEIGHALG(61)HW(9)M | 0 | 614,216 | | | | | | | | | | | | | |
| TrSC58v2_9136_HEIGH TrGP63 | 4 | gene | Tandem | 60 | 73.244 | 75.793 | plus | 849 | SP | 0,608 | SP | 42-43 / 0.6714 | GPI-Anchored | 823 | G | 0,243 | 0 | 0 | 781 | Complete | HEIGHALG(61)HW(9)M | 0 | 680,855 | | | | | | | | 3,65 | 0,9443 | 0,9919 | 1 | 0 | g9139_HEIGH |
| TrSC58v2_9137_HEIGH TrGP63 | 4 | gene | Tandem | 60 | 77.750 | 79.210 | plus | 486 | SP | 0,608 | SP | 42-43 / 0.6714 | Not GPI-Anchored | - | * | 0,985 | 0 | 0 | 444 | Complete | HEIGHALG(61)HW(9)M | 0 | 569,148 | | | | | | | | | | | | | |
| TrSC58v2_9139_HEIGH TrGP63 | 4 | gene | Tandem | 60 | 82.330 | 84.957 | plus | 875 | SP | 0,608 | SP | 42-43 / 0.6714 | GPI-Anchored | 849 | G | 0,243 | 0 | 0 | 807 | Complete | HEIGHALG(61)HW(9)M | 0 | 677,003 | | | | | | | | | | | | | |
| TrSC58v2_9275_HEMAH TrGP63 | 11 | gene | | 65 | 43.048 | 45.177 | minus | 709 | Other | 0,834 | SP | 11 | GPI-Anchored | 681 | D | 0,213 | 5 | 0,75 | 670 | Complete | HEMAHALG(60)HW(9)M | 3,02E-123 | 377,654 | | | | | | | | 13,96 | 1 | 0,999 | 3 | 2 | |
| TrSC58v2_9319_HEIGH TrGP63 | 4 | gene | | 68 | 8.603 | 10.453 | minus | 616 | Other | 1,000 | SP | 10 | GPI-Anchored | 593 | S | 0,691 | 5 | 0,86 | 583 | Complete | HEIGHILG(61)HW(9)M | 0 | 677,003 | | | | | | | | | | | | | |
| TrSC58v2_9356_HAIAH TrGP63 | 10 | gene | | 69 | 30.152 | 31.837 | plus | 561 | Other | 1,000 | SP | 10 | GPI-Anchored | 535 | D | 0,279 | 2 | 0,38 | 525 | Complete | HAIAHALG(57)HLW(9)M | 2,46E-118 | 360,32 | | | | | | | | | | | | | |
| TrSC58v2_9363_HAIAH TrGP63 | 10 | gene | | 69 | 42.223 | 43.749 | plus | 528 | SP | 1,000 | SP | 29-30 / 0.9556 | Not GPI-Anchored | - | * | 0,985 | 2 | 0,4 | 499 | Complete | HAIAHALG(57)HLW(9)M | 6,59E-107 | 329,504 | 1,000 | 0,999 | 7 | 675 | 210 | 465 | | | | | | | |
| TrSC58v2_9386_HEMAH TrGP63 | 11 | gene | | 71 | 7.478 | 9.508 | minus | 676 | Other | 1,000 | SP | 11 | GPI-Anchored | 650 | S | 0,569 | 7 | 1,1 | 639 | Complete | HEMAHALG(60)HW(9)M | 2,29E-120 | 370,385 | | | | | | | | | | | | | |
| TrSC58v2_9430_HEIAH TrGP63 | 12 | gene | | 75 | 6.273 | 8.057 | plus | 594 | Other | 1,000 | SP | 10 | GPI-Anchored | 570 | C | 0,51 | 4 | 0,71 | 560 | Complete | HEIAHALG(60)HW(9)M | 3,61E-128 | 387,719 | | | | | | | | | | | | | |
| TrSC58v2_9446_HETGH TrGP63 | 4 | gene | | 78 | 2.144 | 4.627 | minus | 827 | SP | 0,832 | SP | 42-43 / 0.7840 | GPI-Anchored | 804 | S | 0,411 | 13 | 1,71 | 762 | Complete | HETGHILG(61)HW(9)M | 0 | 697,804 | | | | | | | | | | | | | |
| TrSC58v2_1335_HEIAH pseudo TrGP63 | 12 | pseudogene -Frameshift | Tandem | 3 | 628.261 | 630.062 | plus | 601 | SP | 1,000 | SP | 30-31 / 0.9685 | GPI-Anchored | 576 | S | 0,432 | 1 | 0,18 | 546 | Complete | HEIAHALG(60)HW(9)M | 4,44E-129 | 390,415 | | | | | | | | 13,61 | 0,8739 | 0,9804 | 1 | 1 | |
| TrSC58v2_1337_HEIAH pseudo TrGP63 | 12 | pseudogene -Frameshift | Tandem | 3 | 630.828 | 632.431 | plus | 534 | SP | 1,000 | SP | 30-31 / 0.9685 | Not GPI-Anchored | - | * | 0,992 | 5 | 0,99 | 504 | Complete | HEIAHALG(59)HW(9)M | 8,07E-128 | 385,022 | | | | | | | | | | | | | |
| TrSC58v2_3699_HEIGH pseudo TrGP63 | 4 | pseudogene -Frameshift | | 9 | 737.270 | 739.332 | plus | 688 | SP | 0,999 | SP | 22-23 / 0.9670 | GPI-Anchored | 665 | S | 0,44 | 6 | 0,93 | 643 | Complete | HEIGHILG(61)HW(9)M | 0 | 671,611 | | | | | | | | | | | | | |
| TrSC58v2_4643_HEIGH pseudo TrGP63 | 4 | pseudogene -Frameshift | Tandem | 13 | 404.439 | 406.606 | plus | 726 | SP | 0,778 | SP | 47-48 / 0.8056 | Not GPI-Anchored | - | * | 0,467 | 2 | 0,29 | 679 | Complete | HEIGHILG(61)HW(9)M | 0 | 685,093 | | | | | | | | | | | | | |
| TrSC58v2_4646_HEIGH pseudo TrGP63 | 4 | pseudogene -Frameshift | Tandem | 13 | 407.179 | 409.421 | plus | 744 | SP | 0,778 | SP | 47-48 / 0.8056 | Not GPI-Anchored | - | * | 0,49 | 2 | 0,29 | 697 | Complete | HEIGHILG(61)HW(9)M | 0 | 626,542 | | | | | | | | | | | | | |
| TrSC58v2_4647_HEIGH pseudo TrGP63 | 4 | pseudogene -Frameshift | Tandem | 13 | 410.142 | 412.195 | plus | 686 | SP | 1,000 | SP | 29-30 / 0.9769 | GPI-Anchored | 663 | S | 0,565 | 2 | 0,32 | 634 | Complete | HEIGHILG(61)HW(9)M | 0 | 684,707 | | | | | | | | | | | | | |
| TrSC58v2_4650_HEIGH pseudo TrGP63 | 4 | pseudogene -Frameshift | Tandem | 13 | 415.550 | 417.618 | plus | 729 | SP | 0,778 | SP | 47-48 / 0.8056 | GPI-Anchored | 706 | S | 0,565 | 2 | 0,3 | 659 | Complete | HEIGHILG(61)HW(9)M | 0 | 684,322 | | | | | | | | | | | | | |
| TrSC58v2_4658_HEIAH pseudo TrGP63 | 9 | pseudogene -Frameshift | Tandem | 13 | 434.204 | 435.599 | plus | 465 | Other | 1,000 | GLOB | - | Not GPI-Anchored | - | * | 0,992 | 3 | BNP | - | Complete | HEIAHALG(62)HW(9)M | 4,98E-101 | 312,17 | | | | | | | | | | | | | |
| TrSC58v2_4845_HEMAH pseudo TrGP63 | 3 | pseudogene -Frameshift | | 14 | 336.938 | 339.042 | minus | 703 | SP | 0,999 | SP | 22-23 / 0.8854 | GPI-Anchored | 680 | S | 0,531 | 2 | 0,3 | 658 | Complete | HEMAHALG(214)HW(9)M | 1,65E-119 | 368,024 | | | | | | | | | | | | | |
| TrSC58v2_5931_HEIGH pseudo TrGP63 | 4 | pseudogene -Frameshift | | 21 | 18.105 | 19.879 | minus | 589 | SP | 0,819 | SP | 42-43 / 0.7796 | GPI-Anchored | 565 | S | 0,719 | 3 | 0,57 | 523 | Complete | HEIGHILG(58)HW(9)M | 0 | 649,654 | | | | | | | | | | | | | |
| TrSC58v2_5932_HEIGH pseudo TrGP63 | 4 | pseudogene -Frameshift | | 21 | 21.616 | 23.357 | minus | 585 | SP | 0,725 | SP | 46-47 / 0.7643 | Not GPI-Anchored | - | * | 0,705 | 3 | 0,56 | 539 | Complete | HEIGHILG(61)HW(9)M | 0 | 642,721 | | | | | | | | | | | | | |
| TrSC58v2_5935_HEIGH pseudo TrGP63 | 4 | pseudogene -Frameshift | | 21 | 25.065 | 26.838 | minus | 590 | SP | 0,819 | SP | 42-43 / 0.7796 | GPI-Anchored | 566 | S | 0,719 | 3 | 0,57 | 524 | Complete | HEIGHILG(60)HW(9)M | 0 | 640,024 | | | | | | | | | | | | | |
| TrSC58v2_6135_HEIAH pseudo TrGP63 | 12 | pseudogene -Frameshift | Tandem | 22 | 135.803 | 137.604 | minus | 601 | SP | 1,000 | SP | 30-31 / 0.9665 | GPI-Anchored | 576 | S | 0,521 | 3 | 0,55 | 546 | Complete | HEIAHALG(60)HW(9)M | 5,07E-124 | 377,318 | | | | | | | | | | | | | |
| TrSC58v2_6864_HEMAH pseudo TrGP63 | 11 | pseudogene -Frameshift | | 27 | 294.007 | 296.111 | minus | 702 | Other | 0,500 | SP | 33 | GPI-Anchored | 676 | S | 0,571 | 7 | 1,09 | 643 | Complete | HEMAHALG(61)HW(9)M | 5,60E-118 | 364,992 | | | | | | | | | | | | | |
| TrSC58v2_6869_HEMAH pseudo TrGP63 | 11 | pseudogene -Frameshift | | 27 | 303.412 | 304.916 | minus | 501 | Other | 1,000 | GLOB | - | Not GPI-Anchored | - | * | 0,991 | 4 | BNP | - | Complete | HEMAHALG(61)HW(9)M | 4,37E-102 | 316,022 | | | | | | | | 12,1 | 0,9907 | 0,9987 | 1 | 1 | |
| TrSC58v2_8189_HAIAH pseudo TrGP63 | 10 | pseudogene -Frameshift | | 42 | 92.276 | 93.984 | minus | 572 | SP | 1,000 | SP | 27-28 / 0.8339 | GPI-Anchored | 546 | D | 0,307 | 4 | 0,77 | 519 | Complete | HAIAHALG(57)YLW(9)M | 5,35E-113 | 346,838 | | | | | | | | | | | | | |
| TrSC58v2_8511_HEIAH pseudo TrGP63 | 6 | pseudogene -Frameshift | | 47 | 64.324 | 65.548 | minus | 410 | SP | 1,000 | SP | 21-22 / 0.4911 | Not GPI-Anchored | - | * | 0,976 | 3 | 0,77 | 389 | C-terminal inco | HEIAHVLG(62)HW(9)M | 1,22E-131 | 388,439 | | | | | | | | | | | | | |
| TrSC58v2_8764_HELAH pseudo TrGP63 | 2 | pseudogene -Frameshift | | 51 | 131.382 | 133.111 | plus | 577 | SP | 0,894 | SP | 32-33 / 0.8281 | Not GPI-Anchored | - | * | 0,995 | 0 | 0 | 545 | Complete | HELAHVLG(63)HW(9)M | 1,07E-110 | 341,06 | | | | | | | | | | | | | |
| TrSC58v2_9176_HEIGH pseudo TrGP63 | 4 | pseudogene -Frameshift | | 62 | 1.433 | 2.433 | minus | 331 | Other | 1,000 | GLOB | - | Not GPI-Anchored | - | * | 0,992 | 0 | BNP | - | N-terminal inco | HEIGHILG(60)HW(9)M | 4,85E-161 | 461,677 | | | | | | | | | | | | | |
| TrSC58v2_9361_HDLAH pseudo TrGP63 | 2 | pseudogene -Frameshift | | 69 | 39.963 | 41.879 | plus | 720 | Other | 1,000 | GLOB | - | Not GPI-Anchored | - | * | 0,987 | 3 | BNP | - | Complete | HDLAHVLG(65)HW(8) | 7,64E-89 | 289,493 | 0,9863 | 0,999 | 1 | 2 | 0 | 2 | g3273 | | | | | | |
| TrSC58v2_2800_XXXXX pseudo TrGP63 | 2 | pseudogene -Truncated | Tandem | 7 | 222.730 | 224.652 | minus | 639 | Other | 1,000 | GLOB | - | Not GPI-Anchored | - | * | 0,991 | 2 | BNP | - | Complete | xxxxxxxxLG(63)HW(8)M | 3,47E-72 | 243,269 | | | | | | | | 9,87 | 0,9276 | 0,9893 | 1 | 0 | g3272.t1, g9362.t1 |
| TrSC58v2_3273_XXXXX pseudo TrGP63 | 2 | pseudogene -Truncated | | 8 | 506.292 | 508.043 | minus | 582 | Other | 1,000 | GLOB | - | Not GPI-Anchored | - | * | 0,774 | 2 | BNP | - | Complete | xxxxxxxxLG(64)HW(8)M | 4,12E-79 | 260,217 | | | | | | | | | | | | | |
| TrSC58v2_8382_HAIAH pseudo TrGP63 | 10 | pseudogene -Truncated | | 45 | 60.300 | 61.922 | minus | 540 | Other | 1,000 | SP | - | GPI-Anchored | 516 | C | 0,396 | 3 | BNP | - | Complete | HAIAHALG(57)HLW(9)M | 8,68E-120 | 363,016 | | | | | | | | 23,15 | 0,9998 | 0,999 | 3 | 0 | |
| TrSC58v2_8500_HFIAH pseudo TrGP63 | 10 | pseudogene -Truncated | | 47 | 35.912 | 37.225 | plus | 437 | SP | 0,666 | SP | 29-30 / 0.3528 | Not GPI-Anchored | - | * | 0,975 | 3 | 0,74 | 408 | Complete | HFIAHALG(57)HLW(9)M | 1,05E-98 | 305,236 | | | | | | | | | | | | | |
| TrSC58v2_9026_HEIGH pseudo TrGP63 | 4 | pseudogene -Truncated | | 57 | 81.423 | 83.021 | plus | 528 | Other | 1,000 | GLOB | - | GPI-Anchored | 505 | S | 0,559 | 0 | BNP | - | Complete | HEIGHALG(167)H(9)M | 6,20E-176 | 507,901 | | | | | | | | | | | | | |
| TrSC58v2_9140_XXXXX pseudo TrGP63 | NE | pseudogene -Truncated | Tandem | 60 | 86.913 | 88.339 | plus | 494 | SP | 0,608 | SP | 42-43 / 0.6714 | Not GPI-Anchored | - | * | 0,994 | 1 | 0,22 | 452 | Complete | Absent(General substitutions) | 4,67E-32 | 129,635 | | | | | | | | | | | | | |
| TrSC58v2_2797_HAIAH TrGP63 | 10 | pseudogene-incomplete domain | Tandem | 7 | 215.036 | 216.253 | minus | 405 | Other | 1,000 | GLOB | - | Not GPI-Anchored | - | * | 0,985 | 3 | BNP | - | N-terminal inco | HAIAHALG(57)HLW(9)M | 1,14E-87 | 275,576 | 0 | 0,999 | 7 | 42 | 15 | 27 | | | | | | | |
| TrSC58v2_2798_----- pseudo TrGP63 | NE | pseudogene-incomplete domain | Tandem | 7 | 217.121 | 218.257 | minus | 379 | Other | 1,000 | GLOB | - | Not GPI-Anchored | - | * | 0,991 | 0 | BNP | NBP | N-terminal inco | Absent(GAP) | 4,90E-07 | 51,3897 |  |  |  |  |  |  |  |  |  |  |  |  |  |
| TrSC58v2_2799_HAIAH TrGP63 | 10 | pseudogene-incomplete domain | Tandem | 7 | 220.506 | 221.687 | minus | 393 | Other | 1,000 | GLOB | - | GPI-Anchored | 367 | D | 0,28 | 4 | BNP | - | N-terminal inco | HAIAHALG(57)HLW(9)M | 1,57E-78 | 251,308 | | | | | | | | | | | | | |
| TrSC58v2_4080_HEIAH pseudo TrGP63 | 11 | pseudogene-incomplete domain | | 11 | 225.597 | 226.226 | plus | 248 | Other | 1,000 | GLOB | - | Not GPI-Anchored | - | * | 0,988 | 0 | BNP | - | N-terminal inco | HEIAHAPG(61)HW(9)V | 5,79E-50 | 171,187 | | | | | | | | | | | | | |
| TrSC58v2_5264_HEIAH pseudo TrGP63 | 9 | pseudogene-incomplete domain | Tandem | 16 | 373.348 | 374.184 | plus | 279 | Other | 1,000 | GLOB | - | Not GPI-Anchored | - | * | 0,988 | 1 | BNP | NBP | C-terminal inco | HEIAHALG(62)HW(9)M | 1,30E-67 | 218,566 | | | | | | | | | | | | | |
| TrSC58v2_5265_----- pseudo TrGP63 | NE | pseudogene-incomplete domain | Tandem | 16 | 374.235 | 375.281 | plus | 348 | Other | 1,000 | GLOB | - | GPI-Anchored | 320 | D | 0,239 | 3 | BNP | - | N-terminal inco | Absent(GAP) | 3,85E-19 | 88,3688 |  |  |  |  |  |  |  |  |  |  |  |  |  |
| TrSC58v2_5267_HEIAH pseudo TrGP63 | 9 | pseudogene-incomplete domain | Tandem | 16 | 378.430 | 379.398 | plus | 323 | SP | 1,000 | SP | 31-32 / 0.9735 | Not GPI-Anchored | - | * | 0,988 | 1 | 0,34 | 292 | C-terminal inco | HEIAHALG(62)HW(9)M | 1,99E-66 | 217,411 | | | | | | | | | | | | | |
| TrSC58v2_5268_----- pseudo TrGP63 | NE | pseudogene-incomplete domain | Tandem | 16 | 379.853 | 380.485 | plus | 210 | Other | 1,000 | GLOB | - | GPI-Anchored | 182 | D | 0,238 | 2 | BNP | - | N-terminal inco | Absent(GAP) | 0,000332845 | 41,0389 |  |  |  |  |  |  |  |  |  |  |  |  |  |
| TrSC58v2_556_RGLAY TrGP63 | 11 | pseudogene-incomplete domain | | 1 | 1.325.516 | 1.326.805 | plus | 429 | Other | 1,000 | SP | 10 | Not GPI-Anchored | - | * | 0,995 | 5 | 1,19 | 419 | C-terminal inco | RGLAYALG(61)HW(9)M | 2,84E-109 | 332,2 | | | | | | | | | | | | | |
| TrSC58v2_8168_HAIAH TrGP63 | 10 | pseudogene-incomplete domain | | 42 | 47.854 | 48.867 | minus | 337 | Other | 1,000 | GLOB | - | GPI-Anchored | 313 | C | 0,34 | 3 | BNP | - | N-terminal inco | HAIAHALG(57)YLW(9)M | 1,79E-64 | 212,788 | | | | | | | | | | | | | |
| TrSC58v2_8469_XXXXX pseudo TrGP63 | 4 | pseudogene-incomplete domain | | 46 | 121.592 | 122.551 | minus | 319 | Other | 1,000 | GLOB | - | GPI-Anchored | 297 | C | 0,243 | 3 | BNP | - | N-terminal inco | Absent(GAP) | 7,01E-101 | 307,212 | | | | | | | | | | | | | |
| TrSC58v2_8923_HEMAH TrGP63 | 11 | pseudogene-incomplete domain | | 54 | 108.966 | 110.147 | minus | 393 | Other | 1,000 | SP | 10 | Not GPI-Anchored | - | * | 0,992 | 4 | 1,04 | 383 | C-terminal inco | HEMAHALG(61)HW(9)M | 6,35E-105 | 320,694 | | | | | | | | | | | | | |
| TrSC58v2_9171_HEIAH pseudo TrGP63 | 9 | pseudogene-incomplete domain | | 61 | 77.708 | 78.676 | plus | 323 | SP | 0,644 | SP | 30-31 / 0.6098 | Not GPI-Anchored | - | * | 0,988 | 1 | 0,34 | 293 | C-terminal inco | HEIAHALG(62)HW(9)M | 1,14E-70 | 229,402 | | | | | | | | | | | | | |
| TrSC58v2_9172_----- pseudo TrGP63 | NE | pseudogene-incomplete domain |  | 61 | 78.727 | 79.947 | plus | 406 | Other | 1,000 | GLOB | - | GPI-Anchored | 380 | S | 0,25 | 3 | BNP | - | N-terminal inco | Absent(GAP) | 1,69E-17 | 84,1316 |  |  |  |  |  |  |  |  |  |  |  |  |  |

GPI-Anchored in red: sequence forecasted as GPI-anchored, however biologicaly not possible if signal peptide prediction is correct.

NE: Non-evaluated, SP: Signal Peptide, GLOB: Glubular
