## Supplementary table S4 for "Comprehensive genomic analysis of *Trypanosoma rangeli* reveals key insights into the biology and evolution of this non-virulent American mammalian trypanosome"

| | TrGP63 Groups | 12 | 12 | 12 | 12 | 12 | 12 | 12 | 12 | 12 | 12 | 12 | 12 | 12 | 11 | 11 | 11 | 11 | 11 | 11 | 11 | 11 | 11 | 11 | 11 |
|---|---|---|---|---|---|---|---|---|---|---|---|---|---|---|---|---|---|---|---|---|---|---|---|---|---|
| 1 | TrSC58v2_1339_HEIAH_TrGP63 | 100,00 | 64,58 | 65,42 | 73,02 | 73,02 | 73,02 | 73,02 | 73,02 | 73,02 | 73,21 | 68,98 | 68,98 | 68,98 | 49,91 | 50,27 | 50,27 | 50,71 | 50,28 | 52,26 | 50,18 | 50,46 | 50,70 | 49,74 | 51,08 |
| 2 | TrSC58v2_2794_HEIAH_TrGP63 | 64,58 | 100,00 | 98,32 | 88,18 | 88,18 | 88,18 | 88,18 | 88,18 | 88,18 | 87,99 | 84,63 | 84,63 | 84,63 | 44,12 | 42,90 | 42,90 | 43,00 | 43,62 | 48,43 | 44,87 | 47,50 | 44,21 | 44,50 | 44,32 |
| 3 | TrSC58v2_9430_HEIAH_TrGP63 | 65,42 | 98,32 | 100,00 | 88,28 | 88,28 | 88,28 | 88,28 | 88,28 | 88,28 | 88,09 | 86,52 | 86,52 | 86,52 | 45,10 | 44,46 | 44,46 | 44,56 | 45,56 | 51,19 | 46,66 | 48,49 | 45,86 | 47,16 | 47,02 |
| 4 | TrSC58v2_1328_HEIAH_TrGP63 | 73,02 | 88,18 | 88,28 | 100,00 | 100,00 | 100,00 | 100,00 | 100,00 | 100,00 | 99,81 | 89,12 | 89,12 | 89,12 | 47,42 | 47,60 | 47,60 | 47,89 | 46,83 | 53,31 | 48,94 | 50,00 | 47,98 | 48,85 | 48,92 |
| 5 | TrSC58v2_1329_HEIAH_TrGP63 | 73,02 | 88,18 | 88,28 | 100,00 | 100,00 | 100,00 | 100,00 | 100,00 | 100,00 | 99,81 | 89,12 | 89,12 | 89,12 | 47,42 | 47,60 | 47,60 | 47,89 | 46,83 | 53,31 | 48,94 | 50,00 | 47,98 | 48,85 | 48,92 |
| 6 | TrSC58v2_1330_HEIAH_TrGP63 | 73,02 | 88,18 | 88,28 | 100,00 | 100,00 | 100,00 | 100,00 | 100,00 | 100,00 | 99,81 | 89,12 | 89,12 | 89,12 | 47,42 | 47,60 | 47,60 | 47,89 | 46,83 | 53,31 | 48,94 | 50,00 | 47,98 | 48,85 | 48,92 |
| 7 | TrSC58v2_1331_HEIAH_TrGP63 | 73,02 | 88,18 | 88,28 | 100,00 | 100,00 | 100,00 | 100,00 | 100,00 | 100,00 | 99,81 | 89,12 | 89,12 | 89,12 | 47,42 | 47,60 | 47,60 | 47,89 | 46,83 | 53,31 | 48,94 | 50,00 | 47,98 | 48,85 | 48,92 |
| 8 | TrSC58v2_1332_HEIAH_TrGP63 | 73,02 | 88,18 | 88,28 | 100,00 | 100,00 | 100,00 | 100,00 | 100,00 | 100,00 | 99,81 | 89,12 | 89,12 | 89,12 | 47,42 | 47,60 | 47,60 | 47,89 | 46,83 | 53,31 | 48,94 | 50,00 | 47,98 | 48,85 | 48,92 |
| 9 | TrSC58v2_1334_HEIAH_TrGP63 | 73,02 | 88,18 | 88,28 | 100,00 | 100,00 | 100,00 | 100,00 | 100,00 | 100,00 | 99,81 | 89,12 | 89,12 | 89,12 | 47,42 | 47,60 | 47,60 | 47,89 | 46,83 | 53,31 | 48,94 | 50,00 | 47,98 | 48,85 | 48,92 |
| 10 | TrSC58v2_1333_HEIAH_TrGP63 | 73,21 | 87,99 | 88,09 | 99,81 | 99,81 | 99,81 | 99,81 | 99,81 | 99,81 | 100,00 | 89,12 | 89,12 | 89,12 | 47,61 | 47,60 | 47,60 | 48,08 | 47,02 | 53,31 | 48,94 | 50,00 | 47,98 | 48,85 | 48,72 |
| 11 | TrSC58v2_6131_HEIAH_TrGP63 | 68,98 | 84,63 | 86,52 | 89,12 | 89,12 | 89,12 | 89,12 | 89,12 | 89,12 | 89,12 | 100,00 | 100,00 | 100,00 | 45,84 | 46,18 | 46,18 | 46,81 | 46,15 | 52,43 | 48,58 | 48,63 | 47,54 | 47,62 | 47,66 |
| 12 | TrSC58v2_6132_HEIAH_TrGP63 | 68,98 | 84,63 | 86,52 | 89,12 | 89,12 | 89,12 | 89,12 | 89,12 | 89,12 | 89,12 | 100,00 | 100,00 | 100,00 | 45,84 | 46,18 | 46,18 | 46,81 | 46,15 | 52,43 | 48,58 | 48,63 | 47,54 | 47,62 | 47,66 |
| 13 | TrSC58v2_6133_HEIAH_TrGP63 | 68,98 | 84,63 | 86,52 | 89,12 | 89,12 | 89,12 | 89,12 | 89,12 | 89,12 | 89,12 | 100,00 | 100,00 | 100,00 | 45,84 | 46,18 | 46,18 | 46,81 | 46,15 | 52,43 | 48,58 | 48,63 | 47,54 | 47,62 | 47,66 |
| 14 | TrSC58v2_8803_HEMAH_TrGP63 | 49,91 | 44,12 | 45,10 | 47,42 | 47,42 | 47,42 | 47,42 | 47,42 | 47,42 | 47,61 | 45,84 | 45,84 | 45,84 | 100,00 | 93,86 | 93,85 | 94,03 | 94,64 | 63,86 | 63,92 | 62,74 | 66,39 | 68,13 | 66,50 |
| 15 | TrSC58v2_8920_HEMAH_TrGP63 | 50,27 | 42,90 | 44,46 | 47,60 | 47,60 | 47,60 | 47,60 | 47,60 | 47,60 | 47,60 | 46,18 | 46,18 | 46,18 | 93,86 | 100,00 | 99,85 | 95,72 | 96,01 | 60,12 | 59,52 | 59,39 | 62,11 | 63,64 | 61,89 |
| 16 | TrSC58v2_9386_HEMAH_TrGP63 | 50,27 | 42,90 | 44,46 | 47,60 | 47,60 | 47,60 | 47,60 | 47,60 | 47,60 | 47,60 | 46,18 | 46,18 | 46,18 | 93,85 | 99,85 | 100,00 | 95,56 | 95,85 | 60,06 | 59,46 | 59,48 | 62,05 | 63,58 | 61,83 |
| 17 | TrSC58v2_6853_HEMAH_TrGP63 | 50,71 | 43,00 | 44,56 | 47,89 | 47,89 | 47,89 | 47,89 | 47,89 | 47,89 | 48,08 | 46,81 | 46,81 | 46,81 | 94,03 | 95,72 | 95,56 | 100,00 | 98,18 | 60,48 | 60,63 | 60,10 | 63,06 | 63,39 | 62,40 |
| 18 | TrSC58v2_8770_HEMAH_TrGP63 | 50,28 | 43,62 | 45,56 | 46,83 | 46,83 | 46,83 | 46,83 | 46,83 | 46,83 | 47,02 | 46,15 | 46,15 | 46,15 | 94,64 | 96,01 | 95,85 | 98,18 | 100,00 | 63,27 | 62,99 | 62,35 | 66,50 | 66,39 | 65,69 |
| 19 | TrSC58v2_6130_HEMTH_TrGP63 | 52,26 | 48,43 | 51,19 | 53,31 | 53,31 | 53,31 | 53,31 | 53,31 | 53,31 | 53,31 | 52,43 | 52,43 | 52,43 | 63,86 | 60,12 | 60,06 | 60,48 | 63,27 | 100,00 | 78,11 | 68,54 | 73,79 | 71,43 | 73,65 |
| 20 | TrSC58v2_560_RGLAY_TrGP63 | 50,18 | 44,87 | 46,66 | 48,94 | 48,94 | 48,94 | 48,94 | 48,94 | 48,94 | 48,94 | 48,58 | 48,58 | 48,58 | 63,92 | 59,52 | 59,46 | 60,63 | 62,99 | 78,11 | 100,00 | 85,07 | 73,77 | 70,03 | 73,81 |
| 21 | TrSC58v2_8766_RGLAY_TrGP63 | 50,46 | 47,50 | 48,49 | 50,00 | 50,00 | 50,00 | 50,00 | 50,00 | 50,00 | 50,00 | 48,63 | 48,63 | 48,63 | 62,74 | 59,39 | 59,48 | 60,10 | 62,35 | 68,54 | 85,07 | 100,00 | 62,84 | 60,97 | 63,52 |
| 22 | TrSC58v2_9126_HEMAH_TrGP63 | 50,70 | 44,21 | 45,86 | 47,98 | 47,98 | 47,98 | 47,98 | 47,98 | 47,98 | 47,98 | 47,54 | 47,54 | 47,54 | 66,39 | 62,11 | 62,05 | 63,06 | 66,50 | 73,79 | 73,77 | 62,84 | 100,00 | 77,05 | 75,95 |
| 23 | TrSC58v2_6495_HEMAH_TrGP63 | 49,74 | 44,50 | 47,16 | 48,85 | 48,85 | 48,85 | 48,85 | 48,85 | 48,85 | 48,85 | 47,62 | 47,62 | 47,62 | 68,13 | 63,64 | 63,58 | 63,39 | 66,39 | 71,43 | 70,03 | 60,97 | 77,05 | 100,00 | 82,91 |
| 24 | TrSC58v2_8644_HEMAH_TrGP63 | 51,08 | 44,32 | 47,02 | 48,92 | 48,92 | 48,92 | 48,92 | 48,92 | 48,92 | 48,72 | 47,66 | 47,66 | 47,66 | 66,50 | 61,89 | 61,83 | 62,40 | 65,69 | 73,65 | 73,81 | 63,52 | 75,95 | 82,91 | 100,00 |
| 25 | TrSC58v2_9275_HEMAH_TrGP63 | 50,18 | 44,81 | 47,37 | 48,72 | 48,72 | 48,72 | 48,72 | 48,72 | 48,72 | 48,72 | 47,66 | 47,66 | 47,66 | 66,00 | 61,43 | 61,37 | 61,95 | 65,18 | 74,64 | 74,53 | 64,80 | 75,95 | 80,93 | 93,23 |
| 26 | TrSC58v2_8384_HELAH_TrGP63 | 40,29 | 39,71 | 40,51 | 39,49 | 39,49 | 39,49 | 39,49 | 39,49 | 39,49 | 39,49 | 40,55 | 40,55 | 40,55 | 41,21 | 41,99 | 41,99 | 41,50 | 41,09 | 41,35 | 42,36 | 43,30 | 41,37 | 41,28 | 42,45 |
| 27 | TrSC58v2_559_HELAH_TrGP63 | 40,70 | 39,20 | 39,72 | 39,76 | 39,76 | 39,76 | 39,76 | 39,76 | 39,76 | 39,76 | 39,96 | 39,96 | 39,96 | 41,03 | 41,76 | 41,76 | 41,11 | 40,59 | 41,50 | 41,15 | 41,83 | 40,19 | 40,37 | 42,52 |
| 28 | TrSC58v2_8165_HAIAH_TrGP63 | 46,29 | 44,85 | 45,04 | 44,29 | 44,29 | 44,29 | 44,29 | 44,29 | 44,29 | 44,29 | 44,76 | 44,76 | 44,76 | 44,44 | 44,81 | 44,81 | 44,72 | 43,91 | 45,25 | 46,17 | 49,13 | 46,67 | 46,38 | 45,47 |
| 29 | TrSC58v2_9363_HAIAH_TrGP63 | 43,90 | 42,28 | 42,68 | 42,89 | 42,89 | 42,89 | 42,89 | 42,89 | 42,89 | 42,89 | 42,48 | 42,48 | 42,48 | 44,99 | 44,97 | 44,97 | 45,49 | 44,07 | 44,22 | 46,01 | 46,42 | 44,38 | 44,27 | 42,62 |
| 30 | TrSC58v2_9356_HAIAH_TrGP63 | 45,05 | 45,25 | 45,62 | 44,51 | 44,51 | 44,51 | 44,51 | 44,51 | 44,51 | 44,51 | 44,49 | 44,49 | 44,49 | 44,65 | 45,19 | 45,27 | 45,29 | 45,54 | 45,16 | 46,22 | 50,29 | 45,10 | 44,55 | 42,99 |
| 31 | TrSC58v2_3271_HAIAH_TrGP63 | 43,58 | 42,81 | 45,04 | 43,90 | 43,90 | 43,90 | 43,90 | 43,90 | 43,90 | 43,90 | 43,04 | 43,04 | 43,04 | 42,02 | 40,19 | 40,26 | 40,45 | 41,59 | 38,88 | 40,52 | 45,48 | 39,07 | 38,17 | 37,23 |
| 32 | TrSC58v2_3280_HAIAH_TrGP63 | 45,42 | 45,90 | 46,27 | 44,71 | 44,71 | 44,71 | 44,71 | 44,71 | 44,71 | 44,71 | 44,86 | 44,86 | 44,86 | 44,46 | 44,81 | 44,90 | 45,10 | 45,44 | 44,61 | 46,41 | 50,48 | 44,18 | 43,81 | 42,23 |
| 33 | TrSC58v2_5263_HEIAH_TrGP63 | 38,00 | 37,53 | 38,49 | 38,86 | 38,86 | 38,86 | 38,86 | 38,86 | 38,86 | 38,86 | 38,14 | 38,14 | 38,14 | 35,68 | 35,00 | 35,00 | 35,13 | 35,85 | 35,48 | 37,25 | 37,68 | 36,63 | 37,69 | 38,51 |
| 34 | TrSC58v2_5266_HEIAH_TrGP63 | 38,22 | 37,73 | 38,70 | 39,09 | 39,09 | 39,09 | 39,09 | 39,09 | 39,09 | 39,09 | 38,35 | 38,35 | 38,35 | 35,68 | 35,00 | 35,00 | 35,13 | 35,85 | 35,48 | 37,25 | 37,68 | 36,63 | 37,69 | 38,51 |
| 35 | TrSC58v2_6663_HEIAH_TrGP63 | 37,38 | 36,67 | 37,32 | 38,54 | 38,54 | 38,54 | 38,54 | 38,54 | 38,54 | 38,54 | 38,26 | 38,26 | 38,26 | 35,78 | 33,61 | 33,45 | 33,56 | 35,28 | 35,45 | 35,21 | 35,21 | 35,44 | 36,18 | 36,95 |
| 36 | TrSC58v2_9131_HEIAH_TrGP63 | 37,52 | 35,31 | 35,98 | 37,53 | 37,53 | 37,53 | 37,53 | 37,53 | 37,53 | 37,53 | 36,88 | 36,88 | 36,88 | 35,11 | 32,83 | 32,83 | 32,61 | 34,40 | 35,18 | 35,58 | 35,90 | 35,65 | 34,97 | 36,93 |
| 37 | TrSC58v2_9025_HEIAH_TrGP63 | 39,67 | 37,97 | 38,24 | 38,65 | 38,65 | 38,65 | 38,65 | 38,65 | 38,65 | 38,65 | 38,16 | 38,16 | 38,16 | 38,12 | 38,14 | 38,14 | 37,67 | 37,68 | 40,08 | 39,00 | 38,89 | 40,04 | 39,30 | 40,28 |
| 38 | TrSC58v2_4259_HEIAH_TrGP63 | 40,88 | 37,72 | 38,10 | 39,92 | 39,92 | 39,92 | 39,92 | 39,92 | 39,92 | 39,92 | 39,85 | 39,85 | 39,85 | 36,31 | 33,96 | 33,79 | 33,73 | 35,44 | 37,60 | 35,68 | 36,12 | 35,30 | 37,02 | 37,69 |
| 39 | TrSC58v2_631_HEIGH_TrGP63 | 40,87 | 39,85 | 39,70 | 42,22 | 42,22 | 42,22 | 42,22 | 42,22 | 42,22 | 42,22 | 40,49 | 40,49 | 40,49 | 38,52 | 37,41 | 37,48 | 36,81 | 39,20 | 39,75 | 37,59 | 37,04 | 37,91 | 38,94 | 39,42 |
| 40 | TrSC58v2_632_HEIGH_TrGP63 | 40,87 | 39,85 | 39,70 | 42,22 | 42,22 | 42,22 | 42,22 | 42,22 | 42,22 | 42,22 | 40,49 | 40,49 | 40,49 | 38,52 | 37,41 | 37,48 | 36,81 | 39,20 | 39,75 | 37,59 | 37,04 | 37,91 | 38,94 | 39,42 |
| 41 | TrSC58v2_633_HEIGH_TrGP63 | 40,87 | 39,85 | 39,70 | 42,22 | 42,22 | 42,22 | 42,22 | 42,22 | 42,22 | 42,22 | 40,49 | 40,49 | 40,49 | 38,52 | 37,41 | 37,48 | 36,81 | 39,20 | 39,75 | 37,59 | 37,04 | 37,91 | 38,94 | 39,42 |
| 42 | TrSC58v2_4929_HEIGH_TrGP63 | 38,89 | 38,49 | 39,04 | 39,96 | 39,96 | 39,96 | 39,96 | 39,96 | 39,96 | 39,96 | 39,30 | 39,30 | 39,30 | 37,28 | 35,66 | 35,73 | 35,41 | 37,58 | 37,21 | 37,36 | 37,38 | 36,30 | 37,12 | 37,00 |
| 43 | TrSC58v2_6472_HEIGH_TrGP63 | 40,46 | 39,49 | 40,26 | 40,94 | 40,94 | 40,94 | 40,94 | 40,94 | 40,94 | 40,94 | 39,50 | 39,50 | 39,50 | 40,96 | 39,05 | 39,12 | 38,62 | 41,24 | 38,10 | 36,71 | 35,60 | 36,52 | 37,76 | 38,08 |
| 44 | TrSC58v2_7196_HEIGH_TrGP63 | 39,70 | 38,16 | 40,48 | 41,82 | 41,82 | 41,82 | 41,82 | 41,82 | 41,82 | 41,82 | 39,33 | 39,33 | 39,33 | 39,53 | 36,95 | 36,84 | 36,39 | 38,78 | 36,56 | 35,46 | 34,69 | 35,91 | 36,25 | 36,18 |
| 45 | TrSC58v2_9139_HEIGH_TrGP63 | 40,15 | 39,56 | 39,49 | 42,42 | 42,42 | 42,42 | 42,42 | 42,42 | 42,42 | 42,42 | 39,77 | 39,77 | 39,77 | 38,85 | 36,82 | 36,88 | 36,58 | 39,15 | 37,25 | 35,33 | 36,14 | 36,75 | 37,88 | 37,54 |
| 46 | TrSC58v2_9136_HEIGH_TrGP63 | 40,15 | 39,74 | 39,85 | 42,42 | 42,42 | 42,42 | 42,42 | 42,42 | 42,42 | 42,42 | 39,77 | 39,77 | 39,77 | 38,85 | 37,22 | 37,28 | 36,98 | 38,95 | 37,84 | 35,71 | 36,61 | 36,98 | 38,13 | 37,80 |
| 47 | TrSC58v2_9135_HEIGH_TrGP63 | 39,51 | 39,11 | 39,22 | 41,75 | 41,75 | 41,75 | 41,75 | 41,75 | 41,75 | 41,75 | 39,51 | 39,51 | 39,51 | 38,18 | 36,77 | 36,77 | 36,52 | 37,04 | 36,98 | 35,05 | 35,62 | 36,96 | 37,79 | 37,66 |
| 48 | TrSC58v2_9137_HEIGH_TrGP63 | 40,65 | 42,29 | 42,29 | 43,22 | 43,22 | 43,22 | 43,22 | 43,22 | 43,22 | 43,22 | 41,59 | 41,59 | 41,59 | 41,16 | 40,89 | 40,89 | 40,56 | 40,15 | 41,24 | 39,53 | 39,07 | 40,81 | 41,93 | 42,07 |
| 49 | TrSC58v2_9032_HEIGH_TrGP63 | 40,12 | 39,60 | 39,60 | 39,92 | 39,92 | 39,92 | 39,92 | 39,92 | 39,92 | 39,92 | 38,91 | 38,91 | 38,91 | 40,20 | 39,76 | 39,76 | 39,68 | 39,09 | 39,02 | 37,94 | 38,37 | 38,00 | 38,89 | 39,84 |
| 50 | TrSC58v2_5291_HEIGH_TrGP63 | 40,64 | 40,04 | 40,37 | 41,94 | 41,94 | 41,94 | 41,94 | 41,94 | 41,94 | 41,94 | 40,08 | 40,08 | 40,08 | 38,14 | 36,65 | 36,71 | 36,76 | 38,98 | 38,06 | 36,49 | 36,45 | 38,14 | 39,36 | 38,93 |
| 51 | TrSC58v2_7524_HEIAH_TrGP63 | 39,38 | 36,74 | 39,17 | 40,62 | 40,62 | 40,62 | 40,62 | 40,62 | 40,62 | 40,62 | 38,54 | 38,54 | 38,54 | 37,84 | 35,02 | 34,92 | 35,13 | 37,63 | 34,20 | 34,00 | 33,39 | 34,79 | 34,39 | 34,53 |
| 52 | TrSC58v2_7598_QKIGH_TrGP63 | 40,64 | 38,47 | 40,15 | 41,13 | 41,13 | 41,13 | 41,13 | 41,13 | 41,13 | 41,13 | 40,19 | 40,19 | 40,19 | 38,75 | 36,04 | 35,93 | 35,98 | 38,35 | 36,78 | 36,00 | 35,92 | 35,90 | 35,92 | 35,20 |
| 53 | TrSC58v2_9319_HEIGH_TrGP63 | 40,83 | 42,91 | 42,94 | 44,06 | 44,06 | 44,06 | 44,06 | 44,06 | 44,06 | 44,06 | 41,51 | 41,51 | 41,51 | 39,08 | 38,53 | 38,60 | 38,28 | 39,80 | 41,03 | 38,20 | 37,31 | 39,46 | 39,36 | 40,11 |
| 54 | TrSC58v2_4655_HEIGH_TrGP63 | 40,62 | 41,36 | 41,24 | 41,44 | 41,44 | 41,44 | 41,44 | 41,44 | 41,44 | 41,44 | 39,92 | 39,92 | 39,92 | 39,18 | 39,14 | 39,14 | 38,85 | 38,00 | 39,96 | 38,04 | 37,63 | 38,22 | 38,21 | 38,74 |
| 55 | TrSC58v2_4642_HEIGH_TrGP63 | 42,64 | 41,26 | 41,39 | 42,42 | 42,42 | 42,42 | 42,42 | 42,42 | 42,42 | 42,42 | 41,79 | 41,79 | 41,79 | 39,07 | 37,72 | 37,79 | 37,83 | 39,17 | 39,15 | 37,68 | 37,41 | 38,17 | 37,83 | 38,48 |
| 56 | TrSC58v2_4641_HEIGH_TrGP63 | 42,34 | 40,39 | 41,19 | 42,42 | 42,42 | 42,42 | 42,42 | 42,42 | 42,42 | 42,42 | 41,32 | 41,32 | 41,32 | 38,79 | 37,02 | 37,09 | 37,13 | 38,72 | 38,46 | 37,06 | 36,88 | 37,35 | 37,21 | 37,63 |
| 57 | TrSC58v2_4649_HEIGH_TrGP63 | 42,64 | 41,26 | 41,39 | 42,42 | 42,42 | 42,42 | 42,42 | 42,42 | 42,42 | 42,42 | 41,79 | 41,79 | 41,79 | 39,07 | 37,72 | 37,79 | 37,83 | 39,17 | 39,15 | 37,68 | 37,41 | 38,17 | 37,92 | 38,62 |
| 58 | TrSC58v2_4654_HEIGH_TrGP63 | 42,34 | 40,04 | 41,19 | 42,42 | 42,42 | 42,42 | 42,42 | 42,42 | 42,42 | 42,42 | 41,32 | 41,32 | 41,32 | 38,64 | 35,80 | 35,86 | 35,91 | 37,29 | 37,72 | 36,35 | 36,59 | 36,47 | 36,51 | 36,89 |
| 59 | TrSC58v2_4653_HEIGH_TrGP63 | 37,28 | 40,23 | 40,52 | 39,66 | 39,66 | 39,66 | 39,66 | 39,66 | 39,66 | 39,66 | 39,37 | 39,37 | 39,37 | 37,61 | 37,82 | 37,82 | 37,71 | 37,43 | 39,20 | 37,32 | 36,75 | 37,78 | 37,07 | 37,64 |
| 60 | TrSC58v2_4652_HEIGH_TrGP63 | 37,59 | 36,28 | 37,77 | 40,81 | 40,81 | 40,81 | 40,81 | 40,81 | 40,81 | 40,81 | 36,78 | 36,78 | 36,78 | 36,80 | 34,78 | 34,83 | 34,22 | 35,45 | 34,41 | 32,68 | 34,09 | 32,30 | 32,91 | 32,67 |
| 61 | TrSC58v2_4656_HEIGH_TrGP63 | 33,33 | 35,04 | 35,34 | 36,53 | 36,53 | 36,53 | 36,53 | 36,53 | 36,53 | 36,53 | 33,83 | 33,83 | 33,83 | 32,78 | 32,78 | 32,85 | 32,51 | 33,26 | 33,26 | 32,09 | 32,33 | 31,21 | 30,91 | 31,61 |
| 62 | TrSC58v2_9446_HETGH_TrGP63 | 41,39 | 38,60 | 40,07 | 41,13 | 41,13 | 41,13 | 41,13 | 41,13 | 41,13 | 41,13 | 39,25 | 39,25 | 39,25 | 39,61 | 37,09 | 37,31 | 36,69 | 38,70 | 37,28 | 35,63 | 35,68 | 36,72 | 36,66 | 37,10 |
| 63 | TrSC58v2_4795_HEIAH_TrGP63 | 41,56 | 40,70 | 40,89 | 41,82 | 41,82 | 41,82 | 41,82 | 41,82 | 41,82 | 41,82 | 40,15 | 40,15 | 40,15 | 40,44 | 38,56 | 38,62 | 38,31 | 40,87 | 38,22 | 36,99 | 36,60 | 37,95 | 38,28 | 38,74 |
| 64 | TrSC58v2_7306_HEIAH_TrGP63 | 42,07 | 40,75 | 40,38 | 41,33 | 41,33 | 41,33 | 41,33 | 41,33 | 41,33 | 41,33 | 39,89 | 39,89 | 39,89 | 40,49 | 39,16 | 39,23 | 38,91 | 40,52 | 38,85 | 37,50 | 36,59 | 39,44 | 40,14 | 39,59 |
| 65 | TrSC58v2_4835_HEMAH_TrGP63 | 41,49 | 40,46 | 41,00 | 40,66 | 40,66 | 40,66 | 40,66 | 40,66 | 40,66 | 40,66 | 41,70 | 41,70 | 41,70 | 38,35 | 38,51 | 38,51 | 38,64 | 38,76 | 39,88 | 39,05 | 39,05 | 37,55 | 37,83 | 39,71 |
| 66 | TrSC58v2_3991_HEIVH_TrGP63 | 45,90 | 42,89 | 42,28 | 43,44 | 43,44 | 43,44 | 43,44 | 43,44 | 43,44 | 43,44 | 44,47 | 44,47 | 44,47 | 41,78 | 41,96 | 41,96 | 42,07 | 41,05 | 40,76 | 40,69 | 40,89 | 40,29 | 39,59 | 40,29 |
| 67 | TrSC58v2_4583_HEIAH_TrGP63 | 39,44 | 40,23 | 40,85 | 41,82 | 41,82 | 41,82 | 41,82 | 41,82 | 41,82 | 41,82 | 39,96 | 39,96 | 39,96 | 40,82 | 40,59 | 40,59 | 40,70 | 41,01 | 40,00 | 39,81 | 39,96 | 40,19 | 39,07 | 41,29 |
| 68 | TrSC58v2_7126_HEIAH_TrGP63 | 41,13 | 39,92 | 39,84 | 40,85 | 40,85 | 40,85 | 40,85 | 40,85 | 40,85 | 40,85 | 41,25 | 41,25 | 41,25 | 42,23 | 42,00 | 42,00 | 42,32 | 42,47 | 42,41 | 40,92 | 41,40 | 41,25 | 40,35 | 40,98 |
| 69 | TrSC58v2_8509_HEIAH_TrGP63 | 42,95 | 44,33 | 44,33 | 43,91 | 43,91 | 43,91 | 43,91 | 43,91 | 43,91 | 43,91 | 43,70 | 43,70 | 43,70 | 40,46 | 40,42 | 40,42 | 40,13 | 40,22 | 41,58 | 41,00 | 41,21 | 40,62 | 41,24 | 42,33 |
| 70 | TrSC58v2_2748_HEIAH_TrGP63 | 44,00 | 44,54 | 44,33 | 43,91 | 43,91 | 43,91 | 43,91 | 43,91 | 43,91 | 43,91 | 43,49 | 43,49 | 43,49 | 41,09 | 41,05 | 41,05 | 40,55 | 40,65 | 41,16 | 40,17 | 40,17 | 41,36 | 40,74 | 41,47 |
| 71 | TrSC58v2_6871_HEIAH_TrGP63 | 44,00 | 44,12 | 43,91 | 43,70 | 43,70 | 43,70 | 43,70 | 43,70 | 43,70 | 43,70 | 43,07 | 43,07 | 43,07 | 40,88 | 40,63 | 40,63 | 40,34 | 40,22 | 41,37 | 40,38 | 39,96 | 41,15 | 40,95 | 41,90 |
| 72 | TrSC58v2_2586_HEIAH_TrGP63 | 44,04 | 41,40 | 41,53 | 41,53 | 41,53 | 41,53 | 41,53 | 41,53 | 41,53 | 41,53 | 40,93 | 40,93 | 40,93 | 40,64 | 40,60 | 40,60 | 40,32 | 40,41 | 40,32 | 38,92 | 39,00 | 40,39 | 39,92 | 41,19 |
| 73 | TrSC58v2_7404_HGIAH_TrGP63 | 45,26 | 43,91 | 43,70 | 43,49 | 43,49 | 43,49 | 43,49 | 43,49 | 43,49 | 43,49 | 42,86 | 42,86 | 42,86 | 41,93 | 41,68 | 41,68 | 41,39 | 41,52 | 42,20 | 41,63 | 41,21 | 42,27 | 41,77 | 43,20 |
| 74 | TrSC58v2_8159_HEIAH_TrGP63 | 43,09 | 42,59 | 42,39 | 42,18 | 42,18 | 42,18 | 42,18 | 42,18 | 42,18 | 42,18 | 42,18 | 42,18 | 42,18 | 38,81 | 38,97 | 38,97 | 38,48 | 38,51 | 39,92 | 37,91 | 38,32 | 40,00 | 39,52 | 41,01 |
| 75 | TrSC58v2_4723_HEIAH_TrGP63 | 44,00 | 42,44 | 42,23 | 42,02 | 42,02 | 42,02 | 42,02 | 42,02 | 42,02 | 42,02 | 41,60 | 41,60 | 41,60 | 38,99 | 38,95 | 38,95 | 38,45 | 38,48 | 39,71 | 38,91 | 38,91 | 39,30 | 39,71 | 40,39 |
| 76 | TrSC58v2_6284_HEIAH_TrGP63 | 45,89 | 43,28 | 43,28 | 43,49 | 43,49 | 43,49 | 43,49 | 43,49 | 43,49 | 43,49 | 42,86 | 42,86 | 42,86 | 40,25 | 40,21 | 40,21 | 39,71 | 40,00 | 40,54 | 38,70 | 39,33 | 40,00 | 39,51 | 41,47 |

| | TrGP63 Groups | 11 | 2 | 2 | 10 | 10 | 10 | 10 | 10 | 9 | 9 | 9 | 9 | 9 | 9 | 4 | 4 | 4 | 4 | 4 | 4 | 4 | 4 | 4 | 4 | 4 | 4 | 4 |
|---|---|---|---|---|---|---|---|---|---|---|---|---|---|---|---|---|---|---|---|---|---|---|---|---|---|---|---|---|
| 1 | TrSC58v2_1339_HEIAH_TrGP63 | 50,18 | 40,29 | 40,70 | 46,29 | 43,90 | 45,05 | 43,58 | 45,42 | 38,00 | 38,22 | 37,38 | 37,52 | 39,67 | 40,88 | 40,87 | 40,87 | 40,87 | 38,89 | 40,46 | 39,70 | 40,15 | 40,15 | 39,51 | 40,65 | 40,12 | 40,64 | 39,38 |
| 2 | TrSC58v2_2794_HEIAH_TrGP63 | 44,81 | 39,71 | 39,20 | 44,85 | 42,28 | 45,25 | 42,81 | 45,90 | 37,53 | 37,73 | 36,67 | 35,31 | 37,97 | 37,72 | 39,85 | 39,85 | 39,85 | 38,49 | 39,49 | 38,16 | 39,56 | 39,74 | 39,11 | 42,29 | 39,60 | 40,04 | 36,74 |
| 3 | TrSC58v2_9430_HEIAH_TrGP63 | 47,37 | 40,51 | 39,72 | 45,04 | 42,68 | 45,62 | 45,04 | 46,27 | 38,49 | 38,70 | 37,32 | 35,98 | 38,24 | 38,10 | 39,70 | 39,70 | 39,70 | 39,04 | 40,26 | 40,48 | 39,49 | 39,85 | 39,22 | 42,29 | 39,60 | 40,37 | 39,17 |
| 4 | TrSC58v2_1328_HEIAH_TrGP63 | 48,72 | 39,49 | 39,76 | 44,29 | 42,89 | 44,51 | 43,90 | 44,71 | 38,86 | 39,09 | 38,54 | 37,53 | 38,65 | 39,92 | 42,22 | 42,22 | 42,22 | 39,96 | 40,94 | 41,82 | 42,42 | 42,42 | 41,75 | 43,22 | 39,92 | 41,94 | 40,62 |
| 5 | TrSC58v2_1329_HEIAH_TrGP63 | 48,72 | 39,49 | 39,76 | 44,29 | 42,89 | 44,51 | 43,90 | 44,71 | 38,86 | 39,09 | 38,54 | 37,53 | 38,65 | 39,92 | 42,22 | 42,22 | 42,22 | 39,96 | 40,94 | 41,82 | 42,42 | 42,42 | 41,75 | 43,22 | 39,92 | 41,94 | 40,62 |
| 6 | TrSC58v2_1330_HEIAH_TrGP63 | 48,72 | 39,49 | 39,76 | 44,29 | 42,89 | 44,51 | 43,90 | 44,71 | 38,86 | 39,09 | 38,54 | 37,53 | 38,65 | 39,92 | 42,22 | 42,22 | 42,22 | 39,96 | 40,94 | 41,82 | 42,42 | 42,42 | 41,75 | 43,22 | 39,92 | 41,94 | 40,62 |
| 7 | TrSC58v2_1331_HEIAH_TrGP63 | 48,72 | 39,49 | 39,76 | 44,29 | 42,89 | 44,51 | 43,90 | 44,71 | 38,86 | 39,09 | 38,54 | 37,53 | 38,65 | 39,92 | 42,22 | 42,22 | 42,22 | 39,96 | 40,94 | 41,82 | 42,42 | 42,42 | 41,75 | 43,22 | 39,92 | 41,94 | 40,62 |
| 8 | TrSC58v2_1332_HEIAH_TrGP63 | 48,72 | 39,49 | 39,76 | 44,29 | 42,89 | 44,51 | 43,90 | 44,71 | 38,86 | 39,09 | 38,54 | 37,53 | 38,65 | 39,92 | 42,22 | 42,22 | 42,22 | 39,96 | 40,94 | 41,82 | 42,42 | 42,42 | 41,75 | 43,22 | 39,92 | 41,94 | 40,62 |
| 9 | TrSC58v2_1334_HEIAH_TrGP63 | 48,72 | 39,49 | 39,76 | 44,29 | 42,89 | 44,51 | 43,90 | 44,71 | 38,86 | 39,09 | 38,54 | 37,53 | 38,65 | 39,92 | 42,22 | 42,22 | 42,22 | 39,96 | 40,94 | 41,82 | 42,42 | 42,42 | 41,75 | 43,22 | 39,92 | 41,94 | 40,62 |
| 10 | TrSC58v2_1333_HEIAH_TrGP63 | 48,72 | 39,49 | 39,76 | 44,29 | 42,89 | 44,51 | 43,90 | 44,71 | 38,86 | 39,09 | 38,54 | 37,53 | 38,65 | 39,92 | 42,22 | 42,22 | 42,22 | 39,96 | 40,94 | 41,82 | 42,42 | 42,42 | 41,75 | 43,22 | 39,92 | 41,94 | 40,62 |
| 11 | TrSC58v2_6131_HEIAH_TrGP63 | 47,66 | 40,55 | 39,96 | 44,76 | 42,48 | 44,49 | 43,04 | 44,86 | 38,14 | 38,35 | 38,26 | 36,88 | 38,16 | 39,85 | 40,49 | 40,49 | 40,49 | 39,30 | 39,50 | 39,33 | 39,77 | 39,77 | 39,51 | 41,59 | 38,91 | 40,08 | 38,54 |
| 12 | TrSC58v2_6132_HEIAH_TrGP63 | 47,66 | 40,55 | 39,96 | 44,76 | 42,48 | 44,49 | 43,04 | 44,86 | 38,14 | 38,35 | 38,26 | 36,88 | 38,16 | 39,85 | 40,49 | 40,49 | 40,49 | 39,30 | 39,50 | 39,33 | 39,77 | 39,77 | 39,51 | 41,59 | 38,91 | 40,08 | 38,54 |
| 13 | TrSC58v2_6133_HEIAH_TrGP63 | 47,66 | 40,55 | 39,96 | 44,76 | 42,48 | 44,49 | 43,04 | 44,86 | 38,14 | 38,35 | 38,26 | 36,88 | 38,16 | 39,85 | 40,49 | 40,49 | 40,49 | 39,30 | 39,50 | 39,33 | 39,77 | 39,77 | 39,51 | 41,59 | 38,91 | 40,08 | 38,54 |
| 14 | TrSC58v2_8803_HEMAH_TrGP63 | 66,00 | 41,21 | 41,03 | 44,44 | 44,99 | 44,65 | 42,02 | 44,46 | 35,68 | 35,68 | 35,78 | 35,11 | 38,12 | 36,31 | 38,52 | 38,52 | 38,52 | 37,28 | 40,96 | 39,53 | 38,85 | 38,85 | 38,18 | 41,16 | 40,20 | 38,14 | 37,84 |
| 15 | TrSC58v2_8920_HEMAH_TrGP63 | 61,43 | 41,99 | 41,76 | 44,81 | 44,97 | 45,19 | 40,19 | 44,81 | 35,00 | 35,00 | 33,61 | 32,83 | 38,14 | 33,96 | 37,41 | 37,41 | 37,41 | 35,66 | 39,05 | 36,95 | 36,82 | 37,22 | 36,77 | 40,89 | 39,76 | 36,65 | 35,02 |
| 16 | TrSC58v2_9386_HEMAH_TrGP63 | 61,37 | 41,99 | 41,76 | 44,81 | 44,97 | 45,27 | 40,26 | 44,90 | 35,00 | 35,00 | 33,45 | 32,83 | 38,14 | 33,79 | 37,48 | 37,48 | 37,48 | 35,73 | 39,12 | 36,84 | 36,88 | 37,28 | 36,77 | 40,89 | 39,76 | 36,71 | 34,92 |
| 17 | TrSC58v2_6853_HEMAH_TrGP63 | 61,95 | 41,50 | 41,11 | 44,72 | 45,49 | 45,29 | 40,45 | 45,10 | 35,13 | 35,13 | 33,56 | 32,61 | 37,67 | 33,73 | 36,81 | 36,81 | 36,81 | 35,41 | 38,62 | 36,39 | 36,58 | 36,98 | 36,52 | 40,56 | 39,68 | 36,76 | 35,13 |
| 18 | TrSC58v2_8770_HEMAH_TrGP63 | 65,18 | 41,09 | 40,59 | 43,91 | 44,07 | 45,54 | 41,59 | 45,44 | 35,85 | 35,85 | 35,28 | 34,40 | 37,68 | 35,44 | 39,20 | 39,20 | 39,20 | 37,58 | 41,24 | 38,78 | 39,15 | 38,95 | 37,04 | 40,15 | 39,09 | 38,98 | 37,63 |
| 19 | TrSC58v2_6130_HEMTH_TrGP63 | 74,64 | 41,35 | 41,50 | 45,25 | 44,22 | 45,16 | 38,88 | 44,61 | 35,48 | 35,48 | 35,45 | 35,18 | 40,08 | 37,60 | 39,75 | 39,75 | 39,75 | 37,21 | 38,10 | 36,56 | 37,25 | 37,84 | 36,98 | 41,24 | 39,02 | 38,06 | 34,20 |
| 20 | TrSC58v2_560_RGLAY_TrGP63 | 74,53 | 42,36 | 41,15 | 46,17 | 46,01 | 46,22 | 40,52 | 46,41 | 37,25 | 37,25 | 35,21 | 35,58 | 39,00 | 35,68 | 37,59 | 37,59 | 37,59 | 37,36 | 36,71 | 35,46 | 35,33 | 35,71 | 35,05 | 39,53 | 37,94 | 36,49 | 34,00 |
| 21 | TrSC58v2_8766_RGLAY_TrGP63 | 64,80 | 43,30 | 41,83 | 49,13 | 46,42 | 50,29 | 45,48 | 50,48 | 37,68 | 37,68 | 35,21 | 35,90 | 38,89 | 36,12 | 37,04 | 37,04 | 37,04 | 37,38 | 35,60 | 34,69 | 36,14 | 36,61 | 35,62 | 39,07 | 38,37 | 36,45 | 33,39 |
| 22 | TrSC58v2_9126_HEMAH_TrGP63 | 75,95 | 41,37 | 40,19 | 46,67 | 44,38 | 45,10 | 39,07 | 44,18 | 36,63 | 36,63 | 35,44 | 35,65 | 40,04 | 35,30 | 37,91 | 37,91 | 37,91 | 36,30 | 36,52 | 35,91 | 36,75 | 36,98 | 36,96 | 40,81 | 38,00 | 38,14 | 34,79 |
| 23 | TrSC58v2_6495_HEMAH_TrGP63 | 80,93 | 41,28 | 40,37 | 46,38 | 44,27 | 44,55 | 38,17 | 43,81 | 37,69 | 37,69 | 36,18 | 34,97 | 39,30 | 37,02 | 38,94 | 38,94 | 38,94 | 37,12 | 37,76 | 36,25 | 37,88 | 38,13 | 37,79 | 41,93 | 38,89 | 39,36 | 34,39 |
| 24 | TrSC58v2_8644_HEMAH_TrGP63 | 93,23 | 42,45 | 42,52 | 45,47 | 42,62 | 42,99 | 37,23 | 42,23 | 38,51 | 38,51 | 36,95 | 36,93 | 40,28 | 37,69 | 39,42 | 39,42 | 39,42 | 37,00 | 38,08 | 36,18 | 37,54 | 37,80 | 37,66 | 42,07 | 39,84 | 38,93 | 34,53 |
| 25 | TrSC58v2_9275_HEMAH_TrGP63 | 100,00 | 42,45 | 42,52 | 45,08 | 43,25 | 43,94 | 38,01 | 43,18 | 37,33 | 37,33 | 36,45 | 36,44 | 40,08 | 37,19 | 39,96 | 39,96 | 39,96 | 37,73 | 38,43 | 36,65 | 38,57 | 38,85 | 38,61 | 43,03 | 40,04 | 39,46 | 34,37 |
| 26 | TrSC58v2_8384_HELAH_TrGP63 | 42,45 | 100,00 | 96,53 | 47,10 | 43,14 | 44,44 | 41,83 | 43,90 | 38,72 | 38,72 | 38,33 | 38,46 | 41,00 | 39,74 | 38,04 | 38,04 | 38,04 | 36,87 | 37,55 | 38,69 | 38,51 | 38,30 | 37,34 | 39,85 | 36,21 | 38,33 | 37,99 |
| 27 | TrSC58v2_559_HELAH_TrGP63 | 42,52 | 96,53 | 100,00 | 45,51 | 41,62 | 43,66 | 41,05 | 42,74 | 39,64 | 39,64 | 37,16 | 37,40 | 39,37 | 39,52 | 37,84 | 37,84 | 37,84 | 36,64 | 37,42 | 38,49 | 38,08 | 37,88 | 37,02 | 38,95 | 35,99 | 38,88 | 37,10 |
| 28 | TrSC58v2_8165_HAIAH_TrGP63 | 45,08 | 47,10 | 45,51 | 100,00 | 72,17 | 75,74 | 74,81 | 76,11 | 41,61 | 41,61 | 40,15 | 40,42 | 43,60 | 41,17 | 40,99 | 40,99 | 40,99 | 39,23 | 38,74 | 39,13 | 41,18 | 41,18 | 41,24 | 43,44 | 40,43 | 40,49 | 38,16 |
| 29 | TrSC58v2_9363_HAIAH_TrGP63 | 43,25 | 43,14 | 41,62 | 72,17 | 100,00 | 91,24 | 92,03 | 91,24 | 38,00 | 38,00 | 37,83 | 36,82 | 39,68 | 39,92 | 39,76 | 39,76 | 39,76 | 38,04 | 38,87 | 39,20 | 39,56 | 39,56 | 39,92 | 41,08 | 37,68 | 38,76 | 38,15 |
| 30 | TrSC58v2_9356_HAIAH_TrGP63 | 43,94 | 44,44 | 43,66 | 75,74 | 91,24 | 100,00 | 94,83 | 95,54 | 39,07 | 39,07 | 39,39 | 38,31 | 42,65 | 40,23 | 38,97 | 38,97 | 38,97 | 38,40 | 39,69 | 40,27 | 39,16 | 39,16 | 38,68 | 40,66 | 38,29 | 38,55 | 37,92 |
| 31 | TrSC58v2_3271_HAIAH_TrGP63 | 38,01 | 41,83 | 41,05 | 74,81 | 92,03 | 94,83 | 100,00 | 96,43 | 37,63 | 37,63 | 35,86 | 34,84 | 40,55 | 36,75 | 37,27 | 37,27 | 37,27 | 35,93 | 36,79 | 34,31 | 35,83 | 36,09 | 35,55 | 40,19 | 37,02 | 36,04 | 31,28 |
| 32 | TrSC58v2_3280_HAIAH_TrGP63 | 43,18 | 43,90 | 42,74 | 76,11 | 91,24 | 95,54 | 96,43 | 100,00 | 38,85 | 38,85 | 39,77 | 38,31 | 42,03 | 39,85 | 38,40 | 38,40 | 38,40 | 37,99 | 39,24 | 39,81 | 39,35 | 39,35 | 38,89 | 40,90 | 38,16 | 37,90 | 37,72 |
| 33 | TrSC58v2_5263_HEIAH_TrGP63 | 37,33 | 38,72 | 39,64 | 41,61 | 38,00 | 39,07 | 37,63 | 38,85 | 100,00 | 99,51 | 77,02 | 74,55 | 62,57 | 58,54 | 33,47 | 33,47 | 33,47 | 33,59 | 33,46 | 34,33 | 33,40 | 33,20 | 33,33 | 37,33 | 35,49 | 34,83 | 33,68 |
| 34 | TrSC58v2_5266_HEIAH_TrGP63 | 37,33 | 38,72 | 39,64 | 41,61 | 38,00 | 39,07 | 37,63 | 38,85 | 99,51 | 100,00 | 77,02 | 74,42 | 62,38 | 58,71 | 33,47 | 33,47 | 33,47 | 33,97 | 33,65 | 34,52 | 33,60 | 33,40 | 33,55 | 37,33 | 35,71 | 35,03 | 33,96 |
| 35 | TrSC58v2_6663_HEIAH_TrGP63 | 36,45 | 38,33 | 37,16 | 40,15 | 37,83 | 39,39 | 35,86 | 39,77 | 77,02 | 77,02 | 100,00 | 81,28 | 60,73 | 65,60 | 32,64 | 32,64 | 32,64 | 33,27 | 33,44 | 31,23 | 30,90 | 31,18 | 30,96 | 35,20 | 33,14 | 32,15 | 30,61 |
| 36 | TrSC58v2_9131_HEIAH_TrGP63 | 36,44 | 38,46 | 37,40 | 40,42 | 36,82 | 38,31 | 34,84 | 38,31 | 74,55 | 74,42 | 81,28 | 100,00 | 64,34 | 64,80 | 32,76 | 32,76 | 32,76 | 34,23 | 32,69 | 30,72 | 30,36 | 30,78 | 30,91 | 35,20 | 33,40 | 31,75 | 29,41 |
| 37 | TrSC58v2_9025_HEIAH_TrGP63 | 40,08 | 41,00 | 39,37 | 43,60 | 39,68 | 42,65 | 40,55 | 42,03 | 62,57 | 62,38 | 60,73 | 64,34 | 100,00 | 73,70 | 32,49 | 32,49 | 32,49 | 34,05 | 33,58 | 32,80 | 32,78 | 32,60 | 31,68 | 35,67 | 32,95 | 32,89 | 31,20 |
| 38 | TrSC58v2_4259_HEIAH_TrGP63 | 37,19 | 39,74 | 39,52 | 41,17 | 39,92 | 40,23 | 36,75 | 39,85 | 58,54 | 58,71 | 65,60 | 64,80 | 73,70 | 100,00 | 32,09 | 32,09 | 32,09 | 31,60 | 31,83 | 29,83 | 30,08 | 30,66 | 32,32 | 37,04 | 33,66 | 32,23 | 28,61 |
| 39 | TrSC58v2_631_HEIGH_TrGP63 | 39,96 | 38,04 | 37,84 | 40,99 | 39,76 | 38,97 | 37,27 | 38,40 | 33,47 | 33,47 | 32,64 | 32,76 | 32,49 | 32,09 | 100,00 | 99,85 | 99,85 | 85,34 | 66,45 | 66,34 | 67,24 | 68,78 | 65,69 | 69,48 | 68,84 | 67,75 | 65,03 |
| 40 | TrSC58v2_632_HEIGH_TrGP63 | 39,96 | 38,04 | 37,84 | 40,99 | 39,76 | 38,97 | 37,27 | 38,40 | 33,47 | 33,47 | 32,64 | 32,76 | 32,49 | 32,09 | 99,85 | 100,00 | 100,00 | 85,34 | 66,45 | 66,34 | 67,40 | 68,94 | 65,86 | 69,69 | 69,01 | 67,91 | 65,03 |
| 41 | TrSC58v2_633_HEIGH_TrGP63 | 39,96 | 38,04 | 37,84 | 40,99 | 39,76 | 38,97 | 37,27 | 38,40 | 33,47 | 33,47 | 32,64 | 32,76 | 32,49 | 32,09 | 99,85 | 100,00 | 100,00 | 85,34 | 66,45 | 66,34 | 67,40 | 68,94 | 65,86 | 69,69 | 69,01 | 67,91 | 65,03 |
| 42 | TrSC58v2_4929_HEIGH_TrGP63 | 37,73 | 36,87 | 36,64 | 39,23 | 38,04 | 38,40 | 35,93 | 37,99 | 33,59 | 33,97 | 33,27 | 34,23 | 34,05 | 31,60 | 85,34 | 85,34 | 85,34 | 100,00 | 63,05 | 61,57 | 63,55 | 64,96 | 58,02 | 66,01 | 66,46 | 64,32 | 60,84 |
| 43 | TrSC58v2_6472_HEIGH_TrGP63 | 38,43 | 37,55 | 37,42 | 38,74 | 38,87 | 39,69 | 36,79 | 39,24 | 33,46 | 33,65 | 33,44 | 32,69 | 33,58 | 31,83 | 66,45 | 66,45 | 66,45 | 63,05 | 100,00 | 89,85 | 64,84 | 65,99 | 60,13 | 69,64 | 67,86 | 71,49 | 72,52 |
| 44 | TrSC58v2_7196_HEIGH_TrGP63 | 36,65 | 38,69 | 38,49 | 39,13 | 39,20 | 40,27 | 34,31 | 39,81 | 34,33 | 34,52 | 31,23 | 30,72 | 32,80 | 29,83 | 66,34 | 66,34 | 66,34 | 61,57 | 89,85 | 100,00 | 61,15 | 63,73 | 53,95 | 68,74 | 66,98 | 73,60 | 74,48 |
| 45 | TrSC58v2_9139_HEIGH_TrGP63 | 38,57 | 38,51 | 38,08 | 41,18 | 39,56 | 39,16 | 35,83 | 39,35 | 33,40 | 33,60 | 30,90 | 30,36 | 32,78 | 30,08 | 67,24 | 67,40 | 67,40 | 63,55 | 64,84 | 61,15 | 100,00 | 94,40 | 76,50 | 97,53 | 68,99 | 67,09 | 60,32 |
| 46 | TrSC58v2_9136_HEIGH_TrGP63 | 38,85 | 38,30 | 37,88 | 41,18 | 39,56 | 39,16 | 36,09 | 39,35 | 33,20 | 33,40 | 31,18 | 30,78 | 32,60 | 30,66 | 68,78 | 68,94 | 68,94 | 64,96 | 65,99 | 63,73 | 94,40 | 100,00 | 74,22 | 97,53 | 68,99 | 67,88 | 62,96 |
| 47 | TrSC58v2_9135_HEIGH_TrGP63 | 38,61 | 37,34 | 37,02 | 41,24 | 39,92 | 38,68 | 35,55 | 38,89 | 33,33 | 33,55 | 30,96 | 30,91 | 31,68 | 32,32 | 65,69 | 65,86 | 65,86 | 58,02 | 60,13 | 53,95 | 76,50 | 74,22 | 100,00 | 97,53 | 65,26 | 61,10 | 50,99 |
| 48 | TrSC58v2_9137_HEIGH_TrGP63 | 43,03 | 39,85 | 38,95 | 43,44 | 41,08 | 40,66 | 40,19 | 40,90 | 37,33 | 37,33 | 35,20 | 35,20 | 35,67 | 37,04 | 69,48 | 69,69 | 69,69 | 66,01 | 69,64 | 68,74 | 97,53 | 97,53 | 97,53 | 100,00 | 66,46 | 71,60 | 68,14 |
| 49 | TrSC58v2_9032_HEIGH_TrGP63 | 40,04 | 36,21 | 35,99 | 40,43 | 37,68 | 38,29 | 37,02 | 38,16 | 35,49 | 35,71 | 33,14 | 33,40 | 32,95 | 33,66 | 68,84 | 69,01 | 69,01 | 66,46 | 67,86 | 66,98 | 68,99 | 68,99 | 65,26 | 66,46 | 100,00 | 68,20 | 65,04 |
| 50 | TrSC58v2_5291_HEIGH_TrGP63 | 39,46 | 38,33 | 38,88 | 40,49 | 38,76 | 38,55 | 36,04 | 37,90 | 34,83 | 35,03 | 32,15 | 31,75 | 32,89 | 32,23 | 67,75 | 67,91 | 67,91 | 64,32 | 71,49 | 73,60 | 67,09 | 67,88 | 61,10 | 71,60 | 68,20 | 100,00 | 70,98 |
| 51 | TrSC58v2_7524_HEIAH_TrGP63 | 34,37 | 37,99 | 37,10 | 38,16 | 38,15 | 37,92 | 31,28 | 37,72 | 33,68 | 33,96 | 30,61 | 29,41 | 31,20 | 28,61 | 65,03 | 65,03 | 65,03 | 60,84 | 72,52 | 74,48 | 60,32 | 62,96 | 50,99 | 68,14 | 65,04 | 70,98 | 100,00 |
| 52 | TrSC58v2_7598_QKIGH_TrGP63 | 35,05 | 38,05 | 37,86 | 39,66 | 39,56 | 39,43 | 34,14 | 39,35 | 32,67 | 32,79 | 29,89 | 28,45 | 29,98 | 28,38 | 71,21 | 71,36 | 71,36 | 64,03 | 71,49 | 67,76 | 65,25 | 68,36 | 56,13 | 70,52 | 69,31 | 70,40 | 64,98 |
| 53 | TrSC58v2_9319_HEIGH_TrGP63 | 40,29 | 38,79 | 38,38 | 37,80 | 37,58 | 37,64 | 36,46 | 37,38 | 34,33 | 34,52 | 34,36 | 33,15 | 34,59 | 33,87 | 70,28 | 70,28 | 70,28 | 67,89 | 69,02 | 67,43 | 70,10 | 71,45 | 68,26 | 70,35 | 71,16 | 69,02 | 67,45 |
| 54 | TrSC58v2_4655_HEIGH_TrGP63 | 39,37 | 39,15 | 39,12 | 40,34 | 38,45 | 39,41 | 38,96 | 38,99 | 33,48 | 33,70 | 33,27 | 33,40 | 34,55 | 33,79 | 71,26 | 71,26 | 71,26 | 66,46 | 70,32 | 69,04 | 72,47 | 72,47 | 73,56 | 74,56 | 68,80 | 71,76 | 72,78 |
| 55 | TrSC58v2_4642_HEIGH_TrGP63 | 39,02 | 39,48 | 39,43 | 40,59 | 39,54 | 39,54 | 37,73 | 39,08 | 32,22 | 32,34 | 32,88 | 32,03 | 34,10 | 31,43 | 71,45 | 71,45 | 71,45 | 67,63 | 64,63 | 62,96 | 68,06 | 68,26 | 66,78 | 74,78 | 72,23 | 66,57 | 64,64 |
| 56 | TrSC58v2_4641_HEIGH_TrGP63 | 38,32 | 39,70 | 39,63 | 40,59 | 39,54 | 39,54 | 36,43 | 39,08 | 32,22 | 32,34 | 32,36 | 31,46 | 33,58 | 30,61 | 71,29 | 71,29 | 71,29 | 68,52 | 64,19 | 60,30 | 66,96 | 67,62 | 65,30 | 74,78 | 72,42 | 65,97 | 61,76 |
| 57 | TrSC58v2_4649_HEIGH_TrGP63 | 39,16 | 39,70 | 39,63 | 40,59 | 39,54 | 39,54 | 37,73 | 39,08 | 33,46 | 33,66 | 33,57 | 33,74 | 34,92 | 33,16 | 71,45 | 71,45 | 71,45 | 68,03 | 67,72 | 66,67 | 71,03 | 71,24 | 69,20 | 74,78 | 72,23 | 69,54 | 70,57 |
| 58 | TrSC58v2_4654_HEIGH_TrGP63 | 37,56 | 39,70 | 39,63 | 40,59 | 39,54 | 39,54 | 35,37 | 39,08 | 32,22 | 32,34 | 31,87 | 30,56 | 33,33 | 30,18 | 71,45 | 71,45 | 71,45 | 68,79 | 64,34 | 59,38 | 66,96 | 67,62 | 65,30 | 74,78 | 72,23 | 66,12 | 60,00 |
| 59 | TrSC58v2_4653_HEIGH_TrGP63 | 38,51 | 38,76 | 37,87 | 38,66 | 36,63 | 36,92 | 37,21 | 36,63 | 38,44 | 38,44 | 36,96 | 38,51 | 37,75 | 37,75 | 71,00 | 71,00 | 71,00 | 70,81 | 70,77 | 68,83 | 70,73 | 70,73 | 70,73 | 71,35 | 66,76 | 70,54 | 76,76 |
| 60 | TrSC58v2_4652_HEIGH_TrGP63 | 33,33 | 38,39 | 38,41 | 38,67 | 39,54 | 36,47 | 32,60 | 36,02 | 30,02 | 30,15 | 28,39 | 28,20 | 32,84 | 28,62 | 63,06 | 63,06 | 63,06 | 57,31 | 59,58 | 55,59 | 61,88 | 62,10 | 69,50 | 74,78 | 67,74 | 60,86 | 56,19 |
| 61 | TrSC58v2_4656_HEIGH_TrGP63 | 32,23 | 37,12 | 36,88 | 35,35 | 36,41 | 32,83 | 32,42 | 32,55 | 30,20 | 30,33 | 29,92 | 29,96 | 31,89 | 28,93 | 61,54 | 61,54 | 61,54 | 58,46 | 59,92 | 58,54 | 62,57 | 62,57 | 71,37 | 70,68 | 64,11 | 61,20 | 61,21 |
| 62 | TrSC58v2_9446_HETGH_TrGP63 | 37,44 | 39,06 | 39,15 | 41,18 | 39,16 | 38,86 | 34,18 | 38,02 | 33,39 | 33,52 | 31,93 | 30,14 | 31,87 | 30,87 | 74,77 | 74,92 | 74,92 | 68,15 | 69,75 | 67,82 | 68,54 | 69,90 | 62,73 | 71,96 | 73,54 | 72,55 | 71,98 |
| 63 | TrSC58v2_4795_HEIAH_TrGP63 | 39,28 | 39,83 | 39,06 | 41,44 | 39,96 | 39,31 | 37,14 | 38,86 | 34,17 | 34,37 | 32,83 | 32,83 | 33,78 | 32,77 | 71,47 | 71,63 | 71,63 | 69,59 | 68,97 | 69,16 | 68,72 | 70,17 | 66,89 | 72,37 | 69,08 | 73,68 | 72,46 |
| 64 | TrSC58v2_7306_HEIAH_TrGP63 | 39,96 | 38,82 | 38,74 | 41,18 | 40,56 | 37,90 | 37,38 | 37,64 | 34,42 | 34,62 | 33,63 | 33,57 | 33,66 | 32,91 | 75,40 | 75,24 | 75,24 | 71,09 | 68,59 | 68,80 | 71,41 | 72,73 | 70,07 | 72,58 | 69,66 | 74,40 | 73,95 |
| 65 | TrSC58v2_4835_HEMAH_TrGP63 | 39,08 | 40,72 | 39,96 | 38,66 | 38,27 | 38,59 | 38,56 | 38,59 | 38,90 | 38,90 | 37,30 | 36,76 | 38,14 | 38,51 | 37,28 | 37,28 | 37,28 | 37,98 | 36,18 | 35,69 | 36,09 | 36,09 | 35,39 | 35,57 | 36,26 | 35,77 | 37,01 |
| 66 | TrSC58v2_3991_HEIVH_TrGP63 | 40,50 | 42,92 | 42,44 | 42,92 | 42,26 | 44,31 | 43,71 | 43,78 | 36,55 | 36,78 | 37,04 | 36,42 | 38,64 | 38,48 | 43,97 | 43,97 | 43,97 | 43,22 | 43,31 | 44,14 | 42,64 | 42,64 | 41,60 | 43,14 | 41,15 | 43,95 | 42,71 |
| 67 | TrSC58v2_4583_HEIAH_TrGP63 | 40,90 | 39,03 | 38,87 | 44,35 | 40,42 | 41,54 | 39,53 | 41,63 | 35,62 | 35,76 | 34,84 | 34,12 | 36,20 | 36,15 | 42,94 | 42,94 | 42,94 | 42,54 | 40,50 | 40,49 | 40,75 | 40,75 | 40,43 | 43,08 | 42,20 | 41,77 | 37,81 |
| 68 | TrSC58v2_7126_HEIAH_TrGP63 | 41,60 | 44,32 | 43,11 | 42,83 | 41,80 | 42,15 | 42,36 | 42,24 | 39,15 | 38,93 | 37,92 | 38,70 | 40,43 | 38,65 | 44,13 | 44,13 | 44,13 | 43,97 | 42,41 | 43,44 | 43,77 | 43,77 | 43,73 | 44,76 | 41,59 | 41,83 | 41,28 |
| 69 | TrSC58v2_8509_HEIAH_TrGP63 | 42,55 | 43,58 | 43,82 | 42,71 | 39,92 | 41,79 | 41,58 | 41,79 | 41,26 | 41,26 | 39,14 | 38,52 | 38,89 | 38,92 | 45,19 | 45,19 | 45,19 | 44,42 | 44,72 | 44,96 | 44,40 | 44,40 | 43,50 | 44,08 | 41,76 | 45,28 | 44,56 |
| 70 | TrSC58v2_2748_HEIAH_TrGP63 | 41,90 | 44,04 | 44,14 | 42,41 | 39,63 | 41,36 | 41,15 | 41,36 | 38,93 | 38,93 | 36,81 | 36,81 | 37,99 | 37,40 | 43,53 | 43,53 | 43,53 | 43,76 | 43,70 | 43,95 | 42,55 | 42,55 | 41,65 | 42,67 | 41,29 | 44,60 | 44,56 |
| 71 | TrSC58v2_6871_HEIAH_TrGP63 | 42,55 | 43,58 | 43,93 | 43,04 | 39,83 | 41,79 | 41,58 | 41,79 | 40,09 | 40,09 | 37,63 | 38,04 | 39,63 | 39,05 | 43,92 | 43,92 | 43,92 | 43,76 | 43,70 | 44,76 | 43,53 | 43,53 | 42,63 | 43,76 | 41,88 | 45,38 | 43,93 |
| 72 | TrSC58v2_2586_HEIAH_TrGP63 | 41,60 | 43,54 | 43,78 | 44,20 | 40,53 | 41,20 | 41,02 | 41,29 | 39,73 | 39,73 | 38,05 | 38,25 | 38,40 | 37,83 | 44,66 | 44,66 | 44,66 | 43,86 | 43,87 | 44,51 | 42,78 | 42,78 | 42,53 | 44,08 | 41,90 | 45,21 | 43,27 |
| 73 | TrSC58v2_7404_HGIAH_TrGP63 | 43,84 | 43,12 | 43,40 | 43,33 | 39,29 | 40,94 | 40,72 | 41,15 | 39,39 | 39,39 | 37,91 | 38,11 | 38,89 | 38,10 | 44,99 | 44,99 | 44,99 | 43,76 | 44,92 | 44,96 | 43,42 | 43,42 | 42,72 | 43,64 | 42,55 | 45,47 | 45,61 |
| 74 | TrSC58v2_8159_HEIAH_TrGP63 | 41,44 | 43,05 | 43,12 | 42,04 | 38,89 | 40,08 | 40,29 | 40,08 | 38,50 | 38,50 | 36,35 | 36,95 | 36,90 | 36,31 | 44,42 | 44,42 | 44,42 | 45,09 | 43,94 | 45,96 | 44,04 | 44,04 | 43,41 | 44,74 | 41,27 | 44,32 | 44,79 |
| 75 | TrSC58v2_4723_HEIAH_TrGP63 | 40,39 | 42,43 | 43,10 | 43,45 | 40,25 | 42,43 | 42,22 | 42,64 | 40,33 | 40,33 | 38,24 | 38,45 | 38,60 | 38,02 | 45,10 | 45,10 | 45,10 | 45,30 | 44,72 | 45,97 | 44,71 | 44,71 | 44,20 | 44,64 | 41,88 | 45,78 | 46,03 |
| 76 | TrSC58v2_6284_HEIAH_TrGP63 | 41,68 | 42,20 | 42,77 | 41,67 | 38,25 | 40,30 | 40,09 | 40,51 | 40,09 | 40,09 | 38,52 | 38,52 | 38,48 | 38,30 | 43,61 | 43,61 | 43,61 | 43,76 | 43,90 | 45,36 | 43,81 | 43,81 | 43,11 | 43,86 | 41,76 | 45,47 | 44,35 |

| | TrGP63 Groups | 4 | 4 | 4 | 4 | 4 | 4 | 4 | 4 | 4 | 4 | 4 | 4 | 4 | 3 | 7 | 5 | 8 | 6 | 6 | 6 | 6 | 6 | 6 | 6 | 6 |
|---|---|---|---|---|---|---|---|---|---|---|---|---|---|---|---|---|---|---|---|---|---|---|---|---|---|---|
| 1 | TrSC58v2_1339_HEIAH_TrGP63 | 40,64 | 40,83 | 40,62 | 42,64 | 42,34 | 42,64 | 42,34 | 37,28 | 37,59 | 33,33 | 41,39 | 41,56 | 42,07 | 41,49 | 45,90 | 39,44 | 41,13 | 42,95 | 44,00 | 44,00 | 44,04 | 45,26 | 43,09 | 44,00 | 45,89 |
| 2 | TrSC58v2_2794_HEIAH_TrGP63 | 38,47 | 42,91 | 41,36 | 41,26 | 40,39 | 41,26 | 40,04 | 40,23 | 36,28 | 35,04 | 38,60 | 40,70 | 40,75 | 40,46 | 42,89 | 40,23 | 39,92 | 44,33 | 44,54 | 44,12 | 41,40 | 43,91 | 42,59 | 42,44 | 43,28 |
| 3 | TrSC58v2_9430_HEIAH_TrGP63 | 40,15 | 42,94 | 41,24 | 41,39 | 41,19 | 41,39 | 41,19 | 40,52 | 37,77 | 35,34 | 40,07 | 40,89 | 40,38 | 41,00 | 42,28 | 40,85 | 39,84 | 44,33 | 44,33 | 43,91 | 41,53 | 43,70 | 42,39 | 42,23 | 43,28 |
| 4 | TrSC58v2_1328_HEIAH_TrGP63 | 41,13 | 44,06 | 41,44 | 42,42 | 42,42 | 42,42 | 42,42 | 39,66 | 40,81 | 36,53 | 41,13 | 41,82 | 41,33 | 40,66 | 43,44 | 41,82 | 40,85 | 43,91 | 43,91 | 43,70 | 41,53 | 43,49 | 42,18 | 42,02 | 43,49 |
| 5 | TrSC58v2_1329_HEIAH_TrGP63 | 41,13 | 44,06 | 41,44 | 42,42 | 42,42 | 42,42 | 42,42 | 39,66 | 40,81 | 36,53 | 41,13 | 41,82 | 41,33 | 40,66 | 43,44 | 41,82 | 40,85 | 43,91 | 43,91 | 43,70 | 41,53 | 43,49 | 42,18 | 42,02 | 43,49 |
| 6 | TrSC58v2_1330_HEIAH_TrGP63 | 41,13 | 44,06 | 41,44 | 42,42 | 42,42 | 42,42 | 42,42 | 39,66 | 40,81 | 36,53 | 41,13 | 41,82 | 41,33 | 40,66 | 43,44 | 41,82 | 40,85 | 43,91 | 43,91 | 43,70 | 41,53 | 43,49 | 42,18 | 42,02 | 43,49 |
| 7 | TrSC58v2_1331_HEIAH_TrGP63 | 41,13 | 44,06 | 41,44 | 42,42 | 42,42 | 42,42 | 42,42 | 39,66 | 40,81 | 36,53 | 41,13 | 41,82 | 41,33 | 40,66 | 43,44 | 41,82 | 40,85 | 43,91 | 43,91 | 43,70 | 41,53 | 43,49 | 42,18 | 42,02 | 43,49 |
| 8 | TrSC58v2_1332_HEIAH_TrGP63 | 41,13 | 44,06 | 41,44 | 42,42 | 42,42 | 42,42 | 42,42 | 39,66 | 40,81 | 36,53 | 41,13 | 41,82 | 41,33 | 40,66 | 43,44 | 41,82 | 40,85 | 43,91 | 43,91 | 43,70 | 41,53 | 43,49 | 42,18 | 42,02 | 43,49 |
| 9 | TrSC58v2_1334_HEIAH_TrGP63 | 41,13 | 44,06 | 41,44 | 42,42 | 42,42 | 42,42 | 42,42 | 39,66 | 40,81 | 36,53 | 41,13 | 41,82 | 41,33 | 40,66 | 43,44 | 41,82 | 40,85 | 43,91 | 43,91 | 43,70 | 41,53 | 43,49 | 42,18 | 42,02 | 43,49 |
| 10 | TrSC58v2_1333_HEIAH_TrGP63 | 41,13 | 44,06 | 41,44 | 42,42 | 42,42 | 42,42 | 42,42 | 39,66 | 40,81 | 36,53 | 41,13 | 41,82 | 41,33 | 40,66 | 43,44 | 41,82 | 40,85 | 43,91 | 43,91 | 43,70 | 41,53 | 43,49 | 42,18 | 42,02 | 43,49 |
| 11 | TrSC58v2_6131_HEIAH_TrGP63 | 40,19 | 41,51 | 39,92 | 41,79 | 41,32 | 41,79 | 41,32 | 39,37 | 36,78 | 33,83 | 39,25 | 40,15 | 39,89 | 41,70 | 44,47 | 39,96 | 41,25 | 43,70 | 43,49 | 43,07 | 40,93 | 42,86 | 42,18 | 41,60 | 42,86 |
| 12 | TrSC58v2_6132_HEIAH_TrGP63 | 40,19 | 41,51 | 39,92 | 41,79 | 41,32 | 41,79 | 41,32 | 39,37 | 36,78 | 33,83 | 39,25 | 40,15 | 39,89 | 41,70 | 44,47 | 39,96 | 41,25 | 43,70 | 43,49 | 43,07 | 40,93 | 42,86 | 42,18 | 41,60 | 42,86 |
| 13 | TrSC58v2_6133_HEIAH_TrGP63 | 40,19 | 41,51 | 39,92 | 41,79 | 41,32 | 41,79 | 41,32 | 39,37 | 36,78 | 33,83 | 39,25 | 40,15 | 39,89 | 41,70 | 44,47 | 39,96 | 41,25 | 43,70 | 43,49 | 43,07 | 40,93 | 42,86 | 42,18 | 41,60 | 42,86 |
| 14 | TrSC58v2_8803_HEMAH_TrGP63 | 38,75 | 39,08 | 39,18 | 39,07 | 38,79 | 39,07 | 38,64 | 37,61 | 36,80 | 32,78 | 39,61 | 40,44 | 40,49 | 38,35 | 41,78 | 40,82 | 42,23 | 40,46 | 41,09 | 40,88 | 40,64 | 41,93 | 38,81 | 38,99 | 40,25 |
| 15 | TrSC58v2_8920_HEMAH_TrGP63 | 36,04 | 38,53 | 39,14 | 37,72 | 37,02 | 37,72 | 35,80 | 37,82 | 34,78 | 32,78 | 37,09 | 38,56 | 39,16 | 38,51 | 41,96 | 40,59 | 42,00 | 40,42 | 41,05 | 40,63 | 40,60 | 41,68 | 38,97 | 38,95 | 40,21 |
| 16 | TrSC58v2_9386_HEMAH_TrGP63 | 35,93 | 38,60 | 39,14 | 37,79 | 37,09 | 37,79 | 35,86 | 37,82 | 34,83 | 32,85 | 37,31 | 38,62 | 39,23 | 38,51 | 41,96 | 40,59 | 42,00 | 40,42 | 41,05 | 40,63 | 40,60 | 41,68 | 38,97 | 38,95 | 40,21 |
| 17 | TrSC58v2_6853_HEMAH_TrGP63 | 35,98 | 38,28 | 38,85 | 37,83 | 37,13 | 37,83 | 35,91 | 37,71 | 34,22 | 32,51 | 36,69 | 38,31 | 38,91 | 38,64 | 42,07 | 40,70 | 42,32 | 40,13 | 40,55 | 40,34 | 40,32 | 41,39 | 38,48 | 38,45 | 39,71 |
| 18 | TrSC58v2_8770_HEMAH_TrGP63 | 38,35 | 39,80 | 38,00 | 39,17 | 38,72 | 39,17 | 37,29 | 37,43 | 35,45 | 33,26 | 38,70 | 40,87 | 40,52 | 38,76 | 41,05 | 41,01 | 42,47 | 40,22 | 40,65 | 40,22 | 40,41 | 41,52 | 38,51 | 38,48 | 40,00 |
| 19 | TrSC58v2_6130_HEMTH_TrGP63 | 36,78 | 41,03 | 39,96 | 39,15 | 38,46 | 39,15 | 37,72 | 39,20 | 34,41 | 33,26 | 37,28 | 38,22 | 38,85 | 39,88 | 40,76 | 40,00 | 42,41 | 41,58 | 41,16 | 41,37 | 40,32 | 42,20 | 39,92 | 39,71 | 40,54 |
| 20 | TrSC58v2_560_RGLAY_TrGP63 | 36,00 | 38,20 | 38,04 | 37,68 | 37,06 | 37,68 | 36,35 | 37,32 | 32,68 | 32,09 | 35,63 | 36,99 | 37,50 | 39,05 | 40,69 | 39,81 | 40,92 | 41,00 | 40,17 | 40,38 | 38,92 | 41,63 | 37,91 | 38,91 | 38,70 |
| 21 | TrSC58v2_8766_RGLAY_TrGP63 | 35,92 | 37,31 | 37,63 | 37,41 | 36,88 | 37,41 | 36,59 | 36,75 | 34,09 | 32,33 | 35,68 | 36,60 | 36,59 | 39,05 | 40,89 | 39,96 | 41,40 | 41,21 | 40,17 | 39,96 | 39,00 | 41,21 | 38,32 | 38,91 | 39,33 |
| 22 | TrSC58v2_9126_HEMAH_TrGP63 | 35,90 | 39,46 | 38,22 | 38,17 | 37,35 | 38,17 | 36,47 | 37,78 | 32,30 | 31,21 | 36,72 | 37,95 | 39,44 | 37,55 | 40,29 | 40,19 | 41,25 | 40,62 | 41,36 | 41,15 | 40,39 | 42,27 | 40,00 | 39,30 | 40,00 |
| 23 | TrSC58v2_6495_HEMAH_TrGP63 | 35,92 | 39,36 | 38,21 | 37,83 | 37,21 | 37,92 | 36,51 | 37,07 | 32,91 | 30,91 | 36,66 | 38,28 | 40,14 | 37,83 | 39,59 | 39,07 | 40,35 | 41,24 | 40,74 | 40,95 | 39,92 | 41,77 | 39,52 | 39,71 | 39,51 |
| 24 | TrSC58v2_8644_HEMAH_TrGP63 | 35,20 | 40,11 | 38,74 | 38,48 | 37,63 | 38,62 | 36,89 | 37,64 | 32,67 | 31,61 | 37,10 | 38,74 | 39,59 | 39,71 | 40,29 | 41,29 | 40,98 | 42,33 | 41,47 | 41,90 | 41,19 | 43,20 | 41,01 | 40,39 | 41,47 |
| 25 | TrSC58v2_9275_HEMAH_TrGP63 | 35,05 | 40,29 | 39,37 | 39,02 | 38,32 | 39,16 | 37,56 | 38,51 | 33,33 | 32,23 | 37,44 | 39,28 | 39,96 | 39,08 | 40,50 | 40,90 | 41,60 | 42,55 | 41,90 | 42,55 | 41,60 | 43,84 | 41,44 | 40,39 | 41,68 |
| 26 | TrSC58v2_8384_HELAH_TrGP63 | 38,05 | 38,79 | 39,15 | 39,48 | 39,70 | 39,70 | 39,70 | 38,76 | 38,39 | 37,12 | 39,06 | 39,83 | 38,82 | 40,72 | 42,92 | 39,03 | 44,32 | 43,58 | 44,04 | 43,58 | 43,54 | 43,12 | 43,05 | 42,43 | 42,20 |
| 27 | TrSC58v2_559_HELAH_TrGP63 | 37,86 | 38,38 | 39,12 | 39,43 | 39,63 | 39,63 | 39,63 | 37,87 | 38,41 | 36,88 | 39,15 | 39,06 | 38,74 | 39,96 | 42,44 | 38,87 | 43,11 | 43,82 | 44,14 | 43,93 | 43,78 | 43,40 | 43,12 | 43,10 | 42,77 |
| 28 | TrSC58v2_8165_HAIAH_TrGP63 | 39,66 | 37,80 | 40,34 | 40,59 | 40,59 | 40,59 | 40,59 | 38,66 | 38,67 | 35,35 | 41,18 | 41,44 | 41,18 | 38,66 | 42,92 | 44,35 | 42,83 | 42,71 | 42,41 | 43,04 | 44,20 | 43,33 | 42,04 | 43,45 | 41,67 |
| 29 | TrSC58v2_9363_HAIAH_TrGP63 | 39,56 | 37,58 | 38,45 | 39,54 | 39,54 | 39,54 | 39,54 | 36,63 | 39,54 | 36,41 | 39,16 | 39,96 | 40,56 | 38,27 | 42,26 | 40,42 | 41,80 | 39,92 | 39,63 | 39,83 | 40,53 | 39,29 | 38,89 | 40,25 | 38,25 |
| 30 | TrSC58v2_9356_HAIAH_TrGP63 | 39,43 | 37,64 | 39,41 | 39,54 | 39,54 | 39,54 | 39,54 | 36,92 | 36,47 | 32,83 | 38,86 | 39,31 | 37,90 | 38,59 | 44,31 | 41,54 | 42,15 | 41,79 | 41,36 | 41,79 | 41,20 | 40,94 | 40,08 | 42,43 | 40,30 |
| 31 | TrSC58v2_3271_HAIAH_TrGP63 | 34,14 | 36,46 | 38,96 | 37,73 | 36,43 | 37,73 | 35,37 | 37,21 | 32,60 | 32,42 | 34,18 | 37,14 | 37,38 | 38,56 | 43,71 | 39,53 | 42,36 | 41,58 | 41,15 | 41,58 | 41,02 | 40,72 | 40,29 | 42,22 | 40,09 |
| 32 | TrSC58v2_3280_HAIAH_TrGP63 | 39,35 | 37,38 | 38,99 | 39,08 | 39,08 | 39,08 | 39,08 | 36,63 | 36,02 | 32,55 | 38,02 | 38,86 | 37,64 | 38,59 | 43,78 | 41,63 | 42,24 | 41,79 | 41,36 | 41,79 | 41,29 | 41,15 | 40,08 | 42,64 | 40,51 |
| 33 | TrSC58v2_5263_HEIAH_TrGP63 | 32,67 | 34,33 | 33,48 | 32,22 | 32,22 | 33,46 | 32,22 | 38,44 | 30,02 | 30,20 | 33,39 | 34,17 | 34,42 | 38,90 | 36,55 | 35,62 | 39,15 | 41,26 | 38,93 | 40,09 | 39,73 | 39,39 | 38,50 | 40,33 | 40,09 |
| 34 | TrSC58v2_5266_HEIAH_TrGP63 | 32,79 | 34,52 | 33,70 | 32,34 | 32,34 | 33,66 | 32,34 | 38,44 | 30,15 | 30,33 | 33,52 | 34,37 | 34,62 | 38,90 | 36,78 | 35,76 | 38,93 | 41,26 | 38,93 | 40,09 | 39,73 | 39,39 | 38,50 | 40,33 | 40,09 |
| 35 | TrSC58v2_6663_HEIAH_TrGP63 | 29,89 | 34,36 | 33,27 | 32,88 | 32,36 | 33,57 | 31,87 | 36,96 | 28,39 | 29,92 | 31,93 | 32,83 | 33,63 | 37,30 | 37,04 | 34,84 | 37,92 | 39,14 | 36,81 | 37,63 | 38,05 | 37,91 | 36,35 | 38,24 | 38,52 |
| 36 | TrSC58v2_9131_HEIAH_TrGP63 | 28,45 | 33,15 | 33,40 | 32,03 | 31,46 | 33,74 | 30,56 | 38,51 | 28,20 | 29,96 | 30,14 | 32,83 | 33,57 | 36,76 | 36,42 | 34,12 | 38,70 | 38,52 | 36,81 | 38,04 | 38,25 | 38,11 | 36,95 | 38,45 | 38,52 |
| 37 | TrSC58v2_9025_HEIAH_TrGP63 | 29,98 | 34,59 | 34,55 | 34,10 | 33,58 | 34,92 | 33,33 | 37,75 | 32,84 | 31,89 | 31,87 | 33,78 | 33,66 | 38,14 | 38,64 | 36,20 | 40,43 | 38,89 | 37,99 | 39,63 | 38,40 | 38,89 | 36,90 | 38,60 | 38,48 |
| 38 | TrSC58v2_4259_HEIAH_TrGP63 | 28,38 | 33,87 | 33,79 | 31,43 | 30,61 | 33,16 | 30,18 | 37,75 | 28,62 | 28,93 | 30,87 | 32,77 | 32,91 | 38,51 | 38,48 | 36,15 | 38,65 | 38,92 | 37,40 | 39,05 | 37,83 | 38,10 | 36,31 | 38,02 | 38,30 |
| 39 | TrSC58v2_631_HEIGH_TrGP63 | 71,21 | 70,28 | 71,26 | 71,45 | 71,29 | 71,45 | 71,45 | 71,00 | 63,06 | 61,54 | 74,77 | 71,47 | 75,40 | 37,28 | 43,97 | 42,94 | 44,13 | 45,19 | 43,53 | 43,92 | 44,66 | 44,99 | 44,42 | 45,10 | 43,61 |
| 40 | TrSC58v2_632_HEIGH_TrGP63 | 71,36 | 70,28 | 71,26 | 71,45 | 71,29 | 71,45 | 71,45 | 71,00 | 63,06 | 61,54 | 74,92 | 71,63 | 75,24 | 37,28 | 43,97 | 42,94 | 44,13 | 45,19 | 43,53 | 43,92 | 44,66 | 44,99 | 44,42 | 45,10 | 43,61 |
| 41 | TrSC58v2_633_HEIGH_TrGP63 | 71,36 | 70,28 | 71,26 | 71,45 | 71,29 | 71,45 | 71,45 | 71,00 | 63,06 | 61,54 | 74,92 | 71,63 | 75,24 | 37,28 | 43,97 | 42,94 | 44,13 | 45,19 | 43,53 | 43,92 | 44,66 | 44,99 | 44,42 | 45,10 | 43,61 |
| 42 | TrSC58v2_4929_HEIGH_TrGP63 | 64,03 | 67,89 | 66,46 | 67,63 | 68,52 | 68,03 | 68,79 | 70,81 | 57,31 | 58,46 | 68,15 | 69,59 | 71,09 | 37,98 | 43,22 | 42,54 | 43,97 | 44,42 | 43,76 | 43,76 | 43,86 | 43,76 | 45,09 | 45,30 | 43,76 |
| 43 | TrSC58v2_6472_HEIGH_TrGP63 | 71,49 | 69,02 | 70,32 | 64,63 | 64,19 | 67,72 | 64,34 | 70,77 | 59,58 | 59,92 | 69,75 | 68,97 | 68,59 | 36,18 | 43,31 | 40,50 | 42,41 | 44,72 | 43,70 | 43,70 | 43,87 | 44,92 | 43,94 | 44,72 | 43,90 |
| 44 | TrSC58v2_7196_HEIGH_TrGP63 | 67,76 | 67,43 | 69,04 | 62,96 | 60,30 | 66,67 | 59,38 | 68,83 | 55,59 | 58,54 | 67,82 | 69,16 | 68,80 | 35,69 | 44,14 | 40,49 | 43,44 | 44,96 | 43,95 | 44,76 | 44,51 | 44,96 | 45,96 | 45,97 | 45,36 |
| 45 | TrSC58v2_9139_HEIGH_TrGP63 | 65,25 | 70,10 | 72,47 | 68,06 | 66,96 | 71,03 | 66,96 | 70,73 | 61,88 | 62,57 | 68,54 | 68,72 | 71,41 | 36,09 | 42,64 | 40,75 | 43,77 | 44,40 | 42,55 | 43,53 | 42,78 | 43,42 | 44,04 | 44,71 | 43,81 |
| 46 | TrSC58v2_9136_HEIGH_TrGP63 | 68,36 | 71,45 | 72,47 | 68,26 | 67,62 | 71,24 | 67,62 | 70,73 | 62,10 | 62,57 | 69,90 | 70,17 | 72,73 | 36,09 | 42,64 | 40,75 | 43,77 | 44,40 | 42,55 | 43,53 | 42,78 | 43,42 | 44,04 | 44,71 | 43,81 |
| 47 | TrSC58v2_9135_HEIGH_TrGP63 | 56,13 | 68,26 | 73,56 | 66,78 | 65,30 | 69,20 | 65,30 | 70,73 | 69,50 | 71,37 | 62,73 | 66,89 | 70,07 | 35,39 | 41,60 | 40,43 | 43,73 | 43,50 | 41,65 | 42,63 | 42,53 | 42,72 | 43,41 | 44,20 | 43,11 |
| 48 | TrSC58v2_9137_HEIGH_TrGP63 | 70,52 | 70,35 | 74,56 | 74,78 | 74,78 | 74,78 | 74,78 | 71,35 | 74,78 | 70,68 | 71,96 | 72,37 | 72,58 | 35,57 | 43,14 | 43,08 | 44,76 | 44,08 | 42,67 | 43,76 | 44,08 | 43,64 | 44,74 | 44,64 | 43,86 |
| 49 | TrSC58v2_9032_HEIGH_TrGP63 | 69,31 | 71,16 | 68,80 | 72,23 | 72,42 | 72,23 | 72,23 | 66,76 | 67,74 | 64,11 | 73,54 | 69,08 | 69,66 | 36,26 | 41,15 | 42,20 | 41,59 | 41,76 | 41,29 | 41,88 | 41,90 | 42,55 | 41,27 | 41,88 | 41,76 |
| 50 | TrSC58v2_5291_HEIGH_TrGP63 | 70,40 | 69,02 | 71,76 | 66,57 | 65,97 | 69,54 | 66,12 | 70,54 | 60,86 | 61,20 | 72,55 | 73,68 | 74,40 | 35,77 | 43,95 | 41,77 | 41,83 | 45,28 | 44,60 | 45,38 | 45,21 | 45,47 | 44,32 | 45,78 | 45,47 |
| 51 | TrSC58v2_7524_HEIAH_TrGP63 | 64,98 | 67,45 | 72,78 | 64,64 | 61,76 | 70,57 | 60,00 | 76,76 | 56,19 | 61,21 | 71,98 | 72,46 | 73,95 | 37,01 | 42,71 | 37,81 | 41,28 | 44,56 | 44,56 | 43,93 | 43,27 | 45,61 | 44,79 | 46,03 | 44,35 |
| 52 | TrSC58v2_7598_QKIGH_TrGP63 | 100,00 | 72,36 | 73,33 | 67,51 | 65,70 | 72,68 | 63,74 | 75,68 | 56,22 | 61,14 | 72,52 | 77,37 | 76,71 | 37,28 | 44,83 | 40,74 | 43,07 | 45,38 | 44,31 | 44,31 | 44,36 | 44,60 | 44,62 | 44,90 | 44,60 |
| 53 | TrSC58v2_9319_HEIGH_TrGP63 | 72,36 | 100,00 | 76,05 | 76,97 | 76,81 | 76,97 | 76,97 | 76,22 | 68,16 | 65,20 | 75,77 | 72,96 | 75,08 | 37,22 | 44,25 | 41,79 | 43,36 | 43,26 | 42,86 | 42,25 | 43,84 | 43,26 | 43,70 | 44,87 | 44,47 |
| 54 | TrSC58v2_4655_HEIGH_TrGP63 | 73,33 | 76,05 | 100,00 | 91,48 | 91,30 | 92,66 | 91,48 | 96,76 | 93,15 | 92,34 | 81,67 | 79,70 | 82,36 | 37,40 | 43,98 | 41,91 | 42,32 | 43,75 | 42,74 | 42,94 | 44,71 | 44,76 | 45,11 | 45,56 | 44,96 |
| 55 | TrSC58v2_4642_HEIGH_TrGP63 | 67,51 | 76,97 | 91,48 | 100,00 | 99,58 | 99,84 | 99,30 | 97,03 | 74,60 | 77,54 | 73,98 | 77,52 | 82,65 | 38,23 | 45,03 | 41,68 | 43,25 | 45,27 | 44,27 | 44,47 | 45,29 | 45,88 | 46,06 | 46,48 | 46,08 |
| 56 | TrSC58v2_4641_HEIGH_TrGP63 | 65,70 | 76,81 | 91,30 | 99,58 | 100,00 | 99,84 | 99,35 | 96,76 | 71,41 | 77,54 | 71,43 | 77,36 | 82,48 | 38,03 | 45,22 | 41,61 | 43,05 | 45,07 | 44,06 | 44,27 | 45,10 | 45,67 | 45,87 | 46,28 | 45,88 |
| 57 | TrSC58v2_4649_HEIGH_TrGP63 | 72,68 | 76,97 | 92,66 | 99,84 | 99,84 | 100,00 | 100,00 | 97,03 | 80,67 | 81,97 | 80,38 | 77,64 | 82,65 | 38,23 | 45,03 | 41,94 | 43,25 | 45,27 | 44,27 | 44,47 | 45,29 | 45,88 | 46,06 | 46,48 | 46,08 |
| 58 | TrSC58v2_4654_HEIGH_TrGP63 | 63,74 | 76,97 | 91,48 | 99,30 | 99,35 | 100,00 | 100,00 | 97,03 | 69,58 | 77,72 | 69,36 | 77,52 | 82,65 | 38,23 | 45,03 | 41,61 | 43,25 | 45,27 | 44,27 | 44,47 | 45,29 | 45,88 | 46,06 | 46,48 | 46,08 |
| 59 | TrSC58v2_4653_HEIGH_TrGP63 | 75,68 | 76,22 | 96,76 | 97,03 | 96,76 | 97,03 | 97,03 | 100,00 | 97,03 | 96,76 | 85,14 | 80,54 | 82,16 | 39,51 | 41,89 | 42,62 | 44,14 | 44,57 | 44,84 | 44,57 | 46,20 | 45,92 | 46,47 | 46,20 | 46,20 |
| 60 | TrSC58v2_4652_HEIGH_TrGP63 | 56,22 | 68,16 | 93,15 | 74,60 | 71,41 | 80,67 | 69,58 | 97,03 | 100,00 | 99,83 | 64,97 | 69,73 | 74,44 | 37,42 | 43,66 | 39,53 | 42,19 | 44,06 | 43,06 | 43,26 | 44,23 | 44,87 | 44,88 | 45,67 | 45,07 |
| 61 | TrSC58v2_4656_HEIGH_TrGP63 | 61,14 | 65,20 | 92,34 | 77,54 | 77,54 | 81,97 | 77,72 | 96,76 | 99,83 | 100,00 | 69,88 | 69,13 | 72,17 | 38,41 | 40,32 | 38,12 | 41,87 | 43,55 | 42,86 | 42,86 | 43,75 | 44,24 | 44,72 | 45,85 | 44,70 |
| 62 | TrSC58v2_9446_HETGH_TrGP63 | 72,52 | 75,77 | 81,67 | 73,98 | 71,43 | 80,38 | 69,36 | 85,14 | 64,97 | 69,88 | 100,00 | 84,32 | 85,49 | 37,87 | 44,05 | 40,81 | 42,50 | 45,58 | 43,92 | 45,10 | 46,27 | 45,97 | 46,54 | 46,67 | 45,58 |
| 63 | TrSC58v2_4795_HEIAH_TrGP63 | 77,37 | 72,96 | 79,70 | 77,52 | 77,36 | 77,64 | 77,52 | 80,54 | 69,73 | 69,13 | 84,32 | 100,00 | 87,06 | 38,14 | 44,14 | 42,83 | 44,30 | 43,61 | 42,75 | 43,53 | 44,25 | 44,01 | 44,89 | 45,49 | 44,01 |
| 64 | TrSC58v2_7306_HEIAH_TrGP63 | 76,71 | 75,08 | 82,36 | 82,65 | 82,48 | 82,65 | 82,65 | 82,16 | 74,44 | 72,17 | 85,49 | 87,06 | 100,00 | 37,28 | 43,86 | 44,27 | 44,02 | 46,37 | 44,51 | 45,29 | 46,46 | 46,56 | 46,15 | 47,45 | 45,38 |
| 65 | TrSC58v2_4835_HEMAH_TrGP63 | 37,28 | 37,22 | 37,40 | 38,23 | 38,03 | 38,23 | 38,23 | 39,51 | 37,42 | 38,41 | 37,87 | 38,14 | 37,28 | 100,00 | 40,58 | 41,08 | 45,03 | 43,81 | 44,94 | 44,33 | 44,97 | 46,45 | 43,51 | 43,72 | 44,02 |
| 66 | TrSC58v2_3991_HEIVH_TrGP63 | 44,83 | 44,25 | 43,98 | 45,03 | 45,22 | 45,03 | 45,03 | 41,89 | 43,66 | 40,32 | 44,05 | 44,14 | 43,86 | 40,58 | 100,00 | 43,38 | 45,23 | 45,78 | 46,39 | 47,19 | 45,82 | 49,00 | 48,02 | 49,00 | 46,99 |
| 67 | TrSC58v2_4583_HEIAH_TrGP63 | 40,74 | 41,79 | 41,91 | 41,68 | 41,61 | 41,94 | 41,61 | 42,62 | 39,53 | 38,12 | 40,81 | 42,83 | 44,27 | 41,08 | 43,38 | 100,00 | 45,79 | 49,11 | 48,62 | 48,02 | 46,50 | 48,12 | 46,12 | 47,43 | 48,71 |
| 68 | TrSC58v2_7126_HEIAH_TrGP63 | 43,07 | 43,36 | 42,32 | 43,25 | 43,05 | 43,25 | 43,25 | 44,14 | 42,19 | 41,87 | 42,50 | 44,30 | 44,02 | 45,03 | 45,23 | 45,79 | 100,00 | 49,61 | 50,29 | 50,88 | 47,34 | 49,22 | 49,33 | 50,29 | 49,02 |
| 69 | TrSC58v2_8509_HEIAH_TrGP63 | 45,38 | 43,26 | 43,75 | 45,27 | 45,07 | 45,27 | 45,27 | 44,57 | 44,06 | 43,55 | 45,58 | 43,61 | 46,37 | 43,81 | 45,78 | 49,11 | 49,61 | 100,00 | 88,52 | 88,72 | 69,79 | 71,54 | 69,79 | 69,84 | 71,73 |
| 70 | TrSC58v2_2748_HEIAH_TrGP63 | 44,31 | 42,86 | 42,74 | 44,27 | 44,06 | 44,27 | 44,27 | 44,84 | 43,06 | 42,86 | 43,92 | 42,75 | 44,51 | 44,94 | 46,39 | 48,62 | 50,29 | 88,52 | 100,00 | 92,43 | 72,18 | 73,15 | 71,01 | 73,59 | 72,57 |
| 71 | TrSC58v2_6871_HEIAH_TrGP63 | 44,31 | 42,25 | 42,94 | 44,47 | 44,27 | 44,47 | 44,47 | 44,57 | 43,26 | 42,86 | 45,10 | 43,53 | 45,29 | 44,33 | 47,19 | 48,02 | 50,88 | 88,72 | 92,43 | 100,00 | 71,01 | 72,18 | 71,60 | 72,82 | 71,98 |
| 72 | TrSC58v2_2586_HEIAH_TrGP63 | 44,36 | 43,84 | 44,71 | 45,29 | 45,10 | 45,29 | 45,29 | 46,20 | 44,23 | 43,75 | 46,27 | 44,25 | 46,46 | 44,97 | 45,82 | 46,50 | 47,34 | 69,79 | 72,18 | 71,01 | 100,00 | 74,12 | 71,62 | 72,96 | 73,74 |
| 73 | TrSC58v2_7404_HGIAH_TrGP63 | 44,60 | 43,26 | 44,76 | 45,88 | 45,67 | 45,88 | 45,88 | 45,92 | 44,87 | 44,24 | 45,97 | 44,01 | 46,56 | 46,45 | 49,00 | 48,12 | 49,22 | 71,54 | 73,15 | 72,18 | 74,12 | 100,00 | 77,63 | 78,02 | 77,43 |
| 74 | TrSC58v2_8159_HEIAH_TrGP63 | 44,62 | 43,70 | 45,11 | 46,06 | 45,87 | 46,06 | 46,06 | 46,47 | 44,88 | 44,72 | 46,54 | 44,89 | 46,15 | 43,51 | 48,02 | 46,12 | 49,33 | 69,79 | 71,01 | 71,60 | 71,62 | 77,63 | 100,00 | 80,54 | 81,13 |
| 75 | TrSC58v2_4723_HEIAH_TrGP63 | 44,90 | 44,87 | 45,56 | 46,48 | 46,28 | 46,48 | 46,48 | 46,20 | 45,67 | 45,85 | 46,67 | 45,49 | 47,45 | 43,72 | 49,00 | 47,43 | 50,29 | 69,84 | 73,59 | 72,82 | 72,96 | 78,02 | 80,54 | 100,00 | 83,07 |
| 76 | TrSC58v2_6284_HEIAH_TrGP63 | 44,60 | 44,47 | 44,96 | 46,08 | 45,88 | 46,08 | 46,08 | 46,20 | 45,07 | 44,70 | 45,58 | 44,01 | 45,38 | 44,02 | 46,99 | 48,71 | 49,02 | 71,73 | 72,57 | 71,98 | 73,74 | 77,43 | 81,13 | 83,07 | 100,00 |
