## Supplementary table S5 for "Comprehensive genomic analysis of *Trypanosoma rangeli* reveals key insights into the biology and evolution of this non-virulent American mammalian trypanosome"

| Models | Group | pTM score | Presence of ion zinc binding motif (HEXXHXXGX...H) | Type | Presence of S1' pocket | Global RMSD (Å) | Helix B RMSD (Å) | P2Rank score | Ligand cavity probability | Presence of GXXAWA motif | Type | Docking Affinity (kcal/mol) | Ligand RMSD (Å) compared to control | Rank | Ligand oriented toward the Zn ion | Percentage of complexes with the ligand oriented toward the Zn ion |
|---|---|---|---|---|---|---|---|---|---|---|---|---|---|---|---|---|
| Control | | - | Yes | HEXXHXXG(61)H | Yes | - | - | 15,78 | 0,76 | Yes | GXXAWA | -5,54 | 0 | 2 | Yes | 34% |
| TrSC58v2_2748 | 6 | 0,9 | Yes | HEXXHXXG(60)H | No | 23,29 | 12,7 | 14,91 | 0,74 | Yes | SXXAWQ | -6,65 | 4,54 | 4 | Yes | 67% |
| TrSC58v2_4795 | 4 | 0,7 | Yes | HEXXHXXG(61)H | No | 24,79 | 11,27 | 14,02 | 0,71 | Yes | DXXAWA | -5,62 | 4,29 | 4 | Yes | 44% |
| TrSC58v2_9135 | 4 | 0,6 | Yes | HEXXHXXG(58)H | Yes | 13,24 | 7,71 | 11,87 | 0,63 | Yes | GXXAWA | -5,78 | 2,97 | 5 | Yes | 56% |
| TrSC58v2_9139 | 4 | 0,57 | Yes | HEXXHXXG(61)H | No | 23,38 | 13,53 | 11,15 | 0,6 | Yes | GXXAWA | -6,04 | 3,58 | 5 | Yes | 44% |
| TrSC58v2_2586 | 6 | 0,83 | Yes | HEXXHXXG(60)H | No | 13,6 | 2,16 | 10,7 | 0,57 | Yes | SXXAWA | -5,56 | 2,41 | 5 | Yes | 44% |
| TrSC58v2_7126 | 8 | 0,81 | Yes | HEXXHXXG(60)H | No | 15,26 | 2,25 | 10,22 | 0,55 | Yes | GXXAWA | -5,86 | 2,04 | 5 | Yes | 44% |
| TrSC58v2_1328 | 12 | 0,85 | Yes | HEXXHXXG(59)H | Yes | 15,37 | 3,24 | 9,56 | 0,51 | Yes | AXXAWA | -6,07 | 2,16 | 4 | No | 11% |
| *T. cruzi* | 1 | 0,73 | Yes | HEXXHXXG(70)H | No | 21,76 | 7,34 | 8,84 | 0,47 | Yes | GXXAWA | -6,03 | 1,94 | 3 | Yes | 56% |
| TrSC58v2_3991 | 7 | 0,85 | Yes | HEXXHXXG(60)H | No | 14,9 | 3,8 | 8,46 | 0,45 | Yes | GXXAWA | -4,98 | 3 | 6 | No | 0% |
| TrSC58v2_6130 | 11 | 0,67 | Yes | HEXXHXXG(60)H | No | 9,11 | 1,14 | 8,23 | 0,44 | Yes | NXXAWA | -6,31 | 4,13 | 2 | No | 0% |
| TrSC58v2_4835 | 3 | 0,72 | Yes | HEXXHXXG(233)H | No | 9,6 | 4,16 | 8,05 | 0,43 | Yes | MXXAWA | -5,83 | 2,89 | 4 | No | 0% |
| TrSC58v2_9356 | 10 | 0,82 | No | HAXXHXXG(57)L | No | 15,57 | 1,77 | 7,93 | 0,42 | No | DXXTWA | -4,47 | 4,81 | 9 | No | 0% |
| TrSC58v2_4583 | 5 | 0,71 | Yes | HEXXHXXG(60)H | No | 15,88 | 2,5 | 7,72 | 0,41 | No | AXXAWG | -5,08 | 2,1 | 7 | No | 0% |
| TrSC58v2_559 | 2 | 0,76 | Yes | HEXXHXXG(61)H | No | 15,84 | 2,48 | 6,95 | 0,36 | No | XXXIFA | -5,58 | 3,44 | 2 | No | 0% |
| TrSC58v2_560 | 11 | 0,67 | No | RGXXYXXG(60)H | No | 7,82 | 11,97 | 4,79 | 0,21 | No | AXXVWA | -5,72 | 4,13 | 5 | No | 0% |
| TrSC58v2_9025 | 9 | 0,75 | Yes | HEXXHXXG(61)H | No | 16,81 | 2,22 | 3,44 | 0,12 | No | XXXVWA | -4,2 | 4,67 | 8 | Yes | 22% |
