## Supplementary table S6 for "Comprehensive genomic analysis of *Trypanosoma rangeli* reveals key insights into the biology and evolution of this non-virulent American mammalian trypanosome"

| | SC58 V.2 genome position | | | | | | | Signal Peptide | | | GPI Anchor | | | Core Protein Characteristics | | O-glycosylation | | N-glycosylation | | Phosphorylation | | Thr | | Ala | | Ser | | Val | | Aromatic | |
|---|---|---|---|---|---|---|---|---|---|---|---|---|---|---|---|---|---|---|---|---|---|---|---|---|---|---|---|---|---|---|---|
| Glycoprotein | Annotated | Scaffold | Status | Start | End | Strand | Full length (aa) | Prediction Likelihood | cleavage site Position | cleavage site Probability | GPI-Anchored | ω-site Position | Likelihood | Core length (aa) | Core Molecular Weight (kDa) | residues | % | residues | % | residues | % | residues | % | residues | % | residues | % | residues | % | residues | % |
| TrMUCg | TrSC58v2_9456 TrMUCg1 | TRSC58v2_s001 | gene, putative | 14.388 | 14.810 | plus | 140 | 0,99941 | 26-27 | 0,809 | YES | 115 | 0,296 | 89 | 8,70 | 60 | 67,42 | 0 | 0,00 | 47 | 52,81 | 49 | 55,06 | 10 | 11,24 | 13 | 14,61 | 2 | 2,25 | 0 | 0,00 |
| TrMUCg | TrSC58v2_9457 TrMUCg2 | TRSC58v2_s001 | gene, putative | 11.778 | 12.200 | plus | 140 | 0,99941 | 26-27 | 0,809 | YES | 115 | 0,296 | 89 | 8,70 | 60 | 67,42 | 0 | 0,00 | 45 | 50,56 | 49 | 55,06 | 10 | 11,24 | 13 | 14,61 | 2 | 2,25 | 0 | 0,00 |
| TrMUCg | TrSC58v2_9455 TrMUCg3 | TRSC58v2_s001 | gene, putative | 9.166 | 9.591 | plus | 141 | 0,99927 | 26-27 | 0,873 | YES | 116 | 0,296 | 90 | 8,87 | 60 | 66,67 | 0 | 0,00 | 44 | 48,89 | 51 | 56,67 | 9 | 10,00 | 12 | 13,33 | 2 | 2,22 | 0 | 0,00 |
| TrMUCg | TrSC58v2_9459 TrMUCg4 | TRSC58v2_s044 | gene, putative | 15.572 | 15.859 | plus | 95 | 0,99979 | 26-27 | 0,981 | YES | 70 | 0,304 | 44 | 4,00 | 18 | 40,91 | 0 | 0,00 | 16 | 36,36 | 14 | 31,82 | 12 | 27,27 | 5 | 11,36 | 1 | 2,27 | 0 | 0,00 |
| TrMUCg | TrSC58v2_9460 TrMUCg5 | TRSC58v2_s044 | gene, putative | 13.154 | 13.441 | plus | 95 | 0,99979 | 26-27 | 0,981 | YES | 70 | 0,304 | 44 | 4,00 | 18 | 40,91 | 0 | 0,00 | 16 | 36,36 | 14 | 31,82 | 12 | 27,27 | 5 | 11,36 | 1 | 2,27 | 0 | 0,00 |
| TrMUCg | TrSC58v2_9461 TrMUCg6 | TRSC58v2_s044 | gene, putative | 10.741 | 11.028 | plus | 95 | 0,99978 | 26-27 | 0,978 | YES | 70 | 0,307 | 44 | 4,03 | 19 | 43,18 | 0 | 0,00 | 17 | 38,64 | 15 | 34,09 | 11 | 25,00 | 5 | 11,36 | 1 | 2,27 | 0 | 0,00 |
| TrMUCg | TrSC58v2_9458 TrMUCg7 | TRSC58v2_s025 | gene, putative | 311.604 | 311.906 | plus | 100 | 0,99980 | 26-27 | 0,982 | YES | 75 | 0,304 | 49 | 4,44 | 21 | 42,86 | 0 | 0,00 | 19 | 38,78 | 17 | 34,69 | 14 | 28,57 | 5 | 10,20 | 1 | 2,04 | 0 | 0,00 |
| TrTASV | TrSC58v2_9464 TrTASV-like2 | TRSC58v2_s020 | pseudogene | 97.576 | 97.809 | minus | 77 | 0,99916 | 31-32 | 0,970 | NO | — | 0,994 | 46 | 5,02 | 0 | 0,00 | 0 | 0,00 | 6 | 13,04 | 3 | 6,52 | 6 | 13,04 | 7 | 15,22 | 4 | 8,70 | 3 | 6,52 |
| TrTASV | TrSC58v2_9465 TrTASV-like6 | TRSC58v2_s020 | pseudogene | 90.219 | 90.452 | minus | 77 | 0,99981 | 32-33 | 0,976 | NO | — | 0,995 | 45 | 4,98 | 1 | 2,22 | 0 | 0,00 | 6 | 13,33 | 2 | 4,44 | 5 | 11,11 | 7 | 15,56 | 4 | 8,89 | 3 | 6,67 |
| TrTASV | TrSC58v2_6391 TrTASV-like16 | TRSC58v2_s024 | pseudogene | 82.066 | 82.413 | plus | 115 | 0,99978 | 30-31 | 0,777 | NO | — | 0,901 | 85 | 9,35 | 1 | 1,18 | 2 | 2,35 | 12 | 14,12 | 6 | 7,06 | 5 | 5,88 | 9 | 10,59 | 12 | 14,12 | 9 | 10,59 |
| TrTASV | TrSC58v2_5804 TrTASV-like1 | TRSC58v2_s020 | gene, putative | 81.966 | 82.949 | minus | 327 | 0,99984 | 31-32 | 0,982 | YES | 300 | 0,338 | 269 | 26,63 | 22 | 8,18 | 0 | 0,00 | 40 | 14,87 | 23 | 8,55 | 51 | 18,96 | 33 | 12,27 | 17 | 6,32 | 18 | 6,69 |
| TrTASV | TrSC58v2_8519 TrTASV-like3 | TRSC58v2_s047 | gene, putative | 94.366 | 95.511 | plus | 381 | 0,99984 | 29-30 | 0,983 | NO | — | 0,993 | 352 | 39,07 | 20 | 5,68 | 3 | 0,85 | 41 | 11,65 | 32 | 9,09 | 25 | 7,10 | 27 | 7,67 | 19 | 5,40 | 28 | 7,96 |
| TrTASV | TrSC58v2_8515 TrTASV-like4 | TRSC58v2_s047 | gene, putative | 84.339 | 85.844 | minus | 501 | 0,99984 | 29-30 | 0,983 | YES | 473 | 0,351 | 444 | 45,54 | 39 | 8,78 | 8 | 1,80 | 60 | 13,51 | 48 | 10,81 | 55 | 12,39 | 45 | 10,14 | 17 | 3,83 | 30 | 6,76 |
| TrTASV | TrSC58v2_9169 TrTASV-like5 | TRSC58v2_s061 | gene, putative | 73.652 | 75.154 | plus | 500 | 0,99993 | 22-23 | 0,668 | YES | 472 | 0,313 | 450 | 46,34 | 41 | 9,11 | 5 | 1,11 | 68 | 15,11 | 46 | 10,22 | 52 | 11,56 | 49 | 10,89 | 20 | 4,44 | 32 | 7,11 |
| TrTASV | TrSC58v2_9181 TrTASV-like7 | TRSC58v2_s062 | gene, putative | 8.958 | 10.436 | minus | 492 | 0,99987 | 29-30 | 0,984 | YES | 465 | 0,275 | 436 | 44,2 | 39 | 8,94 | 7 | 1,61 | 64 | 14,68 | 45 | 10,32 | 53 | 12,16 | 51 | 11,70 | 17 | 3,90 | 26 | 5,96 |
| TrTASV | TrSC58v2_8529 TrTASV-like8 | TRSC58v2_s047 | gene, putative | 117.637 | 1.186 | plus | 325 | 0,99990 | 30-31 | 0,488 | YES | 298 | 0,31 | 268 | 26,82 | 24 | 8,96 | 0 | 0,00 | 49 | 18,28 | 26 | 9,70 | 41 | 15,30 | 34 | 12,69 | 18 | 6,72 | 16 | 5,97 |
| TrTASV | TrSC58v2_8521 TrTASV-like9 | TRSC58v2_s047 | gene, putative | 98.395 | 99.375 | plus | 326 | 0,99988 | 30-31 | 0,673 | YES | 299 | 0,355 | 269 | 26,94 | 22 | 8,18 | 0 | 0,00 | 49 | 18,22 | 26 | 9,67 | 41 | 15,24 | 35 | 13,01 | 16 | 5,95 | 16 | 5,95 |
| TrTASV | TrSC58v2_8538 TrTASV-like10 | TRSC58v2_s047 | gene, putative | 146.774 | 147.685 | plus | 303 | 0,99985 | 26-27 | 0,978 | YES | 271 | 0,344 | 245 | 25,06 | 28 | 11,43 | 2 | 0,82 | 37 | 15,10 | 24 | 9,80 | 24 | 9,80 | 33 | 13,47 | 14 | 5,71 | 14 | 5,71 |
| TrTASV | TrSC58v2_9164 TrTASV-like11 | TRSC58v2_s061 | gene, putative | 63.120 | 64.295 | plus | 391 | 0,99999 | 22-23 | 0,680 | YES | 363 | 0,361 | 341 | 34,77 | 38 | 11,14 | 6 | 1,76 | 48 | 14,08 | 28 | 8,21 | 43 | 12,61 | 42 | 12,32 | 18 | 5,28 | 20 | 5,87 |
| TrTASV | TrSC58v2_9162 TrTASV-like12 | TRSC58v2_s061 | gene, putative | 59.510 | 60.685 | plus | 391 | 0,99999 | 22-23 | 0,680 | YES | 363 | 0,351 | 341 | 34,72 | 37 | 10,85 | 6 | 1,76 | 49 | 14,37 | 27 | 7,92 | 44 | 12,90 | 43 | 12,61 | 18 | 5,28 | 20 | 5,87 |
| TrTASV | TrSC58v2_9462 TrTASV-like13 | TRSC58v2_s047 | gene, putative | 87.735 | 88.715 | minus | 326 | 0,99984 | 32-33 | 0,797 | YES | 296 | 0,36 | 264 | 25,01 | 26 | 9,85 | 0 | 0,00 | 35 | 13,26 | 19 | 7,20 | 45 | 17,05 | 38 | 14,39 | 13 | 4,92 | 10 | 3,79 |
| TrTASV | TrSC58v2_9463 TrTASV-like14 | TRSC58v2_s061 | gene, putative | 84.448 | 86.040 | plus | 530 | 0,99983 | 34-35 | 0,471 | NO | — | 0,984 | 496 | 52,99 | 35 | 7,06 | 2 | 0,40 | 47 | 9,48 | 31 | 6,25 | 53 | 10,69 | 39 | 7,86 | 31 | 6,25 | 60 | 12,10 |
| TrTASV | TrSC58v2_8693 TrTASV-like15 | TRSC58v2_s050 | gene, putative | 110.729 | 111.592 | plus | 287 | 0,99986 | 29-30 | 0,980 | NO | — | 0,961 | 258 | 27,46 | 7 | 2,71 | 2 | 0,78 | 25 | 9,69 | 15 | 5,81 | 33 | 12,79 | 19 | 7,36 | 15 | 5,81 | 17 | 6,59 |
| TrTASV | TrSC58v2_6411 TrTASV-like17 | TRSC58v2_s024 | gene, putative | 132.715 | 133.746 | plus | 343 | 0,99986 | 30-31 | 0,805 | YES | 315 | 0,35 | 285 | 28,87 | 25 | 8,77 | 2 | 0,70 | 50 | 17,54 | 31 | 10,88 | 43 | 15,09 | 31 | 10,88 | 25 | 8,77 | 20 | 7,02 |

The selected glycoproteins for Biological assessment in the Choachí strain are highlighted in yellow.
