## Supplementary table S7 for "Comprehensive genomic analysis of *Trypanosoma rangeli* reveals key insights into the biology and evolution of this non-virulent American mammalian trypanosome"

| Strain | Original host | Geographical origin | KP1* | Lineage** |
|---|---|---|---|---|
| Choachí | *Rhodnius prolixus* | Colombia | + | A |
| SC58 | *Phyllomys dasythrix* | Brazil | - | D |
| PIT10 | *Panstrongylus megistus* | Brazil | - | ND |
| R1625 | *Homo sapiens* | El Salvador | + | C |
