## Supplementary table S8 for "Comprehensive genomic analysis of *Trypanosoma rangeli* reveals key insights into the biology and evolution of this non-virulent American mammalian trypanosome"

| Primers | | | | |
|---|---|---|---|---|
| Used in | Name | Sequence (5'- 3') | Restriction site | Amplicon (bp) |
| Amplification and cloning | *Tc*MUC I - without/GPI F | ***TCTAGAT***GATGATGACTTGCCGTCTGCTG | *Xba* I | 314 |
| | *Tc*MUC I - without/GPI R | ***GGTACC***TTTGCGAAGAAGTGGCGGTG | *Kpn* I | |
| | *Tc*MUCII without/GPI F | ***TCTAGA***TGATGATGACATGCCGTCTGCTG | *Xba* I | 596 |
| | *Tc*MUCII without/GPI R | ***AAGCTT***CATTTCGCGAAGACGGGACG | *Hind* III | |
| | *TrTASV-like1* - without/GPI F | ***TCTAGA***TGGCGATGGCGATGGCGATG | *Xba* I | 897 |
| | *TrTASV-like1* - without/GPI R | ***TCTAGA***CGCCTTGCTTTTTGTGTGTTTC | *Xba* I | |
| | *Tr*MUCg3 - without/GPI F | ***TCTAGA***TGAAGCTGAGGAGGTGCC | *Xba* I | 353 |
| | *Tr*MUCg3 - without/GPI R | ***GGTACC***CTTCCTCTCAGCCCTCTTCG | *Kpn* I | |
| Vector checking | 3UTRgp82-F | GAAATTGCCGTGAGGACTTC | | used in several combinations |
| | 3UTRgp82-R | GCTCCGCCATCGTCGTGG | | |
| | NeoR F | GATGGATTGCACGCAGGTTC | | |
| | NeoR R | TCAGAAGAACTCGTCAAGAAG | | |
| | HX1 F | GGCGGTTAACGAGTTTCTTC | | |
| | HX1 R | AAGACATCATAAAAGAAGTTG | | |
| | M13 R | TCACACAGGAAACAGCTATGAC | | |
| | mNG R | TATCCTCCTCGCCCTTGCTC | | |
| | 3utr gapdh | AGGTTGTTCTGCAGCGTTG | | |
| Genome integration assessment | HX1 R | AAGACATCATAAAAGAAGTTG | | 1.700 |
| | TubulinF | TCAAGAGCAGTAAGAACAGATG | | |
